# Complete genomic and epigenetic maps of human centromeres

**DOI:** 10.1101/2021.07.12.452052

**Authors:** Nicolas Altemose, Glennis A. Logsdon, Andrey V. Bzikadze, Pragya Sidhwani, Sasha A. Langley, Gina V. Caldas, Savannah J. Hoyt, Lev Uralsky, Fedor D. Ryabov, Colin J. Shew, Michael E.G. Sauria, Matthew Borchers, Ariel Gershman, Alla Mikheenko, Valery A. Shepelev, Tatiana Dvorkina, Olga Kunyavskaya, Mitchell R. Vollger, Arang Rhie, Ann M. McCartney, Mobin Asri, Ryan Lorig-Roach, Kishwar Shafin, Sergey Aganezov, Daniel Olson, Leonardo Gomes de Lima, Tamara Potapova, Gabrielle A. Hartley, Marina Haukness, Peter Kerpedjiev, Fedor Gusev, Kristof Tigyi, Shelise Brooks, Alice Young, Sergey Nurk, Sergey Koren, Sofie R. Salama, Benedict Paten, Evgeny I. Rogaev, Aaron Streets, Gary H. Karpen, Abby F. Dernburg, Beth A. Sullivan, Aaron F. Straight, Travis J. Wheeler, Jennifer L. Gerton, Evan E. Eichler, Adam M. Phillippy, Winston Timp, Megan Y. Dennis, Rachel J. O’Neill, Justin M. Zook, Michael C. Schatz, Pavel A. Pevzner, Mark Diekhans, Charles H. Langley, Ivan A. Alexandrov, Karen H. Miga

**Affiliations:** Department of Molecular and Cell Biology, University of California, Berkeley, Berkeley, CA, USA; Department of Genome Sciences, University of Washington School of Medicine, Seattle, WA, USA; Graduate Program in Bioinformatics and Systems Biology, University of California San Diego, La Jolla, CA, USA; Department of Biochemistry, Stanford University, Stanford, California, USA; Institute for Systems Genomics, University of Connecticut, Storrs, CT, USA; Department of Molecular and Cell Biology, University of Connecticut, Storrs, CT, USA; Sirius University of Science and Technology, Sochi, Russia; Vavilov Institute of General Genetics, Moscow, Russia; Moscow Polytechnic University, Moscow, Russia; Genome Center, MIND Institute, and Department of Biochemistry and Molecular Medicine, School of Medicine, University of California, Davis, Davis, CA, USA; Department of Biology, Johns Hopkins University, Baltimore, MD, USA; Stowers Institute for Medical Research, Kansas City, MO, USA; Department of Molecular Biology and Genetics, Johns Hopkins University, Baltimore, MD, USA; Center for Algorithmic Biotechnology, Institute of Translational Biomedicine, Saint Petersburg State University, Saint Petersburg, Russia; Genome Informatics Section, Computational and Statistical Genomics Branch, National Human Genome Research Institute, National Institutes of Health, Bethesda, MD, USA; UC Santa Cruz Genomics Institute, University of California Santa Cruz, Santa Cruz, CA, USA; Department of Computer Science, Johns Hopkins University, Baltimore, MD, USA; Department of Computer Science, University of Montana, Missoula, MT; Reservoir Genomics LLC, Oakland, CA; Howard Hughes Medical Institute, Chevy Chase, MD, USA; NIH Intramural Sequencing Center, National Human Genome Research Institute, National Institutes of Health, Bethesda, MD, USA; Department of Psychiatry, University of Massachusetts Medical School, Worcester, MA, USA; Faculty of Biology, Lomonosov Moscow State University, Moscow, Russia; Department of Bioengineering, University of California, Berkeley, Berkeley, CA, USA; Chan Zuckerberg Biohub, San Francisco, CA, USA; BioEngineering & BioMedical Sciences Department, Lawrence Berkeley National Laboratory, Berkeley, CA, USA; Institute for Quantitative Biosciences (QB3), University of California, Berkeley, Berkeley, CA, US; Department of Molecular Genetics and Microbiology, Duke University School of Medicine, Durham, NC, USA; Department of Biomedical Engineering, Johns Hopkins University, Baltimore, MD, USA; Biosystems and Biomaterials Division, National Institute of Standards and Technology, Gaithersburg, MD; Department of Computer Science and Engineering, University of California at San Diego, San Diego, CA, USA; Department of Evolution and Ecology, University of California Davis, Davis, CA, USA; Research Center of Biotechnology of the Russian Academy of Sciences, Moscow, Russia; Department of Biomolecular Engineering, University of California Santa Cruz, CA, USA

## Abstract

Existing human genome assemblies have almost entirely excluded highly repetitive sequences within and near centromeres, limiting our understanding of their sequence, evolution, and essential role in chromosome segregation. Here, we present an extensive study of newly assembled peri/centromeric sequences representing 6.2% (189.9 Mb) of the first complete, telomere-to-telomere human genome assembly (T2T-CHM13). We discovered novel patterns of peri/centromeric repeat organization, variation, and evolution at both large and small length scales. We also found that inner kinetochore proteins tend to overlap the most recently duplicated subregions within centromeres. Finally, we compared chromosome X centromeres across a diverse panel of individuals and uncovered structural, epigenetic, and sequence variation at single-base resolution across these regions. In total, this work provides an unprecedented atlas of human centromeres to guide future studies of their complex and critical functions as well as their unique evolutionary dynamics.

**One-sentence summary:** Deep characterization of fully assembled human centromeres reveals their architecture and fine-scale organization, variation, and evolution.

## Introduction

The human genome reference sequence has remained incomplete for two decades. Genome assembly efforts to date have excluded an estimated 5-10% of the human genome, most of which is found in and around each chromosome’s highly repetitive centromere, owing to a fundamental inability to assemble across long, repetitive sequences using short DNA sequencing reads (*1*, *2*). Centromeres function to ensure proper distribution of genetic material to daughter cells during cell division, making them critical for genome stability, fertility, and healthy development (*3*). Nearly everything known about the sequence composition of human centromeres and their surrounding regions, called pericentromeres, stems from individual experimental observations (*4–7*), low-resolution classical mapping techniques (*8*, *9*), analyses of unassembled sequencing reads (*10–13*), or recent studies of centromeric sequences on individual chromosomes (*14–16*). As a result, millions of bases in each chromosome’s peri/centromere have remained largely uncharacterized and have been omitted from essentially all contemporary genetic and epigenetic studies. Emerging long-read sequencing and assembly methods have now enabled the Telomere-to-Telomere Consortium to produce the first complete assembly of an entire human genome (T2T-CHM13) (*2*). This effort relied on careful measures to correctly assemble, polish, and validate entire centromeric and pericentromeric repeat arrays for the first time (*2*, *17*). By deeply characterizing these newly assembled sequences, we present the first high-resolution, genome-wide atlas of the sequence content and organization of human peri/centromeric regions.

Centromeres provide a robust assembly and attachment point for kinetochore proteins, which physically couple each chromosome to the mitotic or meiotic spindle (*3*). Compromised centromere function can lead to nondisjunction, a major cause of somatic and germline disease (*18*, *19*). In many eukaryotes, the centromere is composed of tandemly repeated DNA sequences, called satellite DNA, but these sequences differ widely among species (*20*, *21*). In humans, centromeres are defined by alpha satellite DNA (αSat), an AT-rich repeat family composed of ~171 bp monomers, which can occur as different subtypes repeated in a head-to-tail orientation for millions of bases (*22*, *23*). In the largest αSat arrays, different monomer subtypes belong to <u>h</u>igher <u>o</u>rder <u>r</u>epeats (HORs); for example, monomer subtypes a,b,c can repeat as abc-abc-abc (*24*, *25*). Each array can contain thousands of nearly identical HORs, but kinetochore proteins bind only a subset of HORs within a single HOR array on each chromosome (*25*). HOR arrays tend to differ in sequence and structure between chromosomes (*26*, *27*) and, like other satellite repeats, they evolve rapidly, expanding and contracting in repeat copy number over time, generating a high degree of polymorphism across individuals (*28–31*). Active (kinetochore-binding) centromeric sequences are embedded within inactive pericentromeric regions, which often include smaller arrays of diverged αSat monomers that lack HORs (*26*, *32*). Pericentromeric regions also contain transposable elements and segmental duplications, which sometimes include expressed genes (*33*, *34*), and frequently contain non-centromeric satellite repeat families (Human Satellites 1-3, beta and gamma satellite, reviewed (*35*)), which have poorly understood functions. Given the unprecedented opportunity to explore these regions in a complete human genome assembly, we investigated the localization of inner kinetochore proteins within active centromeres and surveyed sequence-based trends in the structure, function, variation, and evolution of peri/centromeric DNA.

### Complete assessment of αSat substructure and evolution

Human peri/centromeric satellite DNAs represent 6.2% of the T2T-CHM13v1.0 genome (~189.9 Mb) (Supplemental Section 1, Table S1,2, Fig. S1), which is roughly equal to the entire length of chromosome 4. Nearly all of this sequence remains unassembled in the current GRCh38/hg38 reference sequence (hereafter, hg38), including pericentromeric satellite DNA families that extend into each of the five acrocentric short arms. Based on decades of individual observations, a framework for the overall structure of a typical human peri/centromeric region has been proposed (Fig. 1A). Using the CHM13 assembly, we tested and largely confirmed this broad framework genome-wide at base-pair resolution, with some notable and surprising exceptions (Fig. 1B,C).

**Fig. 1.**
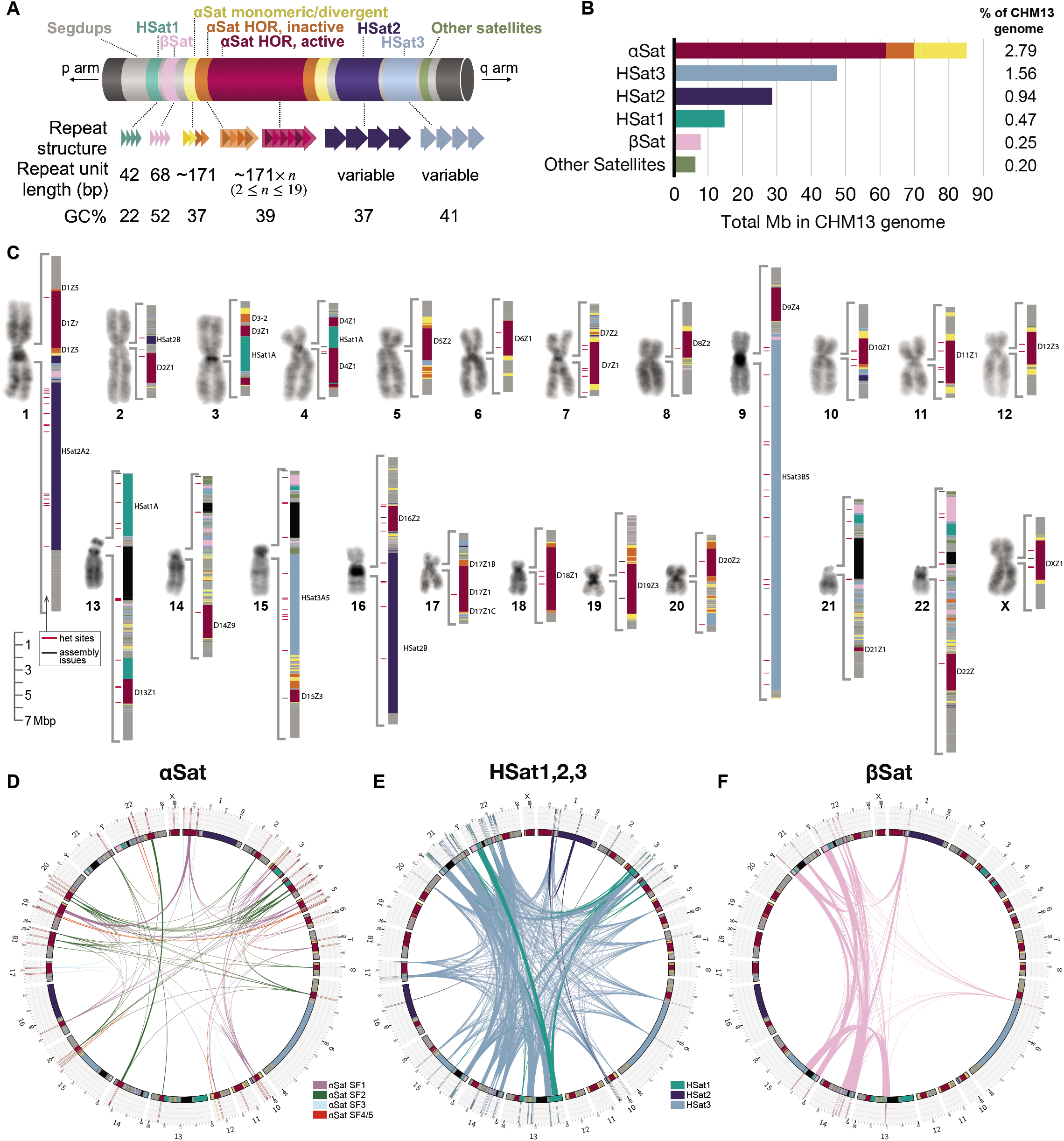
Overview of all peri/centromeric regions in CHM13. (**A**) Schematic illustrating the typical components of a human peri/centromeric region, including the canonical repeat structure, repeat unit length, and GC% of each satellite family. Each family’s assigned color is used consistently throughout the remaining figures. (**B**) Barplot showing the relative total lengths of each major satellite family genome-wide. Alpha satellite (αSat) is broken into active HORs (red), inactive HORs (orange), and divergent/monomeric (yellow). (**C**) Micrographs of representative DAPI-stained chromosomes from CHM13 metaphase spreads, next to color-coded maps of each centromere region. Large satellite arrays are labeled. To the left of each map is a track indicating the locations of potential assembly issues (gray) or sites that are likely to be heterozygous in the CHM13 cell line (red). (**D-F**) Circos plots showing genome-wide relationships in sequence content for 3 different satellite families. Connecting line widths indicate the proportion of 75-mers that are shared between arrays. αSat lines are colored by their suprachromosomal family (SF) assignment. Radial barplots indicate specificity of 75-mers proportionally, with white indicating 75-mers unique to the array, light gray indicating 75-mers shared with other centromeric regions, and black indicating 75-mers shared with regions outside of centromeres.

Consistent with prior studies (reviewed by (*36*)), all centromeric regions contain long tracts, or arrays, of alpha satellite (αSat) monomers, which are organized in a head-to-tail orientation spanning millions of bases (85 Mb total genome-wide). A genome-wide assessment of CHM13 αSat monomers revealed a broad range of pairwise sequence identity, with limited deviation in repeat unit length (median 171 bp, with 99.5% within the range 140-187 bp). We next compared these CHM13 monomers with those found in hg38, which are mostly limited to 59 Mb of centromere reference models that lack biologically meaningful long-range repeat structures (11). We found these monomer sets to be largely concordant with respect to both pairwise sequence identity (≥98%) and chromosomal localization, with only a small number of repeats specific to each respective assembly (CHM13 vs hg38) (Supplemental Section 1, Table S3).

Using previously described methods (*32*, *37*, *38*), we performed complete, monomer-by-monomer classification of all αSat into 20 distinct suprachromosomal families (SFs; Supplemental Section 1). Each family is composed of SF-specific monomer classes (Table S4). SFs annotated include novel SFs 01 and 02, which unite the sequences previously identified as archaic SF1 and SF2, respectively (*37*, *38*), and SFs13-18, which represent small pieces of the most ancient αSat and occur far from the centromere, presumably at the sites of long-defunct ancient centromeres (Table S5). Within each centromeric region, we identify between 1 and 9 HOR arrays, totaling 70 Mb genome-wide (62 Mb active HORs, 8 Mb inactive HORs, Fig. 1). Although 18 out of 23 chromosomes contain multiple, distinct HOR arrays, only one HOR array per chromosome binds inner kinetochore proteins and is thus designated as active with respect to centromere function (Table S3) (*25*). All other HOR arrays on the same chromosome are considered inactive, although it is possible for these inactive arrays to be competent for centromere function, as previously observed for ‘epialleles’ on chromosome 17 and for other cases where CENP-A is present on different HOR arrays between the two homologs (*40*, *41*). The active array on each chromosome ranges in size from 5 Mb on chr18 down to 340 kb on chr21, which is near the low end of normal variation for this centromere (*39*). Adjacent to many highly homogeneous arrays are regions of highly divergent αSat HORs, in which HOR periodicity is somewhat or even completely destroyed (*38*), as well as highly divergent monomeric layers (*32*), together totaling 15.2 Mb in CHM13.

Utilizing HOR monomers and HORs primarily inferred from the hg38 centromere reference models, we provided initial annotation for all HOR arrays in CHM13 at the monomer level, revealing 80 HORs and >1000 different monomers in HORs. All HOR arrays including HORs that were previously named received new names according to a new naming system especially designed for αSat HORs (*38*) (Table S3). Most of the HORs are represented in both the T2T-CHM13 assembly and the hg38 centromere reference models (Table S3). We confirmed the comprehensiveness and accuracy of this HOR/monomer annotation and of the CHM13 reference itself by observing the following expected features: (i) complete coverage of SF1-3 arrays by HOR annotations, (ii) the absence of significant contamination of one HOR array by monomers of the other, and by (iii) concordance with known monomer arrangements in canonical HORs. The annotated CHM13 assembly provides a unique opportunity to analyze αSat across an entire genome, which we detail below.

### New pericentromeric satellite families and detailed study of the largest human satellite arrays

Outside of the αSat arrays, we generated detailed maps of each pericentromeric region, encompassing 104.7 Mb of non-αSat sequences (Supplemental Section 1). Classical human satellites 2 and 3 (HSat2,3, totaling 28.7 and 47.6 Mb, respectively) constitute the largest contiguous satellite arrays found in the human genome, with large arrays on chromosomes 1, 9, and 16 (13.2, 27.6, and 12.7 Mb respectively). HSat2 and HSat3 are derived from a simple ancestral (CATTC)n repeat that diverged into distinct families and 14 previously characterized subfamilies (*10*, *42*). HSat1 describes two distinct sequence families that were discovered within the same AT-rich fraction of genomic DNA isolated by classic separation methods (*42*, *43*). We now provide a new naming system for these two HSat1 families to clarify their identity and origin: HSat1A (formerly “SAR”), which is a 42 bp repeat constituting the most AT-rich regions of the genome, and HSat1B (formerly “HSATI”) which is a composite of AT-rich sequences and Alu fragments, found almost entirely on the Y chromosome (*44*).

Beta satellite (βSat) represents the next-largest family after αSat and HSat1-3 (7.7 Mb genome-wide). It is enriched on the acrocentric short arms (*2*, *45*), and within the pericentromeric regions of 11 chromosomes, and is defined by a 68 bp repeat unit (*46*). βSat can be further subdivided into simple arrays and beta-composite arrays, in which βSat repeats are interspersed with LSau elements (*47–49*). Gamma satellite DNA (γSat), a well-characterized 220 bp tandem repeat on chromosomes 8 and X (*15*, *16*), was identified within all acrocentric short arms and six pericentromeric regions (*50*, *51*) (630 kb total). Although both βSat and γSat represent smaller satellite families in the human genome, they are more GC-rich than other satellites (βSat, 52%; γSat, 72%; αSat, 39%) and contain dense CpG methylation (Fig. S2). All remaining annotated pericentromeric satellite DNAs (collectively referenced as ‘p-censat’) total 5.55 Mb, with 1.19 Mb representing new satellite array predictions (*49*). Non-satellite bases observed between adjacent arrays and extending into the p-arms and q-arms are considered ‘centric transition’ regions, which largely represent long tracts of segmental duplications, including expressed genes (Supplemental Section 2) (*2*, *52*).

### Novel chromosomal localizations and polymorphisms of satellite subfamilies

Distinct repeat arrays from the same satellite family show varying degrees of similarity with each other. For example, centromeres on chromosomes 13/21, 14/22 and 1/5/19 have near-identical HORs that have confounded studies in the past (25, 36–38). T2T-CHM13 is the first assembly to successfully assign each of these active arrays to its specific chromosome, permitting a more comprehensive assessment of their respective repeat structures and sequence composition. As HOR 1/5/19 presents an especially problematic case because of dimeric expansions (*38*), we demonstrated that the arrays were chromosome-specific with reference to flow-sorted chromosome libraries (Supplemental Section 3, Fig. S3). We next developed separate consensus HORs, an HMM-based automatic HOR annotation tool, and browser tracks for each array and discovered that each chromosome has a distinct haplotype characterized by chromosome-specific sequence differences (3-20 base changes per HOR) and sometimes structural variants.

To provide a genome-wide view of the overall sequence similarity between different αSat arrays, we obtained the full set of 75-mer sequences within each array and searched for exact matches to the rest of the genome (Fig. 1D), readily identifying the hierarchical evolutionary relationships between subsets of αSat arrays (which can be organized into SFs and sub-SFs; (reviewed in (*36*)). This hierarchical subfamily organization is also observed for HSats and βSats, although their inter-array divergence levels appear lower than for αSats overall (Fig. 1E-F).

Using standards developed in previous work (*10*, *26*, *32*, *37*, *38*), we assigned the largest satellite DNA families (αSat, HSat3, and HSat2) into their respective sequence subfamilies, determining some of their chromosomal localizations for the first time. For example, we identified a 280 kb HSat3 array on chr17 and found that it belongs to subfamily B1, which had never previously been localized to a particular chromosome (*10*). This subfamily is entirely specific to chr17. While we found novel chromosomal localizations of several HSat3 subfamilies (Fig. 2), we also noticed a conspicuous lack of HSat3B2 on CHM13 chr1, contrary to expectations (*10*). To examine whether this was true for other individuals, we searched for contigs overlapping the chr1 pericentromeric region across 16 haplotype-resolved draft assemblies from genetically diverse individuals, from the Human Pangenome Reference Consortium (HPRC) (*53*) (Supplemental Section 4). This revealed that the chr1 HSat3B2 array in CHM13 belongs to a haplotype with 400 kb polymorphic deletion, which we detected in 29% (8/28) of those examined (Fig. 2A).

**Fig. 2.**
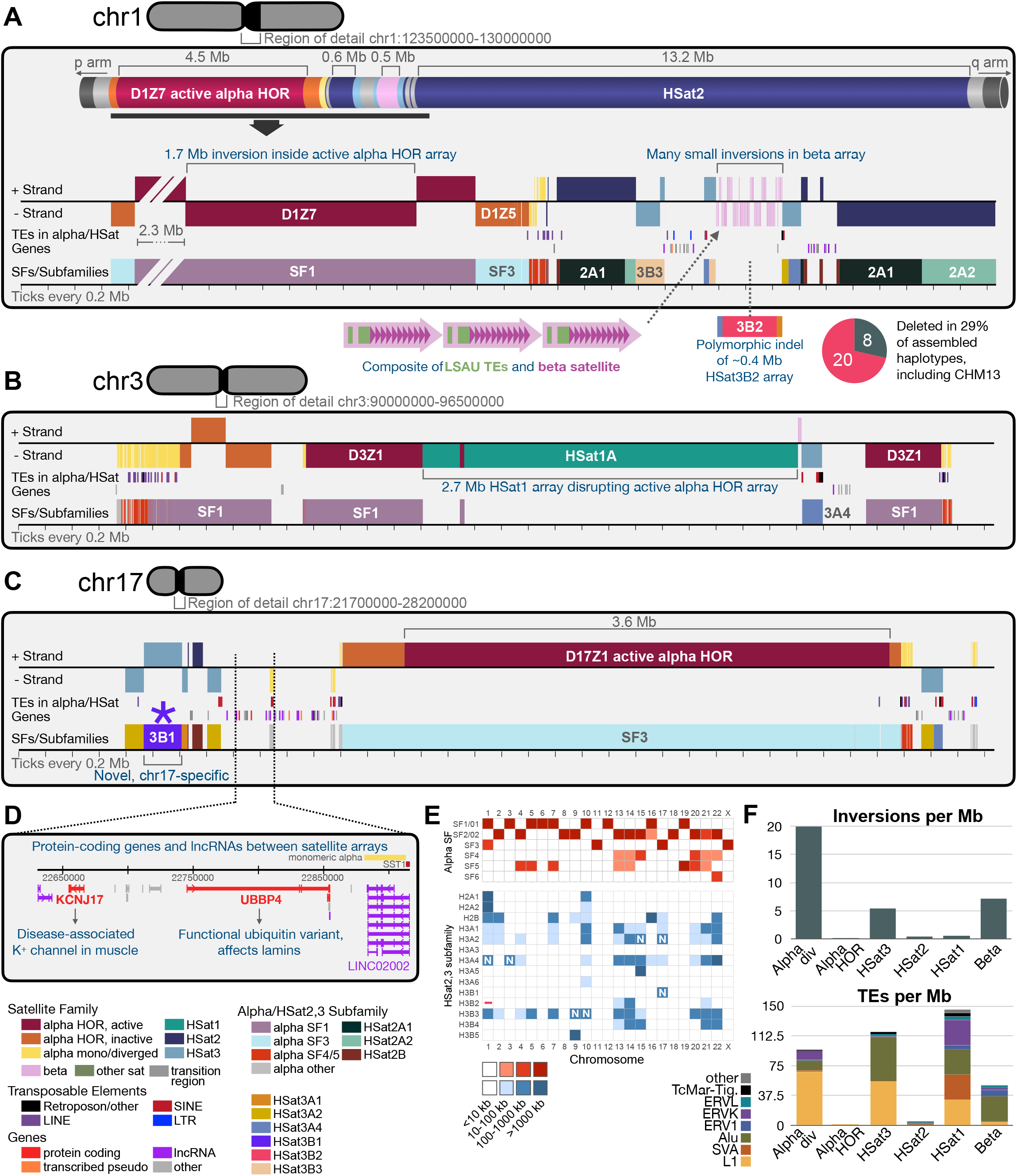
Novel discoveries in three peri/centromeric regions. (**A**) A close view of the transition region between the large αSat and HSat2 regions on chr1. The first track shows satellite family classifications, color coded as in Fig. 1 and shown at the bottom left of the figure. Strand (relative to a canonical polarity for each satellite family) is indicated by the positioning of each rectangle above (+ strand) or below (- strand) the line. The second track indicates the positions of transposable elements (TEs) that overlap αSat or HSat1,2,3, colored by TE type. The third track indicates the transcription start sites of pericentromeric gene annotations, colored by type. The fourth track indicates the subfamily assignments for HSat2 and HSat3 (according to subfamilies defined in (*10*), as well as suprachromosomal family (SF) assignments for αSat arrays. Large arrays are labeled. (**B**) As in (A) but for the peri/centromeric region of chr3. (**C**) As in (A) but for the peri/centromeric region of chr17, with the newly localized HSat3B1 array indicated with an asterisk. (**D**) A zoomed-in view of gene annotations between the αSat and HSat3 arrays on chr17, with genes colored by classification. (**E**) A heatmap showing the major and minor localizations of each αSat HOR SF (upper, red) and each HSat2,3 subfamily (lower, blue). Novel localizations are indicated with the letter N. The polymorphic chr1 HSat3B2 array is marked with a ‘-’. Note: HSat3A3 and 3A6 are almost entirely found on chrY, which is not present in CHM13. (**F**) Barplots illustrating the number of inversion breakpoints (strand switches) or the number and type of TEs detected per megabase within different satellite families genome-wide. “div” = divergent alpha satellite (divergent HORs + monomeric).

### Novel structural rearrangements and genes in peri/centromeric regions

Annotating strand orientations across entire satellite arrays revealed several novel and unexpected anomalies (Fig. 2, Table S6,7). While diverged αSats are known to contain many sequence inversions (*54*), we quantified this phenomenon genome-wide for the first time (Fig. 2F), and found a 1.7 Mb inversion inside the active αSat HOR array on chr1 (Fig. 2A), along with inversions in inactive HORs on chromosomes 3, 16, and 20 (Fig. 2B). Surprisingly, the large pericentromeric HSat3B5 array on chr9 and the beta satellite arrays on chr1 and the acrocentrics (Fig. 2A and Fig. S4) contain over 200 inversion breakpoints. Apart from inversions, two multi-megabase HSat1A arrays appear to have inserted and expanded within the active HOR arrays of chromosomes 3 and 4 (Fig. 2B), and given the concordant strand orientations of the flanking αSat arrays, these apparent insertions were unlikely to have arisen from inversions. The large insertions and inversions within active HOR arrays are particularly surprising (Fig 2, chrs 1, 3, 4), because they reveal dynamism within an area of the genome previously considered highly homogeneous (*30*, *55*). We sought to investigate these further by searching for evidence of these insertion/inversion breakpoints in the set of HPRC draft assemblies (Fig. S5). We found that the chr1 active HOR inversion is polymorphic across individuals, evident in about half of ascertainable haplotypes (11/24), while the HSat1A insertions on chr3 and chr4 were evident in all ascertainable haplotypes (32/32 and 33/33, respectively; Fig. S6). We also found evidence for an ancient duplication event that predated African ape divergence and involved a large segment of the ancient chr6 centromere plus about 1 Mb of adjacent p-arm sequence. This duplication has created a new centromere locus that hosts the current active cen6 HOR array. The duplication is visible as two nested centromere-flanking intra-chromosomal segmental duplications about 1 Mb in size and an old ~200 kb αSat array that follows the q-arm duplication and presumably contains the decayed remnants of the old centromere (Supplemental Section 5, Table S8).

Like inversions and insertions, transposable elements (TEs) are virtually absent from homogeneous HOR arrays but are enriched in divergent αSat (Fig. 2F) (*56*, *57*). The CHM13 assembly also revealed that certain novel satellites are composed entirely of combinations of TEs, which we refer to as “composite satellites” (*49*) (Hoyt et al. 2021). Consistent with individual published observations (*44*, *47*, *58*), we also found that other satellites, such as HSat1, HSat3, and βSat, often include fragments of ancient TEs as part of their repeating units (Fig. 2A,F)—a phenomenon we rarely observe in αSat HOR arrays (Fig. S7).

Finally, we compared our pericentromeric maps to gene annotations (Table S9,10). One region on chr17, located between the large HSat3 and αSat arrays (Fig. 2D), contains two protein-coding genes: *KCNJ17*, which encodes a disease-associated potassium channel in muscle cells (*59*), and *UBBP4*, which encodes a functional ubiquitin variant that may play a role in regulating nuclear lamins (*60*). Notably, *KCNJ17* is missing from GRCh38, which causes inaccurate and missed variant calls in homologous genes *KCNJ12* and *KCNJ18* (*61*). Interestingly, this region also contains a long noncoding RNA annotation (*LINC02002*), which starts inside an SST1 element and continues into an adjacent 33 kb array of divergent αSat (Fig. 2D). Unexpectedly, we also identified a processed paralog of an apoptosis-related protein-coding gene, *BCLAF1* (BCL2 Associated Transcription Factor 1), as part of a segmental duplication embedded within an inactive HOR array on chr16 (Fig. S8).

### New methods uncover the fine repeat structure of satellite DNA arrays

To further chart the structure of peri/centromeric regions at high resolution, we compared individual repeat units within and between different satellite arrays. We decomposed each αSat HOR array first into individual monomers and then into entire HORs, revealing the positions of full-size canonical HORs and structural variant HORs resulting from insertions or deletions (Supplemental Section 6, Table S11). These indels are often not in register with arbitrarily chosen monomer start sites, creating hybrid monomers. For example, if the canonical HOR structure is abc-abc-abc, a deletion variant might occur as ac-ac-ac (i.e. an in-register deletion of b) or a/bc-a/bc-a/bc where a/b is a hybrid (i.e. an out-ofregister deletion overlapping the junction between a and b). HOR structures were characterized by two different approaches: we applied hg38-based manual HOR inference to the CHM13 assembly (*38*), and performed de-novo inference from CHM13 assembly (Supplemental Section 6). Both approaches yielded the same canonical HORs for active arrays (*62–64*). We also searched for these canonical and variant HOR types in HiFi sequencing reads from 16 genetically diverse individuals. While some chromosomes, such as chr7, are composed almost entirely of canonical HOR units, other chromosomes, such as chr10, contain many structural variant HOR types, with high variation in the relative frequency of these variants across individuals (Fig. 3A and Fig. S9).

**Fig. 3.**
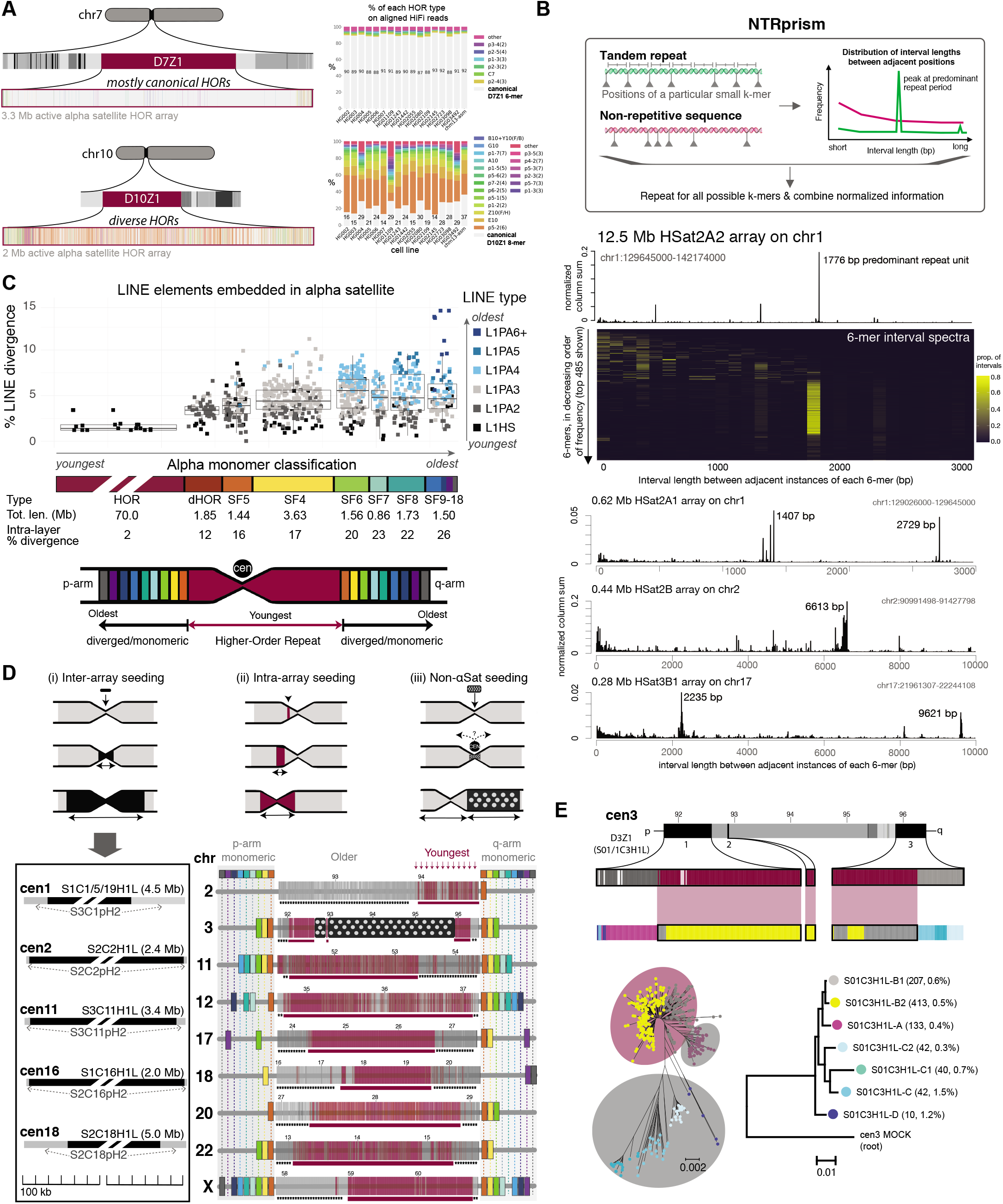
Genome-wide evidence of layered expansions in centromeric arrays. (**A**) On the left are tracks showing HOR structural variant positions across the active αSat arrays on chr7 and chr10. Canonical HORs are shown in light gray, and variants are shown in various colors. On the right are barplots for these same chromosomes showing the proportions of the same structural variant types among HORs identified on HiFi sequencing reads from 16 diverse human cell lines. (**B**) Illustration and demonstration of the NTRprism method for identifying repeat periodicities within any satellite DNA family, including novel periodicities identified for 3 arrays below. (**C**) Comparing the age and divergence of LINE transposable elements to the respective age and divergence of the αSat suprachromosomal families (SFs) that they are embedded in. (**D**) Three different modes of centromere displacement supported by the CHM13 assembly. (i) Five centromeres in which the active HOR array is surrounded by inactive HOR arrays from the same origin, consistent with insertion and expansion of the active array. (ii-iii) Active HOR arrays from 9 chromosomes, illustrating the ordering of monomeric suprachromosomal families surrounding the array on the p and q arms (rainbow colors), along with the locations of two major HOR-haplotypes (red and gray) within each array, supporting the recent expansion of a new HOR-hap from the center of the array. The HSat1 insertion and expansion in cen3 is denoted with polka dots. (**E**) A zoomed-in view of the major αSat HORs in cen 3 (red and gray, as in D), further subdivided into finer HOR-hap clusters, showing further HOR-hap symmetry. Bottom left: a radial tree showing the phylogenetic relationships between all HORs, colored by fine HOR-hap assignments as in the track above. Red and Gray ellipses show the major HOR-hap assignments of each subclade. Bottom right: a phylogenetic tree built from HOR-hap consensus sequences, rooted with a ‘mock’ cen3 repeat representing an ancestral sequence.

Repeat structure decomposition of other satellite families is less straightforward because, unlike αSat, some families have inconsistent or unknown repeat unit sizes. For example, although both HSat2 and HSat3 are thought to have evolved from an ancient simple repeat of (CATTC)n, they have long since diverged on different chromosomes and, at least in some cases, they have been shown to be composed of longer repeat units on the order of multiple kilobases (*10*). We propose calling these longer repeat units nested tandem repeats (NTRs), to distinguish them from higher order repeats, which are composed of discrete numbers of monomers of similar lengths. To expand our ability to annotate repeat structure within newly assembled satellite DNA arrays, we created NTRprism, a versatile algorithm for discovering and visualizing satellite repeat periodicity (Fig. 3B and Fig. S10). NTRprism is somewhat analogous to classical restriction digest experiments that revealed repeat periodicities in certain satellite families (*65*), but it is greatly enhanced by the ability to computationally examine all possible k-mers, not just those targeted by restriction enzymes. Using this tool, we discovered new HORs in HSat1 and βSat arrays, as well as new NTRs in multiple HSat2,3 arrays (Fig. 3B and Fig. S10). We also applied this tool in smaller windows across individual arrays, showing that repeat periodicity can vary across an array, consistent with NTRs evolving and expanding hyper-locally in some cases (Fig. S10).

### Genome-wide evidence of layered expansions in centromeric arrays

Previous αSat studies have hypothesized a layered expansion model for centromeric αSat arrays (*36*), in which distinct new repeats periodically emerge and expand within an active array, displacing the older repeats sideways and becoming the new site of kinetochore assembly. Over time, distinct layers of progressively older and more divergent repeats are expected to expand out on either side, flanking the active centromere core with mirror symmetry (Fig. 3C). The repeats that seed new layers may originate from outside the array (e.g. by insertion of an αSat sequence from another chromosome) or from mutations within the same array (Fig. 3D). As the new centromere core expands, the flanks rapidly shrink and accumulate mutations, inversions, TE insertions, and other satellite expansions (*16*, *32*, *38*). Previous efforts to document this layered expansion pattern have focused on divergent αSat compartments that surround HOR arrays (*32*). Here, we performed a study of active αSat arrays in their entirety, together with adjacent flanking regions, to survey the degree of peri/centromeric symmetry, divergence, and decay signatures, providing the first detailed, genome-wide evidence in support of this model.

First, in agreement with prior studies, we observed a symmetrical flanking arrangement of two types of divergent αSat: divergent HORs (dHORs) (Table S12), and monomeric αSats (Table S13), which represent ancient, decayed centromeres of primate ancestors (*32*). We classified divergent αSat into distinct SFs and dHOR families, and demonstrated how these sequences accumulate mutations, inversions, and TE insertions over time (Table S14). In monomeric αSats, L1s are common, and the age of the oldest L1s increases with the age of the αSat layer (Fig. 3C). Although L1s are extremely rare in αSat HOR arrays, when they do occur, they are always the youngest L1Hs elements, which are known to be still active in humans (Fig. 3C, Table S15) (*49*, *66*). In agreement with previous studies, we document a gradient of size (Fig. 3C), and a gradient of intra-array divergence (17 to 26%) preceded by a steep (~10%) increase that marks the transition between HOR arrays and outer layers (Fig. 3C), (*16*, *32*, *38*). In total, molecular dating of the flanking αSats by divergence (grouped by SF-layers), shared occurrence in primate lineages (*32*), and the age of embedded L1 elements revealed an age gradient away from the central active array (Fig. 3C,D, Table S15).

We next asked if the layered expansion pattern overlaps the active arrays themselves. As shown in Fig. 3D, the sequences seeding the expanding satellite array can be either introduced from within (intra-array seeding) (32) or from an external HOR (or non-HOR) array (inter-array seeding) (*67*, *68*) (Table S16). In total, we document five cases of inter-array type symmetry (Fig. 3D) of which only one was known before (*69*). In some cases of the inter-array model, the active HOR array originates from a different SF than the flanking inactive array (chrs 1 and 16) (Fig. 3D). This, together with a well-studied case of inter-array symmetry in cen17 (*70*) provides evidence of how entire arrays have been displaced recently in favor of an introduced sequence.

Moreover, detailed study of active arrays in their entirety provided evidence of intra-array symmetry, defined by classification of HORs by their shared sequence variants. Such variants were known for decades (*30*, *71*, *72*) and recently were noted in the first completely assembled centromeres from chromosomes X (*73*, *74*) and 8 (*16*), where the central part of the active array was found to contain HOR variants slightly different from those on the flanks. To test if this array structure is typical, we aligned individual HOR units within the same array and clustered them into “HOR-haplotypes” or “HOR-haps” (Supplemental Section 6). Initial broad classifications of entire arrays into 2-4 distinct HOR-haps revealed that active HOR arrays are also composed of distinct layers, which typically expand from the middle (dark red versus grey, Fig. 3D).

Further classification of sequences within each broad HOR-hap identified additional substructure, and evidence for symmetric patterns (Fig. 3E). To examine whether the middle HOR-haps are likely to be the youngest evolutionarily, we built phylogenetic trees of consensus HOR-haps (Fig. 3E) and rooted them using reconstructed “mock” SF-ancestral sequences built from consensus monomers for each SF. We also performed complete phylogenetic analysis of all HORs. The identification of evolutionarily younger and older HOR-haps was supported by both methods, as shown for chr3 (Fig. 3E). In addition, the intra-array divergence in central HOR-haps is usually slightly lower than in the flanking arrays, indicating that the central HOR-haps have expanded more recently. Together, these findings present strong genome-wide evidence for a layered expansion pattern within active arrays.

### Satellite array organization at sites of kinetochore assembly

Human centromeres are defined epigenetically as the specific subregion bound by inner kinetochore proteins within each active αSat HOR array (*21*, *75*). Centromeres contain a combination of epigenetic marks distinguishing them from the surrounding pericentromeric heterochromatin, including the presence of the histone variant CENP-A (*76*, *77*), “centrochromatin”-associated histone modifications (*78*), and reduced CpG methylation (*15*, *16*, *79*). To study HOR organization at sites of kinetochore assembly, we identified discrete regions of CENP-A enrichment within each active array using published native CHM13 ChIP-seq (NChIP) data (*16*) along with CUT&RUN (*80*) data generated in this study (Supplemental Section 7).

Consistent with previous studies, CENP-A binding is almost exclusively localized within αSat HOR arrays, with one active array per chromosome (*25*) (Table S17). To demarcate specific subsets of ‘active HORs’, we developed a new repeat-sensitive short-read alignment method (Fig. 4A) using a collection of informative markers across each active array to precisely map each overlapping read. We identified unique 100 bp marker sequences covering 5.9% of all bases in all active arrays in CHM13. These markers are non-uniformly distributed showing depletion of coverage at the sites of centromere protein enrichment (Fig. S11), which we determine to be due to recent, local duplications. To increase the comprehensiveness of our short-read mapping strategy, we also included “region-specific markers” observed at two or more locations within a set maximum distance from each other. These regionspecific markers allowed us to ask if sequences specific to a given region showed evidence of enrichment, broadening our coverage of the array. We studied centromere protein enrichment patterns using a combination of these single-copy and region-specific markers, either by directly determining the enrichment of these sequences across read datasets (reference-independent (*81*), Fig. S12) or by filtering read alignments based on overlap (marker-spanning alignments) (Supplemental Section 7). Use of these two orthogonal methods allowed us to determine the span of constitutively bound CENP-A within each array and delineate active HOR arrays containing each chromosome’s centromere (Table S17).

**Fig. 4.**
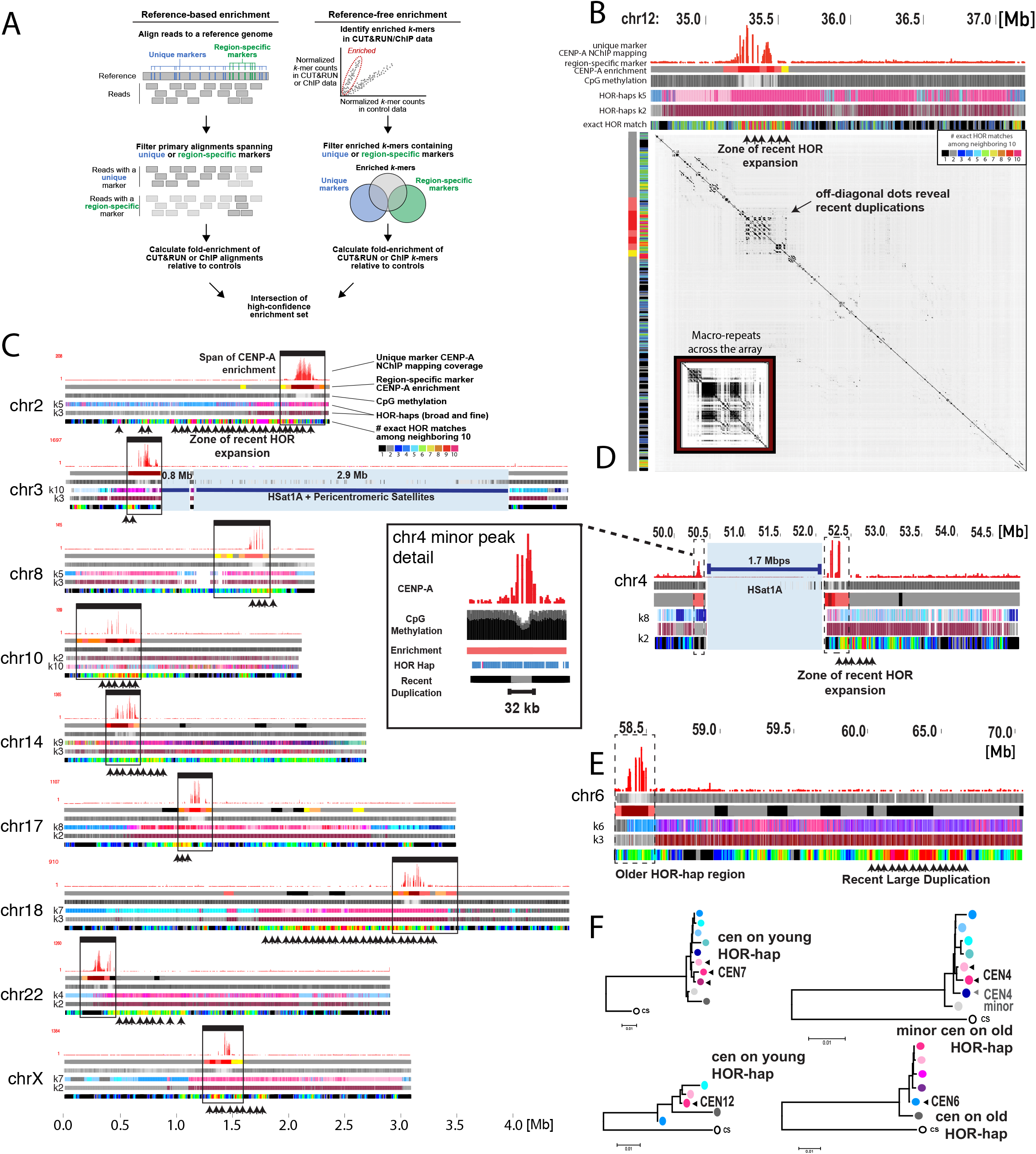
Kinetochore proteins tend to bind recently expanded centromeric subregions. (**A**) Schematic illustrating two approaches to define enrichment of kinetochore proteins in centromeric regions using short-read data from NChIP/CUT&RUN. (**B**) The chr12 αSat HOR array, with megabase coordinates listed at the top. Track 1: coverage from alignment-based marker-assisted mapping of CENP-A NChIP reads. Track 2: a heatmap representing reference-free region-specific marker enrichment in _ bp bins. Track 3: heatmap showing relative CpG methylation frequency. Tracks 4/5: HOR-haplotypes defined by kmeans clustering with k=7 or k=2. Track 6: heatmap representing the number of identical copies of each HOR occurring within the adjacent 10 repeat units, as a measure of recent duplication events. Bottom: A dotplot from a self-alignment of the array, illustrating the local nature of recent duplications, with arrows pointing to a zone of recent duplication. The inset shows a smaller view of the entire array visualized as a dotplot with a smaller word size (allowing for detection of older duplications), showing evidence of large macro-repeats across the array. (**C**) As in (B) but for 9 different centromeres, with the full span of CENP-A enrichment framed by black windows. (**D**) As in (B) tracks 1, 3, 2, 4, & 6 but for chr4, with an inset highlighting a secondary CENP-A enrichment site and minor CpG depletion site on the other side of the interrupting HSat1 array. (**E**) As in (B) tracks 1, 3, 2, 4, 5, & 6 but for chr6, which, unlike most chromosomes, has its CENP-A enrichment site over an older HOR-hap region. (**F**) Horhap consensus trees as in Fig. 3E, with the location of the CENP-A binding region indicated with an arrow.

In agreement with previous studies, we found the strongest CENP-A enrichment near sites reported to be depleted in CpG methylation, or centromere dip regions (CDRs) (*16*, *79*). Notably, some chromosomes show evidence for multiple peaks within each CDR region, which could represent interspersed domains, variation in the organization of CENP-A nucleosomes across the two homologous chromosomes, or polymorphic organization across the population of cells (*79*). Here, we extend these findings and report that the complete span of the centromere region, as defined by the CENP-A enrichment patterns, extends outside of CDRs by hundreds of kilobases across all chromosomes (Fig. 4C). Furthermore, on some chromosomes, we detected smaller regions of centromere protein enrichment outside of the primary CDR, with some overlapping a minor, secondary CDR (chromosome 4) or no CDR at all (chromosome 18) (Fig. 4C, Fig. S13). In total, these findings issue the first map of human centromeres in a complete genome. In doing so, we identified subregions within each HOR array that are competent to support kinetochore assembly and centromere function.

In relation to the layered expansion pattern, we found that CENP-A is commonly enriched in the youngest HOR-haplotypes within the majority of arrays (Table S17). Furthermore, for each centromere region we increased the number of HOR-hap clusters to study more refined groupings of HORs that are enriched for CENP-A and associated with the sites of kinetochore assembly (Fig. 4B-E). In the active array on chromosome 12, we identified CENP-A enrichment on one of two large macro-repeat structures, both presenting similar HOR-hap sequences (Fig. 4B, Fig. S14). Constructing phylogenetic trees from consensus repeats revealed a subset of HORs that are evolutionarily derived from the ancestral SF-specific class monomers (as shown in Fig. 4F for chromosomes 4, 6, 7, and 12, Fig. S15). Further investigation into the region of CENP-A enrichment on chromosome 12 revealed a zone of recent HOR expansions (i.e. eight sites of recent duplications within a ~365 kb region, (Supplemental Section 7, Fig. S16) that coincides with the CDR and distinguishes one macro-repeat region from the other. We observed similar zones of recent expansion that overlap recent HOR-haps on most other chromosomes (Fig. 4C, shown for example on centromere 7 in Fig. S15), although we identified a few notable exceptions to this general trend. For example, on chromosome 4, which has two CENP-A regions occurring on either side of a 1.7 Mb HSat1A array, we found that the larger CENP-A region spans a slightly younger HOR-hap and the smaller CENP-A region spans an older HOR-hap (Fig. 4D,F). Similarly, we observe CENP-A enrichment within an older HOR-hap layer on centromere 6, over a megabase away from the site of a recent duplication event (Fig. 4E,F). In summary, we provide support of the layered expansion model, and observe that human centromeres are commonly positioned over the youngest layers within each array and that these layers are prone to recent duplications.

### Genetic and epigenetic variation across human X centromeres

Satellite DNA arrays are known to be highly variable in size across individuals; in fact, the extremes of this size variation are often plainly visible under the microscope in chromosomal karyotypes, and they have been described for decades, yet the clinical significance of these variants remains unknown and largely unexplored (*82*, *83*). More recent studies have provided low-resolution sequencing-based evidence for variability in both satellite array lengths and the frequency of certain sequence and structural variants within human populations (*10–12*), suggesting accelerated sequence evolution in these regions compared to the rest of the genome. However, satellite array variation and evolution remain poorly understood at base-level resolution due to the lack of complete centromere assemblies.

To address this, we deeply characterized and compared centromere array assemblies from chrX across seven diverse males, thus capturing the full extent of biologically important sequence variation (Fig. 5A, Supplemental Section 8, Fig. S17). We assigned repeats to seven HOR-haps, revealing both localized and broad variation within each array. For example, we identified large, tandem duplications (spanning hundreds of kilobases) in two assemblies relative to CHM13 (HG01109, PUR and HG03492, PJL, Fig. S18). Four of the seven arrays contain zones of recent duplication in the younger HOR-hap, in a similar position to that of CHM13, with all remaining assemblies showing a trend of recent duplication within a shared region closer to the p-arm (spanning different subsets of more divergent and less derived older HOR-haps). Notably, we found evidence for an additional HOR-hap type in ancient lineages (*84*) that did not participate in the late pleistocene emigration of modern humans from Africa, (Fig. 5A, dark red), representing an independent core of expanding centromeric sequence.

**Fig. 5.**
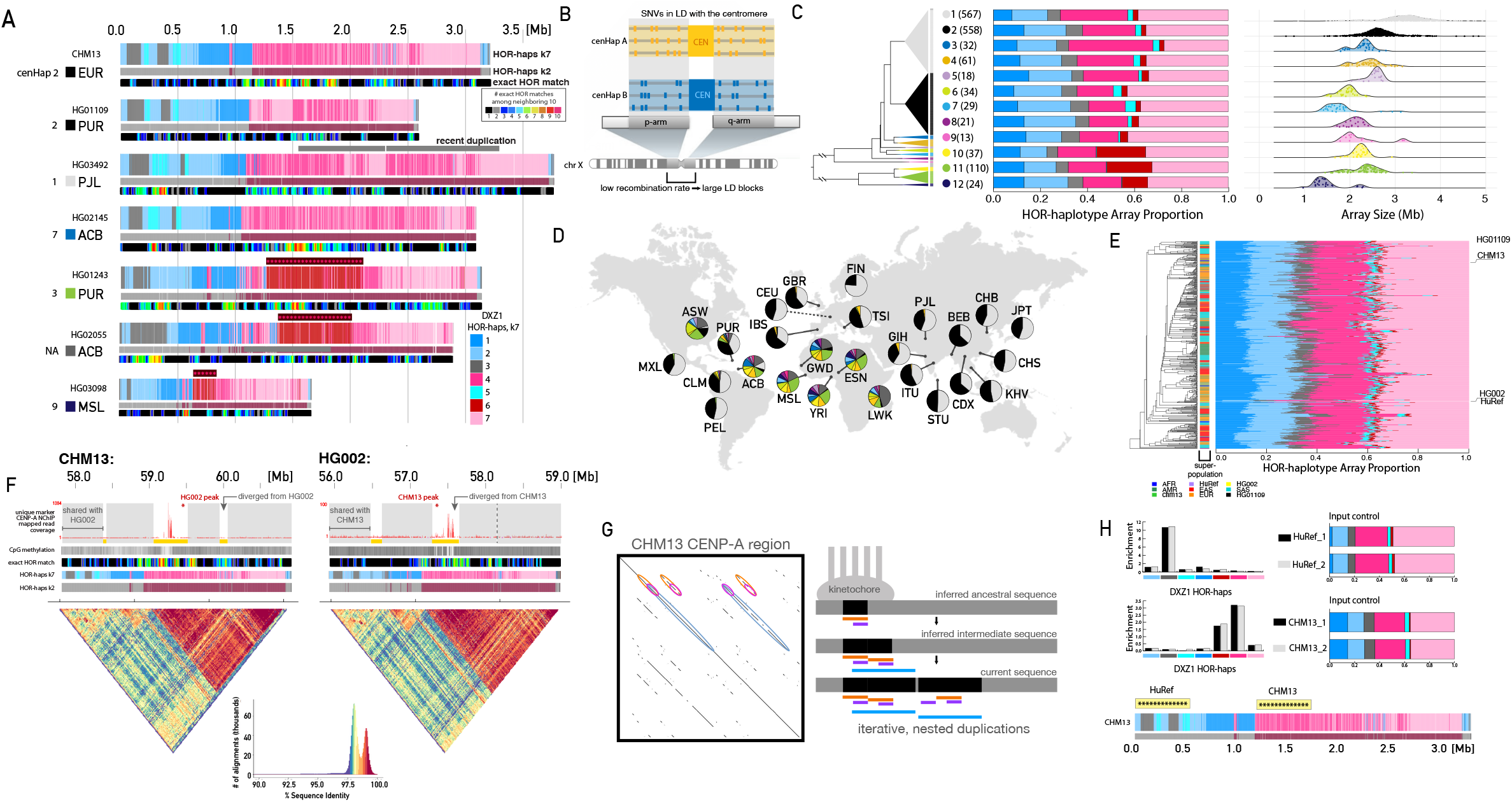
Evidence for substantial genetic and epigenetic variation in a human centromere. (**A**) Comparing the active αSat HOR array on chrX (DXZ1) between CHM13 (top) and 6 HiFi read assemblies from diverse cell lines. Tracks indicate HOR-hap classifications with k=7 (top) and k=2 (middle) along with recent HOR duplication events (bottom, as in Fig. 4B). An African-specific HOR-hap (dark red) is emphasized with red asterisks. A recent duplication in HG03492 is indicated with gray bars above. (**B**) A schematic illustrating the concept of cenhaps, in which the low local recombination rate in the centromere proximal regions lead to the evolution of single nucleotide variants into large, centromere-spanning haplotypes. (**C**) Left: tree illustrating the relationships of 12 cenhaps defined using short-read data from 1599 genetically diverse males from the 1000 Genomes Project. The height of each cenhap triangle is proportional to the number of individuals in that cenhap. Middle: barplots illustrating the average HOR-hap compositions for all individuals within each cenhap. Right: ridgeline plots indicating the distribution of estimated total array sizes for all individuals within each cenhap, with individual values represented as jittered points. (**D**) World map showing the locations of all populations represented among the 1599 males, with pie charts indicating the proportion of cenhap assignments within each population, with the same colors used in (C). (**E**) A detailed tree showing the relationships of all cenhap 2 individuals inferred from their single-nucleotide variant data, along with a barplot as in (C) for each individual and a colorbar indicating the super-population assignment of each individual. (**F**) Comparison of the DXZ1 assembly for CHM13 and HG002, which are both in cenhap2. Tracks are as in (A), with the addition of gray shading to the top track to indicate regions that align closely between the two individuals, and yellow indicating high divergence between the two individuals. Dotted line indicates the homologous site of a chm13 expansion on the HG002 array. Bottom: dotplots representing the % identity of self-alignments within the array, with a color-key and histogram below. (**G**) Left: a zoom-in of the chm13 kinetochore region with a self-alignment dotplot (exact match word size 2000) revealing patterns of recent nested duplication. Right: a model for the recent evolution in this region. (**H**) Left: comparison of CENP-A NChIP enrichment from CHM13 and HuRef cell lines in DXZ1 HOR-haps, for 2 replicates (gray/black). Right: comparison of HOR-hap assignments from input controls. Bottom: asterisks highlighting the CENP-A enriched regions from HuRef and CHM13, with respect to the CHM13 array. Colors represent HOR-haps as in (A).

Next, we studied how this variation within αSats relates to variation across single-nucleotide markers that are in linkage disequilibrium (LD) with the centromere, i.e. markers that tend to be co-inherited with the centromere. Because meiotic recombination rates are extremely low in pericentromeric regions (due to the “centromere effect”; (*85*)), centromeres are embedded in long haplotypes, which are called cenhaps (Fig 5B; (*84*)). Cenhaps are identified by first clustering pericentromeric single-nucleotide variants into phylogenetic trees, and then splitting them into large clades of shared descent. Here, we divided a group of 1599 males genotyped using published short-read sequencing data (*86*) into 12 cenhaps (with 98 individuals remaining unclassified; Fig. 5C, Fig. S19, Table S18). We also defined array-specific and HOR-hap-specific k-mer markers allowing us to utilize short-read sequencing data to estimate the absolute size of each individual’s chrX centromere array (Supplemental Section 8, Table S19) (*11*, *84*)), along with the relative proportion of that individual’s array assigned to each HOR-hap. The results revealed that different cenhaps have different αSat array size distributions as well as different average HOR-hap compositions (Fig. 5C, Fig. S20). As shown in Fig. 5D, two of the 12 cenhaps, 1 and 2, are very common outside of Africa (overall, 49% and 47%, respectively), while the other cenhaps exhibit a range of frequencies across the samples from Africa as well as those with recent African admixture (ASW, PUR, CLM, ACB). This pattern is consistent with the accepted demographic bottleneck associated with early human migration (*87*). The observed concentration of cenhap and αSat variation in African individuals underlines the need for greater representation of African genomes in pan-genome assembly efforts.

To explore the variation within one of the large cenhap groups (cenhap 2), we compared fine-scale cenhap phylogenies and HOR-hap assignments across 567 individual X chromosomes, revealing a degree of further substructure and variation in the αSat array on a more recent evolutionary timescale (Fig. 5E). To dissect this further, we compared two finished centromere assemblies from CHM13 and HG002, a cell line whose chrX array had been constructed using T2T assembly methods, and whose array structure had been experimentally validated by both pulse-field gel electrophoresis Southern blots and by digital droplet PCR (*2*). CHM13 and HG002 have similar array sizes and belong to cenhap 2 (Fig. 5E). We also studied patterns of CENP-A CUT&RUN enrichment in HG002 relative to CHM13 (*79*). We found both genomes to be highly concordant across the array, apart from three regions, where we observe recent amplifications and/or deletions of repeats (Fig. 5F). Notably, the region with the most pronounced structural differences between CHM13 and HG002 coincides with the strongest CENP-A enrichment in both arrays. Therefore, even though inner kinetochore proteins are present in both arrays over CDRs and young HOR-haps, the HOR sequences enriched with CENP-A represent local duplication events that are not shared and distinguish the two arrays (marked in yellow, Fig. 5G, Fig. S21).

Finally, we asked if CENP-A enrichment patterns were consistently found in the younger HOR-haps, as observed in CHM13 and HG002, across publicly available ChIP-seq datasets (Fig. S22). Using the T2T-CHM13 X array as a reference, we mapped these available datasets and determined CENP-A enrichment for each X HOR-hap relative to the matched input DNA. Notably, in several individuals we observed CENP-A enrichment within the older HOR-hap subregion, proximal to the p-arm, indicating the presence of a centromere X epiallele (as shown for three XY individuals, HuRef (*88*) in Fig. 5H, and also supported with data from HT1080b (*89*) and MS4221 (*90*)). Further, we compared two independent CUT&RUN experiments from the RPE-1 cell line (XX) (*91*) and found consistent evidence for heterozygous positions of CENP-A within the same cell line, with enrichment on both older and newer HOR-haps, indicating that the two X homologs carry different functional epialleles. Three additional 46,XX cell lines (IMS13q, PDNC4, K562 (*92*)) were determined to be consistent with CHM13, providing evidence that the same CENP-A+ HOR-hap is shared across both homologous X chromosomes in each line. In total, these findings uncover frequent variation in the position of the X centromere, indicating that individuals may have heterozygous and homozygous epialleles within the population. Further, these observations highlight the need to study both epigenetic and genetic variation in centromeric regions, across both related and unrelated individuals and across populations of cells over time, to better define the trends and exceptions regarding centromeric epiallele positioning and inheritance.

## Discussion

This work has produced detailed maps of previously unassembled centromeric and pericentromeric regions, which represent the largest fraction of newly introduced sequence in the complete T2T-CHM13 reference assembly (*2*). We produced detailed annotations and resources to facilitate further analysis of these complicated loci by the community. In doing so, we revealed surprising large and small scale variations in the organization and composition of active centromeres. The most dramatic variants include the interruption of two active centromeric arrays by a different satellite repeat family (on chrs 3 and 4), and a large inversion in another active array (on chr 1). We also found strong genome-wide evidence for a layered expansion model of centromere evolution, supported by ancient evolutionary patterns in the divergent satellites that flank the active centromere, as well as by recent sequence expansions within the active centromere. We defined sets of short markers specific to each array that can be used for mapping short sequencing reads and for interrogating peri/centromeric structure and function, such as designing oligo-FISH probes and guide RNAs for CRISPR-based experiments. We also demonstrated the utility of these markers to accurately localize short reads from protein-DNA interaction mapping experiments and whole-genome shotgun sequencing datasets. Furthermore, we developed a new method, NTRprism, for visualizing and quantifying tandem repeat periodicity in any satellite family, and we used this to discover novel repeat structure within multiple HSat arrays.

Our new tools and resources allowed us to characterize satellite array variation to new depths, uncovering a large polymorphic deletion of an entire HSat3 array, along with a novel expansion of a particular chrX alpha satellite HOR-haplotype within African populations. Additionally, we found a recent duplication in the chrX HOR array, representing hundreds of kilobases, that is common in individuals from a specific centromere-spanning haplotype group (cenhap 1), which can explain why individuals harboring this cenhap have larger average array sizes compared to other cenhaps. The evidence for such large duplications in human history was revealed by our assessment of centromeric macro-repeats, including those on chromosomes 12, 6, an X. The high degree of polymorphism in these regions underlines the need to produce telomere-to-telomere assemblies from many diverse individuals, to fully capture the extent of human variation in these regions and to shed light on their recent evolution and the functional consequences of this evolution. Achieving this goal will require an ability to produce accurate, complete, phased assemblies from diploid individuals. Centromeric regions would seem to present the greatest challenge for phased assembly due to their repetitive nature, but their high degree of variation may assist these efforts. Now, equipped with the T2T-CHM13 assembly and the approaches we developed here to study and compare the most challenging repetitive regions in the genome, we are optimistic that future high-quality, phased, diploid, T2T assemblies are within reach.

Finally, apart from genetic variation in these regions, we identified epigenetic variation in the location of centromere proteins within an array, as has been described previously on other chromosomes (*40*, *41*, *93*, *94*). Future investigations need to study centromeric protein localization at fine scales across many individuals, in order to better understand centromeric establishment and propagation, and how this relates to the underlying genetic variation found within each array. In light of our observation that CENP-A tends to localize to the most recently expanded HORs genome-wide, many questions remain about the evolutionary and molecular mechanisms responsible for the relationship between the kinetochore and the layered expansion patterns of satellite DNAs. It’s possible that satellite expansions occur neutrally, and more recently expanded subregions coincidentally attract the kinetochore—this is feasible in light of the evolutionary patterns we observe within non-centromeric satellite arrays. Another possibility is what we refer to as the “kinetochore selection hypothesis,” in which the kinetochore plays a causal role in amplifying particular HOR variants to which it preferentially binds (*36*). These models are not mutually exclusive, and they are also compatible with models of centromere drive or other molecular drive models (*95*, *96*). Experiments in model organisms have demonstrated that extreme array sequence variants increase meiotic and mitotic nondisjunction rates and can promote both mutational drive and/or (female) meiotic drive (*97–99*). Similar drive mechanisms, along with selection for variants that promote high-fidelity chromosome transmission, may also play a role in shaping centromeric sequence diversity in the human population.

Exploring these models will require careful experimental systems and methods for precisely measuring interactions between kinetochore proteins and repetitive DNA, as well as how these interactions affect the fidelity of chromosome transmission. While the short-read mapping methods that we developed enable the use of existing protocols like NChIP (*100*) and CUT&RUN (*80*) to provide sensitive protein-DNA interaction information at broad scales within satellite arrays, we anticipate that new long-read methods for mapping protein-DNA interactions will be essential for providing high-resolution binding footprint information, including in regions that lack single-copy or region-specific markers (*101*). We anticipate a future in which we will soon have pan-genome and pan-epigenome references in all human peri/centromeric regions, finally making them accessible for careful study using modern genomic tools.

## Supporting information

High Resolution cenHap pdf

## Acknowledgements

This work was supported, in part, by the Intramural Research Program of the National Human Genome Research Institute, National Institutes of Health (SN, SK, AR, AMM, SB, AY and AMP), the Intramural funding at the National Institute of Standards and Technology (JMZ), HHMI Hanna H. Gray Fellowship (NA), Damon Runyon Postdoctoral Fellowship, Pew Latin American Fellowship (GVC). Grants from the U.S. National Institutes of Health (NIH/NIGMS F32 GM134558 to GAL; NIH/NHGRI R01 HG00990905 to PS and AFS; NIH R01GM123312-02 to SJH, GAH, RJO; NIH R21CA240199, NSF 1643825 to RJO; NIH/NHGRI F31HG011205 to CJS; NIH/NHGRI R01 HG009190 to AG and WT; NIH/NHGRI R01HG010485, U41HG010972, U01HG010961 to KS; NIH/NIGMS R01GM132600, P20GM103546 and NIH/NHGRI U24HG010136 to DO and TJW; NIH/NHGRI R01 HG010329 to SRS; NIH/NHGRI R01HG010485, U41HG010972, U01HG010961, U24HG011853, OT2OD026682 to BP; NIH/GM R35 GM139653 and R01 GM117420 to GHK; NIH R01 GM124041, R01 GM129263, R21 CA238758 to BAS; NIH/NHGRI R01HG002385, R01HG010169 NIH/NHGRI U01 1U01HG010971 to EEE; NIH/OD/NIMH DP2 OD025824 to MYD; NIH/NHGRI U24HG010263; NHGRI U24HG006620; NCI U01CA253481; NIDDK R24 DK106766-01A1 to MCS; NIH/NHGRI U41HG007234 to MD; NIH/NHGRI R01 1R01HG011274-01, NIH/NHGRI R21 1R21HG010548-01 to KHM), National Science Foundation (NSF 1613806 to SJH, GAH, RJO; NSF DBI-1627442, NSF IOS-1732253, NSF IOS-1758800 to MCS); Mark Foundation for Cancer Research to SA and MCS (19-033-ASP); Russian Science Federation RSF 19-75-30039 (analysis of genomic repeats) to IAA; Russian Foundation for Basic Research (RFBR 18-29-13051) to LU; LU is supported by Sirius University; St. Petersburg State University ((grant ID PURE 73023573) to AM, TD, IAA and (grant ID PURE 51555639) to OK); NIH R01AG054712 to EIR; Ministry of Science and Higher Education of the Russian Federation (075-10-2020-116 (13.1902.21.0023)) to FG; Connecticut Innovations to RJO; Stowers Institute for Medical Research to JLG; AS is a Chan Zuckerberg Biohub Investigator; EEE and AFD are investigators of the Howard Hughes Medical Institute. Certain commercial equipment, instruments, or materials are identified to specify adequately experimental conditions or reported results. Such identification does not imply recommendation or endorsement by the National Institute of Standards and Technology, nor does it imply that the equipment, instruments, or materials identified are necessarily the best available for the purpose. This work utilized the computational resources of the NIH HPC Biowulf cluster (https://hpc.nih.gov).

## Author contributions

αSat sequence characterization: AVB, LU, FDR, AM, VAS, TD, OK, FG, EIR, PAP, IAA, KHM; Pericentromeric satellite characterization: NA, GAL, SJH, MEGS, DO, TJW, LGDL, AMP, RJO, KHM CUT&RUN experiments, mapping, and enrichment analyses: GVC, NA, GAL, PS, SJH, AMM, AR, MEGS, KT, SRS, AS, AFS, BAS, AFD, GHK, AG, WT, KHM; cenhap analysis and interpretation: NA, SAL, CHL, IAA, KHM; array length prediction: MB, JLG, MCS, JMZ; Methylation Analysis: AG, WT; Chromosome imaging and flow sorting: TP, JLG, SB, AY, AMP; Dotplot analysis: LU, FDR, MRV, RL, PK, AMP, IAA; Transposable element analysis: NA, SJH, GAH, RJO, LU, IAA; CHM13 satellite assembly and het analysis: NA, GAL, SN, SK, AR, AMM, AMP; UCSC genome browser and annotation workflow: MD; HiFi assemblies and quality assessment of diverse panel: NA, MA, RL, KS, AM, AVB, SA, JMZ, MCS, BP, EEE, AMP, gene annotation and expression: CJS, MRV, MH, MYD, MD; manuscript writing: NA, IAA, KHM, with input from all authors

## Data and materials availability

Sequence data are available through https://www.ncbi.nlm.nih.gov/bioproject/559484

Human Pangenome Reference Consortium (HPRC) generated long and accurate HiFi reads for sixteen human samples HG002, HG003, HG004, HG005, HG006, HG007, HG01243, HG02055, HG02109, HG02723, HG03492, HG01109, HG01442,HG02080,HG02145, and HG03098. We refer to these datasets as HPRC samples. (https://github.com/human-pangenomics/hpgp-data).

Data tracks and satellite annotations can be visualized on the UCSC Genome Browser (*102*, *103*): http://genome.ucsc.edu/cgi-bin/hgTracks?genome=t2t-chm13-v1.0&hubUrl= http://t2t.gi.ucsc.edu/chm13/hub/hub.txt

Annotation data tables and other supporting data and analysis workflows: https://github.com/kmiga/t2t_censat/

## References

1. E. E. Eichler, R. A. Clark, X. She, An assessment of the sequence gaps: unfinished business in a finished human genome. Nat. Rev. Genet. 5, 345–354 (2004).

2. S. Nurk, S. Koren, A. Rhie, M. Rautiainen, A. V. Bzikadze, A. Mikheenko, M. R. Vollger, N. Altemose, L. Uralsky, A. Gershman, S. Aganezov, S. J. Hoyt, M. Diekhans, G. A. Logsdon, M. Alonge, S. E. Antonarakis, M. Borchers, G. G. Bouffard, S. Y. Brooks, G. V. Caldas, H. Cheng, C.-S. Chin, W. Chow, L. G. de Lima, P. C. Dishuck, R. Durbin, T. Dvorkina, I. T. Fiddes, G. Formenti, R. S. Fulton, A. Fungtammasan, E. Garrison, P. G. S. Grady, T. A. Graves-Lindsay, I. M. Hall, N. F. Hansen, G. A. Hartley, M. Haukness, K. Howe, M. W. Hunkapiller, C. Jain, M. Jain, E. D. Jarvis, P. Kerpedjiev, M. Kirsche, M. Kolmogorov, J. Korlach, M. Kremitzki, H. Li, V. V. Maduro, T. Marschall, A. M. McCartney, J. McDaniel, D. E. Miller, J. C. Mullikin, E. W. Myers, N. D. Olson, B. Paten, P. Peluso, P. A. Pevzner, D. Porubsky, T. Potapova, E. I. Rogaev, J. A. Rosenfeld, S. L. Salzberg, V. A. Schneider, F. J. Sedlazeck, K. Shafin, C. J. Shew, A. Shumate, Y. Sims, A. F. A. Smit, D. C. Soto, I. Sović, J. M. Storer, A. Streets, B. A. Sullivan, F. Thibaud-Nissen, J. Torrance, J. Wagner, B. P. Walenz, A. Wenger, J. M. D. Wood, C. Xiao, S. M. Yan, A. C. Young, S. Zarate, U. Surti, R. C. McCoy, M. Y. Dennis, I. A. Alexandrov, J. L. Gerton, R. J. O’Neill, W. Timp, J. M. Zook, M. C. Schatz, E. E. Eichler, K. H. Miga, A. M. Phillippy, The complete sequence of a human genome. bioRxiv (2021), p. 2021.05.26.445798.

3. K. L. McKinley, I. M. Cheeseman, The molecular basis for centromere identity and function. Nat. Rev. Mol. Cell Biol. 17, 16–29 (2016).

4. R. Wevrick, H. F. Willard, Physical map of the centromeric region of human chromosome 7: relationship between two distinct alpha satellite arrays. Nucleic Acids Res. 19, 2295–2301 (1991).

5. M. S. Jackson, P. Slijepcevic, B. A. Ponder, The organisation of repetitive sequences in the pericentromeric region of human chromosome 10. Nucleic Acids Res. 21, 5865–5874 (1993).

6. H. E. Trowell, A. Nagy, B. Vissel, K. H. Choo, Long-range analyses of the centromeric regions of human chromosomes 13, 14 and 21: identification of a narrow domain containing two key centromeric DNA elements. Hum. Mol. Genet. 2, 1639–1649 (1993).

7. C. Tyler-Smith, Structure of repeated sequences in the centromeric region of the human Y chromosome. Development. 101 Suppl, 93–100 (1987).

8. I. Tagarro, A. M. Fernández-Peralta, J. J. González-Aguilera, Chromosomal localization of human satellites 2 and 3 by a FISH method using oligonucleotides as probes. Hum. Genet. 93, 383–388 (1994).

9. N. Archidiacono, R. Antonacci, R. Marzella, P. Finelli, A. Lonoce, M. Rocchi, Comparative mapping of human alphoid sequences in great apes using fluorescence in situ hybridization. Genomics. 25, 477–484 (1995).

10. N. Altemose, K. H. Miga, M. Maggioni, H. F. Willard, Genomic characterization of large heterochromatic gaps in the human genome assembly. PLoS Comput. Biol. 10, e1003628 (2014).

11. K. H. Miga, Y. Newton, M. Jain, N. Altemose, H. F. Willard, W. J. Kent, Centromere reference models for human chromosomes X and Y satellite arrays. Genome Res. 24, 697–707 (2014).

12. Y. Suzuki, E. W. Myers, S. Morishita, Rapid and ongoing evolution of repetitive sequence structures in human centromeres. Sci Adv. 6 (2020), doi:10.1126/sciadv.abd9230.

13. C. Alkan, M. Ventura, N. Archidiacono, M. Rocchi, S. C. Sahinalp, E. E. Eichler, Organization and evolution of primate centromeric DNA from whole-genome shotgun sequence data. PLoS Comput. Biol. 3, 1807–1818 (2007).

14. M. Jain, H. E. Olsen, D. J. Turner, D. Stoddart, K. V. Bulazel, B. Paten, D. Haussler, H. F. Willard, M. Akeson, K. H. Miga, Linear assembly of a human centromere on the Y chromosome. Nat. Biotechnol. 36, 321–323 (2018).

15. K. H. Miga, S. Koren, A. Rhie, M. R. Vollger, A. Gershman, A. Bzikadze, S. Brooks, E. Howe, D. Porubsky, G. A. Logsdon, V. A. Schneider, T. Potapova, J. Wood, W. Chow, J. Armstrong, J. Fredrickson, E. Pak, K. Tigyi, M. Kremitzki, C. Markovic, V. Maduro, A. Dutra, G. G. Bouffard, A. M. Chang, N. F. Hansen, A. B. Wilfert, F. Thibaud-Nissen, A. D. Schmitt, J.-M. Belton, S. Selvaraj, M. Y. Dennis, D. C. Soto, R. Sahasrabudhe, G. Kaya, J. Quick, N. J. Loman, N. Holmes, M. Loose, U. Surti, R. A. Risques, T. A. Graves Lindsay, R. Fulton, I. Hall, B. Paten, K. Howe, W. Timp, A. Young, J. C. Mullikin, P. A. Pevzner, J. L. Gerton, B. A. Sullivan, E. E. Eichler, A. M. Phillippy, Telomere-to-telomere assembly of a complete human X chromosome. Nature. 585, 79–84 (2020).

16. G. A. Logsdon, M. R. Vollger, P. Hsieh, Y. Mao, M. A. Liskovykh, S. Koren, S. Nurk, L. Mercuri, P. C. Dishuck, A. Rhie, L. G. de Lima, T. Dvorkina, D. Porubsky, W. T. Harvey, A. Mikheenko, A. V. Bzikadze, M. Kremitzki, T. A. Graves-Lindsay, C. Jain, K. Hoekzema, S. C. Murali, K. M. Munson, C. Baker, M. Sorensen, A. M. Lewis, U. Surti, J. L. Gerton, V. Larionov, M. Ventura, K. H. Miga, A. M. Phillippy, E. E. Eichler, The structure, function and evolution of a complete human chromosome 8. Nature. 593, 101–107 (2021).

17. A. M. Mc Cartney, Chasing perfection: validation and polishing strategies for telomere-to-telomere genome assemblies. in prep (2021).

18. R. M. Naylor, J. M. van Deursen, Aneuploidy in Cancer and Aging. Annu. Rev. Genet. 50, 45–66 (2016).

19. S. I. Nagaoka, T. J. Hassold, P. A. Hunt, Human aneuploidy: mechanisms and new insights into an age-old problem. Nat. Rev. Genet. 13, 493–504 (2012).

20. S. Henikoff, K. Ahmad, H. S. Malik, The centromere paradox: stable inheritance with rapidly evolving DNA. Science. 293, 1098–1102 (2001).

21. G. H. Karpen, R. C. Allshire, The case for epigenetic effects on centromere identity and function. Trends Genet. 13, 489–496 (1997).

22. J. C. Wu, L. Manuelidis, Sequence definition and organization of a human repeated DNA. J. Mol. Biol. 142, 363–386 (1980).

23. H. F. Willard, The genomics of long tandem arrays of satellite DNA in the human genome. Genome. 31, 737–744 (1989).

24. H. F. Willard, J. S. Waye, Hierarchical order in chromosome-specific human alpha satellite DNA. Trends Genet. 3, 192–198 (1987).

25. S. M. McNulty, B. A. Sullivan, Alpha satellite DNA biology: finding function in the recesses of the genome. Chromosome Res. 26, 115–138 (2018).

26. I. Alexandrov, A. Kazakov, I. Tumeneva, V. Shepelev, Y. Yurov, Alpha-satellite DNA of primates: old and new families. Chromosoma. 110, 253–266 (2001).

27. H. F. Willard, Chromosome-specific organization of human alpha satellite DNA. Am. J. Hum. Genet. 37, 524–532 (1985).

28. K. H. Miga, Centromeric Satellite DNAs: Hidden Sequence Variation in the Human Population. Genes. 10 (2019), doi:10.3390/genes10050352.

29. R. Oakey, C. Tyler-Smith, Y chromosome DNA haplotyping suggests that most European and Asian men are descended from one of two males. Genomics. 7, 325–330 (1990).

30. P. E. Warburton, H. F. Willard, Interhomologue sequence variation of alpha satellite DNA from human chromosome 17: evidence for concerted evolution along haplotypic lineages. J. Mol. Evol. 41, 1006–1015 (1995).

31. M. M. Mahtani, H. F. Willard, Pulsed-field gel analysis of alpha-satellite DNA at the human X chromosome centromere: high-frequency polymorphisms and array size estimate. Genomics. 7, 607–613 (1990).

32. V. A. Shepelev, A. A. Alexandrov, Y. B. Yurov, I. A. Alexandrov, The evolutionary origin of man can be traced in the layers of defunct ancestral alpha satellites flanking the active centromeres of human chromosomes. PLoS Genet. 5, e1000641 (2009).

33. X. She, J. E. Horvath, Z. Jiang, G. Liu, T. S. Furey, L. Christ, R. Clark, T. Graves, C. L. Gulden, C. Alkan, J. A. Bailey, C. Sahinalp, M. Rocchi, D. Haussler, R. K. Wilson, W. Miller, S. Schwartz, E. E. Eichler, The structure and evolution of centromeric transition regions within the human genome. Nature. 430, 857–864 (2004).

34. G. Genovese, R. E. Handsaker, H. Li, N. Altemose, A. M. Lindgren, K. Chambert, B. Pasaniuc, A. L. Price, D. Reich, C. C. Morton, M. R. Pollak, J. G. Wilson, S. A. McCarroll, Using population admixture to help complete maps of the human genome. Nat. Genet. 45, 406–14, 414e1–2 (2013).

35. C. Lee, R. Wevrick, R. B. Fisher, M. A. Ferguson-Smith, C. C. Lin, Human centromeric DNAs. Hum. Genet. 100, 291–304 (1997).

36. Karen H. Miga and Ivan A. Alexandrov, Variation and evolution of human centromeres: A field guide and perspective. Annu. Rev. Genet. (2021).

37. V. A. Shepelev, L. I. Uralsky, A. A. Alexandrov, Y. B. Yurov, E. I. Rogaev, I. A. Alexandrov, Annotation of suprachromosomal families reveals uncommon types of alpha satellite organization in pericentromeric regions of hg38 human genome assembly. Genom Data. 5, 139–146 (2015).

38. L. I. Uralsky, V. A. Shepelev, A. A. Alexandrov, Y. B. Yurov, E. I. Rogaev, I. A. Alexandrov, Classification and monomer-by-monomer annotation dataset of suprachromosomal family 1 alpha satellite higher-order repeats in hg38 human genome assembly. Data Brief. 24, 103708 (2019).

39. A. W. Lo, G. C. Liao, M. Rocchi, K. H. Choo, Extreme reduction of chromosome-specific alphasatellite array is unusually common in human chromosome 21. Genome Res. 9, 895–908 (1999).

40. K. A. Maloney, L. L. Sullivan, J. E. Matheny, E. D. Strome, S. L. Merrett, A. Ferris, B. A. Sullivan, Functional epialleles at an endogenous human centromere. Proc. Natl. Acad. Sci. U. S. A. 109, 13704–13709 (2012).

41. K. E. Hayden, E. D. Strome, S. L. Merrett, H.-R. Lee, M. K. Rudd, H. F. Willard, Sequences associated with centromere competency in the human genome. Mol. Cell. Biol. 33, 763–772 (2013).

42. J. Prosser, M. Frommer, C. Paul, P. C. Vincent, Sequence relationships of three human satellite DNAs. J. Mol. Biol. 187, 145–155 (1986).

43. G. Corneo, E. Ginelli, E. Polli, A satellite DNA isolated from human tissues. J. Mol. Biol. 23, 619–622 (1967).

44. M. Frommer, J. Prosser, P. C. Vincent, Human satellite I sequences include a male specific 2.47 kb tandemly repeated unit containing one Alu family member per repeat. Nucleic Acids Res. 12, 2887–2900 (1984).

45. G. M. Greig, H. F. Willard, Beta satellite DNA: characterization and localization of two subfamilies from the distal and proximal short arms of the human acrocentric chromosomes. Genomics. 12, 573–580 (1992).

46. J. S. Waye, H. F. Willard, Human beta satellite DNA: genomic organization and sequence definition of a class of highly repetitive tandem DNA. Proc. Natl. Acad. Sci. U. S. A. 86, 6250–6254 (1989).

47. A. Agresti, R. Meneveri, A. G. Siccardi, A. Marozzi, G. Corneo, S. Gaudi, E. Ginelli, Linkage in human heterochromatin between highly divergent Sau3A repeats and a new family of repeated DNA sequences (HaeIII family). J. Mol. Biol. 205, 625–631 (1989).

48. R. Meneveri, A. Agresti, A. Marozzi, S. Saccone, M. Rocchi, N. Archidiacono, G. Corneo, G. Della Valle, E. Ginelli, Molecular organization and chromosomal location of human GC-rich heterochromatic blocks. Gene. 123, 227–234 (1993).

49. S. Hoyt, From telomere to telomere: the transcriptional and epigenetic state of human repeat elements. bioRxiv (in review) (2021).

50. C. C. Lin, R. Sasi, C. Lee, Y. S. Fan, D. Court, Isolation and identification of a novel tandemly repeated DNA sequence in the centromeric region of human chromosome 8. Chromosoma. 102, 333–339 (1993).

51. C. Lee, X. Li, E. W. Jabs, D. Court, C. C. Lin, Human gamma X satellite DNA: an X chromosome specific centromeric DNA sequence. Chromosoma. 104, 103–112 (1995).

52. M. R. Vollger, X. Guitart, P. C. Dishuck, L. Mercuri, W. T. Harvey, A. Gershman, M. Diekhans, A. Sulovari, K. M. Munson, A. M. Lewis, K. Hoekzema, D. Porubsky, R. Li, S. Nurk, S. Koren, K. H. Miga, A. M. Phillippy, W. Timp, M. Ventura, E. E. Eichler, Segmental duplications and their variation in a complete human genome. bioRxiv (2021), p. 2021.05.26.445678.

53. HPRC, Human Pangenome Reference Consortium. T2T Diversity Panel (2021), (available at https://github.com/human-pangenomics/hpgp-data).

54. M. K. Rudd, H. F. Willard, Analysis of the centromeric regions of the human genome assembly. Trends Genet. 20, 529–533 (2004).

55. P. E. Warburton, H. F. Willard, Genomic analysis of sequence variation in tandemly repeated DNA. Evidence for localized homogeneous sequence domains within arrays of alpha-satellite DNA. J. Mol. Biol. 216, 3–16 (1990).

56. A. E. Kazakov, V. A. Shepelev, I. G. Tumeneva, A. A. Alexandrov, Y. B. Yurov, I. A. Alexandrov, Interspersed repeats are found predominantly in the “old” α satellite families. Genomics. 82, 619–627 (2003).

57. M. G. Schueler, A. W. Higgins, M. K. Rudd, K. Gustashaw, H. F. Willard, Genomic and genetic definition of a functional human centromere. Science. 294, 109–115 (2001).

58. R. Bandyopadhyay, C. McQuillan, S. L. Page, K. H. Choo, L. G. Shaffer, Identification and characterization of satellite III subfamilies to the acrocentric chromosomes. Chromosome Res. 9, 223–233 (2001).

59. D. P. Ryan, M. R. D. da Silva, T. W. Soong, B. Fontaine, M. R. Donaldson, A. W. C. Kung, W. Jongjaroenprasert, M. C. Liang, D. H. C. Khoo, J. S. Cheah, S. C. Ho, H. S. Bernstein, R. M. B. Maciel, R. H. Brown Jr, L. J. Ptácek, Mutations in potassium channel Kir2.6 cause susceptibility to thyrotoxic hypokalemic periodic paralysis. Cell. 140, 88–98 (2010).

60. M.-L. Dubois, A. Meller, S. Samandi, M. Brunelle, J. Frion, M. A. Brunet, A. Toupin, M. C. Beaudoin, J.-F. Jacques, D. Lévesque, M. S. Scott, P. Lavigne, X. Roucou, F.-M. Boisvert, UBB pseudogene 4 encodes functional ubiquitin variants. Nat. Commun. 11, 1306 (2020).

61. Sergey Aganezov, Stephanie M. Yan, Daniela C. Soto, Melanie Kirsche, Samantha Zarate, A complete reference genome improves analysis of human genetic variation. bioRxiv (in review) (2021).

62. T. Dvorkina, A. V. Bzikadze, P. A. Pevzner, The string decomposition problem and its applications to centromere analysis and assembly. Bioinformatics. 36, i93–i101 (2020).

63. Tatiana Dvorkina, Olga Kunyavskaya, Andrey V. Bzikadze, Ivan Alexandrov, Pavel A. Pevzner, CentromereArchitect: inference and analysis of the architecture of centromeres. Bioinformatics (2021).

64. Olga Kunyavskaya, Tatiana Dvorkina, Andrey V. Bzikadze, Ivan Alexandrov, Pavel A. Pevzner, HORmon: automated annotation of human centromeres. in prep (2021).

65. L. Manuelidis, Repeating restriction fragments of human DNA. Nucleic Acids Res. 3, 3063–3076 (1976).

66. C. R. Beck, P. Collier, C. Macfarlane, M. Malig, J. M. Kidd, E. E. Eichler, R. M. Badge, J. V. Moran, LINE-1 retrotransposition activity in human genomes. Cell. 141, 1159–1170 (2010).

67. I. A. Alexandrov, S. P. Mitkevich, Y. B. Yurov, The phylogeny of human chromosome specific alpha satellites. Chromosoma. 96, 443–453 (1988).

68. H. F. Willard, J. S. Waye, Chromosome-specific subsets of human alpha satellite DNA: analysis of sequence divergence within and between chromosomal subsets and evidence for an ancestral pentameric repeat. J. Mol. Evol. 25, 207–214 (1987).

69. P. Finelli, R. Antonacci, R. Marzella, A. Lonoce, N. Archidiacono, M. Rocchi, Structural organization of multiple alphoid subsets coexisting on human chromosomes 1, 4, 5, 7, 9, 15, 18, and 19. Genomics. 38, 325–330 (1996).

70. M. K. Rudd, G. A. Wray, H. F. Willard, The evolutionary dynamics of alpha-satellite. Genome Res. 16, 88–96 (2006).

71. S. J. Durfy, H. F. Willard, Concerted evolution of primate alpha satellite DNA. Evidence for an ancestral sequence shared by gorilla and human X chromosome alpha satellite. J. Mol. Biol. 216, 555–566 (1990).

72. P. E. Warburton, R. Wevrick, M. M. Mahtani, H. F. Willard, Pulsed-Field and Two-Dimensional Gel Electrophoresis of Long Arrays of Tandemly Repeated DNA: Analysis of Human Centromeric Alpha Satellite. Pulsed-Field Gel Electrophoresis, pp. 299–318.

73. A. V. Bzikadze, P. A. Pevzner, Automated assembly of centromeres from ultra-long error-prone reads. Nat. Biotechnol. 38, 1309–1316 (2020).

74. K. H. Miga, Centromere studies in the era of “telomere-to-telomere” genomics. Exp. Cell Res. 394, 112127 (2020).

75. M. D. Blower, B. A. Sullivan, G. H. Karpen, Conserved organization of centromeric chromatin in flies and humans. Dev. Cell. 2, 319–330 (2002).

76. A. A. Van Hooser, I. I. Ouspenski, H. C. Gregson, D. A. Starr, T. J. Yen, M. L. Goldberg, K. Yokomori, W. C. Earnshaw, K. F. Sullivan, B. R. Brinkley, Specification of kinetochore-forming chromatin by the histone H3 variant CENP-A. J. Cell Sci. 114, 3529–3542 (2001).

77. B. E. Black, D. W. Cleveland, Epigenetic centromere propagation and the nature of CENP-a nucleosomes. Cell. 144, 471–479 (2011).

78. B. A. Sullivan, G. H. Karpen, Centromeric chromatin exhibits a histone modification pattern that is distinct from both euchromatin and heterochromatin. Nat. Struct. Mol. Biol. 11, 1076–1083 (2004).

79. A. Gershman, M. E. G. Sauria, P. W. Hook, S. J. Hoyt, R. Razaghi, S. Koren, N. Altemose, G. V. Caldas, M. R. Vollger, G. A. Logsdon, A. Rhie, E. E. Eichler, M. C. Schatz, R. J. O’Neill, A. M. Phillippy, K. H. Miga, W. Timp, Epigenetic Patterns in a Complete Human Genome. bioRxiv (2021), p. 2021.05.26.443420.

80. P. J. Skene, S. Henikoff, An efficient targeted nuclease strategy for high-resolution mapping of DNA binding sites. Elife. 6 (2017), doi:10.7554/eLife.21856.

81. O. K. Smith, C. Limouse, K. A. Fryer, N. A. Teran, K. Sundararajan, R. Heald, A. F. Straight, Identification and characterization of centromeric sequences in Xenopus laevis. Genome Res. 31, 958–967 (2021).

82. Y. B. Yurov, S. P. Mitkevich, I. A. Alexandrov, Application of cloned satellite DNA sequences to molecular-cytogenetic analysis of constitutive heterochromatin heteromorphisms in man. Hum. Genet. 76, 157–164 (1987).

83. B. Erdtmann, Aspects of evaluation, significance, and evolution of human C-band heteromorphism. Hum. Genet. 61, 281–294 (1982).

84. S. A. Langley, K. H. Miga, G. H. Karpen, C. H. Langley, Haplotypes spanning centromeric regions reveal persistence of large blocks of archaic DNA. Elife. 8 (2019), doi:10.7554/eLife.42989.

85. M. Nambiar, G. R. Smith, Repression of harmful meiotic recombination in centromeric regions. Semin. Cell Dev. Biol. 54, 188–197 (2016).

86. M. Byrska-Bishop, U. S. Evani, X. Zhao, A. O. Basile, H. J. Abel, A. A. Regier, A. Corvelo, W. E. Clarke, R. Musunuri, K. Nagulapalli, S. Fairley, A. Runnels, L. Winterkorn, E. Lowy-Gallego, The Human Genome Structural Variation Consortium, P. Flicek, S. Germer, H. Brand, I. M. Hall, M. E. Talkowski, G. Narzisi, M. C. Zody, High coverage whole genome sequencing of the expanded 1000 Genomes Project cohort including 602 trios. bioRxiv (2021), p. 2021.02.06.430068.

87. P. H. Sudmant, T. Rausch, E. J. Gardner, R. E. Handsaker, A. Abyzov, J. Huddleston, Y. Zhang, K. Ye, G. Jun, M. H.-Y. Fritz, M. K. Konkel, A. Malhotra, A. M. Stütz, X. Shi, F. P. Casale, J. Chen, F. Hormozdiari, G. Dayama, K. Chen, M. Malig, M. J. P. Chaisson, K. Walter, S. Meiers, S. Kashin, E. Garrison, A. Auton, H. Y. K. Lam, X. J. Mu, C. Alkan, D. Antaki, T. Bae, E. Cerveira, P. Chines, Z. Chong, L. Clarke, E. Dal, L. Ding, S. Emery, X. Fan, M. Gujral, F. Kahveci, J. M. Kidd, Y. Kong, E.-W. Lameijer, S. McCarthy, P. Flicek, R. A. Gibbs, G. Marth, C. E. Mason, A. Menelaou, D. M. Muzny, B. J. Nelson, A. Noor, N. F. Parrish, M. Pendleton, A. Quitadamo, B. Raeder, E. E. Schadt, M. Romanovitch, A. Schlattl, R. Sebra, A. A. Shabalin, A. Untergasser, J. A. Walker, M. Wang, F. Yu, C. Zhang, J. Zhang, X. Zheng-Bradley, W. Zhou, T. Zichner, J. Sebat, M. A. Batzer, S. A. McCarroll, 1000 Genomes Project Consortium, R. E. Mills, M. B. Gerstein, A. Bashir, O. Stegle, S. E. Devine, C. Lee, E. E. Eichler, J. O. Korbel, An integrated map of structural variation in 2,504 human genomes. Nature. 526, 75–81 (2015).

88. J. G. Henikoff, J. Thakur, S. Kasinathan, S. Henikoff, A unique chromatin complex occupies young α-satellite arrays of human centromeres. Sci Adv. 1 (2015), doi:10.1126/sciadv.1400234.

89. J. Thakur, S. Henikoff, CENPT bridges adjacent CENPA nucleosomes on young human α-satellite dimers. Genome Res. 26, 1178–1187 (2016).

90. D. Hasson, T. Panchenko, K. J. Salimian, M. U. Salman, N. Sekulic, A. Alonso, P. E. Warburton, B. E. Black, The octamer is the major form of CENP-A nucleosomes at human centromeres. Nat. Struct. Mol. Biol. 20, 687–695 (2013).

91. M. Dumont, R. Gamba, P. Gestraud, S. Klaasen, J. T. Worrall, S. G. De Vries, V. Boudreau, C. Salinas-Luypaert, P. S. Maddox, S. M. Lens, G. J. Kops, S. E. McClelland, K. H. Miga, D. Fachinetti, Human chromosome-specific aneuploidy is influenced by DNA-dependent centromeric features. EMBO J. 39, e102924 (2020).

92. S. J. Falk, L. Y. Guo, N. Sekulic, E. M. Smoak, T. Mani, G. A. Logsdon, K. Gupta, L. E. T. Jansen, G. D. Van Duyne, S. A. Vinogradov, M. A. Lampson, B. E. Black, CENP-C reshapes and stabilizes CENP-A nucleosomes at the centromere. Science. 348, 699–703 (2015).

93. M. E. Kuo, L. L. Sullivan, K. Chew, B. A. Sullivan, Genomic variation within alpha satellite DNA influences centromere location on human chromosomes with metastable epialleles. Genome (2016) (available at http://genome.cshlp.org/content/26/10/1301.short).

94. M. E. Aldrup-MacDonald, M. E. Kuo, L. L. Sullivan, K. Chew, B. A. Sullivan, Genomic variation within alpha satellite DNA influences centromere location on human chromosomes with metastable epialleles. Genome Res. 26, 1301–1311 (2016).

95. H. S. Malik, S. Henikoff, Adaptive evolution of Cid, a centromere-specific histone in Drosophila. Genetics. 157, 1293–1298 (2001).

96. W. R. Rice, A Game of Thrones at Human Centromeres II. A new molecular/evolutionary model. Cold Spring Harbor Laboratory (2019), p. 731471.

97. L. Chmátal, S. I. Gabriel, G. P. Mitsainas, J. Martínez-Vargas, J. Ventura, J. B. Searle, R. M. Schultz, M. A. Lampson, Centromere Strength Provides the Cell Biological Basis for Meiotic Drive and Karyotype Evolution in Mice. Curr. Biol. 24, 2295–2300 (2014).

98. L. Fishman, J. K. Kelly, Centromere-associated meiotic drive and female fitness variation in Mimulus. Evolution. 69 (2015), pp. 1208–1218.

99. L. E. Kursel, H. S. Malik, The cellular mechanisms and consequences of centromere drive. Curr. Opin. Cell Biol. 52, 58–65 (2018).

100. P. J. Park, ChIP–seq: advantages and challenges of a maturing technology. Nat. Rev. Genet. 10, 669–680 (2009).

101. N. Altemose, A. Maslan, O. K. Smith, K. Sundararajan, R. R. Brown, A. M. Detweiler, N. Neff, K. H. Miga, A. F. Straight, A. Streets, DiMeLo-seq: a long-read, single-molecule method for mapping protein-DNA interactions genome-wide. bioRxiv 2021.07.06.451383 (2021), doi:10.1101/2021.07.06.451383.

102. B. J. Raney, T. R. Dreszer, G. P. Barber, H. Clawson, P. A. Fujita, T. Wang, N. Nguyen, B. Paten, A. S. Zweig, D. Karolchik, W. J. Kent, Track data hubs enable visualization of user-defined genome-wide annotations on the UCSC Genome Browser. Bioinformatics. 30, 1003–1005 (2014).

103. W. J. Kent, C. W. Sugnet, T. S. Furey, K. M. Roskin, T. H. Pringle, A. M. Zahler, D. Haussler, The human genome browser at UCSC. Genome Res. 12, 996–1006 (2002).

