## Supplementary figures and images for "Complete genomic and epigenetic maps of human centromeres"

### High Resolution cenHap pdf

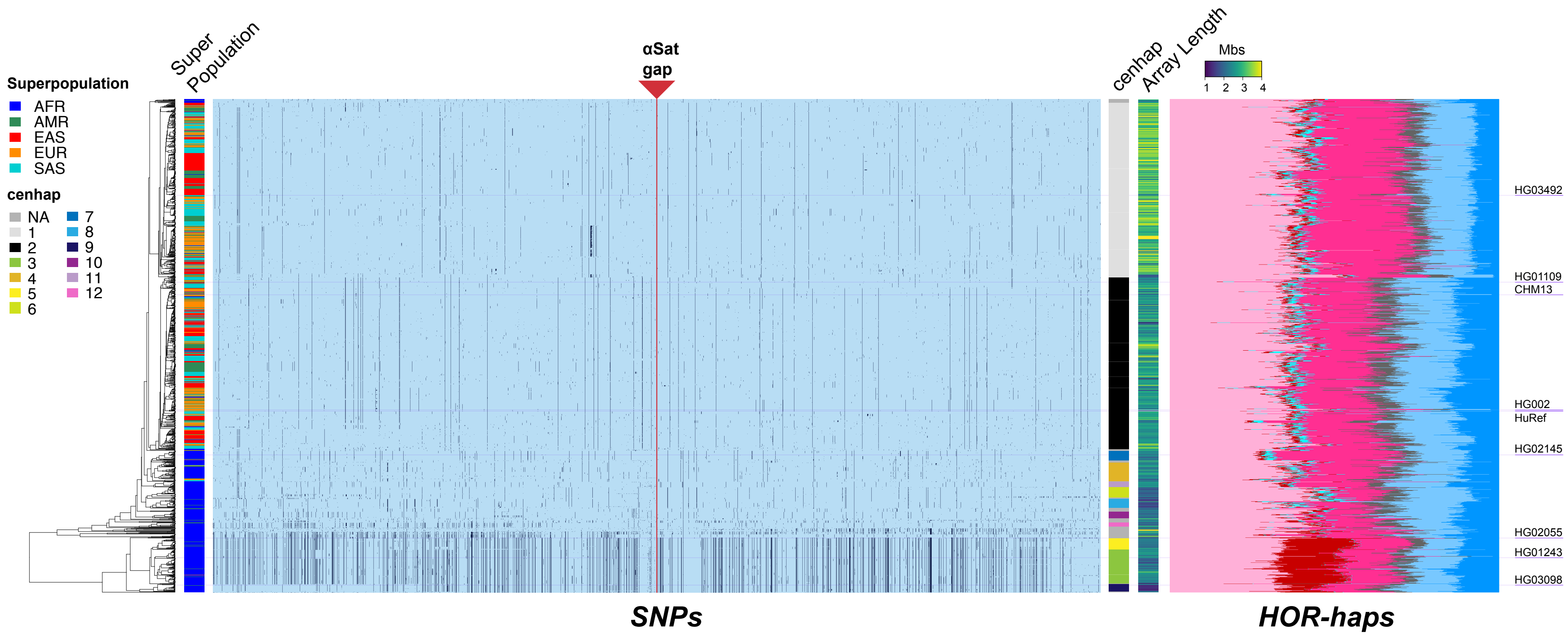
