## Supplemental Online Material and Methods for "Complete genomic and epigenetic maps of human centromeres"

This PDF file includes:
Materials and Methods **Section I to Section 8**
Figs. **S1 to S22**
Captions for Tables **S1 to S19**
**References**

Other supplementary material for this manuscript includes:
Tables **S1, S3, S5, S6, S8, S10, S11, S12, S15, S17, S18, S19**
High-resolution cenhap image (pdf)

### Materials and Methods

**Section I. Sequence content and organization of centromeric regions** **3**
Overview of cenSat annotation track. 3
Classification of alpha satellite monomers into suprachromosomal families 6
Classification of alpha satellite monomers into HOR monomers 14
Comparison against GRCh38 reference models 16
Annotation of Human Satellite 2 and 3 16
Semi-manual HSat2,3 subfamily assignment 16
Description of HSat1 annotation 17
Beta (β-) and gamma satellite consensus sequence generation 17
**Section 2. Centromeric genes and their expression** **17**
Gene expression quantification 18
**Section 3: Chromosome-specific Array Comparison** **18**
CHM13 chromosome isolation and sorting. 18
Resolving the shared ASat arrays S2C13/21H1L, S2C14/22H1L and S1C1/5/19H1L 20
cen13 - cen21 20
cen14 - cen22 23
cen1 - cen5 - cen19 24
Flow Sorted Chromosome Comparison with CHM13v1.0 assemblies 29
Monomer classification/identification was performed by HumAS-HMMER-HOR 29
Array specific k-mer comparison 29
**Section 4: Study of haplotype phased assemblies** **30**
Diversity Panel of HiFi Data used in HOR Structural Variant analysis 30
Detecting inversions, insertions, and deletions in hifiasm assemblies 30
**Section 5: The evolution of human centromere 6** **31**
Bird's eye view of the whole cen6 region 32
A close-up of the old centromere region in chr 6. 33
A close-up of the old centromere region in chr 2q (for comparison). 33
**Section 6: HOR unit and NRS prediction** **35**
HOR Structrual Variant (StV) prediction using CHM13 hor-monomer annotation 36
Automated annotation of "live" alpha satellite arrays. 37
HOR annotation of HiFi reads. 37
HOR decompositions of HiFi reads. 38
Calling repeat periodicity with NTRprism 39
HOR haplotype classification 40
Phylogenetic tree evaluation of evolutionarily young HOR-haps 41
**Section 7: CENP-A CUT&RUN and enrichment study** **44**
CUT&RUN Protocol for CHM13 44
Native CENP-A ChIP-seq analysis 45

Reference-free enrichment of k-mer protocols 46
Regional K-mers and Enrichment Estimates 46
Identifying false positive marker deserts due to multi-mapping 47
Marker assisted mapping protocol 47
**Section 8: Population study of haploid X chromosomes 49**
DXZ1 array characterization of HPRC haploid DXZ1 assemblies 49
Dot plots generated by an exhaustive search for exact matches between any two sequences 53
Prediction of recent repeat expansions using dot-plots 54
X Centromere HOR-hap proportion estimates in 1000 Genomes WGS Data and Epiallele ChIP-Seq & CUT&RUN datasets. 55
DXZ1 array size estimates in XY diversity panel 57
TandemTools evaluation of HPRC_PLUS genomes 58

**Supplementary Figures 59**

**Supplemental Tables and Table Legends 84**

**References 92**

### Section I. Sequence content and organization of centromeric regions

#### Overview of cenSat annotation track.

The primary focus of the cenSat annotation is to characterize the boundaries of satellite array elements across human centromeric regions. The broad definition of “centromeric regions” on each chromosome includes the satellite rich regions and 5 Mb of sequence on the p-arm and q-arm. This definition is supported by the observation (*1*) that pericentromeric segmental duplications are enriched in a span of 5 Mb on either side of the centromere-assigned gapped region. Notably, these genomic coordinates for centromeric regions in CHM13v.1.0 were also used in data analyses in Vollger et al. (*2*), and we prioritized consistency to ensure a shared definition of pericentromeric segmental duplications. Although we do not consider acrocentric short arms “centromeric” the vast majority of satellite DNAs present are highly enriched in peri/centromeric regions (e.g. HSat1-3, Beta satellites (βSat), Alpha satellites (αSat)). Therefore, acrocentromeric short arms are included in the censat annotation track in their entirety.

The cenSat annotation track is used to summarize across a wide range of detailed satellite annotation tools and results to provide discrete locations of the start and stop of each array. Notably, this annotation does not include peri/centromeric satellites that are outside of the boundaries of the centomeric regions, thereby omitting information in the arms. These data are reported by separate analyses (e.g. in Table S5). In this process, every base within a given centromeric region (as noted, including acrocentric short arms) is annotated into one of 10 non, overlapping categories based on the following criteria:

1) Alpha satellite Higher Order Repeat Array (**“hor”**): HOR-monomer and SF-monomer annotation tracks (as described in Table S3) for CHM13.v1.0 were used to identify location, and precise start and end of each array.

2) Alpha satellite divergent Higher Order Repeat Array (“**dhor**”): By default, sequences containing annotation for recent SF (ie. SF 1-3) were initially considered “hor”. If, on closer manual inspection, these regions did not contain ordered in a structured tandem repeat unit or represented mixtures of monomers we labeled regions as diverged, or dhor. These regions are typically found flanking either side of HOR arrays.

3) Alpha satellite monomeric (“**mon**”): monomeric regions were determined by the absence of data from the HOR-mon annotation track and evidence of older-SF classification using the SF-mon track.

4) Human Satellites 2 (“**hsat2**”): Classical human satellites II are often poorly characterized by repeatmasker libraries/strategies. Therefore we employed our own evaluation of these sequences within the CHM13 genome (as described below: Annotation of Human Satellite 2 and 3) and annotation of HSat2 subfamilies in characterizing each array. We listed multiple families (comma delimited) In cases where more than one HSat2 subfamily(*3*) was reported within the same array.

5) Human Satellites 3 (“**hsat3**”): Similarly, classical human satellite III is also poorly characterized by repeatmasker libraries/strategies. Once again, as performed for HSat2, we referenced our separate annotation of Human Satellite 2 and 3 to report the precise start and end of these arrays and subfamily assignments.

6) Human Satellites 1 (“**hsat1**”): HSat1 arrays were defined from the RepeatMasker (*4*)track, with all “SAR” annotations classified as HSat1A and “HSATI” annotations classified as HSat1B. Annotations of the same type and strand within 1 kb were merged into contiguous blocks.

7) Beta Satellites (**“bsat**”): Beta satellite arrays were determined from the RepeatMasker track and from data provided by repeatmodeler and composite satellite predictions (*5*). Annotations were labeled as bsat (if the array was ‘pure’ or contained primarily the canonical repeat), “LSAU-BSAT_Composite” if determined to be organized into a larger repeat structure with frequent LSAU embeds. In several locations we noted the components of the LSAU-BSAT_Composite repeat - minus beta satellite (and never with any other satellite family). In these cases we documented bsat(“LSAU-BSAT_Composite,no_beta”) to recognize

this shared repeat class.

8) Gamma satellites ("**gsat**"): Gamma satellite arrays were determined from the RepeatMasker track, where we noted any subfamily information provided (e.g. GSATII, GSATX). On several occasions we observed "GSATII/TAR1" and reported these events as one censat annotation. Additionally, we reported all occurrences of "GSATII,rnd-6_family-1431" or the presence of a novel repeat class (*5*) associated with four gsat locations in the CHM13 centromeric regions.

9) Peri/centromeric satellites ("**p-censat**"): We labeled all remaining, smaller satellite classes as p-censats, with description provided within parentheses. Sequence classes in these regions were determined from either RepeatMasker information (e.g. CER, SATR, and SST1), novel repeat modeler predictions and T2T repeat annotation (*5*)) (e.g. ACRO composite, and novelSatellites - or satellites often predicted from regions that lacked all repeatmasker annotation), and smaller satellite regions flagged by ULTRA tandem repeat software (*6*).

10) Stretches of non-satellite bases in the centromeric regions were labeled "**ct**", for centric transition regions. These regions are enriched with segmental duplications when found near and directly between satellite DNAs. We also used "ct" to denote the regions extending into the p and q arms (as noted with "p-arm" or "q-arm" in the annotation to distinguish from other ct regions which are more likely providing repeat based information).

11) The current cenSat annotation data is made available for CHM13v1.0. At this time the rDNA arrays on each acrocentric short arm had limited copy representation and presented the largest remaining gaps in the genome (*7*, *8*). The censat annotation notes each of the rDNA gapped regions (gap-rDNA) and also provides individual annotations for existing rDNA repeat units on the acrocentrics (as provided in (*8*)

Naming nomenclature for the censat annotation track provides a prefix for one of the 11 categories, an underscore followed by a chromosome id (chromosome _ ordering number, p-arm to q-arm) followed by any description of the repeat element in parentheses.

To note: the censat annotation aimed to describe entire satellite arrays. Therefore we did not break the array when it was interrupted by TE elements, rather we reported the boundaries of the entire array with TEs interspersed within. Further, we did not use this annotation to represent inversions within a given array and make no assumptions about strandedness.And in many cases the arrays are observed to shift between forward and reverse orientation within a given censat annotation (like beta satellite arrays on chromosome 1).

##### Classification of alpha satellite monomers into suprachromosomal families

SF classification into 20 SFs (Table S4) was initially created and trained on hg38 genome assembly and then applied to the T2T CHM13 v1.0 assembly without a change. It was performed as described earlier (*9*) except the HMMER platform (*10*, *11*) was used instead of PERCON. Annotation success in both hg38 and CHM13 was evident as all SF assignments provided by Shepelev et al., 2015 (*9*) PERCON annotation and the assignments made by phylogenetic analysis and manual mapping in Shepelev et al., 2009 (*12*) and Uralsky et al., 2019 (*13*) matched without exception. The bed files and the αSat SF annotation UCSC Browser tracks were prepared as previously described (*13*). Below we provide the details of the SF-HMMER tool construction.

Position one of the αSat monomer by tradition was set arbitrarily at BamHI cleavage site in DXZ1 X-specific HOR (*14*). The same standard matrices for SF-specific monomers as in Shepelev et al., 2015 (*9*)were used with two important additions. First, the HMMs for J3-J7 and D3-D9 and FD monomer classes characteristic of SF01 (formerly archaic SF1), SF02 (formerly archaic SF2) and SF2, respectively (Table S4), were added. Identification of sequences which belong to SF01 and 02 was described in Shepelev et al., 2015 (*9*)and Uralsky et al., 2019 (*13*). Monomeric types were identified using the haplotyping procedure described earlier (*9*, *13*). HMM profiles were formed by pooling all monomer sequences available for a certain monomeric type (as listed in Shepelev et al., 2015 (*9*) and Uralsky et al., 2019 (*13*)), HORs and dHORs were represented by consensus monomers.  Second, the HMM profiles for monomeric layers previously treated collectively as SF4+ (monomeric type M1+ (*9*)[Shepelev et al., 2015]) were distributed into 8 (Ga through Aa in Table S4) monomeric classes identified initially in chromosome X as reported in Shepelev et al., 2009(*12*) (Figure S2, Supplementary tables and Text S1) with slight modifications. Specifically, based on monomer haplotyping results, the Aa monomer class was split into two, Aa and more ancestral Ia which differed by just 3 positions. Also, the HMM for Ka monomer class was modified to include additional monomers from chromosomes 5 and 19 as the Ka layer in cenX was very small. To these SFs identified in prior studies the new ones for newly identified ancient layers Fa (SF15), Ea (SF16), Qa (SF17) and Pa & Ta (SF18 dimeric family) were added (Table S4). The new αSat SFs were identified as regions of incomplete or mosaic coverage by existing monomer types in hg38 assembly, which were extracted, aligned and haplotyped as described (*9*, *12*). SFs 15-18 were only represented by small pieces mostly located at sites far from existing centromeric arrays which possibly represent extinct ancient centromeres as suggested by their conserved position in primate lineages (see Table 1 and below). These locations were first identified in hg38 assembly and were verified for the T2T CHM13 v1.0 assembly (listed in Table S5). No new locations for the most ancient αSat SFs were found in CHM13. All monomers available for each novel SF in hg38 were extracted and used for HMMs. As the SF-specific monomeric classes represent ancestral layers of αSat sequences they should work in non-human primates as well as in the human genome (*12*)(Shepelev et al., 2009). Therefore a test run of the SF-classification tool was performed on available primate assemblies. As a result, a novel SF15 (monomeric type La) was identified and the respective HMM was created using Macaca mulatta sequences (chr15:110,729,891-110,731,376 and chr19:23,590,591-23,609,030 in rheMac3 assembly) and added to the set. Haplotyping procedure and the position on the phylogenetic tree of consensus monomers (see Figure 1 below) has put it between SF14 (Ia) and SF16 (Fa), as the Ia-specific mutations were already present and the Fa-specific mutations were yet absent. Locations of the most ancient αSat arrays in primates are detailed in Table 1 below. As expected, no arrays of La monomers were found in hg38 and CHM13 indicating deletion in human lineage. Such other deletions in primate lineages are obvious in Table 1.

Table 1. Location of ancient and relic centromeres in primates.

| Human orthologous site | chr12:107,851,722-107,857,592 | chr13:21,291,489-21,292,178 | chr13:24,519,087-24,528,185 | chr13:23,419,704-23,425,052 | chr3:125,655,489-125,670,097 | chr7:64,392,156-64,399,727 | chr22:22,133,960-22,139,864 | chr3:125,655,489-125,670,097 | chr9:38,471,199-38,671,199 (AS deleted)* | chr19:24,040,207-24,240,207 (AS deleted) | chr19:24,040,207-24,240,207 (AS deleted) | chrX:58,061,497-58,078,750 | chr1:103,860,487-103,874,558 | chr3:80,946,630-80,969,727 | chr4:12,274,746-12,290,926 | chr5:23,885,174-23,905,739 | chr12:10,763,306-10,863,306 (AS deleted) | chr16:35,651,906-35,706,487 | chr19:27,660,701-27,733,289 | chrX:58,145,309-58,186,125 |
|---|---|---|---|---|---|---|---|---|---|---|---|---|---|---|---|---|---|---|---|---|
| Chr. band | 12q23.3 | 13q12.11 | 13q12.12 | 13q12.12 | 3q21.2 | 7q11.21 | 22q11.22 | 3q21.2 | 9p13.1 | 19p12 | 19p12 | Xp11.1 | 1p21.1 | 3p12.2 | 4p15.33 | 5p14.2 | 12p13.2 | 16p11.1 | 19q11 | Xp11.1 |
| Species/ AS Monomer class | Pa & Ta | | | Qa | Ea | | | Fa | La | | Ia | | Aa | | | | | | | |
| Color | | | | | | | | | | | | | | | | | | | | |
| Marmoset | ▪ | | ▪ | ▪ | | | | | | | | | ▪ | ▪ | ▪ | ▪ | | | | |
| Macaca | ▪ | | ▪ | ▪ | | ▪ | ▪ | ▪ | ▪ | ▪ | ▪ | □ | ▪ | ▪ | ▪ | ▪ | ▪ | ▪ | | □ |
| Gibbon | ▪ | | ▪ | ▪ | ▪ | ▪ | | ▪ | | ▪ | | | ▪ | | ▪ | ▪ | | | | ▪ |
| Orangutan | ▪ | | ▪ | ▪ | ▪ | | ▪ | ▪ | ▪ | | | ▪ | ▪ | ▪ | ▪ | ▪ | | | ▪ | ▪ |
| Gorilla | ▪ | | ▪ | ▪ | ▪ | ▪ | ▪ | ▪ | | | | ▪ | ▪ | ▪ | ▪ | ▪ | | | | ▪ |
| Chimp | ▪ | ▪ | | ▪ | ▪ | ▪ | ▪ | ▪ | | | | ▪ | ▪ | ▪ | ▪ | ▪ | | ▪ | ▪ | ▪ |
| Human | ▪ | ▪ | ▪ | ▪ | ▪ | ▪ | ▪ | ▪ | | | | ▪ | ▪ | ▪ | ▪ | ▪ | | ▪ | ▪ | ▪ |
| SD nearby | − | + | + | − | + | + | + | + | + | + | + | + | + | − | − | − | + | + | + | + |
| Other Sat. nearby | − | − | − | − | + | + | + | + | + | + | + | + | − | − | − | − | − | + | + | + |

* The second half of the human orthologous site is on chr9:97,027,038-97,227,038 (AS deleted; 9q22.33)

Consensus sequences for all SF-monomeric classes were derived for hg38 using more than 50% majority rule initially. After CHM13 T2T v1.0 assembly was annotated by the SF-HMMER tool, the monomers according to monomeric classes were extracted, aligned, filtered to 150 bp length, and a set of CHM13-specific consensus (simple majority rule) was developed (shown below) for all monomeric layers. No significant differences, other than differences in polymorphic positions where of two major nucleotides one or the other could win by small margin, were found.

**Simple majority CHM13 T2T v1.0-based consensus sequences for SF-class monomers of monomeric SFs.** R1&R2, Ga and Ha were derived only from the regions annotated as “mon” in CenSat annotation, so that they do not cover the HORs which belong to respective SFs, to avoid the bias. Other monomeric classes which are not involved in known HORs were extracted genome-wide, as many of their locations are not covered with CenSat annotation. All monomers are aligned as a single file.

```
>R1_cons.11Jul21_CHM13
AATCT-GCAAGTGGATATTTG-GACTGCTTTGAGGCCTTCG-TTGGAAACGGGAATATCTTCACATAAAAACTAGACAGAA
GCATTCTCAGA-AACTTCTTTGTGATGTGTGCATTCAACTCACAGAGTTGAACCTTTCTTTTGATAGAGCAGTTTTGAAA
CACTCTTTTTGTAG
>R2_cons.11Jul21_CHM13
AATCT-GCAAGTGGATATTTG-GAGCGCTTTGAGGCCTATG-GTGGAAAAGGAAATATCTTCACATAAAAACTAGACAGAA
GCATTCTCAGA-AACTTCTTTGTGATGTGTGCATTCAACTCACAGAGTTGAACCTTTCTTTTGATAGAGCAGTTTTGAAA
CACTCTTTTTGTAG
>Ga_cons.9Jun21_CHM13
AATCT-GCAAGTGGATATTTG-GAGCGCTTTGAGGCCTATG-GTGGAAAAGGAAATATCTTCACATAAAAACTACACAGAA
GCATTCTGAGA-AACTTCTTTGTGATGTGTGCATTCATCTCACAGAGTTGAACCTTTCTTTTGATTGAGCAGTTTTGAAA
CACTCTTTTTGTAG
>Ha_cons.8Jun21_CHM13
AATCT-GCAAAGGGATATTTGTGAGCCCTTTGAGGCCTATG-GTGAAAAAGGAAATATCTTCACATAAAAACTAGACAGAA
GCTTTCTGAGA-AACTTCTTTGTGATGTGTGCATTCATCTCACAGAGTTGAACCTTTCTTTTGATTGAGCAGTTTGGAAA
CAGTCTTTTTGTAG
>Ka_cons.8Jun21_CHM13
AATCT-GCGAAGGGATATTTGGGAGCGCATTGAGGCCTATG-GTGAAAAAGGAAATATCTTCAGATAAAAACTAGAAAGA
AGCTTTCTGAGA-AACTGCTTTGTGATGTGTGCATTCATCTCACAGAGTTAAACCTTTCTTTTGATTGAGCAGTTTGGAA
ACACTGTTTTTGTAG
>Oa_cons.8Jun21_CHM13
ATTCT-GCGAATGGACATTTGGGAGCTCATTGAGGCCAATG-GCGAAAAAGTGAATATCCCAGGATAAAAACTAGAAGGA
AGCTATCTGAGA-AACCGCTTTGTGATGTGTGCATTCATCTCACAGAGTTAAACCTTTCTTTTCATTCAGCAGTTTGGAA
ACACTGTTTTTGTAG
>Na_cons.8Jun21_CHM13
AATCT-GCGAAGGGATATTTGGGAGCGCATTGAGGCCTATG-GTGAAAAAGGAAATATCTTCAGATAAAAACTAGAAAGA
AGCTTTCTGAGA-AACTGCTTTGTGATGTGTGCATTCATCTCACAGAGTTAAACCTTTCTTTGGATTCAGCAGTTTGGAA
ACACTGTTTTTGTCC
>Ca_cons.8Jun21_CHM13
AATCT-GTGAAGGGACATTTGGGAGCCCATTGAGGCCTATG-GTGAAAAACTGAATATCCCCAGATAAAAACTAGAAAGA
AGCTATCTGTGA-AACTGCTTTGTGATGTGTGGATTCATCTCACAGAGTTAAACCTTTCTTTTGATTCAGCAGGTTGGAA
ACACTCTTTTTGTAG
>Ba_cons.8Jun21_CHM13
AATCT-GCGAAGGGACATTTCAGAGCCCACTGAGGCCTATA-GTGAAAAACTGAATATCCCATGATAAAAACTAGAAAGA
AGCTATCTGTGA-AACTGCTTTGTGATGTGTGGATTCAGCTCACAGAGTTAAACCTTTCTTTTGATTCAGCAGGTTGGAA
ACACTCTTTTTGTAG
>Ja_cons.8Jun21_CHM13
AATCT-ACGAAGGGACATTTCAGAGCCCATTGAGGCCTATA-GTGAAAAACCGAATATCCCATGATAAAAACTAGAAACAA
GCTATCTGTGA-AAATGCTTTGTGATGTGTGGATTCATCTCACAGAGTTAAACCTTTGTTTTGATTCAGCAGGTTGGAAA
CACTCTTTTTGTAG
```

>Aa_cons.8Jun21_CHM13
AATCT-ACGAAGGGACATTTCTGAGCCCATTGAGGCCTATA-GTGAAAAACCGAATATCCCGCGATAAAAACTAGAAACA
AGCTATCTGTGA-AAATGCTTTGTGATGTGTGGATTCATCTCACAGAATGGAACCTGTGTTTTGATTCACCAGGTTGGAA
ACACTCTTTTTGTAG
>Ia_cons.8Jun21_CHM13
AATCT-ACGAAGGGACATTTCTGAGCCCATTGAGGCCTATA-AGGAAAAACCGAATATCCAGCGATAAAAACTAGAAACA
AGCTATCTGTGA-AAATGCTTTGTGATGTGTGGATTCATCTCACAGAATGGAACCTGTGTTTTGATTCAGCAGGTTGGAA
ACACTCTTTTTGTAG
>La_cons_absent_in_humans
AATCT-ACNAAGTGACATTTCTGAGCCCATTGAGGCCTATA-AGNAAAAATNGAATATCCAGCNATAAAAACTAGAAACAA
GCTATNTGTGA-AAATGCTTTGTGATGTGCTGTTTCATATCACAGAATGGAACCTGTGTTTTGATTCACCAGGTTCCAAA
CACTCTTTTTGTAG
>Fa_cons.8Jun21_CHM13
AATCT-ATGAAGTGACATTTCTGAGCCCATTGAGGCCTATT-AGGAAAAAATGAATATCCAGCCCTAAAAACTAGAAACAA
GCTATCTGTGA-AAATGCTTTGTGATGTGCTGTTTCATATCACAGAATGGAACCTGTGTTTTGATTCACCAGGTTCCAAA
CACTCTTTTTGTAG
>Ea_cons.8Jun21_CHM13
AATCT-AAGAAGTGACATTTCTGAGCCTATTGAGCCCTTAT-AGGAACATACGAATATCCAGCCCTAAAAACTAGAAACAA
GCTATCTGTGA-AAATGCTTTGTGATGTGCTGTTTTATATCACAGAATGGAACCTGTGTTTTGATTCAACAGGTTCCAAAC
ACTCTTTTTGTAG
>Qa_cons.8Jun21_CHM13
AATCT-AAGGAATGACATTTCAGAGGCTATT-AGGCCTATATATGAACATACGAATATCCAGCCCTAATAACTAAAAACAAG
CTATGTGCGA-AGAAGCTTTGTGATGTGCTGTTTTATATCACTGAATGGAACCTGTGTTTTCATTAAACAAGTTCCAAACA
CACGTTTTGTAG
>Pa_cons.8Jun21_CHM13
AAGCTCAGAAAAAAACATTTCAAAGCCCAGTTAGTGCTTCC-AGGAAGATATGCATATACAGCCCTAAATGCTAAAAGCA
AGATATGGCACACAAATAATGTATGATGTGCTGTTTTCTTTCACTGAATTCAACATGTATTAACTAAGAGCAGTTTTCAAA
GACACATCTTGTCA
>Ta_cons.8Jun21_CHM13
AATCT-CAGAAAACACAATTCTGAGCATAGT-AGGCCCACA-CAGAAAATGCGAATATCTAGCCCTAAAAGGTAAACGCAA
GCTTTGTGCCA-AAAACCTAGATGATGTGCTGTTTTCTTTCACTGAATTCAACATGTTTTGACATTGATCACTTTTCAAAA
ACACGTTTTGTAC

**Consensus monomers for the “new” SFs1-3 (including 01 and 02) generated in hg38 (50% rule).** In these SFs, which do not have a monomeric component and are entirely HOR-based, a different way of consensus derivation had to be employed as described in Shepelev et al., 2015 and Uralsky et al., 2019 (*13*). Consensus sequences for all known constituent HORs were derived first, and partitioned to monomers. Monomers were HMMER-classified into SF-classes and additionally haplotyped (checked for the presence of SF-specific mutations). Then consensus of consensus sequences was derived for each class. All monomers were aligned as a single file with consensus monomers for monomeric SFs provided above.

>J1_cons.hg38
AATTT-GCAAGTGGAGATTTC-AAGCGCTTTGAGGCCAATG-GTAGAAAAGGAAATATCTTCGTATAAAAACTAGACAGAA
TCATTCTCAGA-AACTACTTTGTGATGTGTGCGTTCAACTCACAGAGTTTAACCTTTCTTTTCATAGAGCAGTTTGGAAA
CACTCTGTTTGTAA
>J2_cons.hg38
AGTCT-GCAAGTGGATATTTG-GACCTCTTTGAGGCCTTCG-TTGGAAACGGG-ATTTCTTCATATAA-TGCTAGACAGAAG
AATTCTCAGT-AACTTCTTTGTGTTGTGTGTATTCAACTCACAGAGTTGAACCTTCCTTTAGACAGAGCAGATTTGAAAC
ACTCTTTTTGTGG

>J3_cons.hg38
AATTT-CCAAGTGGATATTTA-GAGCGCTTTGAGGCCTATG-GTAGAAAAGGAAATATCTTCATATAAAAACTAGACAGAAT
CATTCTCAGA-AACTACTTTGTGATGTGTGCGTTCAACTCACAGAGTTTAACCTTTCTTTTGATAGAGCAGTTTTGAAACA
CTCTTTTTGTAG
>J4_cons.hg38
AATCT-GCAAGTGGATATTTG-GACCTCTTTGAGGCCTTCG-TTGGAAACGGGTATTTCTTCATAATAAAACTAGACAGAA
GAATTCTCAGA-AACTTCTTTGTGATGTGTGCATTCAACTCACAGAGTTGAACCTTCCTTTTGATAGAGCAGTTTTGAAA
CACTCTTTTTGTAG
>J5_cons.hg38
AATTT-GCAAGTGTATATTTA-GAGCGCTTTGAGGCCTATG-GTAGAAAAGGAAATATCTTCCCATAAAACCTAGACAGAAG
CATTCTCAGA-AACTACTTTGTGATGTTTGCATTCAACTCACAGAGTTGAACATTCCTCTTGATAGAGCAGTTTTGAAACA
CTCTTTTTGTAG
>J6_cons.hg38
AGTCT-GCATGTGGATATTTT-GAGCGCTTTGAGGTATTCT-TTGGAAACGGGAATGTCTTCACATAAAAGGTAGACAGAA
GTGTTCTCAGA-AACTTCTTTGTGATGTCTGTGTTCAACTCACAGAGTTTAACCTTTCTTTTGATAGAGCGGTTTTGTAAC
ACTCTTTTTGTAG
>D2_cons.hg38
TATCT-GGAAGTGGACATTTG-GAGCGCTTTCAGGCCTATG-GTGAAAAAGGAAATATCTTCCCATAAAAACTAGACAGAA
GCATTCTCAGA-AACTTGTTTGTGATGTGTGTACTCAACTAACAGAGTTGAACCTTTCTTTTGATAGAGCAGTTTTGAAA
CACTCTTTTTGTAG
>D1_cons.hg38
AATCT-GCAAGTGGATATTTG-GATAGCTTTGAGGATTTCG-TTGGAAACGGGAATATCTTCATATAAAATCTAGACAGAAG
CATTCTCAGA-AACTTCTTTGTGATGTTTGCATTCAAGTCACAGAGTTGAACATTCCCTTTCATAGAGCAGGTTTGAAAC
ACTCTTTTTGTAG
>FD_D1D2_hybrid_cons.hg38
AATCT-GCAAGTGGATATTTG-GATAGCTTTAAGGATTTCG-TTGGAAACGGGAATATCTTCATGTAAAATCTAGACAGAAG
CATTCTCAGA-AACTTCTTTGTGATGTGTGTCCTCAACTAACAGAGTTCAACCTTTCTTTTGATACAGCAGTTTGGAAAC
ACTCTTTTTGTAG
>D3_2GA_cons.hg38
AGTCT-GCAAGTGGATATTTG-GATAGCTTTGAGGATTTCG-TTGGAAACGGGAATATCTTCACATAAAAACTAGACAGAA
GCATTCTCAGA-AACTTCTTTGTGATGTTTGCATTCAACTCACAGAGTTGAACATTCCCTTTCATAGAGCAGTTTTGAAA
CACTCTTTTTGTAG
>D4_cons.hg38
TATCT-GGAAGTGGACATTTG-GAGCGCTTTGAGGCCTATG-GTGAAAAAGGAAATATCTTCACATAAAAACTAGACAGAA
GCATTCTCAGA-AACTTCTTTGTGATGTGTGTACTCAACTCACAGAGTTGAACCTTTCTTTTGATACAGCAGTTTTGAAA
CACTCTTTTTGTAG
>D5_169TA_cons.hg38
AATCT-GCAAGTGGATATTTG-GATAGCTTTGAGGCTTTCG-TTGGAAACGGGAATATCTTCACATAAAAACTAGACAGAA
GCATTCTCAGA-AACTTCTTTGTGATGCTTGCATTCAACTCACAGAGTTGAACATTCCTTTTCATAGAGCAGTTTTGAAA
CACTCTTTTTGTAG
>D6_cons.hg38
AATCT-GCAAGTGGACATTTG-GAGCGATTTGAGACCTATG-GTGAAAAAGGAAATATCTTCACATAAAAAGTAGACAGAA
GCATTCTCAGA-AACTACTTTGTGATGTGTGTACTCAACTCACAGAGTTAAACCTTTCCTTTGATACAGCAGTTTTGAAA
CACTCTTCTTGTAG
>D7_cons.hg38
AATTT-GCAAGTGGATATTTG-GACAGCTTTGAGGCCTTCG-CTGGAAACGGGTATATCTTCACATAAAAACTAGACAGAA
GCATTCTCAGA-AACTTCTTTGTGATGCTTGCATTCAACTCACAGAGTTGAACATTCCTTTTCATAGAGCAGTTTTGAAA
CACTCTTTTTGTAG
>D8_cons.hg38
AATCT-GTAAGTGGAAACTTG-GAGCGCTTTGAGGCCTATG-GTGAAAAAGGAAATATCTTCCCATAAAAACTAGACAGAA
GAATTCTCAGA-AACTTCTTTGTGATGTGTGTACTCAACTCACAGAGTTGAACTTTTCTTTTGATAGAGCAGTTTTGAAAC
ACTCTTTTTGTAG
>D9_cons.hg38
AATTT-ACAAGTGGATATTAG-GACAGCATTGAGGATTTCG-TTGGAAACGGGAATATCTTCACATAA-AACTAGACAGAAG
CATTCTCAGA-AACTTCTTTGTGATGTGTGCATTCAACTCACAGAGTTGAAACTTTCTTTTGATAGAGCAGATTGGAAAC
ACTCTTTTTGTAG

```
>W1_cons.hg38
AATCT-GCAAGTGGATATTTG-GACCTCTCTGAGGATTTCG-TTGGAAACGGGATAAACTTCACATAA---CTAAACAGAAG
CATTCTCAGA-AACTTCTTTGTGATGTTTGCATTCAACTCACAGAGTTGAACCTTCCTTT-GATAGTTCAGGTTTGAAACA
CTCTTTTTGTAG
>W2_cons.hg38
AATCT-GCAAGTGGATATTTG-GACCACTTTGTGGCCTTCG-TTCGAAACGGGTATATCTTCACATCAAACCTAGACAGAA
GCATTCTCAGA-ANGTTTTCTGTGATGACTGCATTCAACTCACAGAGTTGAACAATCCTNTTGATGGAGCAGTTTTGAAA
CTCTCTTTCTTTGG
>W3_cons.hg38
AATCT-GCAAGTGGATATGTG-GACCTCTTTGAAGATTTCG-TTGGAAACGGGATCATCTTCACATAAAAACTAAACAGAA
GCATTCTCAGA-AACTNCTTTGTGATGTTTGTGTTCAACTCCCAGAGTTGAACTTTCCTTTTGANAGAGCAGCTNTGAAA
CACTCTTTTTCTAG
>W4_cons.hg38
AATCT-GCAAGTGGACATTTG-GAGGGCTTTGAGGCCTGTG-GTGGAAAAGGAAATATCTTCACATAAAAACTAGATAGA
AGCATTCTCAGA-AACTACTTTGTGATGATTGCATTCAACTCACAGAGTTGAACATTCCTTTTGATAGAGCAGTTTGGAA
ACACTCTTTTTGTAG
>W5_cons.hg38
AATCT-GCAAGTGGAGATTTG-GACCGCTTTGAGGCCTANG-GTAGTAAAGGAAATAACTTCATATAAAAACTAGACAGAA
GCATTCTCAGA-AAATTCTTTGTGATGATTGAGTTTAACTCACAGAGCTGAACATTCCTTTNGATGGAGCAGTTTCNAAA
CACACTTTTTGTAG
```

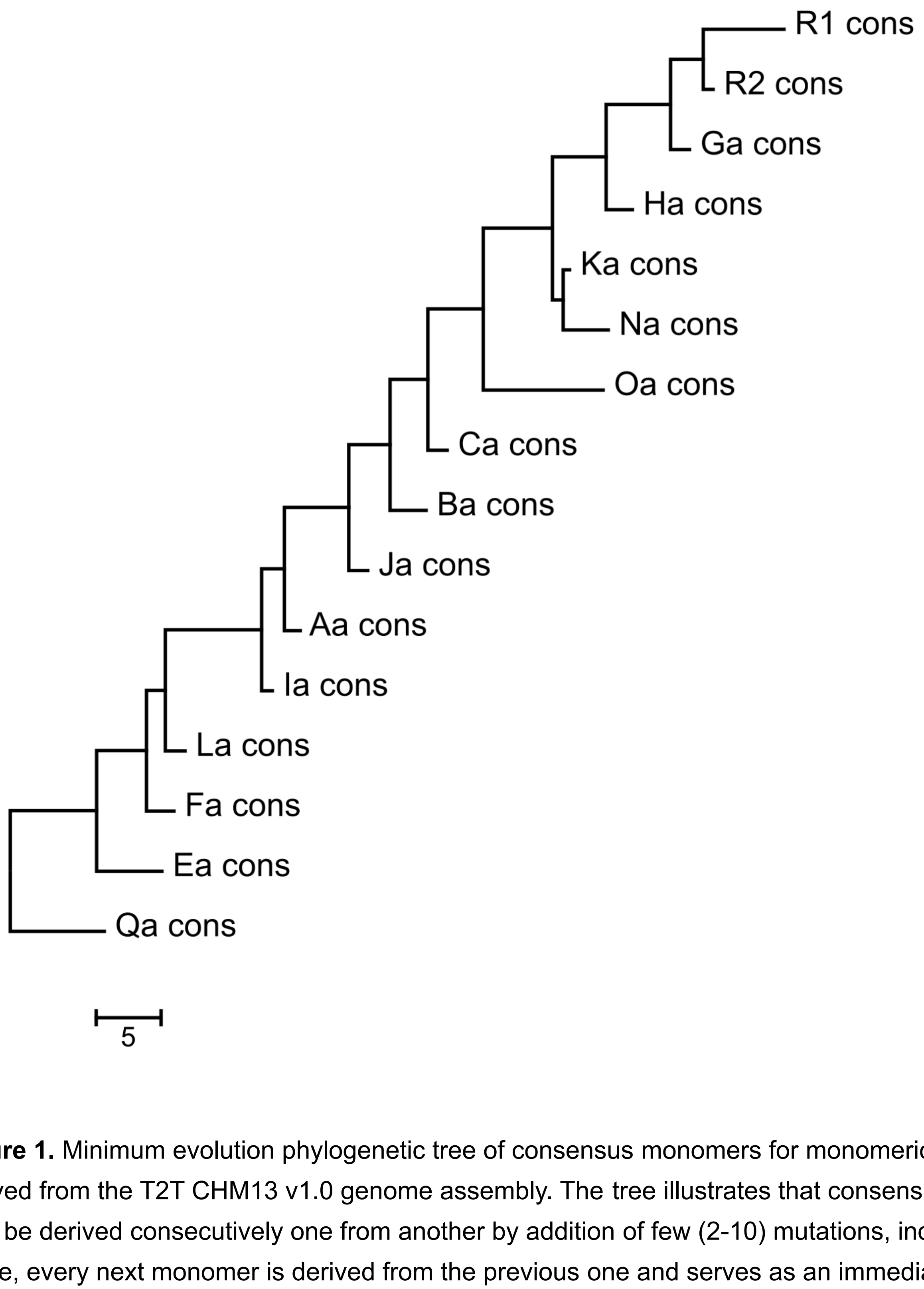

**Figure 1.** Minimum evolution phylogenetic tree of consensus monomers for monomeric layers derived from the T2T CHM13 v1.0 genome assembly. The tree illustrates that consensus sequences may be derived consecutively one from another by addition of few (2-10) mutations, indicating that, as a rule, every next monomer is derived from the previous one and serves as an immediate ancestor to the next one.

##### Classification of alpha satellite monomers into HOR monomers

Classification and annotation of αSat HOR monomers was performed as previously described in full detail in (*13*) for αSat of SF1. Remaining SFs which contain HORs (namely SFs 2, 3, 4, 5 and 6; (*9*) were processed in exactly the same way using hg38 genome assembly. The only change was the introduction of a larger set of competing SF-class HMMs. Specifically, the HMMs for all classes listed in Table S4, except for those that belong to SF01 and 02 were used. This change was introduced just for consistency and did not cause any significant difference in annotation results. All operations including construction of HMM profiles, use of HMMER platform and preparation of BED files and annotation tracks for UCSC Human Genome Browser were performed as described. Resulting annotation of hg38 HOR arrays was evaluated for consistency and accuracy as described (*13*). The complete list of HORs and their characteristics are shown in Table S3. Below we provide brief comments on assessment of this initial annotation which validated its further application to the newly obtained CHM13 T2T v1.0 complete assembly.

The annotation of AS HORs which we report here has been executed using a very simplistic approach and is essentially self explanatory. The correctness of annotation was self-evident as indicated by completeness of coverage, no significant cross-contamination of HOR arrays and numerical order (as opposed to chaos) in the HOR-track which meant that monomers in canonical HORs tended to go in correct numerical succession. These three features were percepted as evidence of good coverage. Only in cases of divergent HORs which form a tiny (about 3%) proportion of all SFs, reduced numerical order was often observed and there it could well be explained by degradation via deletions and other rearrangements. Specifically, the near complete coverage of SFs 1-3 (formed entirely by HORs) by HOR-specific HMM profiles was a demonstration of annotation success given that a satisfactory numerical order was achieved in the HOR reference models (RM) (*15*). The background of false hits outside HOR arrays in the track was uniformly low (see Discussion in Uralsky et al., 2019 (*13*)), but for isolated hits with the monomers of the SF5 divergent HORs (dHORs) which was an expected result of our low-threshold strategy and the high divergence characteristic of dHORs (*13*). It could be reduced by increasing our score to length ratio threshold (currently 0.7; see discussion in Uralsky et al., 2019 (*13*)) but only at the expense of reduced coverage in divergent HOR arrays. As of now, the coverage by the monomers of divergent HORs should only be considered genuine, if it is continuous. The ‘clouds’ of isolated and separated hits over SF5 arrays or the clouds of mosaic hits which belong to different dHORs are not considered to be the evidence of a dHOR array, only relatively continuous coverage by the same dHOR constitutes a dHOR array. This situation is no different from what was described in Uralsky et al., 2019 (*13*).

As a whole, our annotation have visualized the known and expected structure of AS arrays, including the large homogenous cores of the live centromeres, and a number of additional smaller homogeneous arrays percepted as inactive centromeric arrays (both represented mainly by RMs in hg38).

Thus, the initial survey of the HMMER-based AS HOR annotation of the hg38 human genome assembly has confirmed that the genome-wide annotation has worked the same way as SF1-only model annotation reported earlier (*13*). The principal features of this annotation were the monomer-by-monomer processing of the HORs and a very low identity threshold used. These features have allowed us to start addressing such previously unidentified problems as hybrid monomers and divergent HORs. We concluded that the manner of annotation we have chosen worked fine for the human genome assembly and could clearly be applied to the first complete T2T assembly of the human genome. It should be stressed that unlike the SF annotation, HOR annotation is only applicable to humans and would not work in the primate genomes which have different HORs (*16*).

Subsequent annotation of the T2T CHM13 v1.0 genome assembly demonstrated all the same validating features (as described in the main text). Just 3 HORs which were present in hg38 were not found in CHM13 (Table S3). These HORs represented by small RMs in hg38 are possibly the false HOR assignments which could be produced by combination of a single tandem duplication and an increased copy number of ASat arrays which are parts of segmental duplications. As discussed in Shepelev et al., 2015 (*9*), it is possible that the RM constructing protocol (*15*) might have assembled RMs using such sequences in a small number of cases. Otherwise, it could be a polymorphism in the human population. Examination of the T2T annotation track have yielded no evidence of any major HORs which are present in CHM13 and absent in hg38, however a large number of low copy number variant HOR arrays are evident on the fringes of larger HOR arrays as indicated by apparent slight contamination by other HORs or discontinuous coverage. These “edge” HORs need to be further studied and classified as novel sister HORs (*17*)(Miga and Alexandrov, 2021) or major haplotypes of the known HORs. In general, any mixed or non-specific HOR classification in CenSat annotation posted for a HOR region (e.g. hor_1_1(S1C1H2,S1C1H3) or hor_1_3(SF1)) is indicative of a presence of an edge HOR. Such low copy arrays are expected to represent the previous generations of the larger HORs and they bear valuable information on evolution stages of HOR development. However, their large number precludes manual analysis. Future development of automated methods of HOR decomposition and analysis (see section 5 – Automated annotation of “live” alpha satellite arrays) should be instrumental in completing this task.

##### Comparison against GRCh38 reference models

Fasta components (or unique, non-redundant alpha monomers) used to generate all centromere reference models in hg38 (*15*) were used to study the relationship with CHM13 (NCBI BioProject (http://www.ncbi.nlm.nih.gov/bioproject) under accession number PRJNA193213 and GenBank (http://www.ncbi.nlm.nih.gov/genbank/) under accession numbers GK000058 and GK000059). Sequences were mapped to the CHM13v1.0 assembly using bwa mem (*18*). The resulting sam files were processed using samtools (*19*) (converting sam to sorted bam) and bedtools (*20*) to obtain bedfiles with edit distance information in each alignment.

Additionally, we analyzed the SF-monomer classes across both CHM13 v1.0 and GRCh38 using methods described above ("Classification of alpha satellite monomers into HOR monomers").

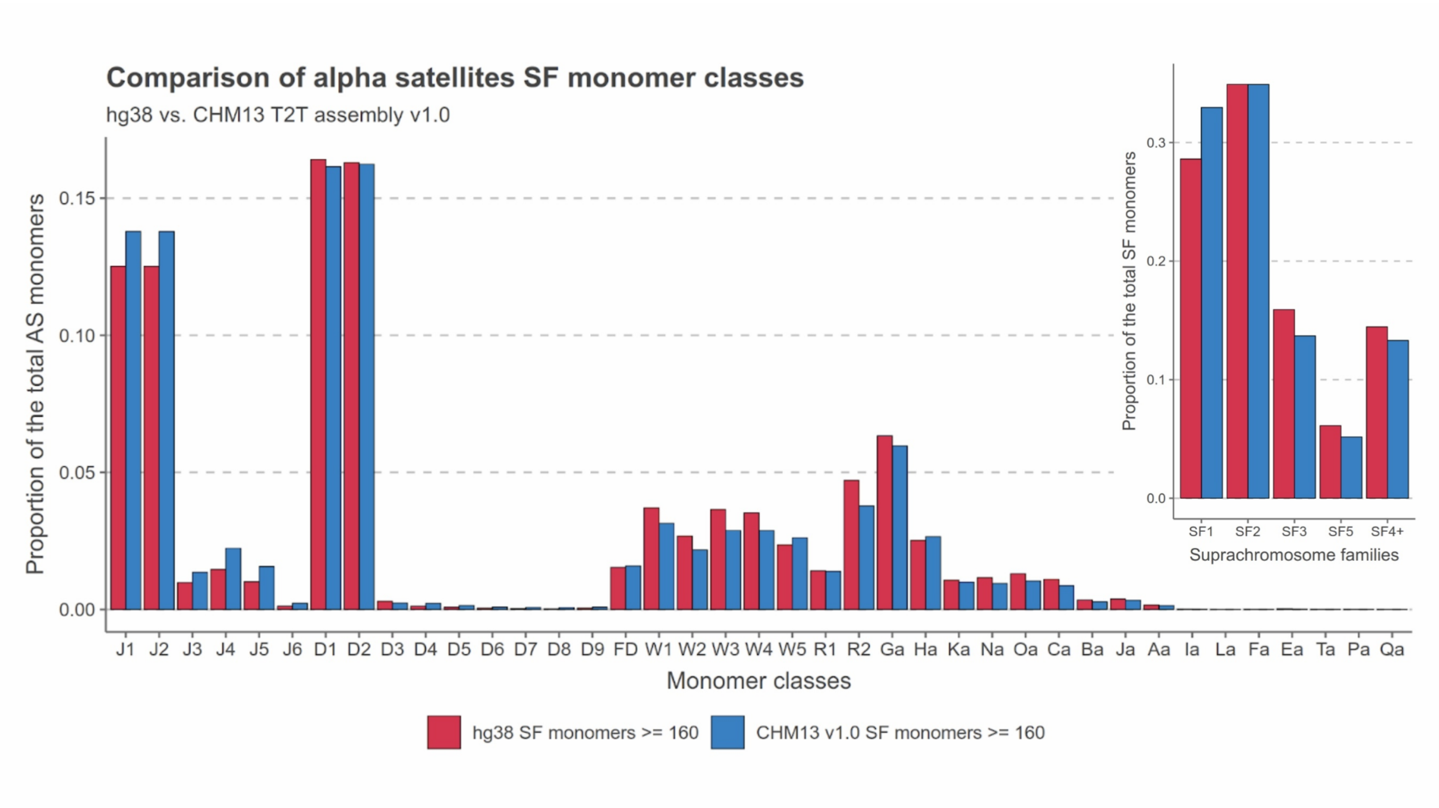

##### Annotation of Human Satellite 2 and 3

Human Satellites 2 and 3 were annotated on the T2T-CHM13 reference as follows. First, all HSat2 and HSat3 classified HuRef Sanger sequencing reads (identified in (*3*)) were obtained (*21*). All 24-mers were counted in each set of reads and ranked by abundance (counted only on the strand containing greater CATTC than GAATG). For HSat2, specific 24-mers were defined as those seen >10 times in HSat2 reads & <10 times in

HSat3 reads (10 times = 2 occurrences on average in genome given 5x HuRef coverage; proportion of HuRef read bases covered: 0.82 for HSat2 reads & 7.3e-05 for HSat3 reads). For HSat3, specific 24-mers were defined as those seen >10 times in HSat3 reads & <10 times in HSat2 reads (proportion of HuRef read bases covered: 0.62 for HSat3 reads & 6.3e-04 for HSat2 reads). Next, a 1-kb window was moved across the T2T-CHM13 assembly (step 1 kb) & each window was pre-screened by counting CATTC or GATTC occurrences. If >15 CATTC or >15 GAATG were counted, then that window would continue to be screened for 24-mers, with the CATTC-rich strand recorded as '+' (validation of pre-screening HuRef reads: captured 99.73% of HSat2 reads and 99.96% of HSat3 reads). If a 1-kb window passed the CATTC/GAATG threshold, HSat2 and HSat3 specific 24-mers were counted on the + strand. If >=100 24-mers were found matching HSat3, the window was classified as HSat3 (validation on HuRef reads: captured 99.7% of HSat3 reads, misclassified 0% of HSat2 reads as HSat3). Otherwise, if >=150 24-mers were found matching HSat2, the window was classified as HSat2 (validation on HuRef reads: captured 99.56% of HSat2 reads, misclassified 0.14% of HSat3 reads as HSat2). Adjacent windows with the same classification and strand were merged. The edges of contiguous blocks were refined by reporting first/last base matching an HSat2,3 specific 24-mer. When overlaps occurred (often in strand flip regions), the starts and ends were adjusted to the center of the overlap region. Any adjacent regions that were of the same type (HSat2 or HSat3) & strand and within 1000 bp were then merged.

##### Semi-manual HSat2,3 subfamily assignment

To assign HSat2,3 arrays (identified by the method above) in CHM13 to the 3 HSat2 subfamilies and 11 HSat3 subfamilies previously identified (*3*), we obtained HuRef reads assigned to each subfamily (*3*) and aligned them to the T2T-CHM13 reference using winnowmap (*22*), then applied a mapping quality filter of 10. In most cases only reads from one subfamily aligned with high coverage to each array. In cases where more than one subfamily aligned to the same array, they nearly always occupied different subregions within the array (Fig. S10A). In the CenSat annotation track, all subfamilies detected in a contiguous array are reported in parentheses, in order by size from largest to smallest. In the more detailed HSat2,3 subfamily assignment track shown in Fig. 2a-c, arrays were split at strand inversion breakpoints and at the borders between different subfamily assignments, and this more detailed subfamily assignment track was used to make Fig. 2e.

##### Description of HSat1 annotation

HSat1 regions and their strand orientation were defined from the RepeatMasker track, with all “SAR” annotations classified as HSat1A and “HSATI” annotations classified as HSat1B. Annotations of the same type and strand within 1 kb were merged into contiguous blocks.

##### Beta (β-) and gamma satellite consensus sequence generation

To generate the β-satellite consensus sequence, the CHM13v1.0 assembly was annotated with RepeatMasker using the new repeat library, and regions containing composite LSAU-BSAT repeats were extracted from the assembly with SAMtools v1.12 (*19*). The LSAU-BSAT-containing regions were assessed with StringDecomposer (*23*)(v1.0.0) to identify β-satellite monomers that had >70% sequence identity to the following β-satellite repeat:

‘GATCACGCAGGTGATGTAACTCTTCTCTATATCTGCCTACTGGTGGCATTGTGGCATATTTCTACACT’. 74,497 β-satellite monomers were identified, with an average length of 67 bp. These monomers were used to derive a consensus sequence.

To generate the -satellite consensus sequence, the CHM13v1.0 assembly was annotated with RepeatMasker using the new repeat library, and regions annotated as “GSAT” were extracted from the assembly with SAMtools v1.12 (*19*). GSAT regions were assessed with StringDecomposer (*23*)(v1.0.0) to identify -satellite monomers that had >75% sequence identity to the following -satellite repeat (DFAM model sequence DF0000148.4):
‘GCTGGGAGCCTCCCAAGGAGGCCTCTCCCATCCCAGAAGCCCCCAGGGCTGTCCCGGGCGGGCTGTAA
AGCCCCAGGCTTTGGAGCAGGGTGCCTGTGTCTCTCGCGGAAGGCCCCCACAAGCGAAAACGGGGCCG
CAGGGTGGCGTGGGCGGGCCGCAGGGACTCAGGGGGACGTTGAGGCAGGCAGAGGGGAGAAGCGGC
GAGACCGCAGGGAAT’. 904 -satellite monomers were identified, with an average length of 215 bp. These monomers were used to derive a consensus sequence

#### Section 2. Centromeric genes and their expression

CHM13 mRNA sequence data

mRNA was extracted from CHM13 cells in two replicates with an RNeasy Mini kit (Qiagen 74104) without DNase treatment. RNA integrity number (RIN) scores were determined to be at least 7.8 by an Agilent Bioanalyzer. PolyA-selected cDNA libraries were generated with the KAPA RNA HyperPrep (Roche KK8540) and sequenced with paired-end, 150-base pair reads on one lane of an Illumina NovaSeq S4 flowcell at the UC Davis DNA Technologies Core.

##### Gene expression quantification

Transcript abundance was estimated in transcripts per million (TPM) with Salmon v1.3.0(*24*), using the CHM13 transcriptome (CATv4 and liftOff, (*8*)) and the v1.0 assembly as a “decoy” sequence to account for reads mapping to unannotated sequences. The “--gcBias” option was enabled. TPM values were aggregated to the gene level using the R package tximport(*25*).

Additionally, RNA-seq reads were trimmed with CutAdapt using a minimum read length of 100 nucleotides and aligned to the CHM13 assembly using bowtie2 with “-k 100.” Alignments were intersected with single-copy 21mer and 51mer markers as described:
https://gitlab.com/SJHoyt/t2t_transposable-elements/-/tree/main/CutnRun_analyses

Read counts were produced for each gene using htseq-count v0.12.3(*26*) with the following parameters: “-t exon --nonunique none -m intersection-nonempty”. Subsequently, TPM values were calculated for each gene using the mean exonic length of all transcripts, as determined by GTFtools (*27*).

Finally, Iso-Seq alignments (N=2 replicates) were obtained from (*28*) and intersected with single-copy 51mer markers. Counts were generated at the gene level, and TPM values were calculated using the total gene length.

Both replicates were combined for all analyses due to high reproducibility (Pearson's *r*>0.95)

#### Section 3: Chromosome-specific Array Comparison

##### CHM13 chromosome isolation and sorting.

To induce mitotic arrest, cells were treated with 100µM Monastrol (Tocris) and 10µM pro-Tame (R&D Systems) for 10-13 hours. Mitotic cells were collected by shake-off. Collected cells were treated with 0.1µg/ml Colcemid for 15 min. Then cells were incubated in hypotonic swelling buffer containing 45mM KCl, 10 mM MgSO4, 0.5mM Spermidine and 0.2mM Spermine for 12 min and collected by centrifugation at 335g for 5min. Chromosomes were isolated using polyamine method (*29*) Briefly, cells were lysed in polyamine buffer (15mM Tris pH 7.5, 80 mM KCl, 2 mM EDTA, 0.5 mM EGTA, 3 mM DTT, 0.25% Triton X-100, 0.2 mM spermine, 0.5 mM spermidine) on ice for 30 minutes. Chromosomes were labeled for 2 hours with 50 µg/ml Chromomycin A3 and 5 µg/ml Hoechst 33258 in the presence of 25 mM sodium sulfite, 10 mM sodium citrate and 10 mM MgSO4. Before sorting, chromosome suspension was filtered through the 8 µm strainer. Chromosomes were sorted on a MoFlo Legacy (Beckman Coulter) at 60-65 psi using 70 µm tip. Hoechst 33258 was excited with UV laser (330-360nm) and the emission was detected in a 409-480 nm range. Chromomycin A3 was excited with the second laser tuned to 458 nm and emission was detected in a 505-605 nm range. Clumps and debris were excluded by gating on forward scatter versus pulse width. Chromosomes were sorted into DNA LoBind collection tubes (Eppendorf). Approximately 37000-123000 sorted events were collected per chromosome.
Ata analysis was performed using FlowJo ( BD Life Sciences).

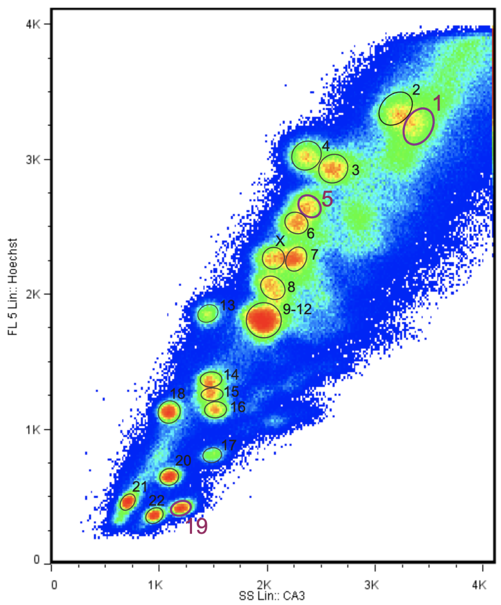

**Figure legend:** Bivariate flow karyotype of chromosomes from CHM13 cell line. Isolated chromosomes were stained with Chromomycin A3 and Hoechst 33258 in polyamine – containing buffer. Hoechst 33258 has propensity to AT bases and Chromomycin A3 has propensity to GC bases, which allows resolving double - stained chromosomes according to their basepair ratio and overall DNA content. Dashed circles denote individual chromosomal populations, solid circles indicate sorted populations.

| Sample name | Number of Chr | Chr size( Mb) | Max amount DNA (ng) | Recovered amount DNA (ng) |
|---|---|---|---|---|
| Chr1 | 51330 | 250 | 14 | 2.25 |
| Chr4 | 55751 | 190 | 12 | 2.53 |
| Chr5 | 59055 | 175 | 11.3 | 3.73 |
| Chr19 | 180,805 | 55 | 10.9 | 0.17 |

Genomic DNA was extracted from sorted chromosomes using the MagAttract HMW DNA Kit (Qiagen). DNA was analyzed and quantified using a FemptoPulse (Agilent). Whole chromosome libraries were prepared from one nanogram DNA using the Accel-NGS 2S DNA Library Kit (Swift Biosciences) with unique dual index adapters according to manufacturer's protocol with the substitution of Kapa HiFi HotStart Ready Mix (Roche) for library amplification. Twelve cycles of PCR were used for samples with 2 ng DNA input and 15 cycles were used for the sample with 0.17 ng input DNA. The libraries were pooled for sequencing on a HiSeq 2500 DNA sequencer in Rapid mode (Illumina, Inc.), generating 336 M paired-end 151 base reads. The raw data was processed using RTA1.18.64 and bwa0.7.12.

#### Resolving the shared ASat arrays S2C13/21H1L, S2C14/22H1L and S1C1/5/19H1L

##### cen13 - cen21

According to the structural variant (StV) track statistics, the most common StVs for cen13 and 21 were S2C13/21H1L_13.1-11, S2C13/21H1L_13.1-7 and S2C13/21H1L_21.1-11. These complete sequences were extracted and aligned via MUSCLE and multiple alignments were used as HMMs with HMMER platform (*10*, *30*) using just these profiles as described for HUMAS-HMMER-HOR (*13*). Chr13 and chr21 assemblies from CHM13 T2T v1.0 were processed and the output file was converted into a bed file and low-score and overlapping hits were filtered as described. Bed files were opened in UCSC Browser (*31*, *32*).

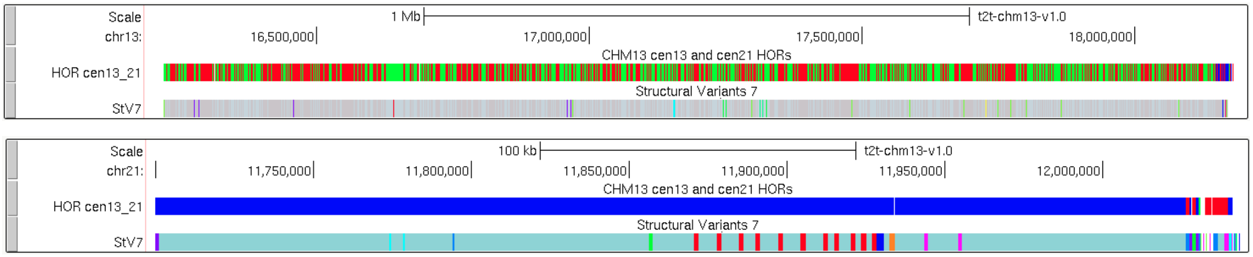

S2C21H1L.1-11 is blue
S2C13H1L.1-7 is green
S2C13H1L.1-11 is red
Cen13 and cen21 HORs were discriminated well except for the right flank on both cen13 and cen21 where the arrays share a collection of identical features including the small sister HOR array of S2C13H1-B (lightly

colored in the close-up below). The general collinearity of the short arms sequences in chr 13 and 21 (*8*) suggests that they basically derive from one chromosome which was duplicated and got two different sets of the long arms. Given that, it seems likely that the right flank of the centromere represents the remnant of the ancestral centromeric array in which the current chromosome-specific HORs existed as HOR-haps (see the main text) along with a sister HOR S2C13H1-B. Later on, on two different chromosomes different HOR-haps got amplified and formed the bulks of the current arrays. The 7-mer StV represents the cen13 HOR-hap and it shares the cen13-specific mutations. Apparently, it has emerged and expanded only in this centromere after the separation.

cen13:

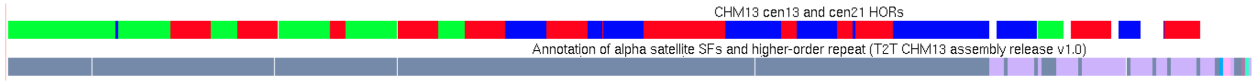

cen21:

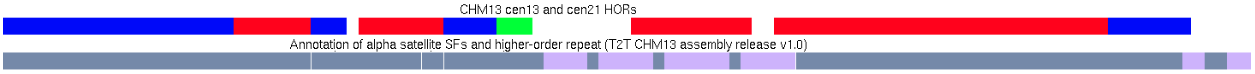

Consensus sequences for S2C13H1L_13 (11mer and 7mer) and S2C13H1L_21 (11mer) were derived and found to differ in 25 positions of the 11-mer HOR, which corresponds to a HOR-hap scale difference (~1%).

>CONS_S2C13/21H1L_21
TACTCCAGAACGAGTGTTTCAAACCTGCTCTATGAAAGGGAATCTTCAACTCTATGAGTT
GAATGCAGACATCAGAAAGAAATTTCTGAGAATGCTGCTGTCTACCTTTTATTTGAATTC
CCGCTTCCAACGAAATCCTCCAAGCTATCCAAATATCCACCTGCATTTTCCACAAAAAGA
GTGTTTCAAAACTG--CTCTATCAATAGAAATGTTCAACTCCTTTGGCTGGGTACACACA
TCACAAACAAGTTTCTGAGAATGCTTCTGTCTAGTTTTTATGGGAAGACGTTCCCTTTTT
CACCAAAGGCATCAAAGCGCTCCAAATGTCCACTTCCAGACACTACAAAAAGAGTGTTTC
AAACGTGCTCTAAGAAAGCGAATGTTCAACTCTGTGACTTGAATGCAGATATCACAAAGT
AGTTTCTGAGAGGGCTTCTGTCTAGATTTTAGATGATGATATTCCCGTTTCCAACGAAAT
CATTAGAGCTATCCAAATATCCACTTACAGTTTCTACAAAAAGAGTGTTTCCAAACTGCT
GCATCAAAAGAGAGGTTCCACTCTGTTAGCTGAGTACACACATCACAAACTTGTTTCTCA
GAATCCTTCTGTCTCGTTTTTATGGGAAGATATTTACTTTTTCACCGTAGGCATCAAAGC
GCTCCAAATGTCCACATCCAGATACTCCAGAAAGAGTGTTTCAAACCTGCTCTATGAAAG
GGAATCTTCAACTCTATGAGTTGAATGCAGACATCAGAAAGAAATTTCTGAGAATGCTGC
TGTCTACCTTTTATTTGAATTCCCGCTTCCAACGAAATCCTCCAAGCTATCCAAATATCC
ACTTGCAGATTCCACAAAAAGAGTGTTTCAAAACTGCTCTCTATCAATGGCAAAGTTCAA
CTCTGTTAGTTGAGGACACATATCACCAACAAGTTTCTGAGAATGCTTCTGTCTATTTTT
TATGGGAAGATATTTCCTTTTTCACCGTAGGCGTCAAGGCGATCGAAATGTCCACTTCCA
CAAACTACAAAAAGAGTGTTTCAAACCTGCTCTATGAAAGGCCATGTTCATCTCTATGAG
TTGAATGGAAATATCCGAAAGAAATTTCTGGGAATGCTGCTGTCTAGTGTTTATACGAAT
TCCCGCTTCCAACGAAATCCTCAAAGCAATCCAAATATCCACTTGCAGAATCCACAAAAA
GAGTGTTTCAAAACTGCTCTATCAATAGAAAGGTTCAACTCTTTTAGTTGAGTACACACA
TCACGAACAAGTTTCTGAGAATGCTTCTGTCTGGCTTTTATTGGAAGACGTTTCCTTTTC
ACCAAAGG---CATCAAAGCGCTCCAAATGTCCACTTCCAGATTCTTCCAAAAGAGTGTT
TCAAACGTGCTCAAAGTAAGGGAATGTTCAACTCTGTGACTTGAATGCAGATATCACCAA

GTAGTTTCTAATAGTGCTTCTGTCTAGATTTTAGATGATGATATTCCCGTTTCCAACGAA
ATCGTTAGAGCTATCCAAATATCCAGTTACAGTTTCTACCAAAAGGGTGTTTCCAAATTG
CTGCATCAAAAGAAAGGTTCAACTCTGTTAGTTGAGGACACACATCACAAAGAA-GTTTG
TGAGAATGCTTCTGTCTAGATTTTGTATGACGATATTCCCTTTTCCAACGATATCGTTAA
AGCAATCTAAATATCAATTTGCAGAATCCACAAAAATAGAGTTTCAAAGCTGCTCTGTAA
AAAGAAAGGTTCCACTCTGTTAGCTGAGTACACACATCACAAACTTGTTTCTGAGAATCC
TTCTGTCTCGTTTTTATGGGAAGATATTTACTTTTCCACCGTAGGCATCAAAGCGCTCCA
AATGTCCACATCCAGAT

>CONS_S2C13/21H1L_13 (11mer)
TACTCCAGAAAGAGTGTTTCAAACCTGCTCTATGAAAGGGAATCTTCAACTCTATGAGTT
GAATGCAGACATCAGAAAGAAATTTCTGAGAATGCTGCTGTCTACCTTTTATTTGAATTC
CCGCTTCCAACGAAATCCTCCAAGCTATCCAAATATCCACCTGCATTTTCCACAAAAAGA
GCGTTTCAAAACTGCTCTCTATCAATAGAAATGTTCAACTCCTTTGGCTGGGTACACACA
TCACAAACAAGTTTCTGAGAATGCTTCTGTCTAGTTTTTATGGGAAGACATTCCCTTTTT
CACCAAAGGCATCAAAGCGCTCCAAATGTCCACTTCCAGACACTACAAAAAGAGTGTTTC
CAACGTGCTCTAAGAAAGCGAATGTTCAACTCTGTGACTTGAATGCAGATATCACAAAGT
AGTTTCTGAGAGGGCTTCTGTCTAGATTTTAGATGATGATATTCCCGTTTCCAACGAAAT
CATTAGAGCTATCCAAATATCCACTTACAGTTTCTACAAAAAGAGTGTTTCCAAACTGCT
GCATCAAAAGAGAGGTTCCACTCTGTTAGCTGAGTACACACATCACAAACTTGTTTCTCA
GAATCCTTCTGTCTCGTTTTTCTGGGAAGATATTTACTTTTTCACCGTAGGCATCAAAGC
GCTCCAAATGTCCACATCCAGATACTCCAGAAAGAGTGTTTCAAACCTGCTCTATGAAAG
GGAATCTTCAACTCTATGAGTTGAATGCAGACATCAGAAAGAAATTTCTGAGAATGCTGC
TGTCTACCTTTTATTTGAATTCCCGCTTCCAACGAAATCCTCCAAGCTATCCAAATATCC
ACTTGCAGATTCCACAAAAAGAGTGTTTCAAAACTGCTCTCTATCAATGGCAAAGTTCAA
CTCTGTTAGTTGAGGACACATATCACCAACAAGTTTCTGAGAATGCTTCTGTCTATTTTT
TATGGGAAGATATTTCCTTTTTCACCGTAGGCGTCAAGGCGATCGAAATGTCCACTTCCA
CAAACTACAAAAAGAGTGTTTCAAACCTGCTCTATGAAAGGCCATGTTCATCTCTATGAG
TCGAATGGAAATATCCGAAAGAAATTTCTGGGAATGCTGCTGTCTAGTTTTTATACGAAT
TCCCGCTTCCAACGAAATCCTCAAAGCAATCCAAATATCCACTTGCAGAATCCACAAAAA
GAGTGTTTCAAAACTGCTCTATCAATAGAAAGGTTCAACTCTTTTAGTTGAGTACACACA
TCACAAACAAGTTTCTGAGAATGCTTCTGTCTGGCTTTTATTGGAAGACGTTTCCTTTTC
ACCAAAGGCATCATCAAAGCGCTCCAAATGTCCACTTCCAGATTCTTCCAAAAGAGTGTT
TGAAACGTGCTCAAAGTAAGGGAATGTTCAACTCTGTGACTTGAATGCAGATATCACCAA
GTAGTTTCTAATAGTGCTTCTGTCTAGATTTTAGATGATGATATTCCCGTTTCCAACGAA
ATCGTTAGAGCTATCCAAATATCCACTTACAGTTGCTACAAAAACAGTGTTTCCAAACTG
CTGCATCAAAAGAAAGGTTCAACTCTGTTAGTTGAGGACACACATCACAAAGAATGTTTG
TGAGAATGCTTCTGTCTAGATTTTGTATGACGATATTCCCTTTTCCAACGATATCATTAA
AGCAATCTAAATATCCATTTGCAGAATCCACAAAAATAGAGTTTCAAAGCTGCTCTGTAA
AAAGAAAGGTTCCACTCTGTTAGCTGAGTACACACATCACAAACTTGTTTCTCAGAATCC
TTCTGTCTCGTTTTTATGGGAAGATATTTACTTTTTCACCGTAGGCATCAAAGCGCTCCA
AATGTCCACATCCAGAT

>CONS_S2C13/21H1L_13 (7mer)

------------------------------------------------------------
------------------------------------------------------------
------------------------------------------------------------
------------------------------------------------------------
------------------------------------------------------------
------------------------------------------------------------
------------------------------------------------------------
------------------------------------------------------------
------------------------------------------------------------
------------------------------------------------------------
------------------------------------------------------------
----------------------TACTCCAGAAAGAGTGTTTCAAACCTGCTCTATGAAAG
GGAATCTTCAACTCTATGAGTTGAATGCAGACATCAGAAAGAAATTTCTGAGAATGCTGC
TGTCTACCTTTTATTTGAATTCCCGCTTCCAACGAAATCCTCCAAGCTATCCAAATATCC
ACTTGCAGATTCCACAAAAAGAGTGTTTCAAAACTGCTCTCTATCAATGGCAAAGTTCAA
CTCTGTTAGTTGAGGACACATATCACCAACAAGTTTCTGAGAATGCTTCTGTCTATTTTT
TATGGGAAGATATTTCCTTTTTCACCGTAGGCGTCAAGGCGATCGAAATGTCCACTTCCA
CAAACTACAAAAAGAGTGTTTCAAACCTGCTCTATGAAAGGCCATGTTCATCTCTATGAG
TCGAATGGAAATATCCGAAAGAAATTTCTGGGAATGCTGCTGTCTAGTTTTTATACGAAT
TCCCGCTTCCAACGAAATCCTCAAAGCAATCCAAATATCCACTTGCAGAATCCACAAAAA
GAGTGTTTCAAAACTGCTCTATCAATAGAAAGGTTCAACTCTTTTAGTTGAGTACACACA
TCACAAACAAGTTTCTGAGAATGCTTCTGTCTGGCTTTTATTGGAAGACGTTTCCTTTTC
ACCAAAGG---CATCAAAGCGCTCCAAATGTCCACTTCCAGATTCTTCCAAAAGAGTGTT
TGAAACGTGCTCAAAGTAAGGGAATGTTCAACTCTGTGACTTGAATGCAGATATCACCAA
GTAGTTTCTAATAGTGCTTCTGTCTAGATTTTAGATGATGATATTCCCGTTTCCAACGAA
ATCGTTAGAGCTATCCAAATATCCACTTACAGTTGCTACAAAAACAGTGTTTCCAAACTG
CTGCATCAAAAGAAAGGTTCAACTCTGTTAGTTGAGGACACACGTCACAAAGAA-GTTTG
TGAGAATGCTTCTGTCTAGATTTTGTATGACGATATTCCCTTTTCCAACGATATCATTAA
AGCAATCTAAATATCCATTTGCAGAATCCACAAAAATAGAGTTTCAAAGCTGCTCTGTAA
AAAGAAAGGTTCCACTCTGTTAGCTGAGTACACACATCACAAACTTGTTTCTCAGAATCC
TTCTGTCTCGTTTTTATGGGAAGATATTTACTTTTTCACCGTAGGCATCAAAGCGCTCCA
AATGTCCACATCCAGAT

#### cen14 - cen22

According to the StV track statistics, the most common StVs for cen14 and 22 were the full-size HORs S2C14/22H1L_14.1-8 and S2C14/22H1L_22.1-8. These sequences were extracted, aligned via MUSCLE and installed in HMMER and used to haplotype chr14 and chr22. Output file was converted into a bed file and low-score and overlapping hits were filtered out. Bed file was opened in UCSC Browser.

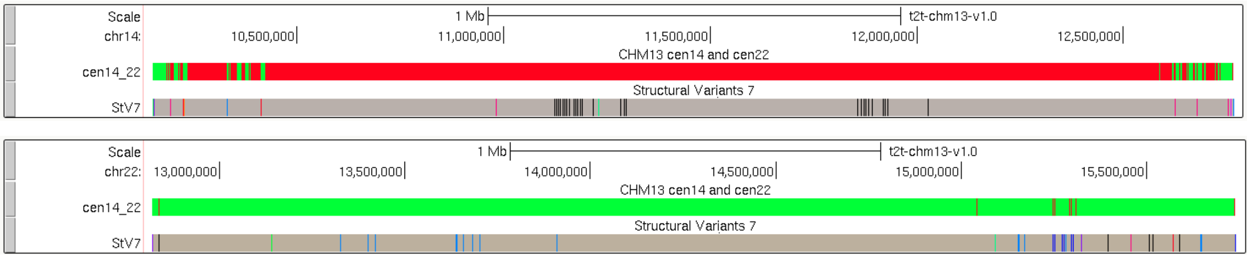

S2C14/22H1L_14.1-8 is red
S2C14/22H1L_22.1-8 is green
Cen14 and cen22 HORs were discriminated well except the flanks where the mixture of the 2 chromosome-specific HOR-haps is observed.
Consensus sequences for the 2 chromosome-specific HOR-haps were generated and found to differ in only 3 positions.

>CONS_S2C14/22H1L_14 (8mer)
CTACAAAAAGAGTGTATCAAAACTGCTCTGTCAAAAGGAAGGTTCTTCTCTGTTAGGTGAGTGCATACGTCA
TAAAGGAGTTTCTGAGAATGTTTCTGTCTAGTGGTTATGGGAAGATATTTGCTTTTTCACCGTAGGCCTCAG
AGCGCTCCAAATATCCACTTGCACATACTACAAAAAGAGTGCTTCAAAGCTGCTCTCTGAAACGGAATGTTC
AACTCTATGAGTTGAATGCAAACATCACAAAGACGTTTCTGAGAATGCTTCTGTCTAGATTTGATATGAAGAT
ATTCCCGTTTCCAACGAAATCTTCAAATCTATCCAAATGTCCACTTGCAGATTCAACAAAAAGTGTTTTTCAG
AACTGCTCTATCAAAAGAAAGATCCACCTCTGTTAGCTGAGTTCACACATCACAAACAAGTTTATGAGAATG
CTTCTGTCTAGTTTTTATTTGAAGATATTTCCTTTCTCACCATAGAGCTGAAAGCTGTCCTAATGTTCACTTCC
AGATACTACAGAAAGAGTGTTTCAAAACTGCTGTACGAAAGGGAATGTTCAACTCTGTGACTTGAATGCACA
CATCACAAAGAAGTTTCTGAGGATGCTGCTGTCTACTTTTTATACGTAATCCCGTTTCCAACGAAATCCTCCA
AGCTATCCAAATATCCACTTGCAGATTCCACAGAAAGACTGTTTCAAAACTGCTCTGTCAATAGAAAGGTTC
AACTCTGTTAGCTGCGTGCATATATCCCAAAGAAGATTCTGAGATTGCTTCTGTCTAGTTTTTATGGGAAGAT
ATTTCCCTTTTCACCGTAGGCGTCAAGGCGCTCCAAATGTCCACTTCCAGATACTACAAAAAGAGTGTTTCA
AACCTACTCTGTGAAAGGGAATATTCAACTCTGTGACTTGAATGCACATATCACAAGGAAGTTTCTGAGAAT
GCTTCTGTCGAGATTTTATATGAAGATATTCCCGTTTCCAACGAAATCCTGAAATCTATCCAAATATCCCCTCG
CAGATTCTACAAAAAGAGTGTTTCAAAACTGCTCTGTAAAAAGAAAGGTTCAACTCTGTTAGTTGAGTACAC
ACATCACAAACAAGTTTCACAGAATGCTTCTTTCTAGCTTGTAGGGGAAGATATTCCCTTTATCACCATGGGC
CTCAAACCGTCCGAAACGTCCACTTCCATATACTACAAAAAGAGCGTTTCAAACCTGCTCTATGAAAGGCAA
TGTTCAACTCTGTGACTTGAATGCAGACATCACAGAGCAGTTTCTGAGAATGCTTCTGTCTAGATTTTATAG
GAAGATATTCCCGTTTCCAACGAAATCTTCACAGCTATCCAAATATCCACTTGCAGATT

>CONS_S2C14/22H1L_22 (8mer)
CTACAAAAAGAGTGTATCAAAACTGCTCTGTCAAAAGGAAGGTTCTTCTCTGTTAGGTGAGTGCATACGTCA
TAAAGGAGTTTCTGAGAATGTTTCTGTCTAGTGGTTATGGGAAGATATTTGCTTTTTCACCGTAGGCCTCAG
AGCGCTCCAAATATCCACTTGCACATACTACAAAAAGAGTGCCTCAAAGCTGCTCTCTGAAACGGAATGTTC
AACTCTATGAGTTGAATGCAAACATCACAAAGACGTTTCTGAGAATGCTTCTGTCTAGATTTGATATGAAGAT
ATTCCCGTTTCCAACGAAATCTTCAAATCTATCCAAATGTCCACTTGCAGATTCAACAAAAAGTGTTTTTCAG
AACTGCTCTATCAAAAGAAAGATCCACCTCTGTTAGCTGAGTTCACACATCACAAACAAGTTTATGAGAATG
CTTCTGTCTAGTTTTTATTTGAAGATATTTCCTTTCTCACCATAGACCTGAAAGCTGTCCTAATGTTCACTTCC
AGATACTACAGAAAGAGTGTTTCAAAACTGCTGTACGAAAGGGAATGTTCAACTCTGTGACTTGAATGCACA

CATCACAAAGAAGTTTCTGAGGATGCTGCTGTCTACTTTTTATACGTAATCCCGTTTCCAACGAAATCCTCCA
AGCTATCCAAATATCCACTTGCAGATTCCACAGAAAGACTGTTTCAAAACTGCTCTGTCAATAGAAAGGTTC
AACTCTGTTAGCTGCGTGCATATATCCCAAAGAAGATTCTGAGATTGCTTCTGTCTAGTTTTTATGGGAAGAT
ATTTCCCTTTTCACCGTAGGCGTCAAGGCGCTCCAAATGTCCACTTCCAGATACTACAAAAAGAGTGTTTCA
AACCTACTCTGTGAAAGGGAATATTCAACTCTGTGACTTGAATGCACATATCACAAAGAAGTTTCTGAGAAT
GCTTCTGTCGAGATTTTATATGAAGATATTCCCGTTTCCAACGAAATCCTGAAATCTATCCAAATATCCCCTCG
CAGATTCTACAAAAAGAGTGTTTCAAAACTGCTCTGTAAAAAGAAAGGTTCAACTCTGTTAGTTGAGTACAC
ACATCACAAACAAGTTTCACAGAATGCTTCTTTCTAGCTTGTAGGGGAAGATATTCCCTTTATCACCATGGGC
CTCAAACCGTCCGAAACGTCCACTTCCATATACTACAAAAAGAGCGTTTCAAACCTGCTCTATGAAAGGCAA
TGTTCAACTCTGTGACTTGAATGCAGACATCACAGAGCAGTTTCTGAGAATGCTTCTGTCTAGATTTTATAG
GAAGATATTCCCGTTTCCAACGAAATCTTCACAGCTATCCAAATATCCACTTGCAGATT

##### cen1 - cen5 - cen19

The active HOR arrays of cen1, cen5 and cen19 mostly consist of non-canonical dimer S1C1/5/19H1L.6/4-5 (dimer proportions in S1C1/5/19H1L HOR arrays: cen1 - 0.69; cen5 - 0.45; cen19 - 0.96). Sequences of dimers were extracted for each chromosome separately from a monomeric track (AS_SF_HOR_Annot.bed). The dimers from cen1 inversion (chr1:124130687-125858867) were reversed. From each set of dimers a random sample of 500 sequences was made and aligned via MUSCLE (*33*) and used as HMMs (one for each chromosome). This did not yield satisfactory discrimination (shown below).

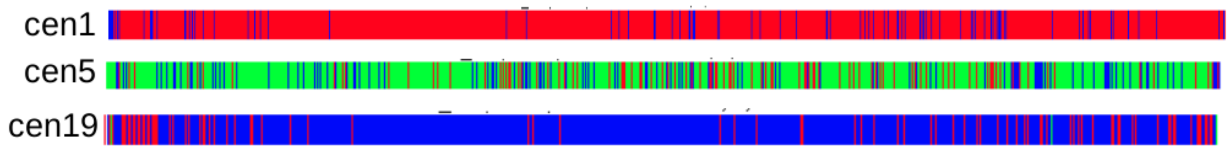

So, multiple alignments were used to build a tree where dimers coming from each chromosome were marked the same colors as in the tracks above. This tree showed that each chromosome has one or more major and well-isolated branch (haplotype), but also a lot of small branches which are also chromosome-specific, but sit nearby on the tree in a wide and mixed zone.

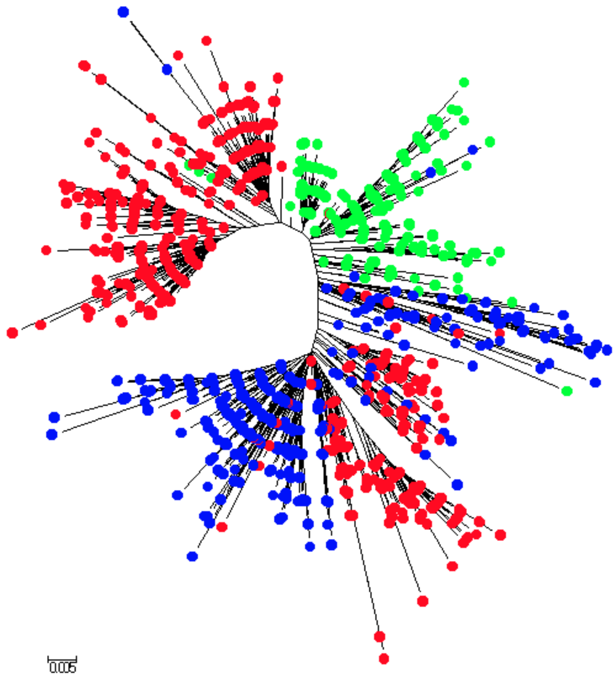

Hence, separate branches from this tree were extracted as sequences and used as HMMs (cen1 - 6 branches, cen5 - 2 branches, cen19 - 6 branches) to haplotype CHM13 T2T v1.0 assembly. The resulting haplotype tracks are shown below.

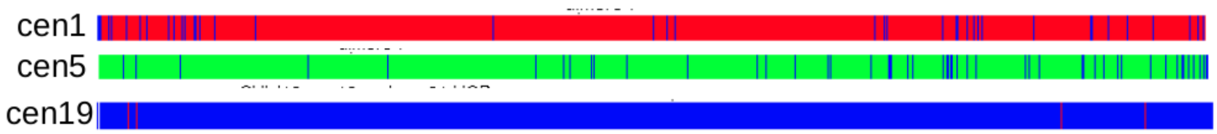

The 3 centromeres appeared to be well resolved proving that the S1C1/5/19 HOR has evolved independently in all 3 locations.

The consensus sequences were derived for each branch and compared to the consensus derived across the complete sequence set:

| consensus | differences from overall consensus |
|---|---|
| CONS-chr1-a | 3 |
| CONS-chr1-b | 7 |
| CONS-chr1-c | 11 |
| CONS-chr1-d | 7 |
| CONS-chr1-e | 7 |
| CONS-chr1-f | 3 |
| CONS-chr19-a | 4 |
| CONS-chr19-b | 3 |
| CONS-chr19-c | 7 |
| CONS-chr19-d | 4 |
| CONS-chr19-e | 11 |
| CONS-chr19-f | 1 |

| | |
|---|---|
| CONS-chr5-a | 6 |
| CONS-chr5-b | 6 |

The minimum evolution tree of consensus dimers was built to assess their relations. It can be seen that the ancestral S1C1/5/19H1 array which gave rise to the current arrays on all 3 chromosomes was likely to have a number of HOR-haps some of which got shared by different centromeres, and some more recent HOR-haps may have developed after the separation, as chromosome-specific sequence variants.

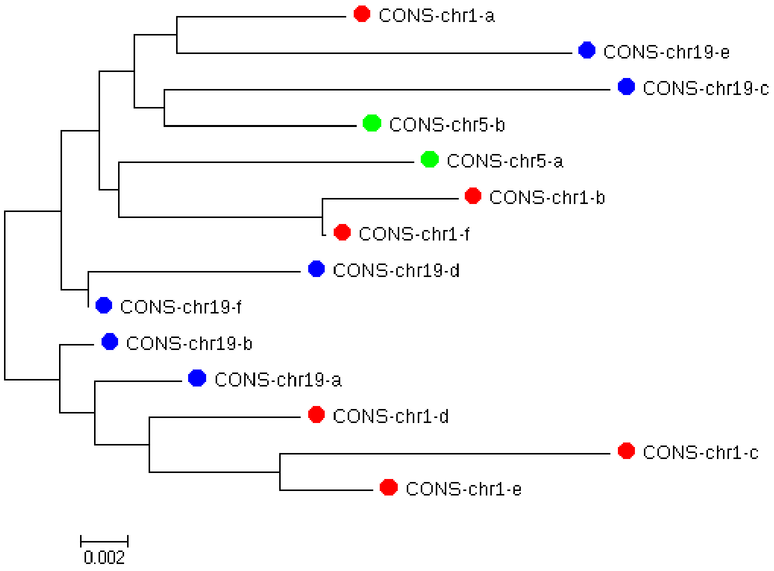

>CONS-chr1-a
AAGTCTGCAAGTGGATATTCAGACCTCCTTGAGGCCTTCGTTGGAAACGGGATTTCTTCATATTCTGCTAGA
CAGAAGAATTCTCAGTAACTTCCTTGTGTTGTGTGTATTCAACTCACAGAGTTGAACGATCCTTTACACAGA
GCAGACTTGAAACACTCTTTTTGTGGAATTTGCAAGTGGAGATTTCAGCCGCTTTGAGGTCAATGGTAGAA
AAGGAAATATCTTCGTATAAAGACTAGACAGAATGATTCTCAGAAACTCCTTTGTGATGTGTGCGTTCAACTC
ACAGAGTTTAACTTTTCTTTTCATAGAGCAGTTAGGAAACACTCTGTTTGTAA-

>CONS-chr1-b
-AGTCTGCAAGTGGATATTCAGACCTCTTTGAGGCCTTCGTTGGAAACGGGATTTCTTCATATTCTGCTAGA
CAGAAGAATTCTCAGTAACTTCCTTGTGTTGTGTGTATTCAACTCACAGAGTTGAACGATCCTTTACACAGA
GCAGACTTGAAACACTCTTTTTGTGGAATTTGCAAGTGGAGATTTCAGCCGCTTTGAGGTCAATAGTAGAAA
AGGAAATATCTTCGTAGAAAAACTAGACAGAATGATTCTCAGAAACTCCTTTGTGATGTGTGTGTTCAACTCA
CAGAGTTTAACCTTTCTTTTCATAGAGCAGTTAGTAAACACTCTGTTTATAAA

>CONS-chr1-c
-AGTCTGCAAGTGGATATTCAGACCTCTTTGAGGCCTTCGTTGGAAACGGGATTTCTTCATATTATGCTAGAC
AGAATAATTCTCAGTAACTTCCTTGTGTTGTGTGTATTCAACTCACAGAGTTGAAGGATCCTTTACAGAGAG
CAGGCTTGAAACACTCTTTTTGTCGAATTTGCAAGTGGAGATTTCAGCCGCTTTGAGGTCAATGGTAGAATA
GGAAATATCTTCTTATAGAAACTAGACAGAATGATTCTCATAAACTCCTTTGTGATGTGTGCGTTCAACTCAC
AGAGTTTAACCTTTCTTTTCATAGAGCAGTTAGGAAACACTCTGTTTGTAA-

```
>CONS-chr1-d
-AGTCTGCAAGTGGATATTCAGACCTCTTTGAGGCCTTCGTTGGAAACGGGATTTCTTCATATTATGCTAGAC
AGAAGAATTCTCAGTAACTTCCTTGTGTTGTGTGTATTCAACTCACAGAGTTGAACGATCCTTTACACAGAG
CAGACTTGAAACATTCTTTTTGTGGAATTTGCAAGTGGAGATTTCAGCCGCTTTGAGGTCAATGGTAGAATA
GGAAATATCTTCCTATAGAAACTAGACAGAACGATTCTCAGAAACTCCTTTGTGATGTGTGCGTTCAACTCA
CAGAGTTTAACCTTTCTTTTCATAGAGCAGTTAGGAAACACTCTGTTTGTAA-

>CONS-chr1-e
AAGTCTGCAAGTGGATATTCAGACCTCTTTGAGGCCTTCGTTGGAAACGGGATTTCTTCATATTATGCTAGA
CAGAATAATTCTCAGTAACTTCCTTGTGTTGTGTGTATTCAACTCACAGAGTTGAACGATCCTTTACACAGAG
CAGACTTGAAACACTCTTTTTGTGGAATTTGCAAGTGGAGATTTCAGCCGCTTTGAGCTCAATGGTAGAATA
GGAAATATCTTCCTATAGAAACTAGACAGAATGATTCTCATAAACTCCTTTGTGATGTGTGCGTTCAACTCAC
AGAGTTTAACCTTTCTTTTCATAGAGCAGTTAGGAAACACTCTGTTTGTAA-

>CONS-chr1-f
AAGTCTGCAAGTGGATATTCAGACCTCTTTGAGGCCTTCGTTGGAAACGGGATTTCTTCATATTCTGCTAGA
CAGAAGAATTCTCAGTAACTTCCTTGTGTTGTGTGTATTCAACTCACAGAGTTGAACGATCCTTTACACAGA
GCAGACTTGAAACACTCTTTTTGTGGAATTTGCAAGTGGAGATTTCAGCCGCTTTGAGGTCAATAGTAGAAA
AGGAAATATCTTCGTAGAAAAACTAGACAGAATGATTCTCAGAAACTCCTTTGTGATGTGTGCGTTCAACTC
ACAGAGTTTAACCTTTCTTTTCATAGAGCAGTTAGTAAACACTCTGTTTGTAA-

>CONS-chr19-a
AAGTCTGCAAGTGGATATTCAGACCTCCTTGAGGCCTTCGTTGGAAACGGGATTTCTTCATATTCTGCTAGA
CAGAAGAATTCTCAGTAACTTCCTTGTGTTGTGTGTATTCAACTCACAGAGTTGAACGATCCTTTACACAGA
GCAGACTTGAAACACTCTTTTTGTGGAATTTGCAAGTGGAGATTTCAGCCGCTTTGAGGTCAATGGTAGAAT
AGGAAATATCTTCCTATAGAAACTAGACAGAATGATTCTCAGAAACTCCTTTGTGATGTGTGCGTTCAACTCA
CAGAGTTTAACCTTTCTTTTCATAGAGCAGTTAGGAAACACTCTGTTTGTAA-

>CONS-chr19-b
AAGTCTGCAAGTGGATATTCAGACCTCTTTGAGGCCTTCGTTGGAAACGGGATTTCTTCATATTATGCTAGA
CAGAAGAATTCTCAGTAACTTCCTTGTGTTGTGTGTATTCAACTCACAGAGTTGAACGATCCTTTACACAGA
GCAGACTTGAAACACTCTTTTTGTGGAATTTGCAAGTGGAGATTTCAGCCGCTTTGAGGTCAATGGTAGAAT
AGGAAATATCTTCCTATAAAAACTAGACAGAATGATTCTCAGAAACTCCTTTGTGATGTGTGCGTTCAACTCA
CAGAGTTTAACCTTTCTTTTCATAGAGCAGTTAGGAAACACTCTGTTTGTAA-

>CONS-chr19-c
AAGTCTGCAAGTGGATATTCAGACATCTTTGAGGCTTTCGTTGGAAACGGGATTTCTTCATATTCTGCTAGA
CAGAAGAATTCTCAGAAACTTCCTTGTGTTGTGTGTTTTCAACTCACAGAGTTCAACGATCCTTTACACAGA
GTAGACTTGAAACACTCTTTTTGTGGAATTGGCAAGTGGAGATTTCAGCCGCTTTGAGGTCAATGGTAGAA
AAGGAAATATCTTCGTATAAAAACTAGACAGAATGATTCTCAGAAACTCCTTTGTGATGTGTGCGTTCAACTC
ACAGAGTTTAACCTTTCTTTTCATAGAGCAGTTAGGAAACACTCTGTTTGTAA-

>CONS-chr19-d
```

AAGTCTGCAAGTGGATATTCAGACCTCTTTGAGGCCTTCGTTGGAAACGGGATTTCTTCATATTATGCTAGA
CAGAAGAATTCTCAGTAACTTCCTTGTGTTGTGTGCATTCAACTCACAGAGTTGAACGATCCTTTACACAGA
GCAGATTAGAAACACTCTTTTTGTGGAATTTGCAAGTGGAGATTTCAGCCGCTTTGAGGTCAATGGTAGAAA
AGGAAATATCTTCGTATAAAAACTAGACAGAATGATTCTCAGAAACTCCTTTGTGATGTGTGCGTTCAACTCA
CAGAGTTTAACCTTTCTTTTCATAGAGCAGTTAGGAAACACTCTGTTTGTAA-

>CONS-chr19-e
--GTCTGCAAGTGGATATTCAGACCTCCTTGAGGCCTTCGTTGGAAACGGGATTTCTTCATATTCTGCTATAC
AGAAGAATTCTCAGAAACTTCCTTGTGTTGTGTGTATTCAACTCACAGAGTTGAACGATCGTTTACACAGAG
CAGACTTGAGACACTCTTTTTGTGGAATTTGTAAGTGGAGATTTCAGCCGCTTTGAGGTCAATGGTAGAAAA
GGAAATATCTTCATATAAAAACTAGACAGAATGATTCTCAGAAACTCCTTTGTGATGTGTGCGTTCAACTCAC
AGAGTTTAACCTTTCTTTTCATAGAGCAGTTAGGAAACACTCTGTTTGT---

>CONS-chr19-f
AAGTCTGCAAGTGGATATTCAGACCTCTTTGAGGCCTTCGTTGGAAACGGGATTTCTTCATATTATGCTAGA
CAGAAGAATTCTCAGTAACTTCCTTGTGTTGTGTGTATTCAACTCACAGAGTTGAACGATCCTTTACACAGA
GCAGACTTGAAACACTCTTTTTGTGGAATTTGCAAGTGGAGATTTCAGCCGCTTTGAGGTCAATGGTAGAA
AAGGAAATATCTTCGTATAAAAACTAGACAGAATGATTCTCAGAAACTCCTTTGTGATGTGTGCGTTCAACTC
ACAGAGTTTAACCTTTCTTTTCATAGAGCAGTTAGGAAACACTCTGTTTGTAA-

>CONS-chr5-a
-ACTCTGCAAGTGGATATTCAGACCTCTTTGAGGCCTTCGTTGGAAACGGGATTTCTTCATACTGTGCTAGA
CAGAAGAATTCTCAGTAACTTCCTTGTGTTGTGTGTATTCAACTCACAGAGTTGAACGATCCTTTACACAGA
GCAGACTTGAAACACTCTTTTTGTGGAATTTGCAAGTGGAGATTTCAGCCGCGTTGAGGTCAATGGTAGAA
AAGGAAATATCTTCGTATAAAAACTAGACAGAATGATTCTCAGAAACTCCTTTGTGATGTGTGTGTTCAACTC
ACAGAGTTTAACCTTTCTTTTCATAGAGCAGTTAGGAAACACTCTGTTTGTAA-

>CONS-chr5-b
--CTCTGCAAGTGGATATTCAGACCTCTTTGAGGCCTTCGTTGGAAACGGGATTTCTTCATATTCTGCTAGAC
AGAAGAATTCTCAGAATCTTCCTTGTGTTGTGTGTATTCAACTCACAGAGTTGAACGATCCTTTACACAGAG
CAGACTTGAAACACTCTTTTTGTGGAATTTGCAAGTGGAGATTTCAGCCGCTTTGAGGTCAATGGTAGAAAA
GGAAATATCTTCGTATAAAAACTAGACAGAATGATTCTCAGAAACTTCTTTGTGATGTGTGCGTTCAACTCAC
AGAGTTTAACCTTTCTTTTCATAGAGCAGTTAGGAAACACTCTGTTTGTAA-

#### Flow Sorted Chromosome Comparison with CHM13v1.0 assemblies

Counts of various monomers (including many hybrids) of AS active array HOR S1C1/5/19H1L in centromeres of three chromosomes (1, 5, and 19), calculated from HOR annotation of CHM13 effectively haploid 22XX cell line complete T2T assembly v1.0 and the same counts performed in Illumina short read libraries prepared

from flow-sorted chromosomes 1, 5 and 19 obtained from the same cell line. Only the near full-length monomers (length >= 150 bp) were counted in all samples to reduce the risk of erroneous monomer classification. Flow-sorted paired-end reads where merged using fastp tool (*34*) (https://github.com/OpenGene/fastp#merge-paired-end-reads).
In flow-sorted samples, 601,588 (chr1), 339,136 (chr5), and 2,281,989 (chr19) S1C1/5/19H1L monomers were identified and 127,496 (chr1), 72,363 (chr5), and 438,341 (chr19) monomers remained after the length filtering. In the assembly, 26,430 (chr1), 14,885 (chr5), and 22,905 (chr19) S1C1/5/19H1L monomers were identified and 26,379 (chr1), 14,885 (chr5), and 22,902 (chr19) monomers remained after the length filtering.
The two samples used where: 151 bp flow-sorted Illumina reads (Globus link: https://app.globus.org/file-manager?origin_id=9db1f0a6-a05a-11ea-8f06-0a21f750d19b&origin_path=%2Fteam-curation%2Fflowsort%2Fraw_reads%2F) and the T2T CHM13 assembly v1.0 (top panels) (Globus link: https://app.globus.org/file-manager?origin_id=9db1f0a6-a05a-11ea-8f06-0a21f750d19b&origin_path=%2Fassemblies%2Frelease%2Fv1.0%2F).

##### Monomer classification/identification was performed by HumAS-HMMER-HOR

(https://github.com/enigene/HumAS-HMMER) described in its partial form (SF1 HORs only) in (*13*), and in its complete form (all known HORs covered) in this paper. It uses the HMMER platfom (*11*) and in its current form it identifies (as the best match in a set of standards) each monomer of about 80 known HORs and about 30 classes of monomers in AS monomeric layers. Note that the standards used for monomers of homogeneous HORs are just one or several representatives for each monomer installed as independent standards, not as multiple alignments. So, the best match for them is just the best ordinary alignment. For divergent HORs, the standard is formed by a number of representative monomers used as multiple alignment and therefore constitutes a true HMM. Conversion of the HMMER output file into a bed file was performed using hmmertblout2bed script (https://github.com/enigene/hmmertblout2bed) as described in (*13*).

##### Array specific k-mer comparison

All k-mer analyses were performed using KMC v3,1.1 (*35*) plus some additional custom functionality which can be found at https://github.com/msauria/KMC. Analysis code can be found at https://github.com/msauria/T2T_Kmer_Analysis.

For 75-mer analysis, all sequences were split based on repeat masker annotations ensuring that no k-mer overlapped any part of a repeat masked sequence. 75-mer databases were created for the complete CHM13v1 genome, each array type, each specific array instance, and all non-centromeric sequences. For each specific array greater than 100 kb, the number of 75-mers was counted for each of the following categories: unique to that specific array, shared with another specific array, specific to the specific array's general array class, shared exclusively between two general array classes (the specific array's class and another), specific to centromeric sequence, and total number of 75-mers.

To create k-mer sets for HOR and BSAT analyses, 100-mer and 21-mer databases were created, respectively, for each specific instance of each of these array classes. For every k-mer found in the array class, all specific array instances containing that k-mer, along with the number of occurrences, were identified. In addition, k-mers were labelled as "general" if they occurred anywhere outside of the target array class or "specific" otherwise.

#### Section 4: Study of haplotype phased assemblies

##### Diversity Panel of HiFi Data used in HOR Structural Variant analysis

Human Pangenome Reference Consortium (HPRC) (*36*) generated long and accurate HiFi reads for sixteen human samples HG002, HG003, HG004, HG005, HG006, HG007, HG01243, HG02055,  HG02109, HG02723, HG03492, HG01109, HG01442,HG02080,HG02145, and HG03098. We refer to these datasets as HPRC samples. (https://github.com/human-pangenomics/hpgp-data).

##### Detecting inversions, insertions, and deletions in hifiasm assemblies

Winnowmap (*22*)alignments between CHM13 and contigs from hifiasm assemblies from 16 HPRC Plus cell lines were obtained (cell lines HG005, HG00733, HG01109, HG01243, HG02080, HG02109, HG02145, HG02723, HG02818, HG03098, HG03486, HG03492, NA18906, NA19240, NA20129, NA21309). To search for a polymorphic deletion containing an HSat3B2 (*3*) array on chr1 (**Fig. 2a**), the identities, sequences, and haplotype assignments of contigs overlapping the chr1 alpha-hsat transition region (chr1:126700000-144000000) were obtained. 27 of the possible 32 haplotypes had long contigs encompassing most or all of the transition region. Each contig sequence was scanned for HSat2,3 subfamily-specific 24-mers on both strands, and adjusted for the relative abundance of each 24-mer set in each subfamily (*3*). 24-mer frequencies were automatically plotted along each contig, with positive signed frequencies indicating matches on the positive strand, and negative signed frequencies indicating matches on the negative strand (Fig. S5). Plots were manually scanned to note the presence or absence of the HSat3B2 array in the context of the other arrays in the transition region.

To detect inversion and insertion breakpoints in HOR arrays on chrs 1, 3, 4, a similar approach was taken, but using sets of 21-mers specific to Alpha, HSat1, HSat2, and HSat3 (previously identified in (*8*)). Contigs aligning to each breakpoint were obtained for all 32 haplotypes. These were scanned for the family-specific 21-mers, which allowed for ‘painting’ of each contig by satellite family and strand to confirm the presence of inversion and insertion breakpoints in the correct contexts and orientations. The results for all individuals are summarized in Fig S6. Sometimes no contig was found overlapping the relevant region, and these are noted as ‘?’, and sometimes only 1 of two breakpoints was spanned, and these are noted as ‘(+)’. Classification of the chr1 inversion required sufficiently long contigs to span most of the D1Z7 array, which were not available for 9 haplotypes.

#### Section 5: The evolution of human centromere 6

Chromosome 6 centromere represents an interesting example of centromere evolution. It was known that in Catarrhini ancestor Chr6 centromere was situated near pos. 26Mb of the modern human chromosome (*37*). In Macaca mulatta, this old centromere went defunct and repositioned to a new place. In humans, old centromere went defunct and a new one emerged near the modern position of human cen6. Such cases are known as Evolutionary New Centromeres (ENC) (*37*)). The complete assembly of the human Chr6 gives us an opportunity to investigate the origin of ENC on Chr6. The sequence of human Chr6 and available contigs of primates was compared with each other by similarity search and dot-plot construction.

The structure of Chr6 centromeric region (as shown below) features two ASat domains. The left one includes the huge S01C6H1L active HOR domain flanked by SF8 mon_6_2 domain and by S01C3/6H1d dhor_6_1 domain which has significant islets of S01CMH1d (chr6:61058623-61413737). The right ASat domain is represented by a short S01CMH1d domain surrounded by two monomeric ASat near-symmetrical flanks which feature large SF8 domains. Symmetrical disposition of monomeric layers and the core of more recent SF01 divergent HORs are very much reminiscent of the dead degraded primate centromere in chr2q. Therefore we suggest that this locus is indeed what remains of another dead centromere, which was active and then abandoned at the time when SF01 formed the active arrays and before the spread of more recent SF1. This suggests the dead centromere belongs to some primate ancestor before the separation of human lineage with gorilla which already has SF1. This aligns well with the dating of this locus by analysis of sharing with primate lineages (Table S8). The left side arm region adjacent to this ASat array (about 1 Mb) and a short fragment of SF8 were involved in a large duplication later modified by inversion of fragment E and some insertions. Interestingly, insertion between Abc and D fragments on the p-arm comes from Chr6p near position 26M which is the site of the old Catarrhini ancestor Chr6 centromere (*37*) and a human clinical neocentromere region (*38*). This duplication has created an alternative ASat array in ~1Mb distance from the first, which apparently has assumed centromeric function and hosted the amplification of another SF01 HOR shared with chr3, and later on the amplification of the currently active SF01 HOR (S01C6H1L).

Bird’s eye view of the whole cen6 region

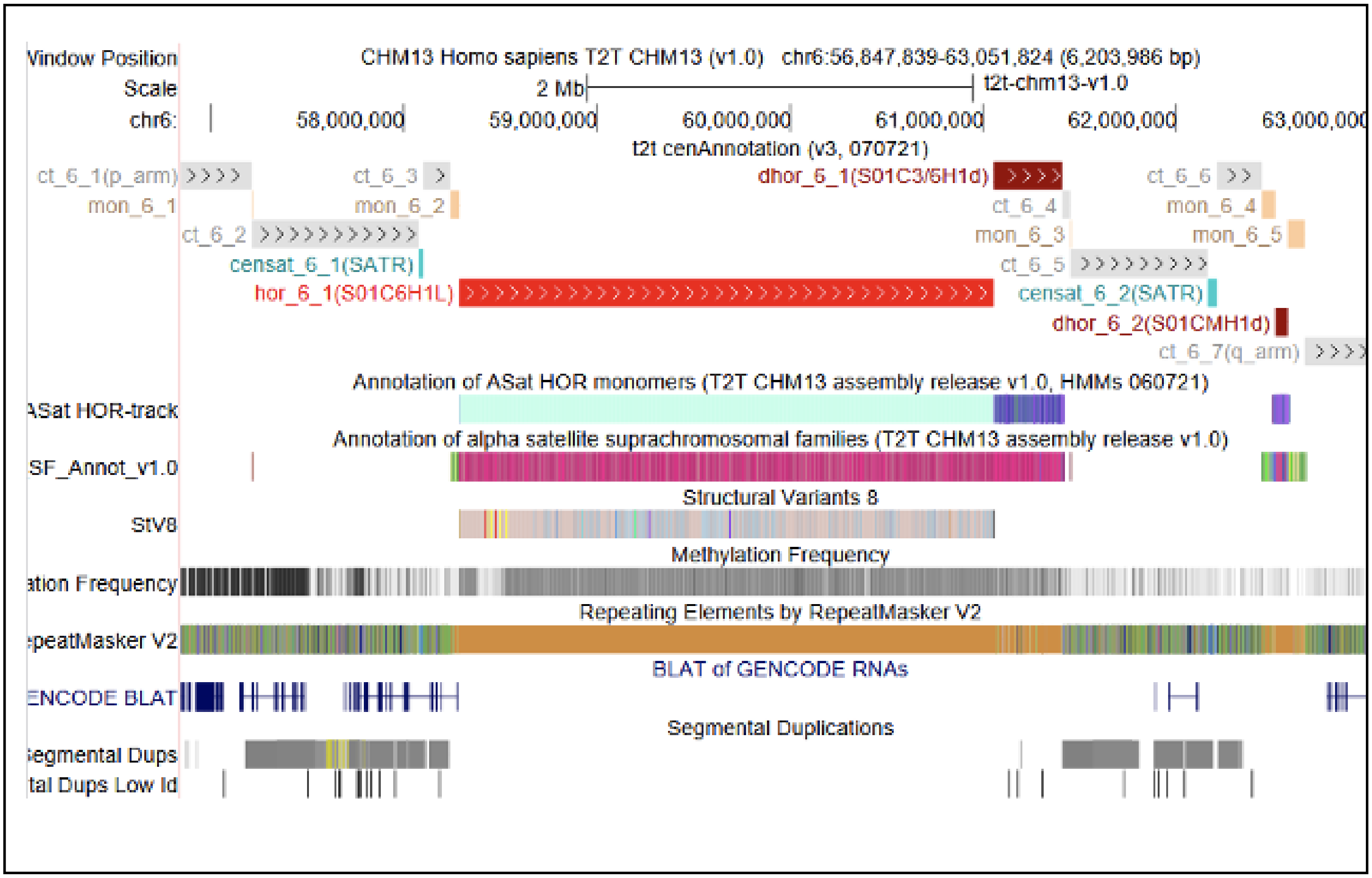

A close-up of the old centromere region in chr 6.

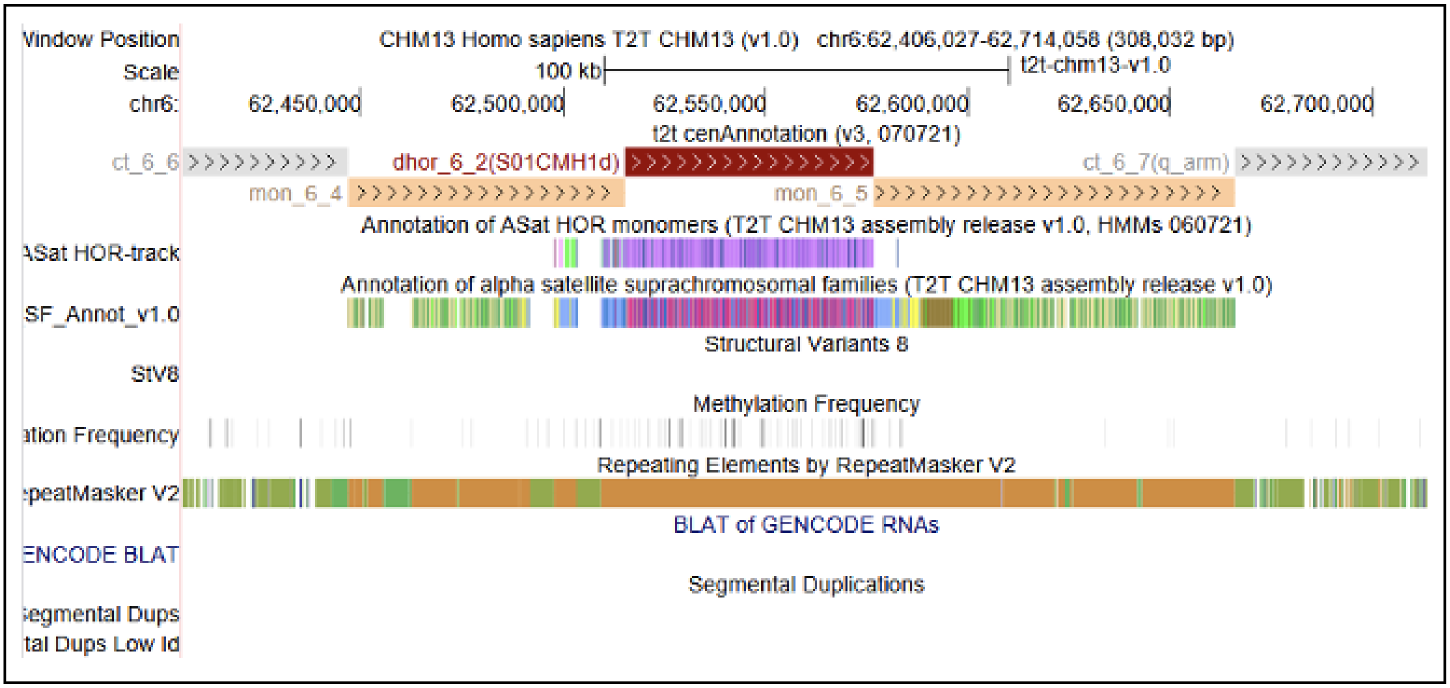

A close-up of the old centromere region in chr 2q (for comparison).

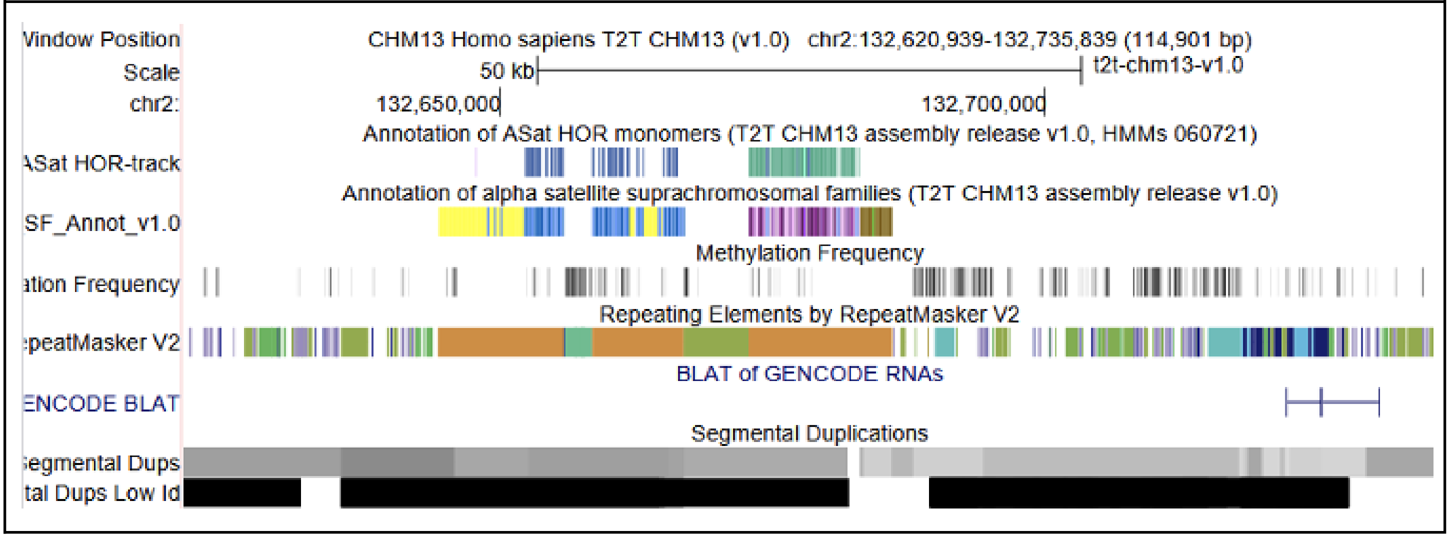

**Legend to Supplementary Figure (below).** In the bottom part, is a dot-plot of the p-arm vs. q-arm of chr 6 which shows the structure and repetition pattern of the elements. In the top part, there is a schematic of a proposed scenario of the human Chr6 centromere evolution. This scenario includes SF8 ASat seeding and expanding in a new place. Next is expansion of the younger monomeric layers down to SF4. This stage is reflected by the current orangutan centromere, the flanks of which are as indicated by the scheme. Next stages include a duplication of the left part of the centromere, inversion of segment E, origin and expansion of S01C6H1L, as described in Section 5: The evolution of human centromere 6.

Letter code.
Letters A, B, C, D, E, F a, b, c, d, e, f denote six repeated fragments detailed in the region. Uppercase and lowercase letters represent orthologous but slightly diverged elements.
Divergence from top to bottom is assumed. F and f orthologous segments include both ASat and non-ASat sequences. Red ovals on dot-plot and vertical arrows represent repeated elements on the p-arm. Copies on q-arm are not connected for figure simplicity.

SCENARIO OF NEW HUMAN CHR6CEN EVOLUTION

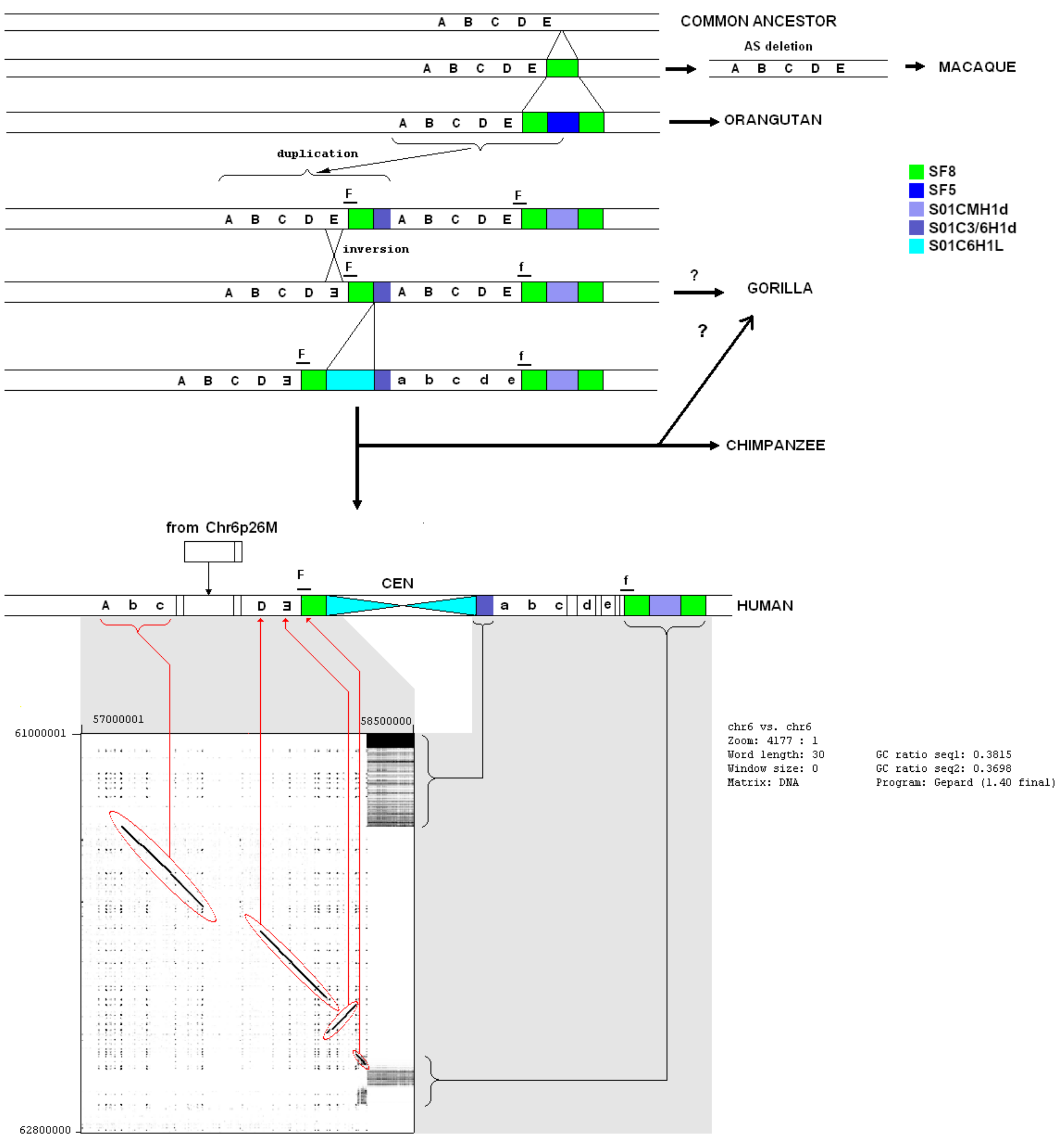

#### Section 6: HOR unit and NRS prediction

##### HOR Structrual Variant (StV) prediction using CHM13 hor-monomer annotation

The StV track was derived from the AS HOR annotation track (HOR-track; AS_HOR_Annot.bed) by using python scripts. The HOR-track shows the coordinates of every monomer in the assembly and its name which indicates to which HOR the monomer belongs and what is the number of this monomer in a HOR (defined as a standard master HOR with a fixed cyclic shift, see the HOR-track description and Uralsky 2019). So, each array is identified by a HOR name (e.g. S3CXH1L for the live array of the X chromosome) and each monomer by its number (e.g. S3CXH1L.1 for monomer #1 of the HOR). Hybrid monomers are indicated with a "/" (e.g. S2C8H1L.4/7, where the first part of the monomer is S2C8H1L.4 and the second is S2C8H1L.4/7).

The main idea of generating the StV map was iterating through AS HOR annotation bed file and cutting after the last monomer in a HOR (the one with maximal number in a given array) or before the first monomer (the one with number #1) if the last monomer is not present in a given StV. For arrays where AS was on reverse strand, a cut was performed after the first monomer or before the last monomer if the first monomer is not present in a given StV. Note that about a half of the live centromeres have AS on direct strand and another half on reverse strand. Some arrays may include an inversion (e.g. chr1:124130688-125857749).

**StV naming**. The first part of the StV name is the HOR name (e.g. S2C8H1L). Then, after "." follows listing of monomer numbers included in StV separated by "_". If monomer numbers go in natural order (or descending natural order for the reverse strand HORs), we join the first and the last monomers in this string with "-" to indicate an interval and do not show the numbers in between (e.g. S3C11H1L.1_2_3 would appear S3C11H1L.1-3). If AS is on reverse strand, the monomer numbers go in reverse (e.g. S3CXH1L.1-12 would appear as S3CXH1L.12-1). Hybrids are always flanked by "_" (e.g. S2C8H1L.1-3_4/7_8-11). Rarely occurring monomers identified as monomers of another HOR or as SF class monomers (usually due to misclassification) are shown by their own name (e.g. S2C15H1L.1-4_S5C1qH6d.2_5-11 or S3C11H1L.1-2_W3_5).

The coordinates of the live HORs were taken from the centromere annotation track (t2t_cenAnnotation.v2.021921.bed). The resulting full-length HORs and StVs were counted and listed in the stats table. Reverse strand HOR and StV names were reversed in the stats table to make it easier to read (e.g. S3CXH1L.12-11_8-1 would become S3CXH1L.1-8_11-12). In centromeres 1, 5 and 19, there are long chains of "_6/4_5" repeats which appear within a HOR (HOR formula S1C1/5/19H1L.1-5(_6/4_5){n}-6). In StV names these repeats were shown as "(_6/4_5){n}" (e.g. S1C1/5/19H1L.1-5(_6/4_5){7}-6 where {7} indicates the number of repeats in a given HOR). In the stats table all numbers in "{}" were replaced by "{n}" because we consider them as one StV, however the statistics for {n} is collected by the script and can be retrieved. The StVs and full-length HORs in every centromere were numbered. A number is at the beginning of StV's name after "#" sign. After a number there is ":" sign (e.g. #117:S3CXH1L.12-11_8-1). Numbering goes from p- to q-side of a chromosome regardless of the AS direction. The StV track was colored randomly but rare StVs (<10% in a centromere) are in bright colors to increase visibility and common StVs and full-length HORs are in dim colors.

##### Automated annotation of “live” alpha satellite arrays.

*Alpha satellite arrays* in “live” human centromeres are tandem repeats that are formed by units repeating thousands of times with limited nucleotide-level variations but extensive variations in copy numbers in the human population (*39*). Each such unit represents a tandem repeat formed by smaller repetitive building blocks (referred to as *monomer-blocks*), thus forming a *nested tandem repeat.* Partitioning all monomer-blocks into *n* clusters of similar monomer-blocks defines *n monomers,* where each monomer represents the consensus of all monomer-blocks in a given cluster. A *canonical order of monomers* (referred to as a *high-order repeat* or *HOR*) is specific for each centromere and is evolutionarily defined as the ancestral order of monomers that has evolved into the complex organization of modern centromeres. In addition to units formed by canonical HORs, there exist units formed by *partial HORs* (substrings of canonical HORs).

Recent evolutionary studies of alpha satellite arrays (*13*, *40*, *41*) revealed the importance of partitioning them into monomers, the problem that was addressed by the StringDecomposer algorithm (*23*). StringDecomposer opened a possibility to *automatically* derive all HORs and annotate alpha satellite arrays, the problem that remains unsolved despite multiple studies in the last four decades (*9*, *13*, *16*, *42–45*). Recently, Kunyavskaya et al., 2021 presented HORmon — a pipeline that resulted in automated derivation of all monomers and HORs in “live” alpha satellite arrays, and annotation of these arrays that is largely consistent with previous semi-manual centromere annotations (*9*, *13*).

Given a nucleotide string *Centromere*, HORmon transforms it into a monocentromere *Centromere** written in the monomer alphabet. As an alternative approach to centromere annotation, HORmon uses the concept of the monomer-graph (*46*). For a given monocentromere, a directed *monomer-graph* is constructed on the vertex-set of all monomers and the edge-set formed by all pairs of consecutive monomers in the monocentromere. The multiplicity of a vertex *M* in the monomer-graph is defined as the number of monomer-blocks corresponding to the monomer *M*. The multiplicity of an edge (*M,M’*) in the monomer-graph is defined as the number of times the monomer *M’* follows the monomer *M* in the monocentromere.

The genome browser tracks with monomer and HOR decompositions of alpha satellite arrays in the CHM13 cell line are available at:
http://genome.ucsc.edu/cgi-bin/hgTracks?genome=t2t-chm13-v1.0&hubUrl=http://t2t.gi.ucsc.edu/chm13/hub/hub.txt.

Accession numbers for extracted nucleotide sequences of HORs can be found in Table NucleotideConsensusOfHORs.

##### HOR annotation of HiFi reads.

While centromeres in a haploid cell line were successfully assembled, centromere assembly in a diploid genome remains an open problem. Therefore, even though HiFi reads were generated for multiple HPRC samples, it remains unclear how to analyze variations in centromere architectures across the human population. We thus modified the HORmon in order to generate monomer-graphs based on HiFi reads rather than complete centromere assemblies. For each HPRC sample, the HiFi reads were mapped to the CHM13 assembly (v1.0) using Winnowmap2 (*22*). Reads mapping to “live” alpha satellite arrays (referred to as centromeric reads) were subsequently recruited (coordinates of arrays were selected according to Kunyavskaya et al., 2021)(*46*).

We utilized the monomer-set generated by HORmon for the CHM13 cell line (18145 monomers) and applied StringDecomposer (*23*)) in order to transform each centromeric read into *monoread* written in the monomer alphabet. To ensure that all considered reads are in fact centromeric, we filtered out reads with average monomer identity below *AvgMonoIdentity* (default value *AvgMonoIdentity* = 95%). HORmon decomposes each monoread into canonical and partial HORs (*46*) resulting in *HORDecomposedReads*. We follow this approach and decompose each monoread in the HPRC samples (rather than an assembly) into the set of HORs generated by HORmon for the CHM13 cell line.

##### HOR decompositions of HiFi reads.

The count of a HOR H in a sample S (referred to as count(H, S)) is defined as the number of its occurrences in HOR decompositions of all monoreads in sample S. For a given centromere C and each sample S, we order HORs by their decreasing counts and refer to the i-th most frequent HOR in C of sample S as HC,S,i. We identify the imin(C,S) as the minimum value of i such that total count of HC,S,i exceeds the threshold MinFraction · |HORDreads(C, S)|, where |HORDreads(C, S)| is the total length of all HORDecomposedReads in centromere C of sample S (the default value MinFraction = 0.99). Finally, we define the set of frequent HORs in a centromere C as a union of HORC,S,imin(C,S) over all samples S.

For each centromere, Figure MostFrequentHORs compares frequencies of the most frequent HORs among HPRC samples and illustrates the variability of the centromere architectures across the human population. It demonstrates that while the architecture of some centromeres (e.g., cenX) is rather stable across all analyzed HPRC genomes; other centromeres (e.g., cen6, cen13, cen16, cen17, and cen21) reveal recent evolutionary innovations:

- **cen6.** The second most frequent HOR p16-12 (15-mer) in CHM13 assembly occupies a large fraction of its length (23 %). However, in HPRC samples, its fraction ranges from 0 to 9 %.
- **cen13.** In CHM13, fractions of canonical HOR and its partial HOR p5-11 are similar (45 and 49% respectively). This HOR is also frequent in samples HG002,4,5,7 (21-34%). However, its frequency is much lower in the rest of the samples (0-9%).
- **cen16.** Sample HG02723 demonstrates unordinary HORs fractions. In other HPRC samples and in CHM13 assembly, the dominant HOR is canonical (72-80%). However, in HG02723, the canonical HOR occupies only 31% of the centromere. Contrary to other samples, HG02723 presents an extraordinary number of four auxiliary and partial HORs: F, p2-9, B, and p4-9. The typical fragment of HOR decompositions where these HORs are prominent is B F p4-9 B F p4-9 p2-9 p2-9 B F p4-9. Importantly, F-blocks typically have low identity (~95% instead of

98-99% for other monomers). It is unclear whether this monomer is novel or is a frequent variant of another known monomer.

- **cen17 (S3C17H1L).** Ten samples (HG004, HG01109, HG01243, HG01442, HG02055, HG02080, HG02109, HG02145, HG02723, HG03492) have high fractions of HORs p30-28 (10-17%), O17+AS17 (11-21%), and p30-25 (6-18%). On the other hand, the fraction of live canonical HORs in these samples averages at 29%, unlike the rest of samples where it averages at 60%.
- **cen21.** HG006 is the only sample that shows a substantially distinct HOR structure from the one presented in the CHM13 assembly. While in all other HPRC samples and the CHM13 assembly, the dominant HORs is canonical (68-83%), its frequency in HG006 is only 29%. Two dominant HORs in HG006 are a single monomer C and a partial 7-mer p8-3. HOR decompositions of reads in HG006 show that these two HORs typically form repetitive structures C p8-3 C p8-3 C p8-3.

Other differences include cen12, and 20. In cen12, two samples HG02055, HG02723 present lower numbers of canonical HORs (31 and 26% respectively with 72% average for other samples) and higher numbers of partial HOR p6-4 (24 and 31% respectively with 1.7% average for other samples). In three samples (HG004, HG007, HG01442), cen20 shows a high fraction of the second most frequent partial HOR p7-16 (average 11.6% and 1.5% for the rest of the samples).

##### Calling repeat periodicity with NTRprism

NTRprism takes as input a DNA sequence, a choice of k (6 by default), a maximum interval length (30,000), and a minimum k-mer count (10). It has linear computational complexity with respect to sequence length and works by computing, for each possible k-mer satisfying the minimum k-mer count (minimum number of times a k-mer must be observed across the input sequence), the empirical distribution of interval lengths between adjacent occurrences of that k-mer. This distribution is normalized to sum to 1 for each k-mer. Then, for each interval length (or for all interval lengths in a bin), the values across all k-mers are summed, and the sum is divided by the total number of k-mers. This is plotted as a Nested Tandem Repeat spectrum. The highest peak in the spectrum is reported as the most frequent periodicity found in the input sequence. To evaluate the performance of the method, it was compared to annotated alpha canonical HOR lengths across the genome, recovering the exact canonical HOR periodicity for nearly every array (Fig. S10B). Discordant results are explained by arrays in which non-canonical HOR structural variants predominate in the array. NTRprism was also run on simulated ‘alpha satellite’ HOR sequences, in which random 171 bp sequences were generated with varying user-specified pairwise percent identities, then repeated in HORs with varying user-specified pairwise percent identities (Fig. S10C). NTRprism was able to recover the correct simulated HOR periodicity in all cases, except when inter-HOR divergence greatly exceeded inter-monomer divergence (20-30% vs 5%, respectively). When provided with a fully randomized, non-periodic sequence, NTRprism reports the expected exponential distribution of interval lengths (Fig. S10D). To visualize how repeat periodicities may shift across an array, an HSat3 array was split into 100-kb windows and these windows were input into NTRprism separately (Fig. S10B). The code for NTRprism will be made available on github.

#### HOR haplotype classification

Annotated HOR libraries (StV data) of active arrays in CHM13 were used to construct sequence-based comparisons and HOR-haplotype classifications. Whenever possible, full-length canonical HOR sequences were used in HOR-hap predictions. However, several arrays represented a mixture of different length HORs, and in these cases we identified a common span, or grouping of monomers that provided consistent sequence information across the majority of the array. Next, we generated a multiple alignment of all HOR fasta for each active array (same orientation) using Kalign software (*47*). Resulting alignment files were used to identify the most common base at every position or consensus sequence. We generated pairwise alignment (needle, EMBOSS (*48*)) against the We generated pairwise alignment (needle, EMBOSS (*48*))against the CHM13 DXZ1 consensus sequence. Each alignment was reformatted into an ordered string of 0's and 1's, where 0 indicates a match to the shared position with the consensus and 1 if there is a difference at that position relative to the consensus. These tables were then read into r (*49*) and plots were made using ggplot2 (*50*). Prediction of the best "k" clusters used "FitKMeans"(max.clusters=20,nstart=25,seed=278613), using Hartigan's Number (visualized using "HartiganPlot") to determine if the k+1st cluster should be added. For all arrays, we found evidence for classification into a small number of k-clusters (ranging in size from k=2 to k=4). These classifications revealed the dark red versus grey expansions (Figure 3D,E). Examples on chromosomes 3 and 4 are shown below.

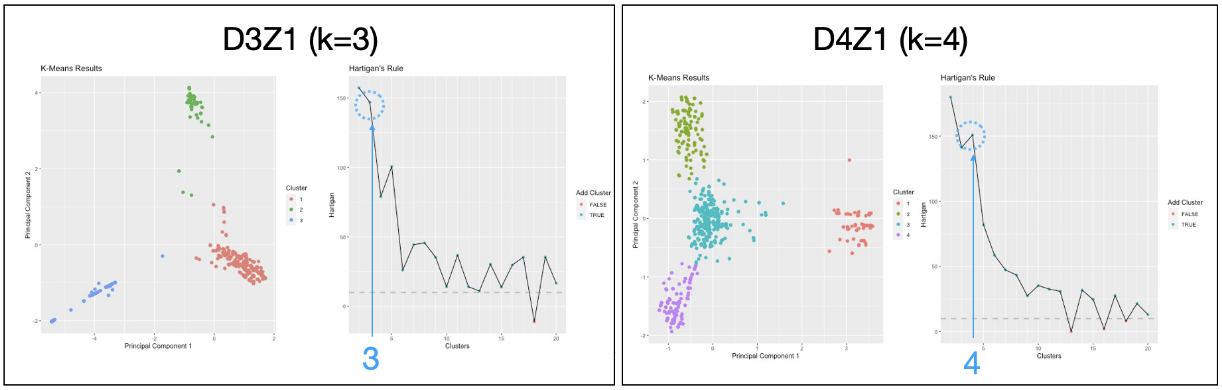

**Figure Legend**: Clustering of the active arrays on chromosome 3 (D3Z1,S01/1C3H1, 2720 bp representing monomers (m)1-4 and m6-17) revealed support for three cluster groups, selecting initial hor-hap groupings of k=3 (showing results for PCA plot of k-means clusters, and Hartigan plot). When studying these cluster patterns relative to the array, cluster 1 (red) was positioned in the center and clusters 2 and 3 were flanking either side (proximal to p-arm and q-arm). Using a similar method we predict four clusters for the entire D4Z1 array.

We also characterized these clusters at higher number "k-clusters" (as featured in Figure 4C) or separated the grouping and clustered them independently of one another (as shown for chromosome 3 in figure 3E, providing clustering information only from the dark red grouping and reclassified again to identify more substructure). Data and results associated semi-automated classification protocols are discussed in the censat github: [https://github.com/kmiga/t2t_censat/](https://github.com/kmiga/t2t_censat/)

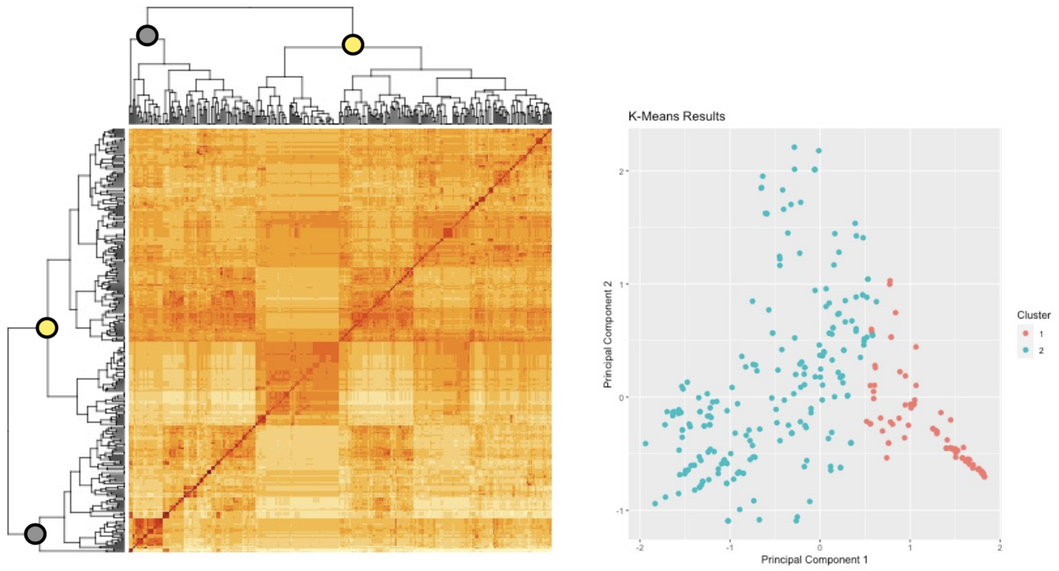

**Figure Legend:** HOR sequences from k-means “group 1” (when classifying into three groups), or the dark red younger HORs in the middle of the array (with group 2,3 flanking closer to p/q arms). Reclassification provides clear evidence for two distinct groups (grey, S01C3H1L-B1 and yellow S01C3H1L-B2), as shown in Figure 3E. Notably, we identify additional substructure within these classifications.

This semi-automated HOR-hap prediction strategy was applied across the genome providing predictions of array structure and organization. The validity of this approach was tested or benchmarked against carefully studied arrays, where all HORs were manually evaluated using phylogenetic trees, discussed below.

Phylogenetic tree evaluation of evolutionarily young HOR-haps

### Phylogenetic tree evaluation of evolutionarily young HOR-haps

#### Analysis of cenX

1. Reverse complement sequence, was prepared of the cenX array (chrX:57,819,765-60,927,196) annotated into S3CXH1L HORs (StV track), cut it into HORs and abnormal HORs were deleted, so that only the full-length HORs remained. Complete HORs were aligned and inspected. There were a number of short

deletions and insertions, some of which appeared in a few copies of a HOR. There were 1475 HORs in the alignment which were analysed further.

2. A minimum evolution tree was made in MEGA5 (*51*). The tree had few well-differentiated branches which corresponded to certain cenX regions and no significant mixing of the HORs from different branches was observed except for yellow and green branches. There was a pure green region on the right side and the region of the yellow-green mix in the center. No pure yellow region was observed.

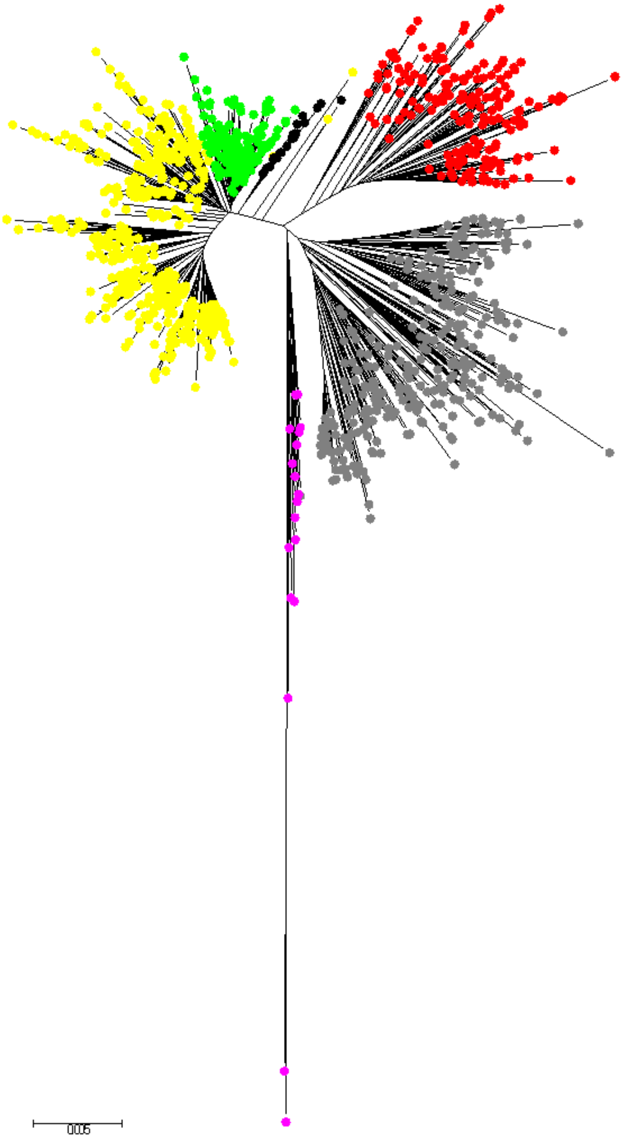

Figure 1. Evolutionary relationships of S3CXH1L haplotypes.

The evolutionary history was inferred using the Minimum Evolution method (*52*). The optimal tree with the sum of branch length = 102.01269531 is shown. The tree is drawn to scale, with branch lengths in the same units as those of the evolutionary distances used to infer the phylogenetic tree. The evolutionary distances were computed using the number of differences method (*53*) and are in the units of the number of base differences per sequence. The ME tree was searched using the Close-Neighbor-Interchange (CNI) algorithm (*53*) at a search level of 1. The Neighbor-joining algorithm (*54*) was used to generate the initial tree. Evolutionary analyses were conducted in MEGA5 (*51*).

3. Next, only the grey branch was extracted and a separate phylogenetic tree was constructed by the same method. A number of well differentiated branches were defined and mapped to the assembly again. All branches occupied separate and well-defined regions in the array as shown in the Hap-track panel below.

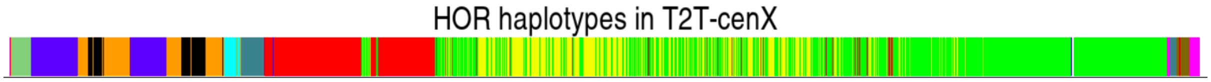

We concluded that the tree was “well regionalized” and that different branches represent different haplotypes of the S3CXH1L HOR, most of them forming at least one compact domain in the assembly.

4. The two minor branches which were marked lilac and black were considered separately. The black branch has 17 HORs interspersed in the yellow region, sometimes in clusters of 2 or 3 copies (eventually termed the YELLOW3 haplotype). Lilac appeared to span the very edges on both flanks (16 HORs on the q-side side and 2 on the p-side). Also, the 2 extra long shoots in the lilac branch were considered specially. They appeared to sit together on the extreme right flank of the array after the lilac HOR-hap. These 2 distal HORs on the q-arm do not have the 15 bp insert characteristic of human DXZ1 (S3CxH1L) (*14*)(Waye and Willard 1985) and have many differences, hence they were considered PRE-LILAC and appear to be represent the ancestor of the entire array linking it to other related SF3 HORs which all do not have the insertion. The third HOR (PLfuse) is a hybrid, to pos  587 it is PRE-LILAC, after it is LILAC. It does have the 15 bp inserts which immediately precedes pos. 587, which is the last PRE-LILAC diagnostic position. So, in fact the insert may already represent the LILAC part (only 1 position contradicts that). The disposition of the described HORs on the right flank is shown in a close-up below.

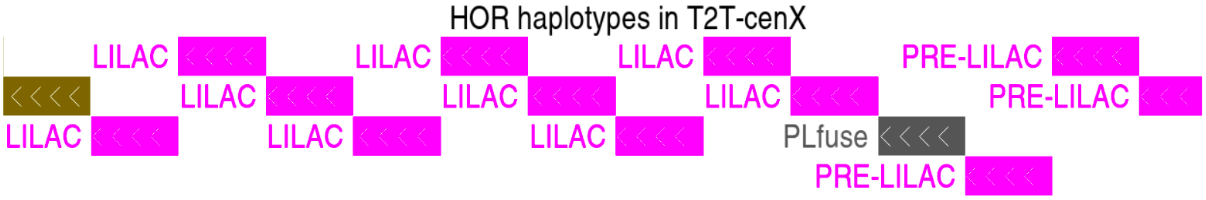

5. Next, the extracted sequences from each branch were installed in HMMER as HMM profiles and automatic haplotyping tool was obtained as described for the HMMER-HOR and -SF tools. It was used to haplotype the chm13 and hg002 cenX assemblies with results shown below in Figure 2A and B. The consensus sequences for all HOR-haps were obtained and the minimum evolution phylogenetic tree of consensus HOR haplotypes was constructed (Fig. 2C) and rooted on the PRE-LILAC haplotype which is known to be the ancestral one as indicated by the absence of 15 bp insertion. It has demonstrated that the YELLOW and GREEN haplotypes were the youngest (more derived ones). The alignment to the methylation track which features the CDR, demonstrated that the kinetochore is located on the young YELLOW2 haplotype in both cell lines.

6. The HOR-hap consensus tree also shows that the haplotypes form 3 groups, named LILAC, GREY and YELLOW (the GREEN therefore was renamed into YELLOWG) which unite the oldest, older and younger HOR-haps, respectively. When multiple sequence alignments used for HMMs were pooled across these three groups, the haplotyping tool and haplotype annotation of reduced resolution were obtained. The reduced resolution patterns have readily visualised the layered expansion pattern within the cenX array (Fig. 2B, upper panel), with both LILAC and GREY older HOR-haps flanking the younger YELLOW on both sides. The statistics for this reduced set of haplotypes is presented in Fig. 2D.

**A**

| Haplotype | CHM13 chrX_v0.7 (score >= 2000) | HG_002 chrX (score >= 2000) |
|---|---|---|
| GREYB | 46 | 55 |
| GREYG | 24 | 24 |
| GREYL | 19 | 11 |
| GREYM | 16 | 17 |
| GREYP | 96 | 168 |
| GREYR | 212 | 206 |
| GREYS | 106 | 97 |
| GREYZ | 37 | 25 |
| LILAC | 15 | 10 |
| PLfuse | 1 | 1 |
| PRE-LILAC | 2 | 2 |
| YELLOW1 | 43 | 29 |
| YELLOW2 | 348 | 327 |
| YELLOW3 | 17 | 8 |
| YELLOWG | 504 | 517 |
| Total | 1486 | 1497 |

**B**

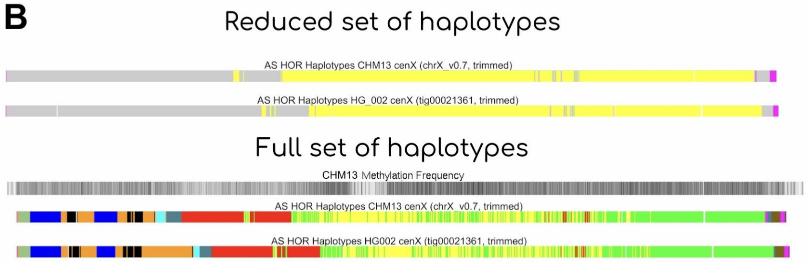

**C**

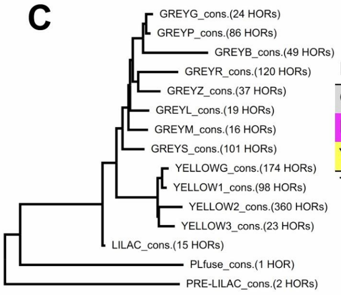

**D**

| Haplotype | CHM13 chrX_v0.7 (score >= 2000) | HG_002 chrX (score >= 2000) |
|---|---|---|
| GREY | 562 | 608 |
| LILAC | 16 | 13 |
| YELLOW | 908 | 876 |
| Total | 1486 | 1497 |

Figure 2. Haplotypes in cenX HORs

7. Finally, the inra-array divergence measurements performed in the multiple alignments for each HOR-hap (shown in Fig 3) demonstrated the decreasing gradient of divergence in PRE-LILAC, LILAC, GREY, LILAC succession.

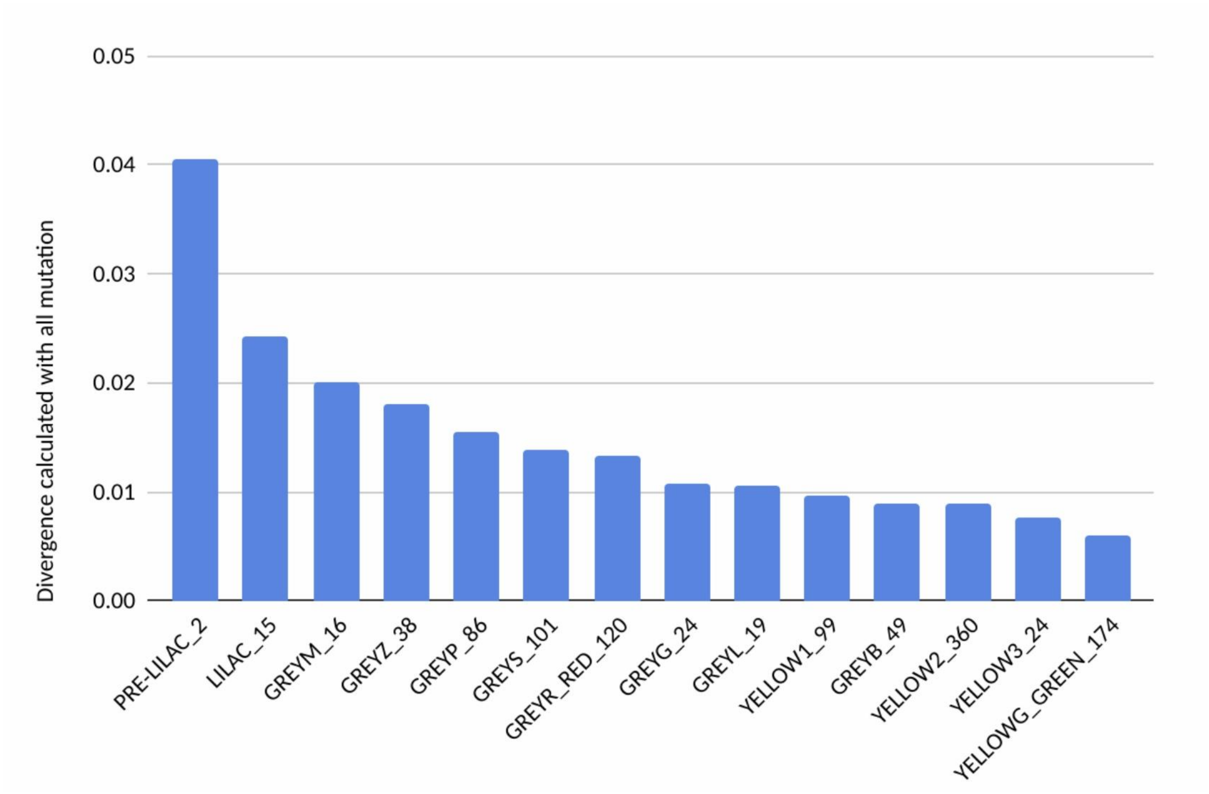

Figure 3. Divergence in cenX haplotypes

Overall, the detailed phylogenetic and divergence analyses in cenX show that expectations of the layered expansion model are met in this centromeric array. This was further confirmed by similar analysis performed for cens 8 and 3.

#### Section 7: CENP-A CUT&RUN and enrichment study

##### CUT&RUN Protocol for CHM13

Thawed Chm13 cells (total passage #18) were washed and resuspended in growth media: DMEM F-12 with 1X Glutamax plus 1x Penicillin/Streptomycin, 10% FBS (Avantor Seradigm Select, VWR 89510-186), 1x ITS, 1x NEAA, and 1x Na pyruvate. Cells were incubated at 37C with 5% CO2 and 100% humidity. Cells were expanded 1:6 every 4 days following detachment with TrypLE Select (Gibco 12563011). When enough cells were obtained, cells were trypsinized, pelleted, and resuspended at 1M cells/ml in Bambanker Standard Serum-Free Freezing Medium (Lymphotec Inc 6801; contains DMSO). Cells were frozen slowly overnight in a Mr. Frosty isopropanol bath at -80C, then transferred to liquid nitrogen for long-term storage. Frozen cells were thawed to room temperature, washed, counted, and then directly harvested for CUT&RUN experiments. HG002 (GM24385 from Coriell) cells were grown in suspension in RPMI-1640 with L-Glutamine, and supplemented with 15% FBS and 1X P/S. Cells were split 1:4 when they reached 1M cells/ml, roughly every 4 days, and they were frozen in RPMI-1640 with 20% FBS and 6% DMSO.

CUT&RUN was carried out as in Thakur and Henikoff 2018 (*55*), with some variations. Frozen pellets of 1,6M HG002 cells or 1,2M CHM cells were thawed on ice and centrifuged at 500g for 5 minutes at 4°C. Cells were washed with cold PBS twice. For nuclear extraction cell pellet was resuspended in 500 mL of Nuclear Extraction Buffer (NEB, 20mM HEPES pH 7.9, 10mM KCl, 0.5mM Spermidine, 0.1% NP40, 20% glycerol, Roche Proteinase Inhibitor tablets) by repipetting gently, and incubated on ice for 5 minutes. Cells were centrifuged and washed with Washing Buffer (WB, 20mM HEPES pH 7.5, 150mM NaCl, 0.5mM Spermidine, 0.1% BSA, 0.05% NP40, Roche Proteinase Inhibitor tablets), blocked in WB containing BSA, and incubated in primary antibody for 2h at 4°C under rotation. Cells were washed twice with WB and incubated with pAG-MNase (Cell Signaling) for 1h at 4°C under rotation. For pAG-MNase digestion, samples were incubated on ice-water (0°C) for 10 minutes, and $CaCl_2$ was added to 2mM as final concentration. Samples were incubated for 30 minutes in cold (0°C). To stop digestion, an equal volume of 2X STOP solution (200mM NaCl, 20mM EDTA, 4mM EGTA, 0.1% NP40) was added. To recover low-salt fragments, samples were incubated for 1h at 4°C under rotation, centrifuged at 500g for 5 minutes and supernatant collected and labeled as low-salt fraction. To recover high-salt fragments, the pellet was resuspended in 150 mL of low-salt solution (175mM NaCl, 20mM EDTA, 4mM EGTA, 0.1% NP40) and then 150 mL of high-salt solution (825mM NaCl, 20mM EDTA, 4mM EGTA, 0.1% NP40) was added. Cells were rotated for 1h at 4°C, centrifuged at 16000g for 5 minutes and supernatant collected and labeled as high-salt fraction. RNase A was added to both fractions following incubation for 20 minutes at 37°C . Samples were treated with Proteinase K for 1h at 65°C (or overnight), and DNA extraction was carried out with MasterPure Complete DNA Isolation kit (Lucigen) as indicated by the manufacturer. Samples were analyzed by a Fragment Analyzer. For library preparation, 1.5 pg of Spike-in Yeast DNA was added (obtained from the Henikoff lab) and NEBNext Ultra II End repair/A-tailing and Ligation kits were used as indicated by the manufacturer. DNA was cleaned using AMPure XP beads and the PCR reaction was carried out using NEBNext UltraII Q5 master mix and NEBNext multiplex oligos for Illumina (12 cycles with annealing/extension for 15 seconds at 65°C). Libraries were cleaned using AMPure XP beads as indicated by the manufacturer. Libraries were sequenced using NovaSeq 6000 150PE Flow Cell SP (minimum 1% PhiX control run). Primary antibodies used were: mouse a CENP-A (Abcam, ab13939), rabbit a CENP-B (Abcam ab25734), guinea pig a CENP-C (MBL, PD030), rabbit a H3K9me3 (Abcam, ab8898).

##### Native CENP-A ChIP-seq analysis

Native CHM13 CENP-A ChIP and input data (*28*) were trimmed with Sickle (https://github.com/najoshi/sickle) to remove low-quality 5' and 3' end bases and Cutadapt (*56*) to remove adapters, as previously described (*28*). Processed data were aligned to the CHM13v1.0 assembly using BWA-MEM (*18*) with the following parameters: bwa mem -k 50 -c 1000000 {index} {read1.fastq.gz} {read2.fastq.gz}. The resulting SAM files were filtered using SAMtools v1.12 (*19*) with FLAG score 2308 to prevent multi-mapping of reads. With this filter, reads mapping to more than one location are randomly assigned a single mapping location, thereby preventing mapping biases in highly identical regions.

#### Reference-free enrichment of k-mer protocols

The reference-free identification of enriched k-mers is based on a pipeline described previously (*57*) and outlined in schematic here. Briefly, 51 bp or 100 bp k-mers were first generated using KMC from trimmed reads (read1 or read2) of the paired-end ChIP-seq or CUT&RUN dataset, for both the dataset of interest as well as IgG control. K-mer counts were then normalized to the total length of the trimmed dataset, and enrichment ratios calculated by dividing normalized k-mer counts of the k-mer in the dataset of interest and IgG control. Enrichment ratio cutoffs were determined as multiples of median absolute deviations away from the median enrichment ratio. Using a scatter plot of normalized k-mer counts in the dataset of interest and IgG, an enrichment ratio threshold is then identified, and all k-mers below the threshold are discarded. Finally, enriched k-mers from reads 1 and 2 are merged, intersected with unique or local k-mers and intersected across experimental replicates.

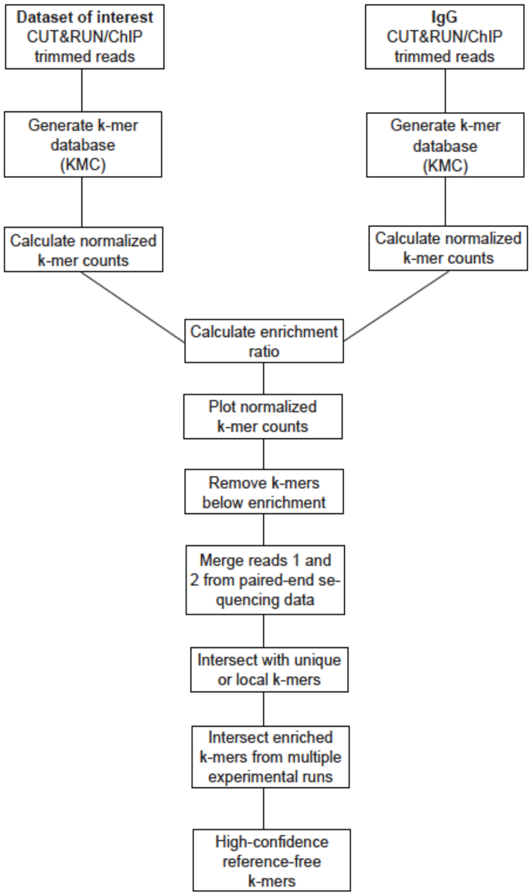

#### Regional K-mers and Enrichment Estimates

Regional k-mers (where k=100 bp), or sequences that are not unique themselves but all copies are organized within a defined local window in the CHM13 genome. For example, if you had a HOR sequence with a variant that is tandemly duplicated (100% identity between adjacent copies), the variant would not be detected by our single copy maker estimates and that region would be ignored by marker-assisted mapping based methods. To address this, we constructed libraries of regional k-mers for every active array in the CHM13 genome, using sliding windows of length 10kb -100kb (with 0.5 slide, ie. 5kb slide for 10kb window). Each array was reformatted and each 100 bp sequence was given a count representing its occurrence (jellyfish, (*58*)). For example, within a given active array, a set of regional 100bp could be observed 3 times. Each window (of length 10kb through 100kb, with windows in 10 kb increments), reported the content of 100-bp and their counts. If the number of counts within a single window exactly matched the total counts for the entire array that set of 100-mers was considered specific to that window, and were noted as providing “regional information”. All regional 100-bp candidates were cross-evaluated genome-wide to ensure no exact matches were identified outside of the active array. Regional kmers were then issued an enrichment value using methods described in “Reference-free enrichment of k-mer protocols”. Bed tracks were then generated across all regional windows (10kb through 100kb), where black indicated windows that lacked regional kmer information, grey indicated windows that contained regional markers, however these markers did not have evidence of enrichment, and orange/red color indicate enrichment with scaling (ie, where scaling represents the number of enriched 100-mers detected within the window and not the enrichment score itself). 100bp regional makers were determined to be enriched if they were >4 fold over (log Transformed ratio of CENP-A data from NChIP or CUT&RUN relative to background).

##### Identifying false positive marker deserts due to multi-mapping

To assess the specificity of reads mapping to regions lacking unique k-mer markers (termed “marker deserts”), reads mapping to these regions were extracted using SAMtools v1.12 (*19*) and remapped to the CHM13v1.0 assembly with the same parameters described above to determine if they align back to their original marker desert region or not. The resulting alignments were filtered using SAMtools v1.12 (*19*) with FLAG score 2308 and converted to a BED file using BEDtools v2.27.1 (*20*). The BED files were used to determine the ratio of reads (or basepairs) re-mapping to the same marker desert region for both CENP-A and input data. Additionally, the BED files were used to determine the enrichment of CENP-A reads (or basepairs) in each marker desert region relative to input data.

##### Marker assisted mapping protocol

CHM13 and HG002 CENP-A CUT&RUN data (IP and IgG) were trimmed with Sickle (https://github.com/najoshi/sickle) to remove low-quality 5' and 3' end bases and Cutadapt (*56*) to remove adapters, as previously described (*28*). Processed data were aligned to the CHM13v1.0 assembly using BWA-MEM (*18*) with the following parameters: bwa mem -k 50 -c 1000000 {index} {read1.fastq.gz} {read2.fastq.gz}. The resulting SAM files were filtered using SAMtools v1.12 (*19*) with FLAG score 2308 to prevent multi-mapping of reads. Marker-assisted mapping of CUT&RUN and NChIP data to the same genome (CHM13 to T2T-CHM13 or HG002 to CHM13 autosomes (chromosomes 1-22), HG002 T2T chromosome X and GRCh38 chromosome Y) was performed using markers of length 21, 51, and 100bp ((*7*, *59*). Following mapping of CutnRun data, follow the steps laid out to retain only those reads that overlap a unique kmer”.

https://gitlab.com/SJHoyt/t2t_transposable-elements/-/tree/main/CutnRun_analyses

**Unique kmer filtering pipeline for CutnRun data**

1: Convert sort.bam to bed for overlapSelect with unique kmers:
Version: bedtools/2.29.0
Command: bedtools bamtobed -i mapped.bam > mapped.bed

2a: Overlap Cut & Run data with Meryl unique 21mers:

Version: GenomeBrowser/20180626
Usage: overlapSelect -overlapBases=[# of bases required to overlap to retain read] unique_kmers.bed reads-aligned.bed reads-aligned.over.unique-kmers.bed
Command: overlapSelect -overlapBases=21 chm13.v1.k21.single.mrg.bed mapped.bed mapped.over.chm13-v1-k21-single-meryl-mrg.bed

2b: Overlap Cut & Run data with Meryl unique 51mers:
Version: GenomeBrowser/20180626
Command: overlapSelect -overlapBases=51 chm13.v1.single.k51.mrg.bed mapped.bed mapped.over.chm13-v1-k51-single-meryl-mrg.bed

2c: Overlap Cut & Run data with unique 100mers (generated by Michael Sauria):
Version: GenomeBrowser/20180626
Command: overlapSelect -overlapBases=100 chm13v1_unique_100mers_reformatted4overlaps.bed mapped.bed mapped.over.chm13-v1-k100-single-mrg.bed

3a: For 21mer overlaps, bed > bedgraph > bigwig for viewing in browser:
Versions: GenomeBrowser/20180626, bedtools/2.29.0
Commands:
bedtools genomecov -bg -i mapped.over.chm13-v1-k21-single-meryl-mrg.bed -g chm13v1.0_seqLengths.genome > mapped.over.chm13-v1-k21-single-meryl-mrg.bedgraph
export LC_COLLATE=C
sort -k1,1 -k2,2n mapped.over.chm13-v1-k21-single-meryl-mrg.bedgraph > mapped.over.chm13-v1-k21-single-meryl-mrg_sorted.bedgraph
bedGraphToBigWig mapped.over.chm13-v1-k21-single-meryl-mrg_sorted.bedgraph chm13v1.0_seqLengths.genome mapped.over.chm13-v1-k21-single-meryl-mrg_sorted.bigwig

3b: For 51mer overlaps, bed > bedgraph > bigwig for viewing in browser:
Versions: GenomeBrowser/20180626, bedtools/2.29.0
Commands:
bedtools genomecov -bg -i mapped.over.chm13-v1-k51-single-meryl-mrg.bed -g chm13v1.0_seqLengths.genome > mapped.over.chm13-v1-k51-single-meryl-mrg.bedgraph
export LC_COLLATE=C
sort -k1,1 -k2,2n mapped.over.chm13-v1-k51-single-meryl-mrg.bedgraph > mapped.over.chm13-v1-k51-single-meryl-mrg_sorted.bedgraph
bedGraphToBigWig mapped.over.chm13-v1-k51-single-meryl-mrg_sorted.bedgraph chm13v1.0_seqLengths.genome mapped.over.chm13-v1-k51-single-meryl-mrg_sorted.bigwig

3c: For 100mer overlaps, bed > bedgraph > bigwig for viewing in browser:
Versions: GenomeBrowser/20180626, bedtools/2.29.0
Commands:
bedtools genomecov -bg -i mapped.over.chm13-v1-k100-single-mrg.bed -g chm13v1.0_seqLengths.genome > mapped.over.chm13-v1-k100-single-mrg.bedgraph
export LC_COLLATE=C

sort -k1,1 -k2,2n mapped.over.chm13-v1-k100-single-mrg.bedgraph > mapped.over.chm13-v1-k100-single-mrg_sorted.bedgraph
bedGraphToBigWig mapped.over.chm13-v1-k100-single-mrg_sorted.bedgraph chm13v1.0_seqLengths.genome mapped.over.chm13-v1-k100-single-mrg_sorted.bigwig

**Filtering through Meryl unique CHM13v1.0 21mers and 51mers:**

- For marker-assisted alignment filtering, we removed alignments that did not span any "marker" k-mer as defined in https://doi.org/10.1101/2021.07.02.450803 for k=21. Similarly, for k=51, k-mers present once in T2T-CHM13v1.0 and between 28 and 107 in the Illumina reads were used.
- Meryl unique kmer generation code: https://github.com/arangrhie/T2T-Polish/tree/master/marker_assisted
- Meryl unique CHM13v1.0 21mers and 51mers: https://s3-us-west-2.amazonaws.com/human-pangenomics/index.html?prefix=T2T/CHM13/assemblies/alignments/marker/

**Filtering through unique CHM13v1.0 100mers**

- Unique CHM13v1.0 100mer generation code: https://github.com/msauria/T2T_Kmer_Analysis
- Unique CHM13v1.0 100mers: https://github.com/msauria/T2T_Kmer_Analysis/blob/main/results/chm13v1_100mer_coverage.bed

Methylation analysis of minor CDRs

CpG methylation values were calculated according to methods outlined in Gershman et al. Methylation frequency for methylation map plots was generated by calculating the fraction of methylated reads to total coverage within 10kb bins in CHM13 with the BSGenome Bioconductor package (https://bioconductor.org/packages/BSgenome). Multiples of three bins were further smoothed with the "rollmean" function from the R package Zoo (https://cran.r-project.org/web/packages/zoo/ index.html). The depth panel represents the number of aligned nanopore reads containing at least one high quality CpG methylation call. Single read plots were generated in IGV using Nanopore_methylation_utilities (https://github.com/timplab/nanopore-methylation-utilities) to integrate methylation information into the alignment BAM file for viewing in the bisulfite mode in IGV.

#### Section 8: Population study of haploid X chromosomes

##### DXZ1 array characterization of HPRC haploid DXZ1 assemblies

Chromosome X haplotype phased assemblies (trio-hifiasm, (*60*)) from six diverse individuals (1000 genomes LCLs) were obtained from the Human Pangenome Reference Consortium (*36*) (https://github.com/human-pangenomics/hpgp-data). Assemblies were constructed using the following workflow:

https://dockstore.org/workflows/github.com/human-pangenomics/hpp_production_workflows/TrioHifiasmAssembly:master?tab=info

Resulting hifiasm assemblies were mapped against CHM13v1.0 using winnowmap (*22*). Contigs spanning the X centromere were identified using samtools (view, “chrX:57820107-60927026”) (*19*). As shown in the summary table below, four large contig assemblies (HG01109, HG01243, HG03098, HG03492) represented the entire DXZ1 array in a single contig with extensions into the p-arm and q-arm. TandemQuast (*61*) evaluation of these four assemblies did not reveal any structural or base-level errors (Figure S16). The remaining two assemblies, HG02055 and HG02145, were defined by 4 and 2 contig assemblies respectively. HG02145 contigs were merged in order, where contigs anchored on either the p-arm or q-arm and had a single break in the middle. HG02055 represented four contigs, with two larger contigs that anchor in either p-arm or q-arm. The two smaller contigs were concatenated end-to-end in the ordering relative to the CHM13v1.0 alignments. In both cases, contigs were concatenated together without assumption of overlap nor confirmation of structural accuracy at site of contig merger (Table below).

**Table Legend:** Description of six HPRC DXZ1 assemblies representing individuals from Puerto Rico (PUR, American), African Caribbean in Barbados (ACB, African), Mende in Sierra Leone (MSL, African), and Punjabi in Lahore, Pakistan (PJL, South Asian). Maternal haplotype phased contig numbers range from 1-4. The contigs identified to span the DXZ1 centromere array in CHM13 are provided with contig ID, start (S) and end (E). The observed length from the assembly is compared to the approximate sizing (discussed below: DXZ1 array size estimates in XY diversity panel ), using ~30x coverage from 1000 Genomes Illumina data (*62*, *63*).

| ID | POP | S_POP | Mat Contigs | DXZ1 contig ID | contig S | contig E | Array Size (Mb) | ILMN Predicted Array Size (Mb) |
|---|---|---|---|---|---|---|---|---|
| HG01109 | PUR | AMR | **1** | HG01109#2#h2tg000048l | 5823121 | 8408816 | 2.59 | 2.20 |
| HG01243 | PUR | AMR | **1** | HG01243#2#h2tg000168l | 3546383 | 6515118 | 2.97 | 2.42 |

| | | | | | | | | |
|---|---|---|---|---|---|---|---|---|
| HG02055 | ACB | AFR | 4 | HG02055#2#h2tg000121l | 4776941 | 4930321 | 0.15 | |
| | ACB | AFR | 4 | HG02055#2#h2tg000281l | 1 | 88728 | 0.09 | |
| | ACB | AFR | 4 | HG02055#2#h2tg000147c | 1 | 61588 | 0.06 | |
| | ACB | AFR | 4 | HG02055#2#h2tg000097l | 1 | 2027495 | 2.03 | 1.90 |
| HG02145 | ACB | AFR | **2** | HG02145#2#h2tg000272l | 839805 | 1764556 | 0.93 | |
| | ACB | AFR | **2** | HG02145#2#h2tg000225l | 1 | 1972981 | 1.97 | 2.47 |
| HG03098 | MSL | AFR | **1** | HG03098#2#h2tg000077l | 4787896 | 6371416 | 1.58 | 1.49 |
| HG03492 | PJL | SAS | **1** | HG03492#2#h2tg000168l | 456931 | 4236957 | 3.78 | 3.56 |

To identify the start and end of all HOR in these assemblies, we used the consensus sequence from the CHM13v1.0 DXZ1 assembly (in both forward and reverse orientation) to generate a HMMER profile (v3.0) (*64*) and performed an alignment search against each assembly. We used the resulting DXZ1 start and end coordinates to populate a bed file, and used bedtools (*20*) (getfasta) to issue a HOR fasta file for each assembly. In total, we had a database of 9104 HOR (12-mer) fasta (all ordered in the same 5'->3' orientation) representing all six HPRC assemblies and CHM13. We generated pairwise alignment (needle, EMBOSS (*48*))against the CHM13 DXZ1 consensus sequence. Each alignment was reformatted into an ordered string of 0's and 1's, where 0 indicates a match to the shared position with the consensus and 1 if there is a difference at that position relative to the consensus. These tables were then read into r (*49*) and plots were made using ggplot2 (*50*). Prediction of the best "k" clusters used "FitKMeans"(max.clusters=20,nstart=25,seed=278613), using Hartigan's Number (visualized using "HartiganPlot") to determine if the k+1st cluster should be added. Seven k-cluster (using kmeans, centers=7) was selected used in this analysis due to support in Hartigan Plot, k-means assignment with PCA plot shown below, and comparisons with hierarchical clustering of the same data (not shown).

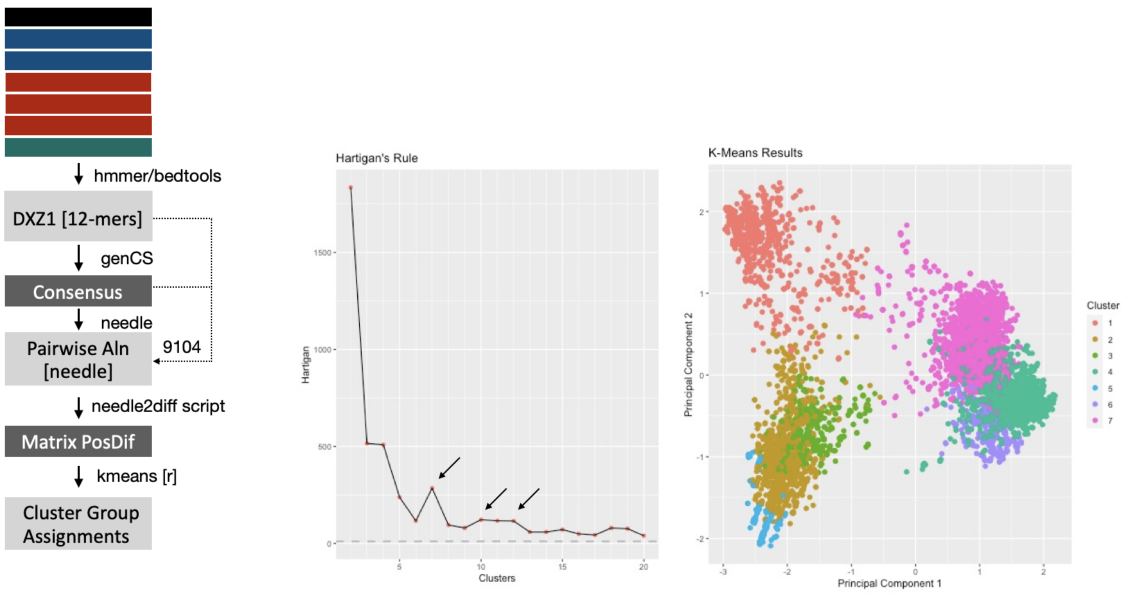

**Figure Legend.** HOR (12-mers, roughly 2057 bp) from all contigs were identified (represented in top left as stacked, multicolored rectangles. Multiple alignment of all HORs (combined in a single file representing all six individual assemblies plus CHM13) was made using kalign (*47*) to provide a consensus sequence. This consensus sequence was then used in pairwise alignments to reformat each sequence into a binary string, where 0's represented a match to the consensus and 1's represented a position of a mismatch. This matrix (rows: each HOR x column: each position relative to consensus populated with either a 0 or 1), was used to estimate k-means clusters. Hartigan plot and k-means PCA plot are shown.

The resulting classification assignments were used to label each HOR coordinate across the arrays. In parallel, direct pairwise comparisons between arrays were made using dot plots (self dot plots are shown in Figure S_, and between CHM13 and each assembly, or merged contig assemblies. Interestingly, comparisons between three contigs: HG0123, HG03098, and HG02055 revealed very little similarity with CHM13 by pairwise comparisons, yet these contigs appeared to be more closely related to one another (as shown below). These contigs are observed to contain HOR-haplotype not identified in CHM13 (shown as dark red in Figure 5a). Methods describing dot plot generation are below.

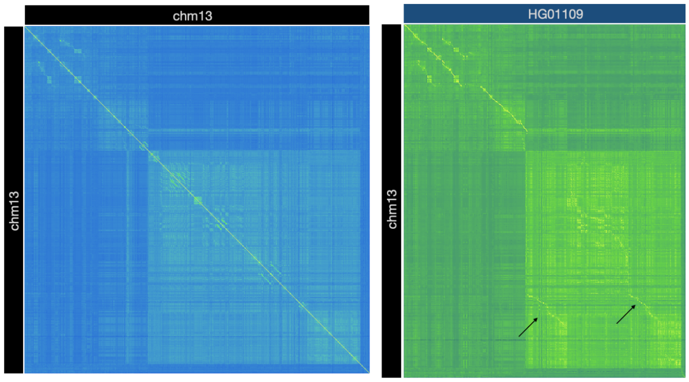

**Figure Legend. Dot plot Comparisons** Self dot plots (shown CHM13 vs CHM13, with other Figure S_) and comparisons against CHM13 were used to survey regions of consistent and inconsistent array structure. Tandem duplication in HG01109 relative to CHM13 is indicated by black arrows. SImilar large clustering of HORs into two groupings can be observed, in support of k=2 clusters. Macro satellite patterns (p-arm proximal) are observed to be shared across assemblies.

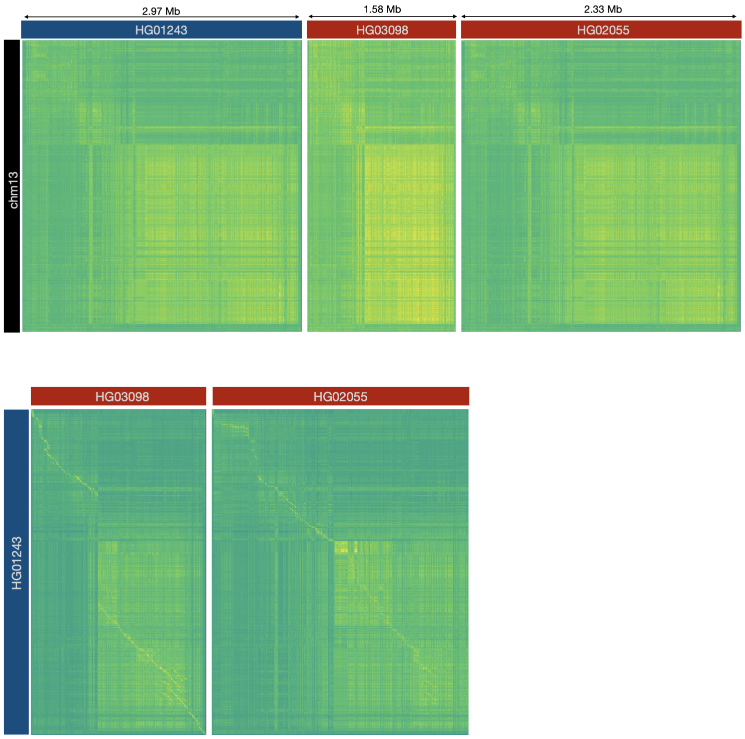

**Figure Legend. Dot plot Comparisons** revealed that three assemblies had very little in common with CHM13 and were more similar to each other (when switching the CHM13 assembly for HG01243 as shown).

#### Dot plots generated by an exhaustive search for exact matches between any two sequences

Dotplots were generated by computing an exhaustive search for exact matches between any 2 sequences, where matches are >= 16bp. A 2D matrix then stores the length of the longest match within square bins that are 1/nth the length of the longest sequence, where n is the number of pixels desired in the output image. The values in the matrix are then log scaled and normalized from 0-1 where 1.0 is the longest observed match. The values are interpreted as a color in the Viridis colormap and rendered as an (n,n)px image.

The code is available at https://github.com/rlorigro/simple-dotplot

**Pairwise sequence identity dot-plots**

To generate pairwise sequence identity dot-plots we used the software package StainedGlass. The input for this program is sequence fragmented into windows (5 kbp) after which all possible pairwise alignments between the fragments are calculated using minimap2 (*65*). The color used in the dot-plot was then determined by the sequence identity of the alignment which was calculated as: $$ ID = 100 \biggl( \frac{M}{M+X+I+D} \biggr) $$ where $ID$ was the percent sequence identity, $M$ the number of matches, $X$ the number of mismatches, $I$ the number of insertion events, and $D$ the number of deletion events. When there were multiple alignments between the same two sequence fragments all alignments other than the one with the most matches were filtered out regardless of their sequence identity. The resulting matrix of percent identity scores was then visualized using ggplot and geom_tile.

All code and documentation is available at: https://mrvollger.github.io/StainedGlass/

**Prediction of recent repeat expansions using dot-plots**

Prediction of recent repeat expansions was performed by making and analyzing the dot-plots. The dot-plots were generated with GENOME PAIR RAPID DOTTER (GEPARD https://cube.univie.ac.at/gepard). The aligned sequences were either two identical sequences or the sequences representing the same AS array in different individuals or cell lines (e.g. CHM13 vs HG002). The word length was chosen to be about equal to the HOR length in the array. Therefore, the dot was placed in the plot if entire HORs were identical in both sequences. The window size was 1 bp. The self alignment dot-plot has the main diagonal and some extra diagonals which represent highly identical (nearly 100% identical) internal repeats in the sequence (recent duplications or multiplications). So, various patterns of short diagonal pieces around the main diagonal which we observed in the plots were interpreted as the signs of recent amplification activity. Such patterns were usually observed in under-kinetochore areas (18-19 of 23 chromosomes), but by no means exclusively in these areas. If we needed to interrogate the lower identity sequence relationships in an array, the word length was made gradually smaller. The word length 500 usually reveals the HOR haplotype patterns which may be used for comparison.

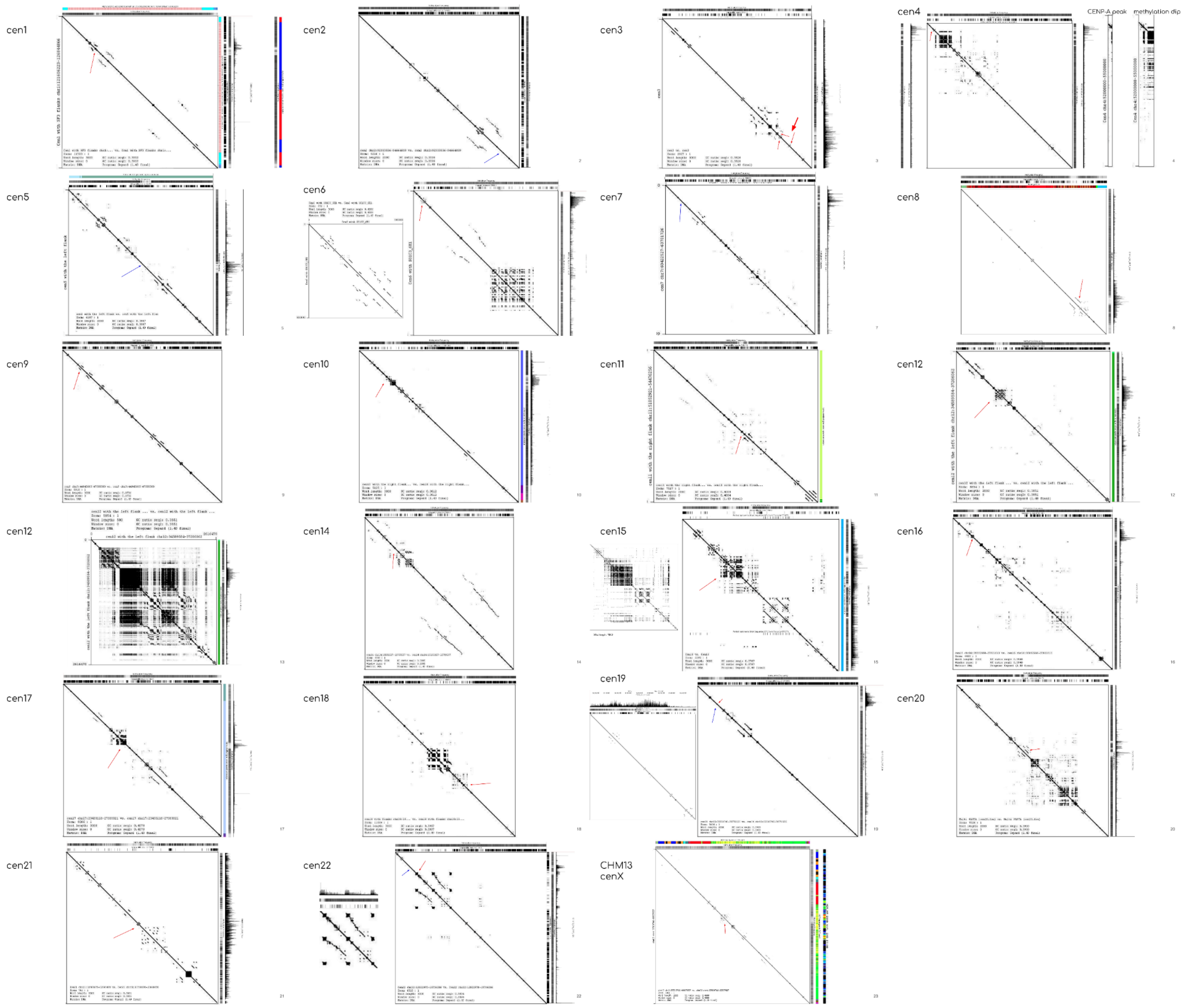

**Figure Legend:** The dot-plots of all live centromeres were generated. As needed, the close ups of under-kinetochore areas or other regions were generated and aligned to haplotype tracks and/or StV tracks with their numbered HORs to allow the fine analysis of the region. By such examination of the cenX close up dot-plot the figure ”Detailed alignment of highly identical under-kinetochore repeats in chm13” was generated.

#### X Centromere HOR-hap proportion estimates in 1000 Genomes WGS Data and Epiallele ChIP-Seq & CUT&RUN datasets.

We took the following steps in characterizing the HOR-haps of each DXZ1 array:

1. **Identification of 51-mers (or 21-mers in case of epiallele data) that are specific to each HOR-haplotype.** Here we used HOR-haps of the X centromere to expand studies of population

diversity (where, k=7 in studies of 1000 Genomes Data, (*62*, *63*)) and epiallele variation (where, k=13 in studies of publicly available CENP-A ChIP-Seq and CUT&RUN experiments. In both applications overlapped in method development. Here we describe the process using k=7 as an example. First, we generated separate DXZ1 (12-mer, canonical) fasta files for each HOR-hap classification (groups 1-7). Each fasta file was reformatted to represent all possible 51 bp sequences (including both strands). Therefore, each HOR-hap classification (groups 1-7) had a file of total 51-bp sequences. Then for each HOR-hap specific 51-bp listing we compared to all other HOR-haps to identify and exclude any exact matches. These identified matches, or exact 51-bp sequences shared between different HOR-hap groupings were determined to be non-specific. All 51-bp sequences are considered 'specific' if they are only found within one HOR-hap grouping. We then estimated the relative frequency of the specific 51-bp markers relative to the total number of markers per HOR-hap grouping (Table 1).

**Table 1**: Listing of specific kmers per each HOR-hap and relative frequency.

| HOR-hap | Specific 51-bp | Total 51-bp | Frequency of Specific 51-mer |
|---|---|---|---|
| 1 | 401704 | 4273414 | 0.0940007216712446 |
| 2 | 689248 | 5441910 | 0.126655530870595 |
| 3 | 282258 | 2596656 | 0.108700574893247 |
| 4 | 231380 | 8641142 | 0.0267765533768569 |
| 5 | 42440 | 858206 | 0.049451996373831 |
| 6 | 50254 | 2500178 | 0.0201001688679766 |
| 7 | 430266 | 12224282 | 0.0351976500542118 |

2. **Method optimization using CHM13 controlled short read datasets.** We used short read data in the CHM13 genome to test and develop our method for reporting HOR-hap proportions. Using the CHM13 X-array HOR-haps (shown below for k=13 classification), we identified specific 51-mer libraries for each grouping as described above and generated a corresponding frequency table. Corresponding paired read DXZ1 libraries were generated either by using alignments to CHM13 PCRFree Illumina alignments (generated via bwa mem v0.7.15 (*18*)) (md5: bb41008d0f5de787d26896fb49027420, https://github.com/marbl/CHM13; where centromere X bams were obtained using samtools (view option with "chrX:57820107-60927026") ) or using simulated short read data at 30x coverage (to match the expected average coverage of the 1000 Genomes Alignments). Simulated Illumina data was obtained using ART Simulation software (*66*); "ART_Illumina (2008-2016), Q Version 2.5.8 (Jun 6, 2016)"; https://www.niehs.nih.gov/research/resources/software/biostatistics/art/index.cfm) (*66*). Next, we screened through each Illumina data set and counted how many times we observed an exact match

to a specific HOR-hap 51-mer. These counts were divided by the observed frequency for specific 51-mers in the CHM13 X-assembly. That is, if (i) the specific 51-mers in HOR-hap group 1 represent 0.082 of the total 51-mers, and (ii) there are 551484 total counts for group 1 in the centromere X reads from the PCRfree Illumina dataset, then group 1 was given the adjusted value of 6727434.013. All adjusted values were added together (group 1 through 13), and then the relative frequency was determined for each. For example, when group 1's adjusted value would be divided by the sum of all adjusted values we observed the value of 0.0111182081482941 (shown in Figure below), with all proportions adding up to 1 as expected. The results of this analysis using Simulated reads, CHM13 Illumina PCR-free alignment data are plotted below in comparison of the expected HOR-hap frequencies determined using the full HOR dataset.

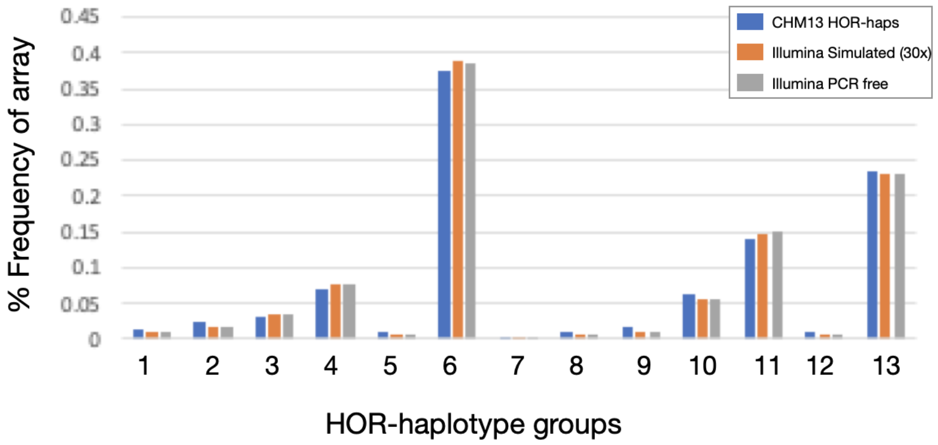

3. **Generated a DXZ1 fasta database for 1599 individuals from the1000 genomes.** 1000 genomes data was aligned to the CHM13v1.0 assembly as described (*62*). Centromere X specific bams were generated for each individual using samtools (view, "chrX:57820107-60927026") (*19*), and bedtools (bamtofastq) (*20*). Paired read data were combined and each fasta was screened for exact matches to specific 51-bp markers for the HOR-haps (classified into seven groups), and the frequency is adjusted using the HOR proportions (Table 1), using method described above. The resulting proportions used in our cenHap classification can be referenced in Table S15.

##### DXZ1 array size estimates in XY diversity panel

41mers which occur exclusively in the DXZ1 array of the CHM13 reference genome were identified. To ensure that we were not including infrequent kmers that are the result of SNPs, and to improve the speed of the pipeline, we only included kmers that occur at least 1000 times within the DXZ1 array. The coverage of array kmers was normalized to a second set of kmers which were identified in a 15 mb region of the X-chromosome.

All kmers chosen for the normalization set occur just a single time in the CHM13 genome (single copy kmers). The 15 mb region was chosen because it is rich in these single copy kmers.

Array kmers were matched to a number of single copy kmers equal to their frequency of occurrence in the array (i.e. an array kmer which occurs 1001 times was matched to 1001 different single copy kmers). To account for sequencing bias based on G/C content, two steps were taken during kmer matching. First, the 15 mb normalization region was split into 10 kb chunks, and only chunks with GC-content within 3% of the mean DXZ1 GC value were considered. Second, the array kmers were matched to single copy kmers based on the G/C content of the kmer itself.

Whole genome sequencing reads were aligned to the X-centromere and the control region, then kmers from each set were counted in the reads. Before performing the array size calculation, a filtering step was necessary for the single copy kmers; while these kmers were identified as single copy in the CHM13 genome, this was not necessarily true for individuals in the 1000 Genomes Project. Single copy kmers which had counts of 0 were assumed to not occur in the genome and were removed. The resulting set of counts was approximately normally distributed, and any kmers which occurred greater than 3 standard deviations from the mean were removed. The array kmer set was then downsized to match the number of valid single copy kmers. Array size was finally estimated by taking the sum of counts of array specific kmers divided by the sum of counts of single copy kmers, then multiplied by the array size in CHM13.

##### TandemTools evaluation of HPRC_PLUS genomes

TandemTools (*61*)was ran on the active chrX alpha satellite HOR array DXZ1 of 4 HiFi read assemblies where HOR array was assembled in a single contig (HG01109, HG01243, HG03098, HG03492). PacBio HiFi reads were mapped to the assembly to perform a search for assembly errors and heterozygous sites. TandemTools compares distances between consecutive rare *k*-mers in the assembly and read mappings (here, *k* = 301 and frequency of a *k*-mer in the assembly do not exceed 20). The analysis did not reveal any consistent discrepancies between the read-set and assemblies except a low covered heterozygous site in the HG01109 assembly.

### Supplementary Figures

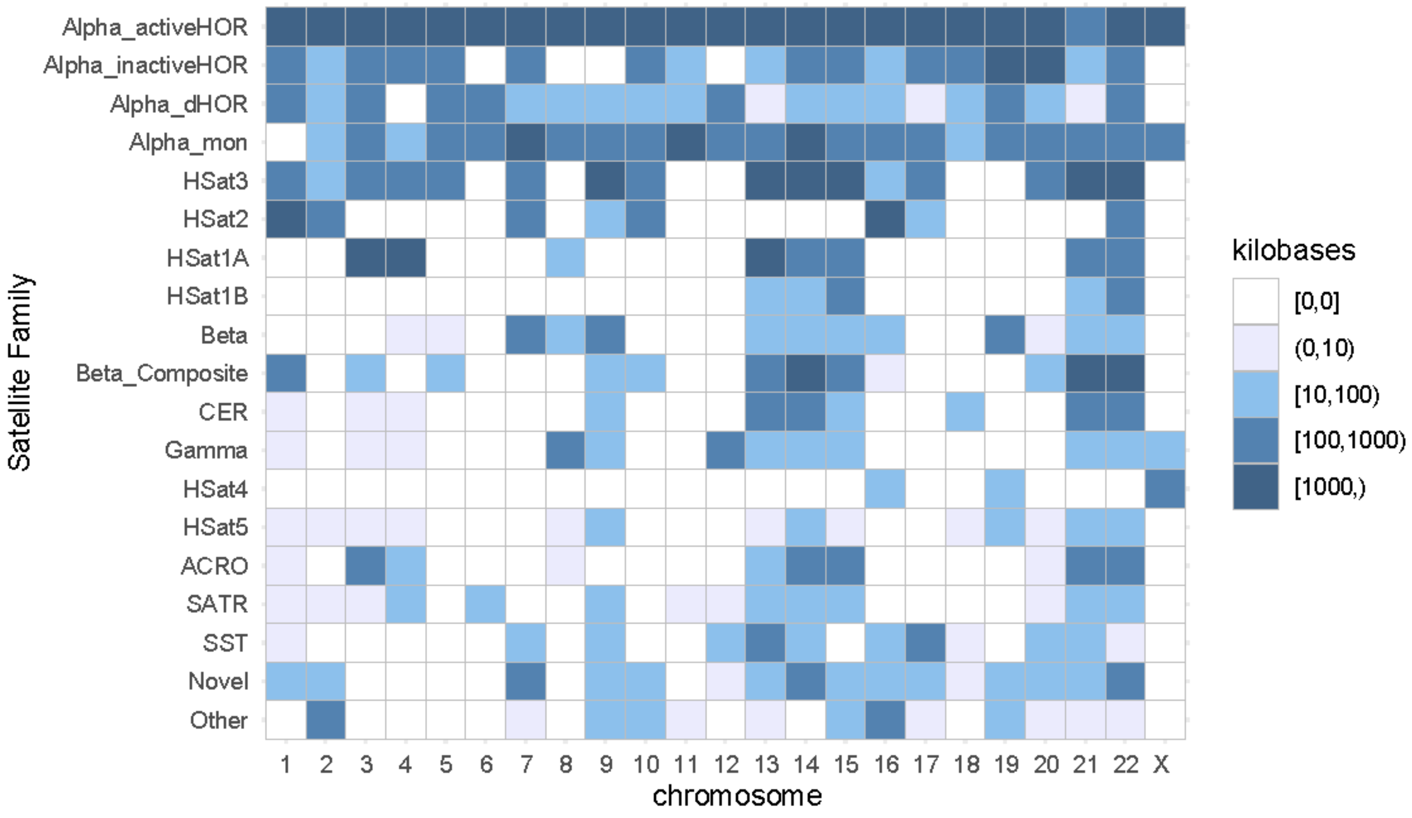

**Figure S1. CenSat chromosome abundance.** Summary of centromere satellite (censat) annotation in CHM13v1.0 with colors indicating the total kilobases for each annotated class across each chromosome.

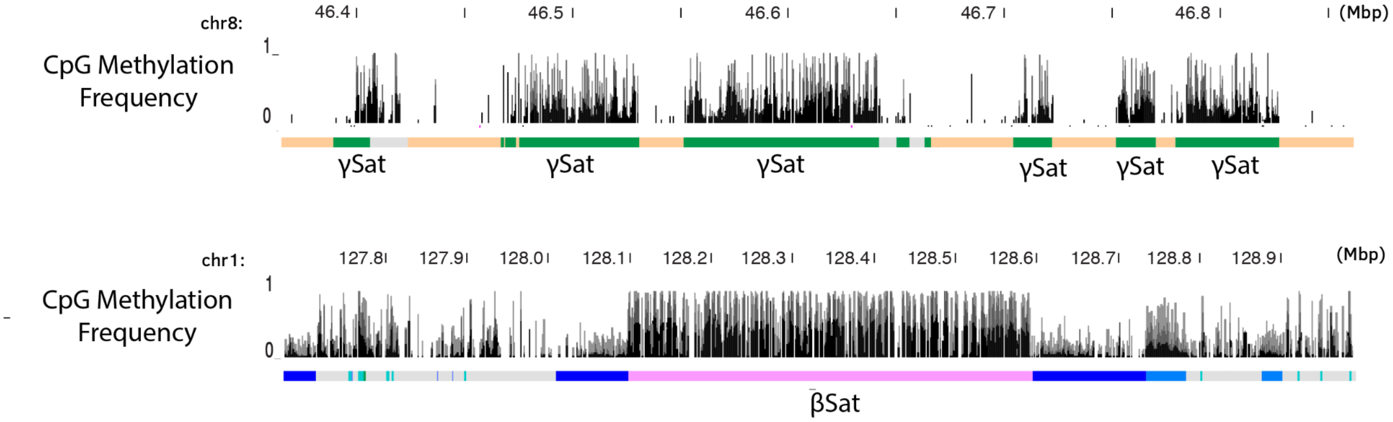

**Figure S2. βSat and γSat contain dense CpG methylation.**

βSat and γSat represent are more GC-rich than other satellites (βSat, 52%; γSat, 72%) and are observed to have dense CpG methylation, as shown for γSat (green) on chromosome 8 ( CHM13v1.0;

chr8:46,365,292-46,861,749), which is found interspersed with monomeric αSat (αSat GC-rich 39%). Additionally, methylation is observed within sites enriched with βSat-LSAU (as shown on CHM13v1.0; chr1:127,674,906-128,992,099). Dark and light blue indicate HSat2,3 classes, grey are centric transitions (non-satellite DNAs), and the low-frequency pericentromeric satellites are represented by teal annotation (chr1 only).

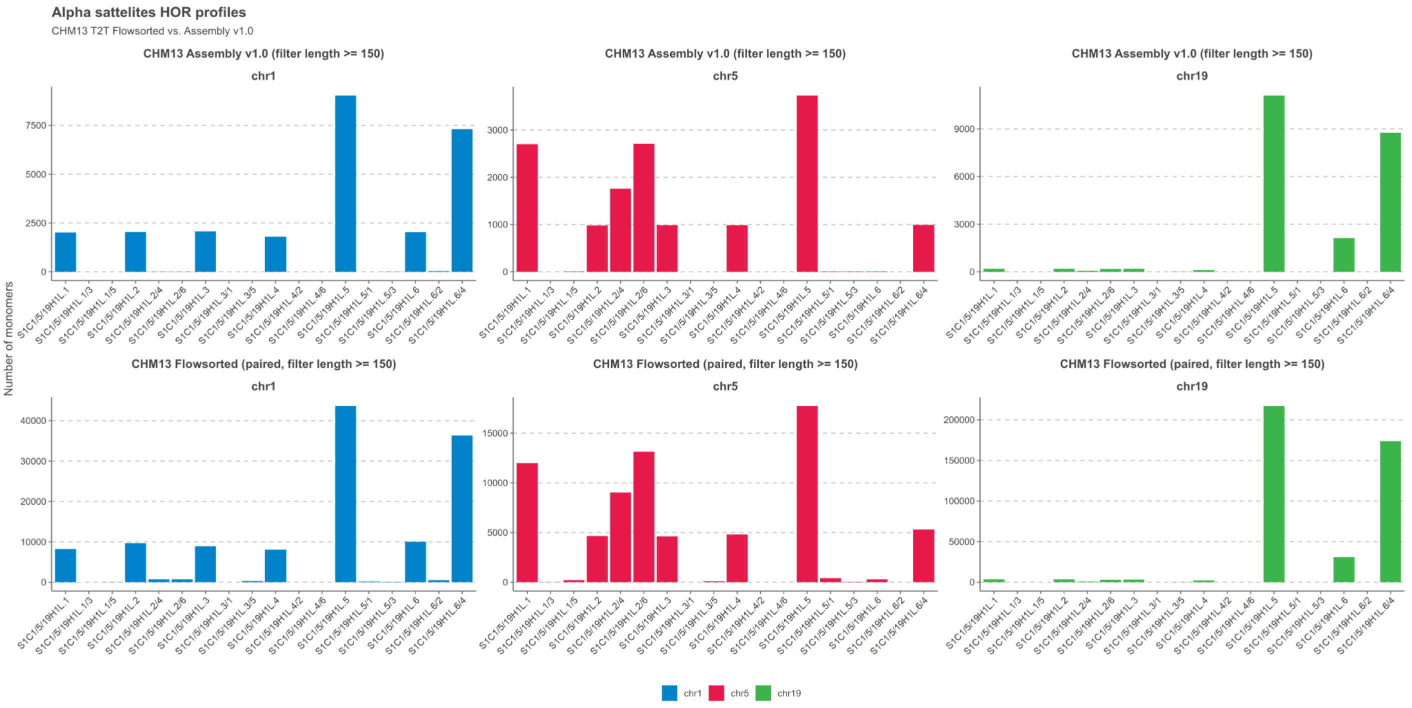

**Figure S3. HOR 1/5/19 arrays are chromosome-specific with reference to flow-sorted chromosome libraries** The figure shows the counts of various monomers (including many hybrids) of AS active array HOR S1C1/5/19H1L in centromeres of three chromosomes (1, 5, and 19), calculated from HOR annotation of CHM13 effectively haploid 22XX cell line complete T2T assembly v1.0 and the same counts performed in Illumina short read libraries prepared from flow-sorted chromosomes 1, 5 and 19 obtained from the same cell line. Monomer profiles appeared to be chromosome-specific and there is a good concordance between the assembly and the flow-sorted material. It suggests that the S1C1/5/19H1L sequences from different chromosomes were correctly segregated into assembled arrays.

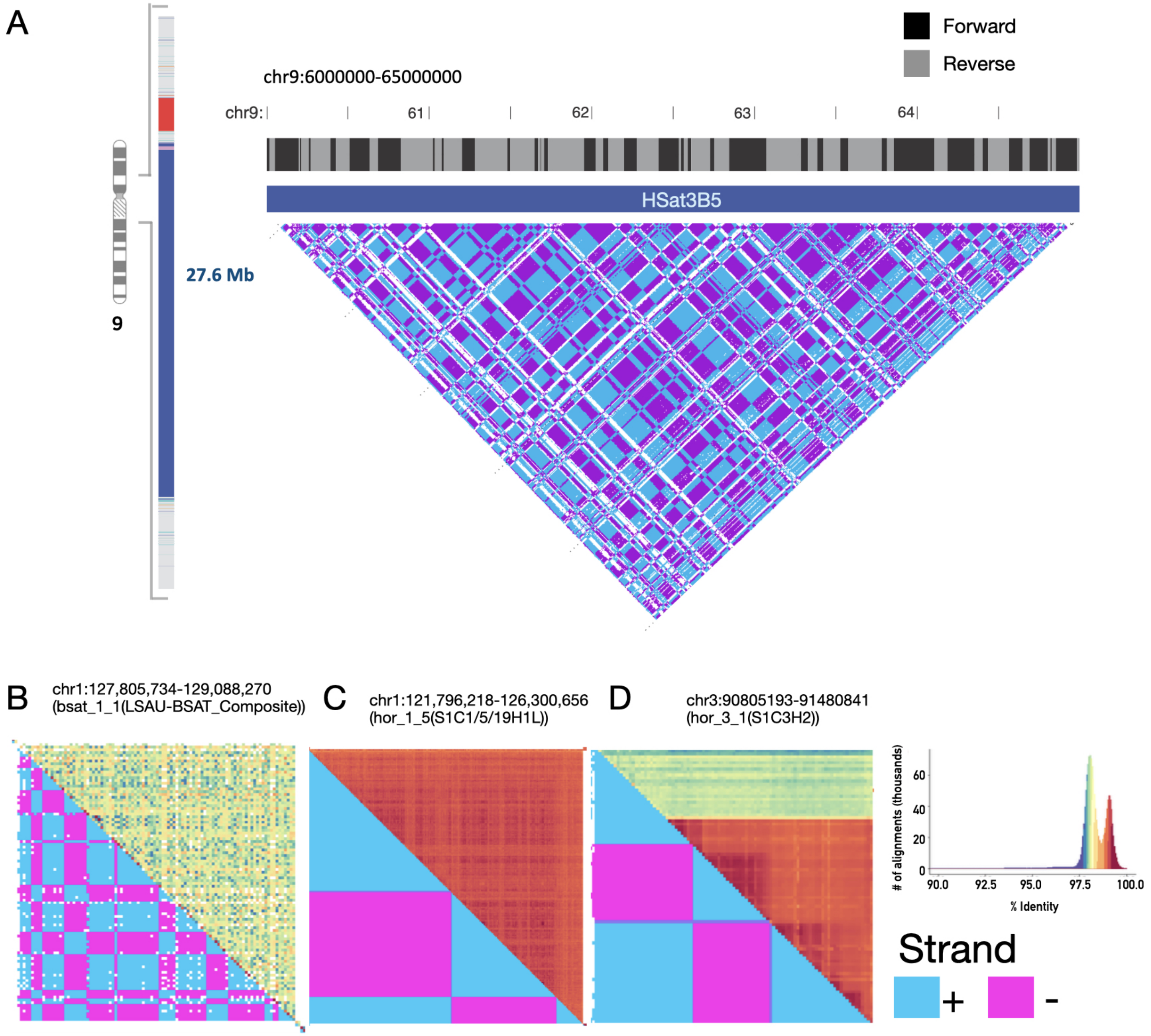
A
Forward
Reverse
chr9:6000000-65000000
chr9:
61
62
63
64
HSat3B5
27.6 Mb
9
B
chr1:127,805,734-129,088,270
(bsat_1_1(LSAU-BSAT_Composite))
C
chr1:121,796,218-126,300,656
(hor_1_5(S1C1/5/19H1L))
D
chr3:90805193-91480841
(hor_3_1(S1C3H2))
### of alignments (thousands)
% Identity
Strand
+
-

**Figure S4. Inversions are identified within many homogeneous arrays of different satellite families (HSat, βSat, and αSat).** Shifts in satellite repeat orientation (as shown in blue, +-strand, and purple, -strand, are observed across several larger arrays in the CHM13 genome. A. The large pericentromeric HSat3B5 (represented by a 5Mb zoom in array on chr9:60-65Mb) is observed to have alternating orientation throughout (dotplot: https://mrvollger.github.io/StainedGlass/, and shown by bed track reporting regions by prevalence of CATTC (Forward) vs GAATG (Reverse)). Inversions are also detected in Beta satellites-LSAU composite arrays (also represented in Figure 2a), active centromere array on chromosome 1 (D1Z7 or S1C1/5/19H1L), and an inactive array on chromosome 3 (S1C3H2).

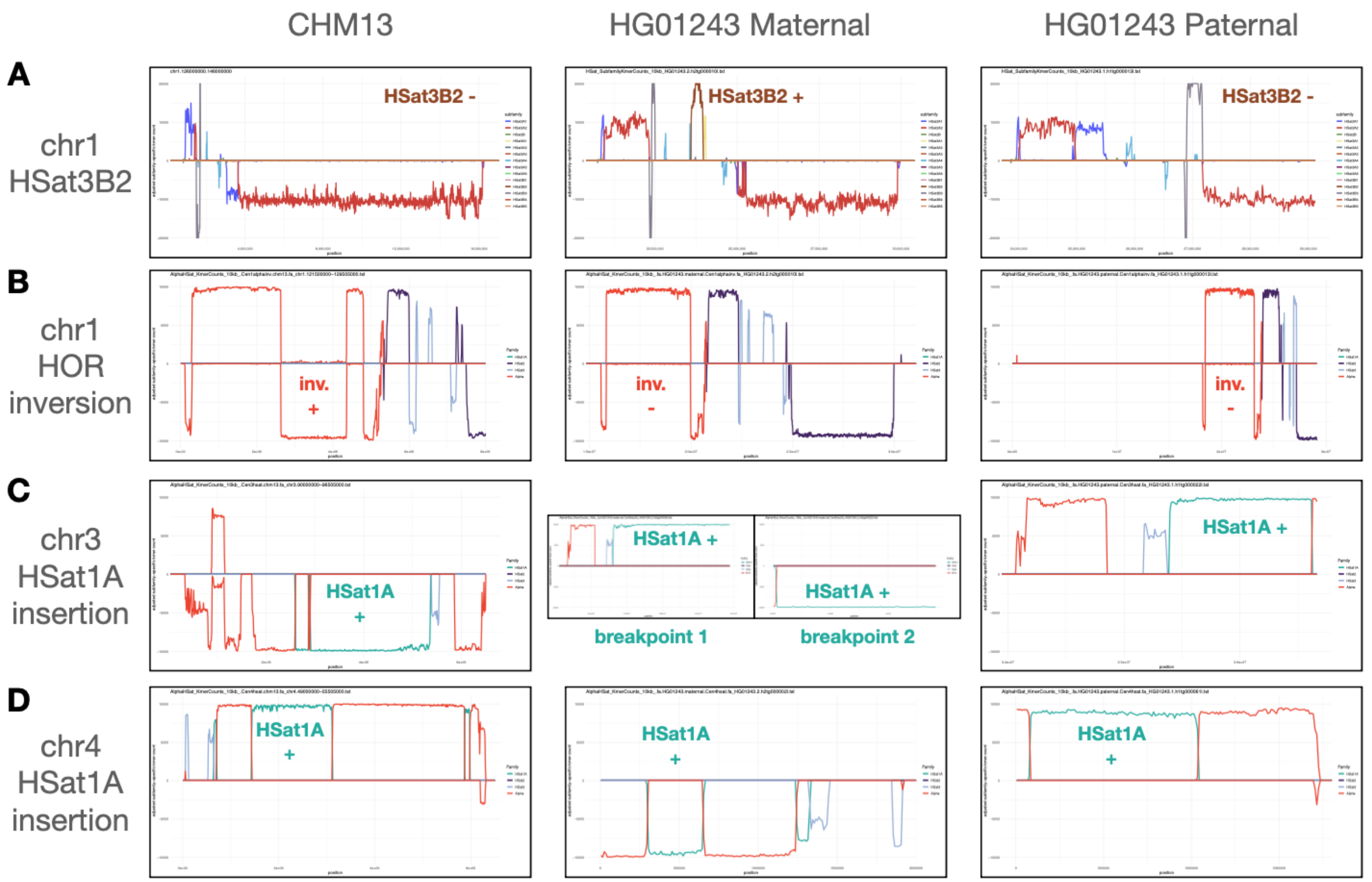

**Figure. S5. Painting hifiasm contigs to reveal satellite in/del and inversion breakpoints. A**) Plots show, for the contig overlapping the chr1 Alpha-HSat2 transition region, the relative abundance of HSat2,3 subfamily-specific 24-mers (each of the 14 subfamilies is plotted in a different color, indicated in plot legend. Lines above the x axis represent 24-mers found on the + strand, and those below the axis represent 24-mers found on the - strand. The brown line corresponds to HSat3, subfamily B2, which is absent in CHM13 and in the paternal haplotype of HG01243, but present in the maternal haplotype. Plots are zoomed into the leftmost and rightmost coordinates with matching 24-mers. **B**) As in (A), but painting using 21-mers that broadly distinguish Alpha, HSat1A, HSat2, and HSat3. Here CHM13 is positive for the large inversion in the

D1Z7 alpha array, but HG01243 is not. Note: the regions of alpha on the (-) strand on the flanks of the array represent a different HOR type, D1Z5. The inversion phenotype classification requires that the assembly span between these D1Z5 flanks, and show a block of intervening D1Z7 on the (-) strand as well. **C-D**) As in (B) but for the HSat1A insertions on chrs 3 and 4. Classification required that both insertion breakpoints be detected, even if on separate contigs, as for the HG01243 maternal haplotype.

| cell line | chr1 HSat3B2 | | | chr1 HOR inv. | | | chr3 HSat1A ins. | | | chr4 HSat1A ins. | | |
|---|---|---|---|---|---|---|---|---|---|---|---|---|
| | P | M | coords | P | M | coords | P | M | coords | P | M | coords |
| CHM13 | - | | chr1:126700000-144000000 | + | | chr1:125850000-126350000 | + | | chr3: 92850000-95400000 | + | | chr4:50400000-52150000 |
| HG005 | - | - | h1tg000101l, h2tg000044l | - | ? | h1tg000002l, h2tg000044l | + | + | h1tg000018l, h2tg000009l | + | + | h1tg000011l, h2tg000008l, h2tg000061l |
| HG00733 | + | + | h1tg000154l, h2tg000050l | ? | + | h1tg000129l, h2tg000155c | + | + | h1tg000048l, h2tg000054l, h2tg000140l | + | + | h1tg000014l, h2tg000001l |
| HG01109 | - | + | h1tg000053l, h2tg000076l | - | - | h1tg000053l, h2tg000076l | + | + | h1tg000013l, h1tg000094l, h2tg000300l, h2tg000031l | + | + | h1tg000014l, h2tg000060l |
| HG01243 | - | + | h1tg000013l, h2tg000010l | - | - | h1tg000013l, h2tg000010l | + | + | h1tg000022l, h2tg000036l, h2tg000223l | + | + | h1tg000061l, h2tg000002l |
| HG02080 | ? | + | h2tg000037l | + | + | h1tg000109l, h2tg000041l | (+) | + | h1tg000004l, h2tg000090l | + | + | h1tg000061l, h2tg000072l |
| HG02109 | + | + | h1tg000131l, h2tg000203l | + | + | h1tg000100l, h2tg000104l | + | + | h1tg000282l, h2tg000194l | + | + | h1tg000051l, h2tg000080l |
| HG02145 | ? | + | h2tg000042l | - | ? | h1tg000216l, h1tg000024l, h2tg000042l | + | + | h1tg000121l, h2tg000046l, h2tg000348l | + | + | h1tg000120l, h2tg000127l |
| HG02723 | ? | - | h2tg000118l | + | ? | h1tg000078l, h2tg000118l | + | + | h1tg000091l, h1tg000323l, h2tg000255l | + | + | h1tg000238l, h2tg000053l |
| HG02818 | + | + | h1tg000101l, h2tg000010l | - | + | h1tg000101l, h2tg000010l | + | + | h1tg000064l, h1tg000075l, h2tg000050l, h2tg000367l | + | + | h1tg000197l, h2tg000370l |
| HG03098 | + | + | h1tg000072l, h2tg000003l | - | - | h1tg000018l, h2tg000003l | + | + | h1tg000077l, h2tg000057l | + | + | h1tg000027l, h2tg000055l |
| HG03486 | + | + | h1tg000040l, h2tg000101l | ? | + | h1tg000040l, h2tg000101l | (+) | + | h1tg000149l, h2tg000041l, h2tg000275l | + | + | h1tg000101l, h2tg000164l |
| HG03492 | - | - | h1tg000141l, h2tg000116l | + | + | h1tg000053l, h1tg000141l, h2tg000116l | + | + | h1tg000040l, h2tg000088l, h2tg000336l | + | + | h1tg000193l, h2tg000051l |
| NA18906 | ? | + | h1tg000016l, h2tg000016l | ? | - | h1tg000052l, h2tg000034l | + | + | h1tg000019l, h2tg000005l, h2tg000146l | + | + | h1tg000041l, h2tg000041l |
| NA19240 | + | ? | h1tg000047l | - | - | h1tg000047l, h2tg000010l | ? | + | h2tg000168l | + | + | h1tg000092l, h2tg000205l |
| NA20129 | + | + | h1tg000089l, h2tg000117l | ? | - | h1tg000081l, h2tg000007l | + | + | h1tg000189l, h2tg000003l | + | + | h1tg000075l, h2tg000272l |
| NA21309 | + | + | h1tg000188l, h2tg000104l | ? | ? | h1tg000188l, h2tg000241l | + | + | h1tg000106l, h2tg000272l | + | + | h1tg000332l, h2tg000194l |

**Figure S6. Full table of satellite in/del and inversion genotypes.** '+' indicates that the rearrangement was found on the maternal (M) or paternal (P) haplotype. '-' indicates that a spanning contig was found that showed no evidence for the rearrangement. '?' indicates that no spanning contig was found, perhaps due to assembly failure in the region (so it provides no evidence either way). '(+)' indicates that only one of the two insertion breakpoints was detected on any contig. The relevant contig names are listed for each individual.

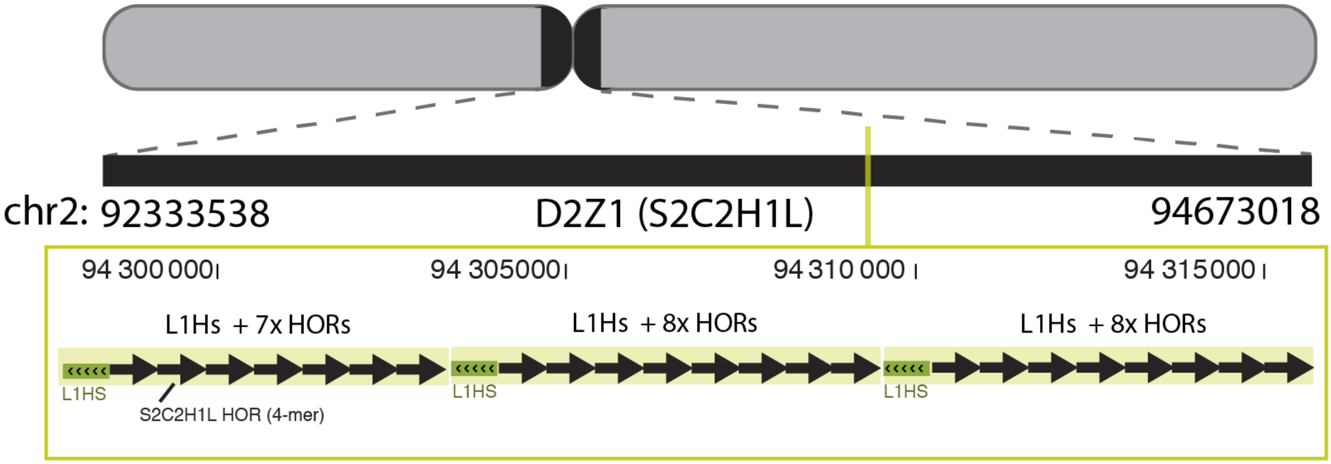

**Figure S7. Rare TE-involved repeat expansion within the active array on chromosome (D2Z1/S2C2H1L).** Transposable elements (TEs) are rarely embedded in homogenous arrays of alpha satellites, and when they are observed it is typically as a single event and rarely are these integrated TEs involved in local repeating units. Here we show a rare exception where we document a local repeat expansion involving a L1Hs/LINE element and 7 or 8 HORs (4-mers, S2C2H1L.1-4) on chromosome 2.

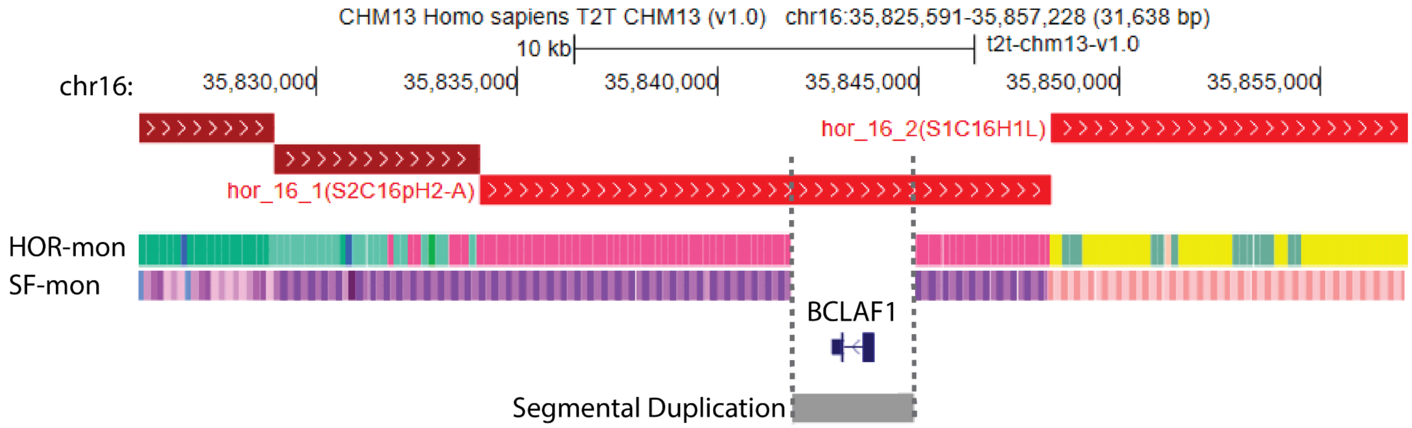

**Figure S8. Evidence of a gene embedded within a HOR array**. Study of the CHM13 centromeric region on chromosome 16 revealed a small 3kb segmental duplication (CHM13v1.0, chr16:35841889-35844926), with one other paralogous region marked on chr5:111458431-111461614 (< 98% identity, with 0.958 fracMatch reported). The gene, *BCLAF1P2-201* (BCL2 Associated Transcription Factor 1 P2, ENSG00000279800.2), is reported as a processed pseudogene with no frame documented for the exon and no evidence for expression by Salmon Transcript-level expression estimates or by Pro-Seq mapped data.

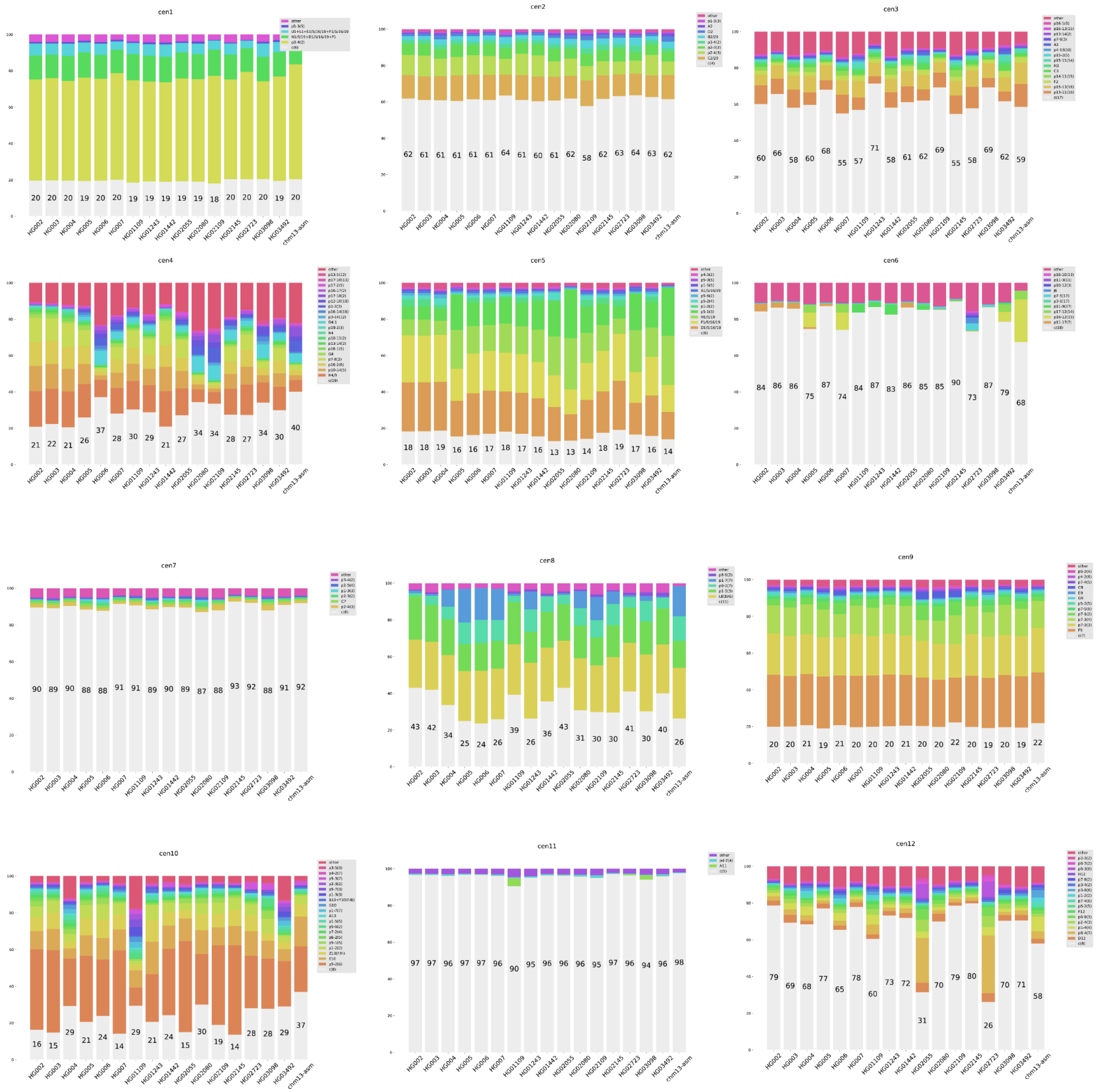
cen1
cen2
cen3
cen4
cen5
cen6
cen7
cen8
cen9
cen10
cen11
cen12

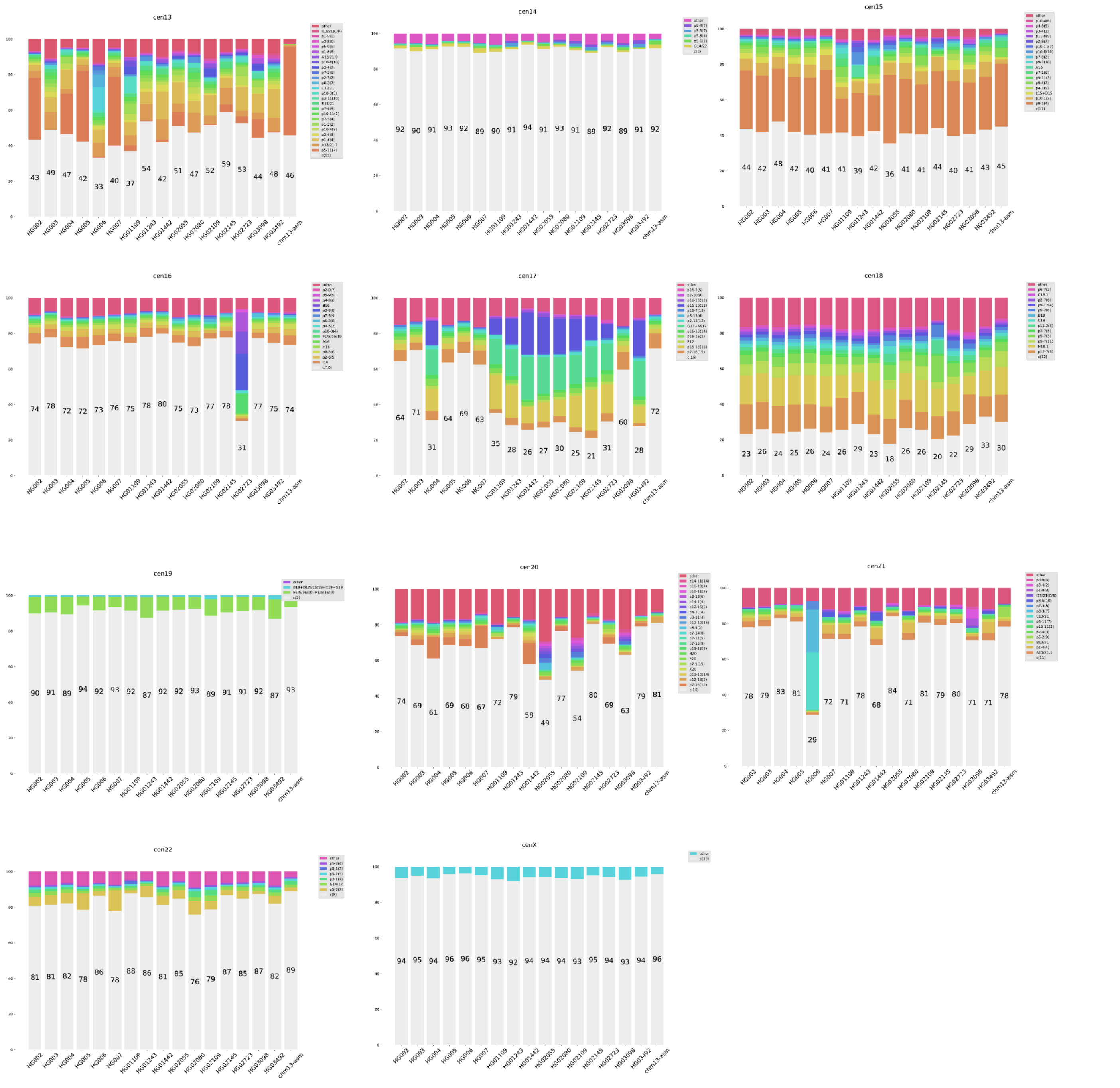

**Figure S9. Most Frequent HORs.** Each barplot corresponds to an individual centromere. Each bar shows the most frequent in a particular sample. Last bar corresponds to the assembly of CHM13. The (canonical) HOR in a particular centromere is marked as c. Each occurrence of a partial HOR that includes monomers from i to j is labeled as pi,j (*46*).

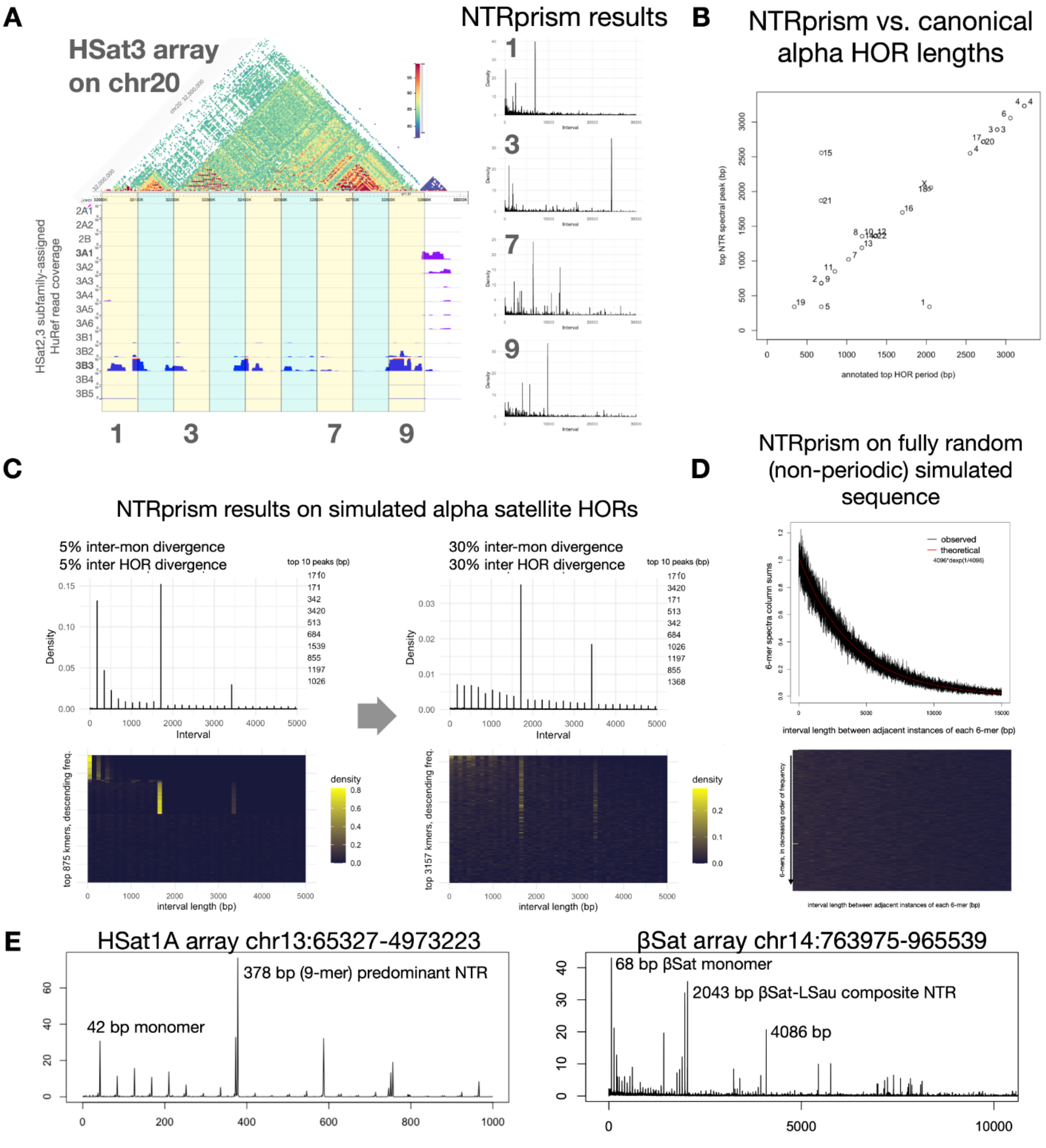

A
HSat3 array on chr20
NTRprism results
HSat2,3 subfamily-assigned HuRef read coverage
2A1
2A2
2B
3A1
3A2
3A3
3A4
3A5
3A6
3B1
3B2
3B3
3B4
3B5
1
3
7
9
B
NTRprism vs. canonical alpha HOR lengths
top NTR spectral peak (bp)
annotated top HOR period (bp)
C
NTRprism results on simulated alpha satellite HORs
5% inter-mon divergence
5% inter HOR divergence
top 10 peaks (bp)
1710
171
342
3420
513
684
1539
855
1197
1026
Density
Interval
30% inter-mon divergence
30% inter HOR divergence
top 10 peaks (bp)
1710
3420
171
513
342
684
1026
1197
855
1368
top 875 kmers, descending freq.
top 3157 kmers, descending freq.
interval length (bp)
density
D
NTRprism on fully random (non-periodic) simulated sequence
observed
theoretical
6-mer spectra column sums
interval length between adjacent instances of each 6-mer (bp)
E
HSat1A array chr13:65327-4973223
378 bp (9-mer) predominant NTR
42 bp monomer
βSat array chr14:763975-965539
68 bp βSat monomer
2043 bp βSat-LSau composite NTR
4086 bp

**Figure S10. HSat2,3 classification and NTRprism analysis. A**) For a block of HSat3 from chr20:32000000-33000000, a browser screenshot illustrating how this sequence was classified into subfamilies. HuRef read alignment tracks indicate the coverage of the region by sets of HuRef reads assigned within each HSat2,3 subfamily. The track with the highest coverage across the array becomes the array's subfamily assignment; in this case, it was assigned as HSat3B3 for the left block of the array, and HSat3A1 for the small array on the right-hand side. A dotplot showing the max % sequence identity for alignments of 5-kb blocks is shown above, illustrating clear separation of the 3B3 and 3A1 components of this larger super-array, as well as evidence of smaller regions of recent sequence expansion within the 3B3 array. To investigate how repeat nested tandem repeat periodicity might vary across the 3B3 array, we split the array into 9 subregions, each roughly 100-kb long, and applied NTRprism to these subregions. On the right-hand side are the resulting NTRprism spectra for a subset of the regions, showing wide variation in the predominant NTR length. **B**) For all live alpha HOR arrays in CHM13,a comparison of NTRprism's periodicity predictions and the annotated canonical HOR length for that array. Chromosomes off the diagonal have abundant non-canonical structural variant HORs that explain their discrepancy. **C**) The results of applying NTRprism to 1 Mb of simulated alpha-satellite-like tandem repeats, for which inter-monomer and inter-HOR divergence values were varied. Even with high sequence divergence, NTRprism uncovers the correct periodicity, but an increase in noise is clearly evident with higher divergence. **D**) The results of applying NTRprism to 1 Mb of fully randomized sequence with no periodicity, showing a close match to the theoretical exponential distribution of inter-k-mer lengths. **E**) left: the results of applying NTRprism to a large HSat1A array on chr13, showing that the canonical 42 bp repeat is organized into 378 bp NTRs (or in this case, HORs). Right: the results for a composite beta satellite array on chr14 showing both the canonical 68-bp beta monomer but also a larger 2043 LSau-beta composite NTR (and its 4086 bp dimer).

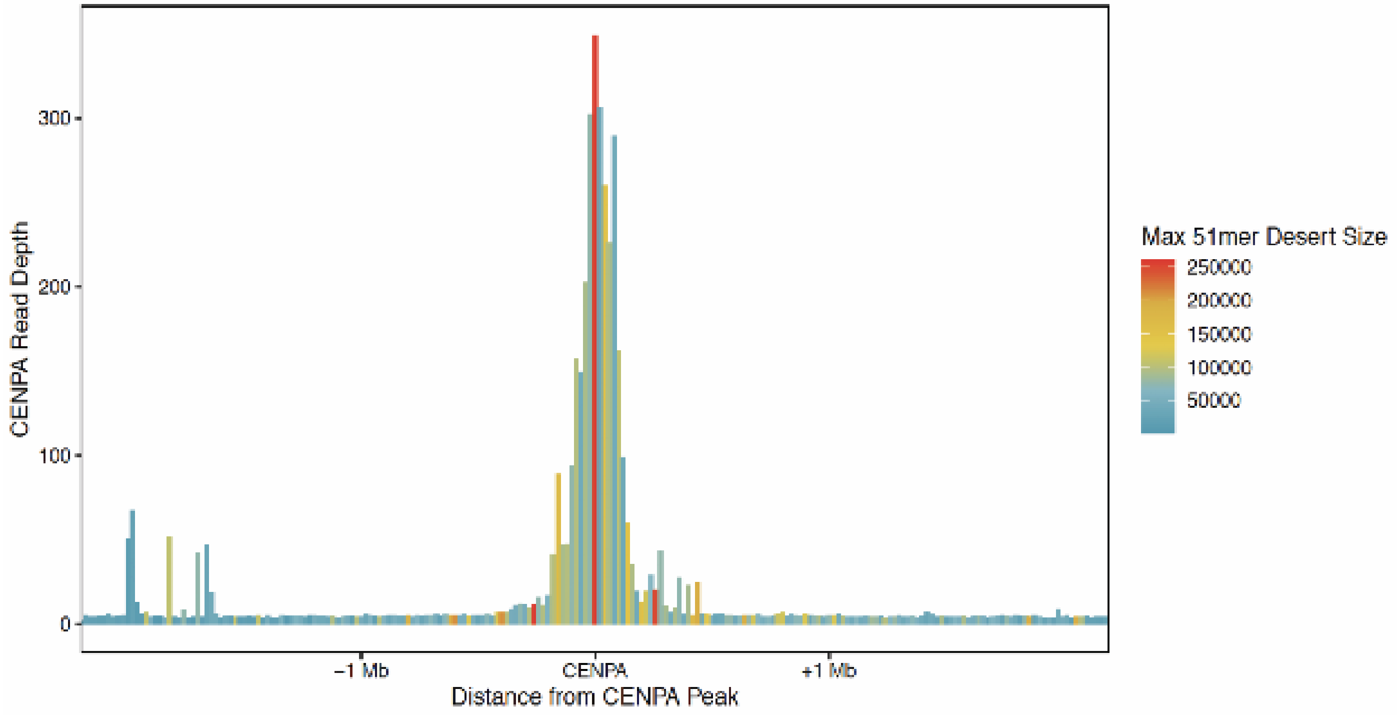

**Figure. S11**. **Unique markers are depleted at the sites of centromere protein enrichment**. Centering on the regions with the highest coverage of CENP-A reads (NChIP- CENP-A data), we identified an increase in the length of “marker desert regions”, or span of sequences that lack a unique 51 bp marker.

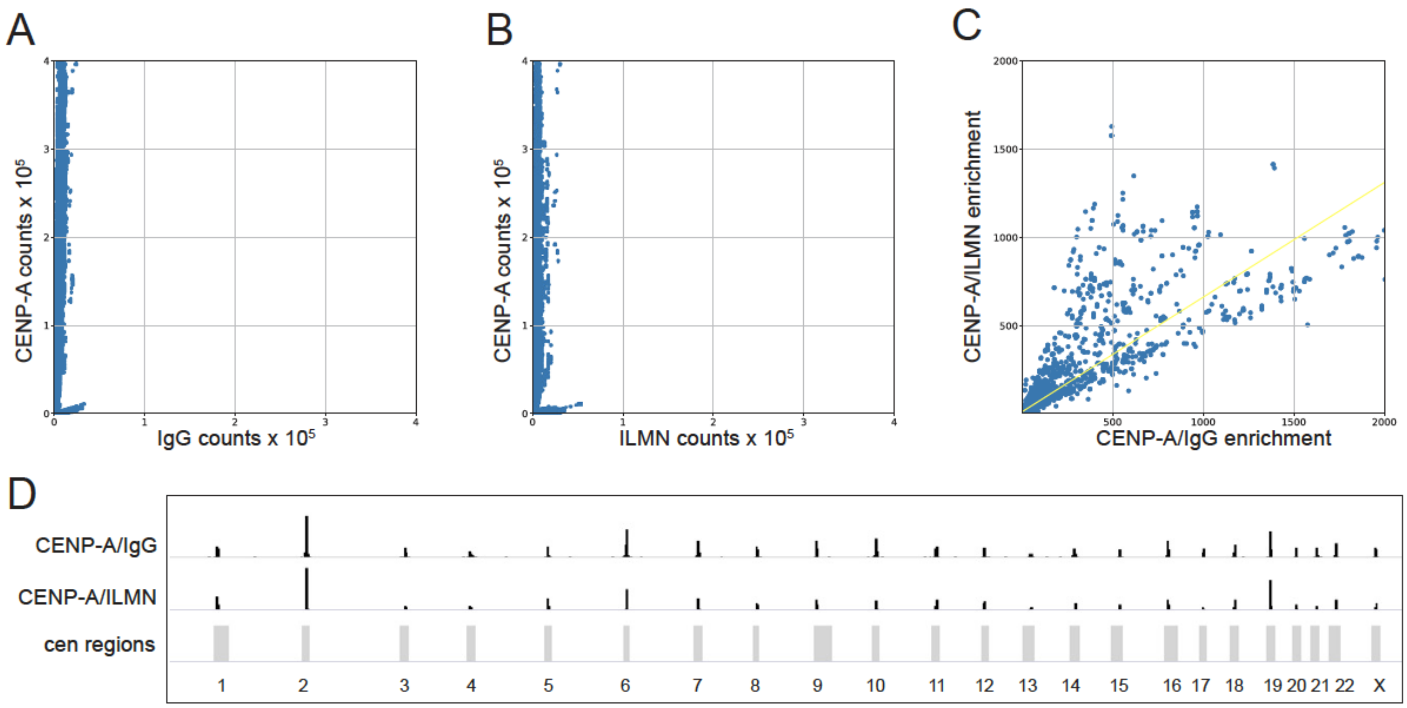

**Figure S12**: Calculating k-mer enrichment over IgG or PCR-free Illumina reads for the reference-free analysis yields similar results . A) Scatter plot of normalized k-mer counts of all 51 bp k-mers found in CENP-A and IgG libraries from read 1 (r1) of paired-end ChIP-seq experiment. B) Scatter plot of normalized k-mer counts of all 51 bpk-mers found in CENP-A from r1 of a paired-end ChIP seq experiment, compared to those from r1 of a PCR-free Illumina sequencing dataset. C) X-Y correlation plot of enrichment ratios of a randomly selected subset of unique 51 bp k-mers found to be enriched in CENP-A from the analyses in both A and B (yellow line is line of best fit, R2=0.866), a subset of 10,000 k-mers common to both datasets is shown. D) Bowtie1 alignment of CENP-A enriched 51 bp k-mers from A and B shows that both sets of enriched k-mers align to centromeric regions.

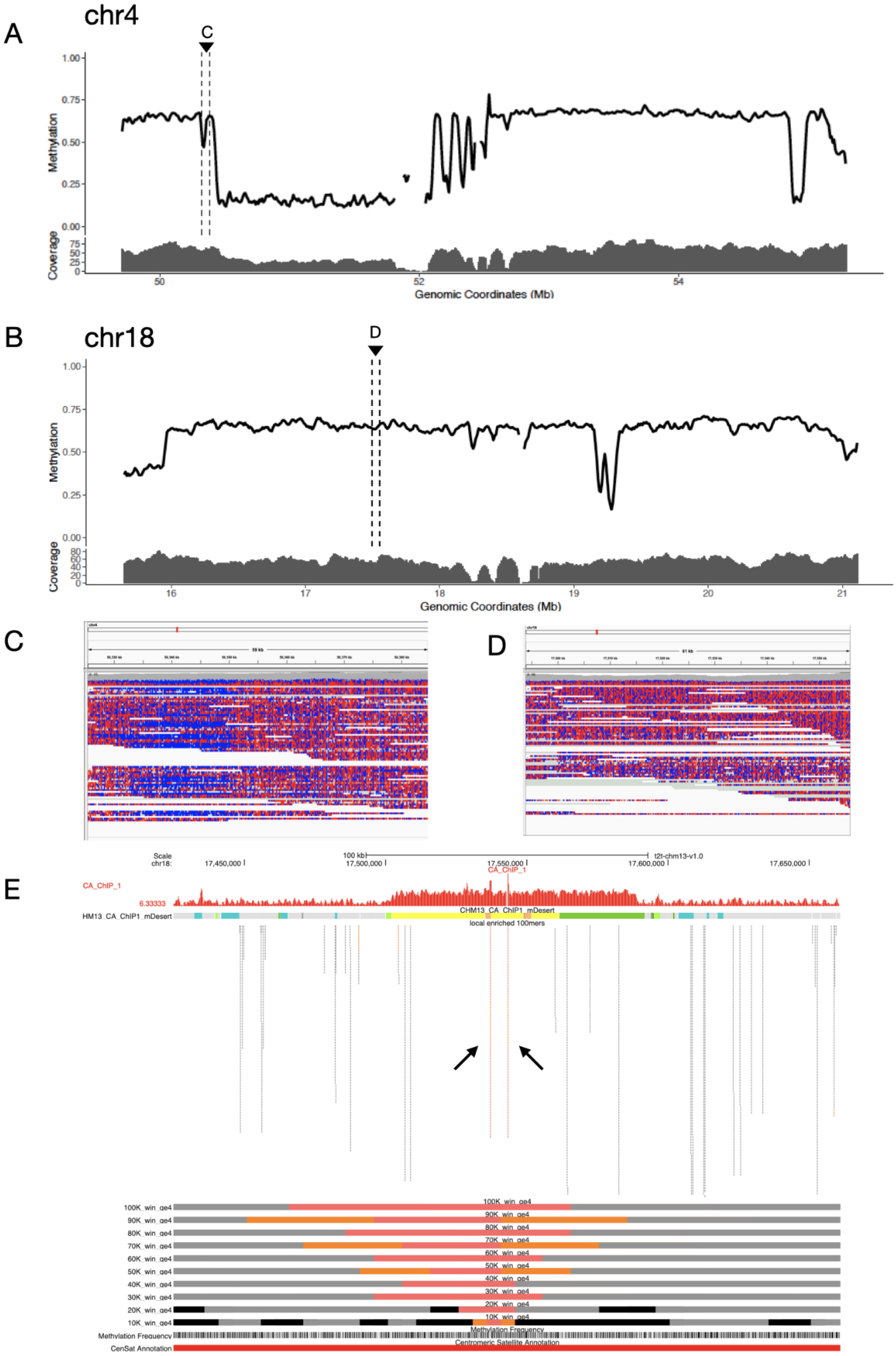

A
chr4
C
Methylation
1.00
0.75
0.50
0.25
0.00
Coverage
75
50
25
0
50
52
54
Genomic Coordinates (Mb)
B
chr18
D
Methylation
1.00
0.75
0.50
0.25
0.00
Coverage
80
60
40
20
0
16
17
18
19
20
21
Genomic Coordinates (Mb)
C
D
E
Scale
chr18:
100 kb
t2t-chm13-v1.0
17,450,000
17,500,000
17,550,000
17,600,000
17,650,000
CA_ChIP_1
6.33333
HM13_CA_ChIP1_mDesert
CHM13_CA_ChIP1_mDesert
local enriched 100mers
100K_win_ge4
90K_win_ge4
80K_win_ge4
70K_win_ge4
60K_win_ge4
50K_win_ge4
40K_win_ge4
30K_win_ge4
20K_win_ge4
10K_win_ge4
Methylation Frequency
CenSat Annotation
Centromeric Satellite Annotation

**Figure S13. Identifying small regions of enrichment outside of the primary CDR region** We detected smaller regions of centromere protein enrichment outside of the primary CDR, with some overlapping a minor, secondary CDR (chromosome 4) or no CDR at all (chromosome 18). **A**) Methylation profile (with read coverage) is provided for the centromeric region on chromosome 4. Dotted lines indicate the second site of CENP-A enrichment with a smaller CDR (expected to present a heterozygous site either between the two homologous chromosomes or high-frequency epigenetic variation within the population of cells). Increasing the resolution to focus on the region marked with “C”, to note data shown in panel C. **B**) Methylation profile and coverage plot across centromere array on chromosome 18. Site where enrichment is determined by region specific markers does not provide evidence for a drop in methylated CpG, as supported by zoomed in panel D (with region marked with dotted lines and noted with a “D”). C) Study of reads spanning the smaller CDR region on chromosome 4 provide evidence for a region without unmethylated bases (blue: unmethylated CpG, red: methylated CpG).In contrast, no apparent signature of unmethylated bases was observed on chromosome 18 (D). E) Illustrates the location of a set of enriched regional k-mers (logTransformed normalized ratio of 100-mers that are unique to chromosome 18 and are only observed within a 10kb window. Read alignment data is shown before marker assisted mapping.

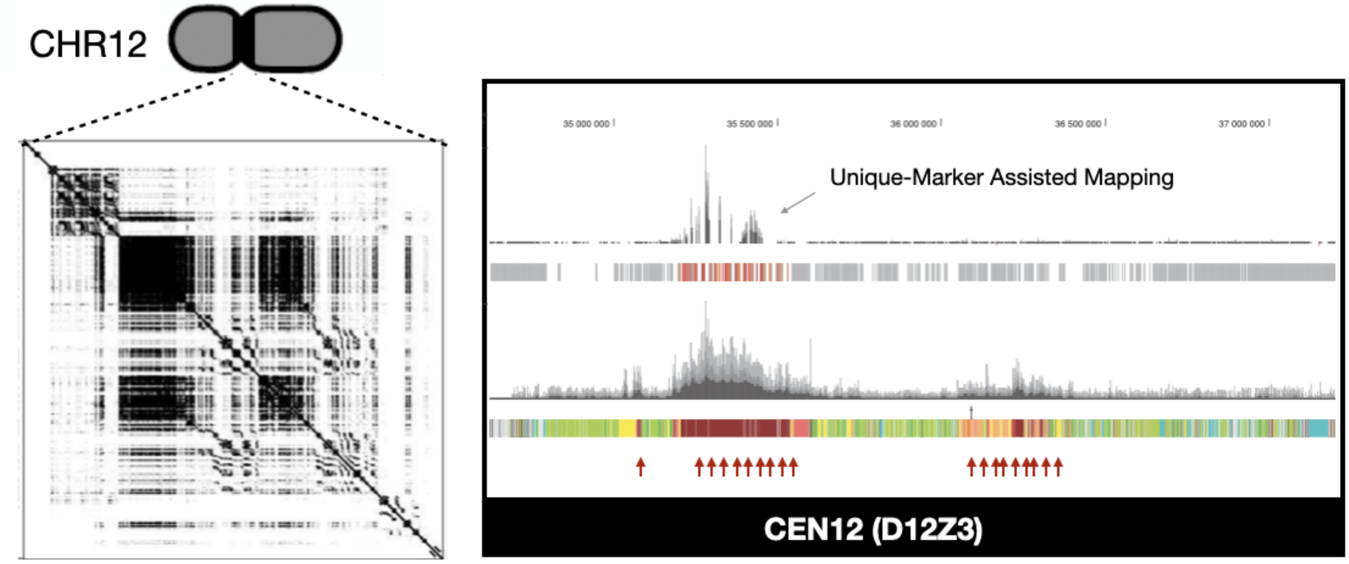

**Figure S14. Unique k-mer mapping strategy can determine CENP-A enrichment specifically to one of two large macro-repeat structures in centromere 12.** Study of the HOR array on chromosome 12 (D12Z3, or S1C12H1L) revealed large repeat structures (as illustrated in the dot plot on right). These repeats challenge short read mapping strategies which may suggest a functional dicentric chromosome, or two locations of CENP-A enrichment in the array (as indicated by the red arrows and heatmap, which shows the density of exact matched alignments - no mismatch, after excluding secondary and partial alignments - with multi-mapping allowed). Top of the left panel, the wiggle track is showing read coverage after filtering, and retaining only sequence alignments that fully span a unique marker (of length 100bp). These data are consistent with regional marker enrichment strategy (shown as increased enrichment (dark red/red/orange represent sequences greater than 4-fold, after taking the log-transformed normalized ratio of 100-mers (reference-free approach) and grey represent region specific 100-mers that do not provide evidence for enrichment.

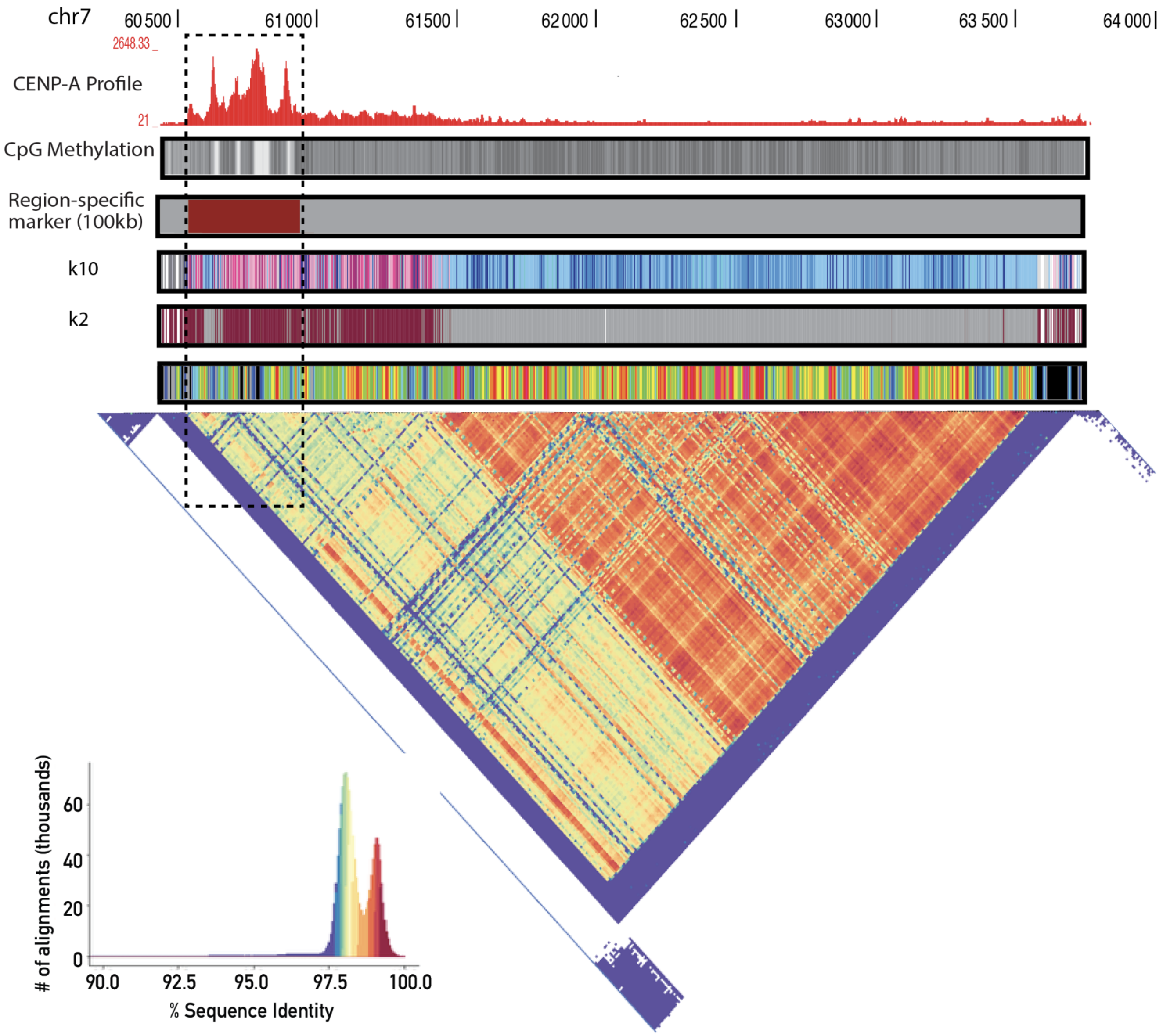

**Figure S15. Centromere 7 array organization.** Kinetochore assembly enrichment is observed over the youngest HOR-haplotype (k=10) in the D7Z1 array.

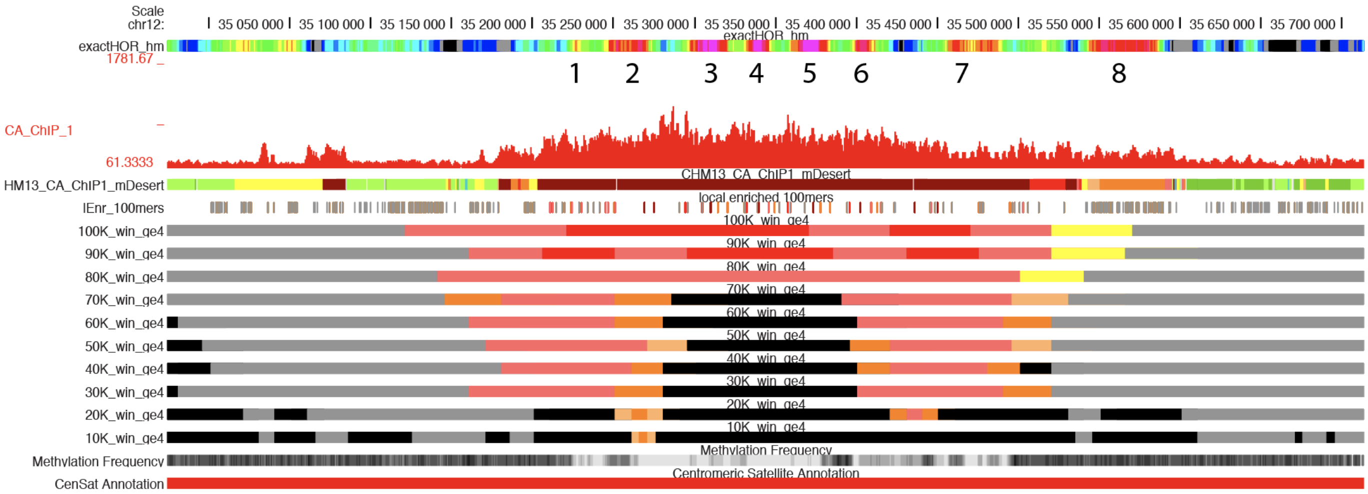

**Figure S16. CENP-A enrichment on chromosome 12 revealed a zone of recent HOR expansions.** Centromere enrichment region on chromosome 12 overlaps eight labeled sites of recent duplication as shown in the exact HOR matching heat map. Where a sliding window (representing 10x the canonical HOR length) reports the number of exact HOR matches. Color density for exact matching: 10/10, bright pink, 9/10, red, 8/10 orange, 7/10 yellow, 6/10 green, 5/10 teal, 4/10 light blue, 3/10 dark blue, 2/10 grey, 1/10 black (with the assumption of each HOR at least having a value of 1/10 for a self-alignment). CENP-A read mapping (unfiltered) is shown as the wiggle track (red). Exact multi-mapping track is shown below, with dark red/red/and orange showing regions that represent both true and false mapping. Using a sliding windows of length 10 kb to 100 kb, we identified region specific 100-mers and marked windows with dark red/red/orange/yellow if they overlap with at least two reference free regional kmers that are enriched >2 logTenriced ratio of NChIP data / matched background control. Windows that lack region-specific markers, and do not offer any information, are shown in black. Methylation profiles are provided as a density track, revealing that the CENP-A enrichment overlaps a CDR (as previously reported, Gershman et al).

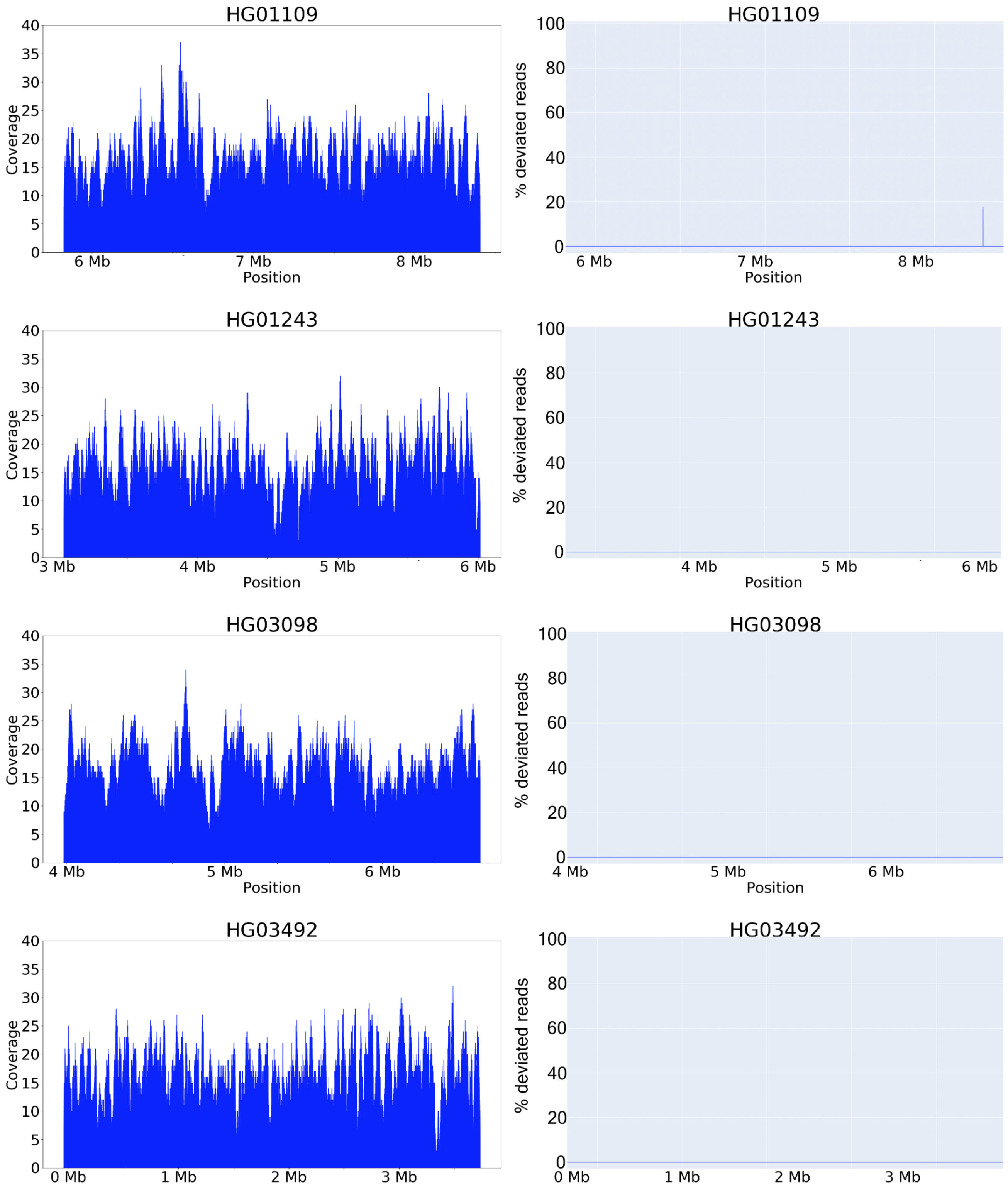

**Figure S17** Evaluating assembled centromere arrays from chrX using TandemTools. Left: coverage plots of 4 HiFi read assemblies. Right: Plots revealing discrepancies between the mapped reads and the assembly. The x-axis is chromosome position, the y-axis is percent of deviated reads.

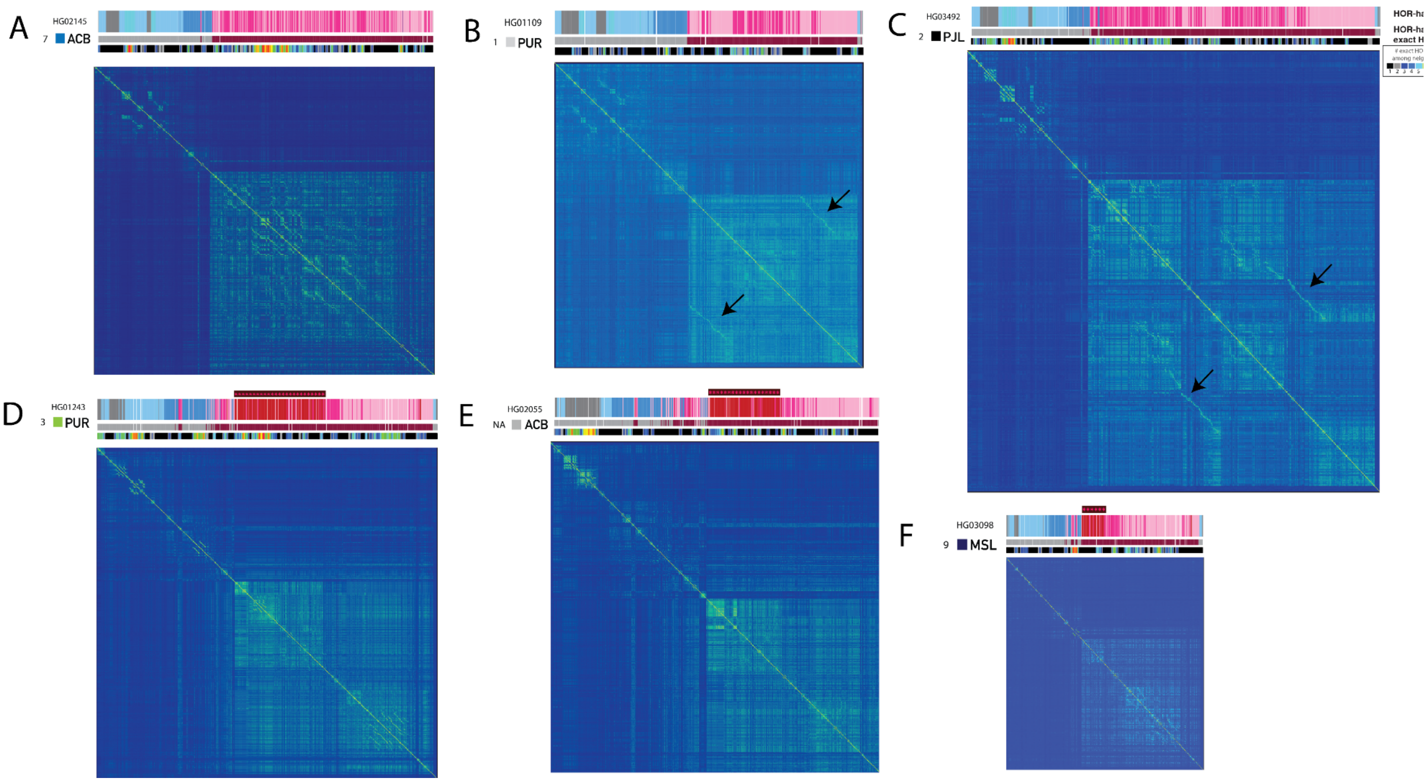

**Figure S18**. **Characterization of assembled centromere arrays from chrX across a panel of XY cell lines.** Self dot plots generated by an exhaustive search for exact matches between any two sequences (described Section 7) across six assembled X arrays reveal large blocks consistent across all lines that support hor haplotype classification (k=2, dark red versus grey). Also these plots identify small regions of recent local duplication (in support of the exact matching regions of duplication heat map track). Larger duplicated regions spanning hundreds of kilobases are noted (black arrow) only in **B** (HG01109 PUR, AMR) and **C** (HG03492, PJL, SAS) and absent in **A** (HG02145 ACB, AFR). Finally, dot plots also visually support the similarity blocks of the recent expansion of the dark red more ancient HOR-hap (k=7, arrays **D-F**).

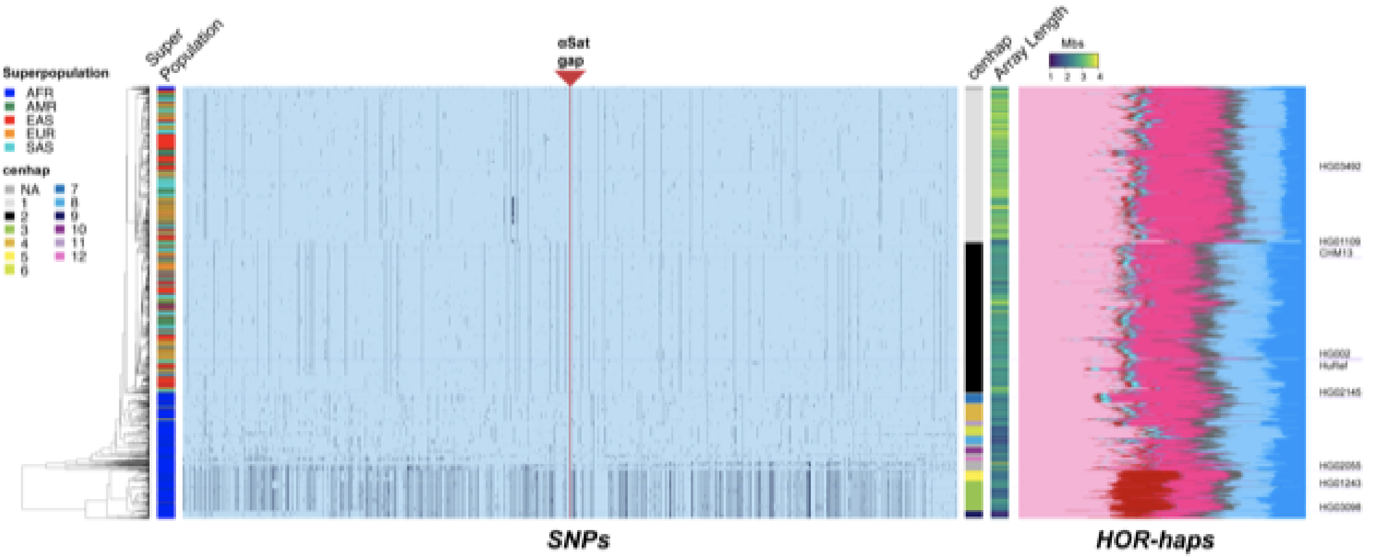

**Figure S19.** Cenhaps of human male chrX in the 1000 Genomes demonstrate robust genotyping and association with αSAT and HOR-hap composition. SNPs in the pericentromeric regions of the p and q arms of chrX were filtered for those occurring at least 10 times among the 3202 individuals (4805 X chromosomes) in the 1000 Genomes samples (Byrska-Bishop, 2021). Genotypes from 4000 SNPs flanking the αSATs gap (p:58261243-58539894, q:62506482-62754579, Hg38) for the 1599 1000 Genomes male genomes were merged with that of CHM13, HuRef and HG002, clustered on these using UMPGA and then 'cut' into 10 large subclades, cenhaps (Langley_2019). Note 98 genomes clustered in small clades that are labelled grey. The red line indicates position of the αSATs, a gap in the Hg38 closed in this report. The dendrogram reflecting the descent of each genome is on the left adjacent to a bar indicating the 'Superpopulations'. The SNP genotype of each genome (major allele in blue and minor allele in black) clearly fall into distinct haplotypes. The two most common cenhaps (light grey and black in the color bar to the right) are common outside of Africa, while much more cenhap diversity is found within Africa, including an anciently diverged clade of cenhaps 9, 10, 11 and 12 (Langley et al. 2019). To the right of the SNPs, a green-yellow bar displays estimated alpha array lengths and shows that cenhaps exhibit systematic differences in total αSAT (see Supplemental Section 7). The variation of HOR-hap composition (on the right) shows not only systematic differences among the 10 cenhaps, but also differences arising more recently (smaller clades). On the far right are the positions in the dendrogram CHM13 and other reference genomes from Figure 5. Note the lines marking their positions across the figure to the dendrogram.

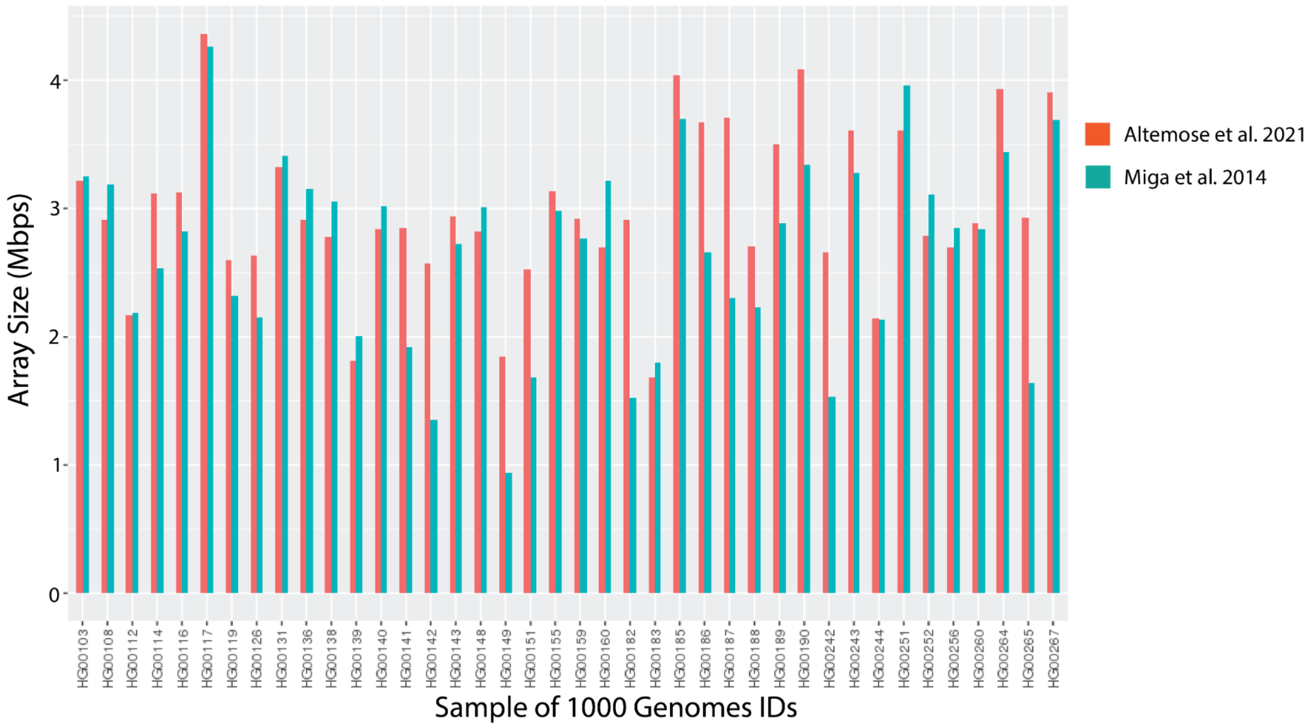

**Figure S20. αSat centromere X array size distributions compared to previous estimates.** Sampling of individuals from 1000 genome data (cite) to provide a comparison of X-array sizing to previous estimates (Miga et al 2014) that used low coverage (~2-4x) 1000 Genomes Data (cite) relative to 24-mers in the HuRef genome. Sizing estimates were relatively consistent, with notable exceptions (e.g. HG00149 and HG00242).

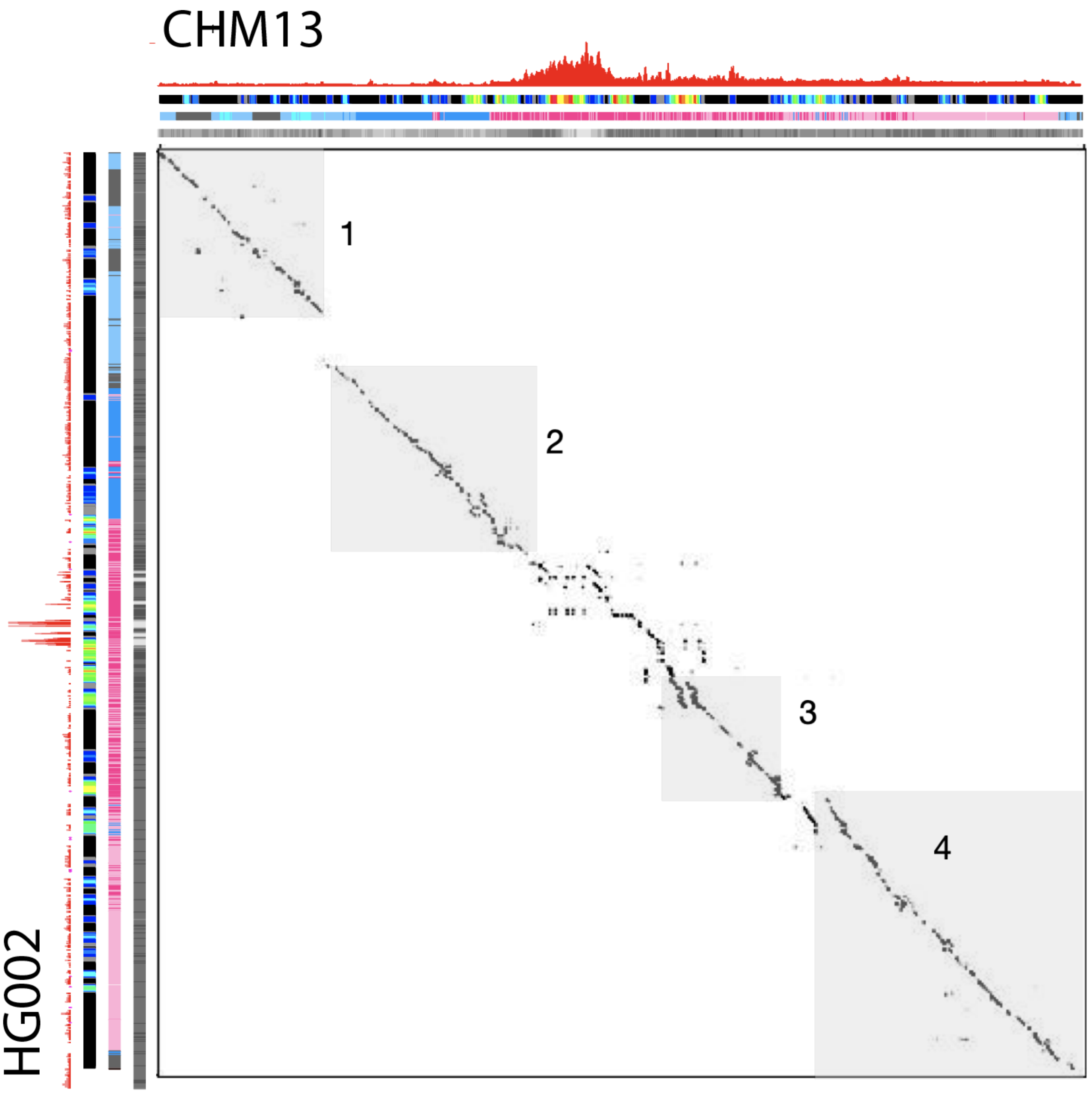

**Fig. S21. Dot plot (word 2000 bp) of HG002 and CHM13 X arrays** Kinetochore proteins are present in both arrays over CDRs and young HOR-haps and the HOR sequences enriched with CENP-A represent local duplication events that are not shared and distinguish the two arrays. The four regions highlighted in Figure 5 are shown as greyed boxes.

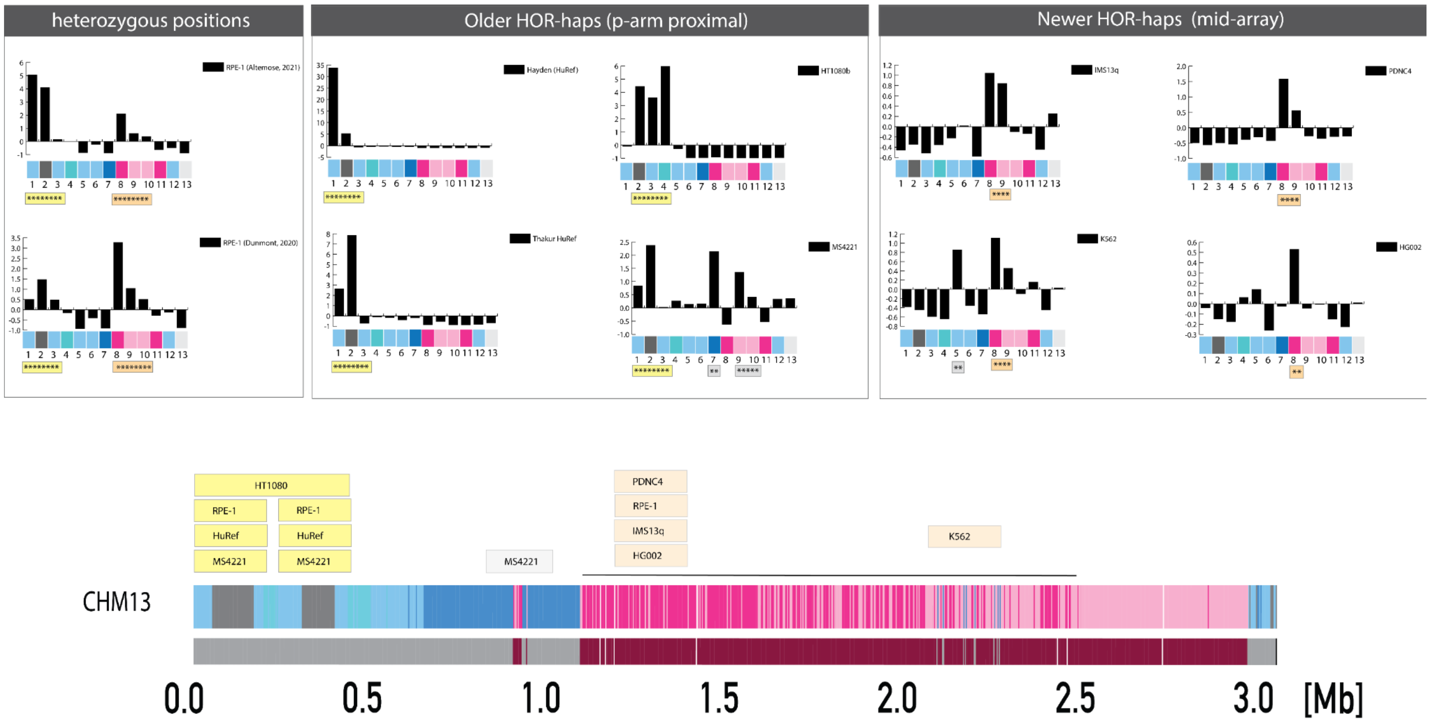

**Figure. S22**. **Using the T2T-CHM13 X array as a reference we studied the relative enrichment patterns of a panel of publicly available CENP-A (NChIP and CUT&RUN) data**. Left to right: we compared two independent CUT&RUN experiments from the RPE-1 cell line (XX) (91) and found consistent evidence for *heterozygous positions* of CENP-A within the same cell line, with enrichment on both older and newer HOR-haps. Next, in several individuals we observed CENP-A enrichment within the older HOR-hap subregion, proximal to the p-arm, indicating the presence of a centromere X epiallele (as shown for three XY individuals, HuRef, HT1080b (89) and MS4221 (90)). And finally, three additional 46,XX cell lines (IMS13q, PDNC4, K562 (92)) were determined to be consistent with CHM13, providing evidence that the same CENP-A+ HOR-hap is shared across both homologous X chromosomes in each line. In total, these findings are summarized by roughly placing the estimated site of enrichment relative to the position in the CHM13 X centromere.

### Supplemental Tables and Table Legends

**Table S1. CenSat Annotations**. [*Table_S1.cenSat_Annotation.tsv*] UCSC bed track describing centromere annotations in CHM13v1.0. columna headers provide 0) chromosome, 1) chromosome start, 2) chromosome end, 3) cenSat naming (where censat classification (e.g. hor) is followed by chromosome id (e.g. 1_10, would indicated chromosome 1 and the 10th time that cenSat category is observed moving p-arm to q-arm), 4) (dummy variable) score is set to 0 by default, 5) orientation is set to "." not specified.

**Table S2. An overview of published satellite DNA families in the human genome.**
Monomeric repeat length refers to the fundamental repeat unit of each family, although higher-order repeats may be much larger.

| Name | Published Chromosomal Localization(s) Classic + (hg38 RM) | Chromosomal Localization(s) in CHM13 | Published Monomeric Repeat Length (bp) | Published consensus sequence | Total size in CHM13 (bp) | NOTES | Ref(s) |
|---|---|---|---|---|---|---|---|
| **Alpha Satellite** | All | All | ~171 | | **85,183,432** | | Rosenberg et al.1978; Manuelidis et al. 1976; Musich et al. 1980 |
| **Human Satellite 3 (HSat3)** | major: 9, Y<br>minor: 1, 5, 10, 13, 14, 15, 17, 20, 21, 22 | >1Mb: 9,13,14,15,21,22<br>>100kb: 1,3,4,5,7,10, 17,20<br>>10kb: 16 | Variable, derived from ancestral (CATTC)n | well-conserved ATTCC pentameric repeats that are occasionally interspersed with the specific 10-bp sequence (ATGTCGGGTTG) | **47,680,005** | | Prosser et al. 1986<br>Cooke & Hindley 1979<br>Altemose et al. 2014 |
| **Human Satellite 2 (HSat2)** | major: 1, 2, 10, 16<br>minor: 7, 15, 17, 22 | >1Mb: 1,16<br>>100kb: 2,7,10,22<br>>10kb: 9,17 | Variable, derived from ancestral (CATTC)n | Enrichment of CATTCGATTG | **28,714,157** | | Prosser et al. 1986<br>Altemose et al. 2014 |
| **Human Satellite 1 (HSat1A) autosomal component** | 3, 4, 13, 14, 15, 21, 22; (8) | >1Mb: 3,4,13<br>>100kb: 14,15,21,22<br>>10kb: 8 | 42 (17+25) | 17-bp (ACATAAAATATCGAAAGT) and 25-bp (ACCCAAAATAGTAGTATTATATACTGT) joined subunits | **13,390,790** | Called SAR by RepeatMasker; Renamed here to HSat1A | Prosser et al. 1986 |

| | | | | | | | |
|---|---|---|---|---|---|---|---|
| **Human Satellite 1 (HSat1B) chrY component** | Y; (2,5,6,10,12,13,16,21,22) | >100kb: 15, 22<br>>10kb: 13,14,21 | 2.47 kb | incomplete | **1,150,150** | Composite of AT-rich and GC-rich components including an Alu fragment; Called HSATI by RepeatMasker; renamed here to HSat1B | Prosser et al. 1984 |
| **Beta Satellite** | 9, Y, 13, 14, 15, 21, 22; (1,2,3,4,5,6,7,10,16,19,20) | >1Mb:14,21,22<br>>100kb: 1,7,9,13,15,19,<br>>10kb: 3,5,8,10,16,20 | 68 | GATCAGTGCAGAGATATGTCACAATGCCCCTGTAGGCAGAGCCTAGACAAGAGTTACATCACCTGGGT | **7,688,273** | Often found in composite repeats with LSAU elements, travels in NTRs | Waye and Willard 1989b |
| **CER** | 13, 14, 15, 21, 22, Y; (2,3,4,5,9,18,20) | >100kb:13,14,21,22<br>>10kb:9,15,18 | 48 | CAGAACACTGCTGCTGGGTTCTGAGTGTTTGTCCCTCACATAGGATTC | **1,082,730** | | Metzdorf et al. 1988 |
| **Gamma Satellite** | 8, X; (1,2,3,4,5,6,9,12,13,14,15,17,18,21,22,Y) | >100kb:8,12<br>>10kb:9,13,14,15,21,22,X | 220 | GCTGGGAGCCTCCCAAGGAGGCCTCTCCCATCCCAGAAGCCCCCAGGGCTGTCCCGGGCGGGCTGTAAAGCCCCAGGCTTTGGAGCAGGGTGCCTGTGTCTCTCGCGGAAGGCCCCCACAAGCGAAAACGGGGCCGCAGGGTGGCGTGGGCGGGCCGCAGGGACTCAGGGGGACGTTGAGGCAGGCAGAGGGGAGAAGCGGCGAGACCGCAGGGAAT | **630,076** | | Lin et al. 1993 |
| **HSat4** | 1,16,19,X; (3,4,6,8,9) | >100kb:X<br>>10kb: 16,19 | 35 | TAGAATGCCTGGGGTCGCCCAGGTGTCTCTACCAT | **240,136** | | Warburton et al. 2008 |
| **HSat5** | (2,3,4,8,9,13,17,18,19,21,22,Y) | >10kb:9,4,19,21,22 | 14-15 | AGGGCCTCACTGACC | **164,966** | found in composites with other satellites | Smit, A.F. RepeatMasker update 2003 |
| **“p-censat” /other** | | | | | **4,062,171** | | |

**Table S3. HOR Annotation and Naming Nomenclature.** [*Table_S3.HOR_table.tsv*] Column 0) provides unique identifier, 1) CHM13v1.0 chromosome, 2) Assignment of Suprachromosomal Family (SF), 3) Old HOR name, 4) New HOR name, 5) Comment, 6) Basic HOR length (mon), 7) Expected order*, 8) Coords of master HOR in hg38 or contig, 9) RM or sample contig, 10) RM size (bp), 11)State 12)) CenpA reads, 13)Homogeneous/Divergent, 14)Age of the new-SF HORs, 15)Reference. *Note on monomer numbering: Mind that not all HORs in our annotation are supposed to have a straight numbering (1, 2, 3, 4, 5, ..n). Some HORs that belong to sister HOR families marked with letter indices (e.g. S1C10H1-B) are expected to have non-straight numbering (for rationale see Uralsky 2019). **Note on monomer order in divergent HORs: in some cases the HOR copies are so ruined that it is not possible to establish the consensus monomer order, only the number of different monomers in an array is clear. So, we can state the "Basic HOR length",but not the "Expected order".

**Table S4. SF-mon Summary Table,** Using previously described methods (Section 1), we performed complete, monomer-by-monomer classification of all αSat into 20 distinct suprachromosomal families. Each family was composed of SF-specific monomer classes.

| Layer | Monomer classes | Type | Monomer length (deletion in pos. 21) | Age group | Last common ancestor | Color in graphs and tracks |
|---|---|---|---|---|---|---|
| **SF1** | J1 & J2 | A+B | 171 | new | Gorilla | pink |
| **SF01** | J3 & J4 & J5 & J6 | A+B | 171 | new | Gorilla | cherry |
| **SF2** | D1 & D2 & FD | A+B | 171 | new | Gorilla | violet |
| **SF02** | D3 & D4 & D5 & D6 & D7 & D8 & D9 | A+B | 171 | new | Gorilla | dark violet |
| **SF3** | W1 & W2 & W3 & W4 & W5 | A+B | 171 | new | Gorilla | azur |
| **SF03** | | | | | | |
| **SF5** | R1 & R2 (irregular) | A+B | 171 | old | Orangutan | blue |
| **SF05** | R2 | A | 171 | old | Orangutan | light blue |
| **SF4** | Ga | A | 171 | old | Gibbon | yellow |
| **SF6** | Ha | A | 172+171 | old | † Proconsul? | brown |
| **SF7** | Ka | A | 172 | ancient | OWM | bright green |
| **SF8** | Oa & Na (dimer) | A | 172 | ancient | | olive & green |
| **SF9** | Ca | A | 172 | ancient | | red |
| **SF10** | Ba | A | 172 | ancient | | orange |
| **SF11** | Ja | A | 172 | ancient | NWM | lilac |
| **SF12** | Aa | A | 172 | most ancient | | grey |
| **SF13** | Ia | A | 172 | most ancient | | pomegranate |
| **SF14** | La | A | 172 | most ancient | | purple |
| **SF15** | Fa | A | 172 | most ancient | | teal |
| **SF16** | Ea | A | 172 | most ancient | | watery blue |
| **SF17** | Qa | | 172 | prehistoric | | |
| **SF18** | Pa & Ta | | 174+172 | prehistoric | | |

(dimer)

**Table S5. Alpha Satellite outside of cenSat annotation regions.** *[Table_**S5.AS_in_arms**.tsv]* This table provides the locations of small pieces of the most ancient αSat and occur far from the centromere, presumably at the sites of long-defunct ancient centromeres. Column information: 1) chromosome ID (CHM13v1.0), 2) chromosome start, 3) chromosome end, 4) HOR or monomer ID name, 5) strand information, 6) RM data, 7) Length (bp), 8) comment

**Table S6.Alpha satellite Inversions hor/mon main table [***Table_S6.Inversions_HOR_dHOR_mon.tsv***]** This table describes the alpha satellite inversions documented in CHM13. Column information: 1) chromosome ID (CHM13v1.0), 2) chromosome start, 3) chromosome end, 4) Mon switching information, 5) Strand switch, 6) Gap (bp), 7) cenSat_Annotation reference, 8) comment (manual curation notes)

**Table S7.** Annotating strand orientations across entire satellite arrays revealed several novel and unexpected anomalies.

| family | size (bp) | inversions | inv./Mb |
|---:|---:|---:|---:|
| **alpha HOR** | **69,986,938** | **7** | **0.10** |
| **alpha div/mon** | **15,196,494** | **256** | **16.85** |
| **HSat3** | **47,648,914** | **259** | **5.44** |
| **HSat2** | **28,705,695** | **12** | **0.42** |
| **HSat1** | **14,504,723** | **8** | **0.55** |
| **beta (*no lsau)** | **6,254,315** | **45** | **7.20** |

**Table S8.** *[Table_S8.cen6_primate_dating.xls]* Evolutionary relationship of a q-arm duplication and decayed remnants of the old centromere with dating information across non-human primates.

**Table S9. Gene counts between satellite arrays.** All gene annotations (*8*) occurring between any satellite annotations larger than 100 kb, broken down by class. “Expressed” indicates non-zero transcripts-per-million values in CHM13 reported by Iso-seq, RNA-seq, or PRO-seq (Hoyt et al. 2021). “Novel” refers to classification of the annotation as novel by (*8*)

| Class | Total | Expressed | Novel |
|---|---|---|---|
| IG_V_gene | 8 | 0 | 2 |

| | | | |
|---|---|---|---|
| IG_V_pseudogene | 16 | 0 | 9 |
| lncRNA | 234 | 89 | 145 |
| miRNA | 29 | 4 | 23 |
| misc_RNA | 13 | 1 | 6 |
| polymorphic_pseudogene | 2 | 0 | 0 |
| processed_pseudogene | 382 | 74 | 244 |
| protein_coding | 21 | 7 | 6 |
| rRNA | 8 | 5 | 0 |
| rRNA_pseudogene | 22 | 1 | 1 |
| snoRNA | 3 | 0 | 2 |
| snRNA | 9 | 0 | 5 |
| sRNA | 1 | 0 | 0 |
| StringTie | 13 | 13 | 6 |
| TEC | 132 | 34 | 119 |
| transcribed_processed_pseudogene | 23 | 9 | 15 |
| transcribed_unprocessed_pseudogene | 39 | 29 | 10 |
| unprocessed_pseudogene | 211 | 63 | 80 |

**Table S10. Salmon and Pro-Seq data for genes genome wide. [***Table_S10.CHM13.CATv4.allTPM.tsv***]** Comma delimited table listing Transcript abundance was estimated in transcripts per million (TPM) with Salmon v1.3.0(16), using the CHM13 transcriptome (CATv4 and liftOff, (8)) and the v1.0 assembly as a "decoy" sequence to account for reads mapping to unannotated sequences.

**Table 11. StV10 Table.** *[Table_S11.StV10.tsv]* We decomposed each αSat HOR array first into individual monomers and then into entire HORs, revealing the positions of full-size canonical HORs and structural variant HORs resulting from insertions or deletions. This table breaks down the naming (column 0) of each HOR structural variant for each HOR array in CHM13, reports the number of these occurrences per array (column 1), and notes hybrid monomer structures observed (column 2).

**Table S12. Symmetry dHOR table. [***Table_S12.symmetry_dHORs.tsv***]** We report a symmetrical flanking arrangement divergent αSat: (dHORs). Column headers are as follows: 1) chromosome ID (CHM13v1.0), 2) chromosome start, 3) chromosome end, 4) HOR naming, 5) Length (bp), 6) chromosome arm (ie p-arm or q-arm), and 7) Comment (manual curation)

**Table S13**. **Symmetry monomeric table**. We report a symmetrical flanking arrangement of monomeric αSats which represent ancient, decayed centromeres of primate ancestors. These data are in agreement with data presented in Figure 3.

| AS Layer cen only | Aa | Ja | Ba | Ca | Oa Na | Ka | Ha | Ga | R2 | R1 R2 | dHOR | Pseudo | LIVE | Pseudo | dHOR | R1 R2 | R2 | Ga | Ha | Ka | Oa Na | Ca | Ba | Ja | Aa |
|---|---|---|---|---|---|---|---|---|---|---|---|---|---|---|---|---|---|---|---|---|---|---|---|---|---|
| 1 | | | | | | | | | | | | | ■ | ■ | ■ | | | | | | | | | | |
| 2 | | | | | | | | | | | | | ■ | ■ | | | | | | | | | | | |
| 3 | | | | | | | ■ | ■ | | ■ | ■ | ■ | ■ | | ■ | ■ | ■ | ■ | ■ | ■ | | | | | |
| 4 | | ■ | ■ | ■ | | | | | | | | | ■ | | | ■ | | | | | | ■ | ■ | ■ | |
| 5 | | | | | ■ | ■ | ■ | ■ | | ■ | | | ■ | | | ■ | | ■ | ■ | ■ | | | | | |
| 6 | | | | | ■ | | | | | | ■ | | ■ | | ■ | | | | | | | | | | |
| 7 | | | | ■ | ■ | ■ | ■ | ■ | ■ | ■ | | ■ | ■ | | | ■ | ■ | ■ | ■ | ■ | ■ | | | | |
| 8 | | | | ■ | | | | | ■ | ■ | ■ | | ■ | | | ■ | ■ | ■ | | | | ■ | ■ | ■ | |
| 9 | | | | | | | | | | | | | ■ | | | | | | | | | | | | |
| 10 | | | | ■ | ■ | ■ | ■ | ■ | ■ | ■ | ■ | | ■ | ■ | ■ | | | ■ | ■ | | ■ | ■ | | | |
| 11 | | | | ■ | ■ | | ■ | ■ | ■ | ■ | ■ | ■ | ■ | ■ | ■ | ■ | ■ | ■ | ■ | | ■ | ■ | | | |
| 12 | | | | | ■ | ■ | | | | | | | ■ | | | | ■ | ■ | ■ | ■ | ■ | ■ | ■ | ■ | |
| 13 | | | | | | | | | | | | | ■ | | | ■ | | ■ | | | | | | | |
| 14 | | | | | | | | | | | | | ■ | | | ■ | | ■ | | | | | | | |
| 15 | | | | | | | ■ | ■ | | | | | ■ | | | ■ | | ■ | | | | | | | |
| 16 | | | | ■ | ■ | | ■ | ■ | | | | | ■ | | | ■ | ■ | ■ | ■ | | | | | ■ | |
| 17 | | ■ | | | | | ■ | | | | | | ■ | | | ■ | | ■ | | | | | | | |
| 18 | | | | | | | | ■ | | ■ | | | ■ | | | | | | | | | | | | |
| 19 | | | | ■ | ■ | ■ | | ■ | | | | | ■ | | | ■ | | ■ | ■ | ■ | | ■ | | | |
| 20 | | | | | | | | | | ■ | | | ■ | | | ■ | | ■ | ■ | | | | | | |
| 21 | | | | | | | ■ | ■ | ■ | ■ | | | ■ | | | ■ | | ■ | | | | | | | |
| 22 | | | | | | | ■ | ■ | | ■ | | | ■ | | | ■ | | ■ | | | | | | | |
| X | ■ | | ■ | ■ | ■ | ■ | ■ | ■ | | ■ | | | ■ | | | ■ | | ■ | ■ | ■ | | ■ | ■ | ■ | |
| | SF12 | SF11 | SF10 | SF9 | SF8 | SF7 | SF6 | SF4 | SF5 | SF5 | SF3 | SF2 | SF1 | SF2 | SF3 | SF5 | SF5 | SF4 | SF6 | SF7 | SF8 | SF9 | SF10 | SF11 | SF12 |

*Note:* Small AS-containing SDs were ignored.

**Table S14. Size and Inter-array Divergence of Alpha Satellite Compartments (CHM13v1.0)**

| Layer | Monomer classes | Size per genome (Mb) | Mean divergence (%) | Comment |
|---|---|---|---|---|
| **HOR active** | SF1,2,3 | 61.600 | 1.7 | |
| **HOR inactive** | SF1-6 | 8.413 | 2.1 | |
| **HOR active+inactive** | | 70.013 | 2 | |
| **dHOR** | SF1-3,5 | 1.851 | 12.3 | |
| **mon** | SF4+ | 13.317 | NA | SF5-18 |
| **SF5** | R1 & R2 (irregular) | 1.442 | 15.8 | Divergence of R1 - 14.9%, R2 - 16.7% |
| **SF4** | Ga | 3.633 | 16.9 | |
| **SF6** | Ha | 1.560 | 19.7 | |
| **SF7** | Ka | 0.857 | 22.9 | |
| **SF8** | Oa & Na (dimer) | 1.735 | 21.8 | Divergence of Oa - 22.3%, Na - 21.2% |
| **SF9** | Ca | 0.760 | 24.9 | |
| **SF10** | Ba | 0.250 | 25.4 | |
| **SF11** | Ja | 0.300 | 25.7 | |
| **SF12** | Aa | 0.128 | 25.8 | |
| **SF13** | Ia | 0.016 | 27.3 | |
| **SF14** | La | 0.000 | NA | Absent in humans |
| **SF15** | Fa | 0.007 | 25.4 | |
| **SF16** | Ea | 0.020 | 28.4 | |
| **SF17** | Qa | 0.003 | 26.4 | |
| **SF18** | Pa & Ta (dimer) | 0.012 | 23.3 | Divergence of Pa - 20.7%, Ta - 25.9% |

**Table S15. interspersed_repeats_HOR, dHOR, and mon (combined) in centromeric, or cenSat annotated regions [*Table_S15.interspersed_repeats_HOR_dHOR_mon.tsv*]** We document embedded transposable elements across alpha satellite HORs, dHOR, and monomeric DNAs. Column header: (1) chromosome ID (CHM13v1.0), (2) chromosome start, (3) chromosome end, (4) repName Strand, (5) repClass, (6) repFamily, (7) Length, (8) HORregion, (9) Comment (manual curation)

**Table S16.** As shown in Fig. 3D, we have documented events of “inter-array seeding” and expansions of different satellite classes genome wide.

| Element | Previous element | Next element | Number of cases | Length (bp) | Length (Mpb) | Frequency of elements in regions (per Mb) |
|---|---|---|---|---|---|---|
| hor | censat | dhor | 1 | 70,011,897 | 70.0 | 0.014283 |
| hor | dhor | hsat1 | 1 | 70,011,897 | 70.0 | 0.014283 |
| hor | hor | bsat | 1 | 70,011,897 | 70.0 | 0.014283 |
| hor | hsat1 | hor | 3 | 70,011,897 | 70.0 | 0.042850 |
| hor | hsat1 | hsat1 | 3 | 70,011,897 | 70.0 | 0.042850 |
| **Total HOR** | | | **9** | 70,011,897 | **70.012** | **0.128550** |
| dhor | censat | mon | 1 | 1,851,153 | 1.9 | 0.540204 |
| dhor | dhor | censat | 2 | 1,851,153 | 1.9 | 1.080408 |
| dhor | dhor | hsat2 | 2 | 1,851,153 | 1.9 | 1.080408 |
| dhor | hsat2 | hor | 2 | 1,851,153 | 1.9 | 1.080408 |
| dhor | hsat2 | hsat2 | 2 | 1,851,153 | 1.9 | 1.080408 |
| dhor | hsat3 | ct | 1 | 1,851,153 | 1.9 | 0.540204 |
| **Total dHOR** | | | **10** | 1,851,153 | **1.851** | **5.402039** |
| mon | bsat | ct | 1 | 13,317,305 | 13.3 | 0.075090 |
| mon | bsat | hor | 1 | 13,317,305 | 13.3 | 0.075090 |
| mon | censat | censat | 5 | 13,317,305 | 13.3 | 0.375451 |
| mon | censat | ct | 8 | 13,317,305 | 13.3 | 0.600722 |
| mon | censat | dhor | 3 | 13,317,305 | 13.3 | 0.225271 |
| mon | ct | bsat | 1 | 13,317,305 | 13.3 | 0.075090 |
| mon | ct | censat | 7 | 13,317,305 | 13.3 | 0.525632 |
| mon | ct | gsat | 2 | 13,317,305 | 13.3 | 0.150181 |
| mon | ct | hsat1 | 1 | 13,317,305 | 13.3 | 0.075090 |
| mon | ct | hsat2 | 1 | 13,317,305 | 13.3 | 0.075090 |
| mon | ct | hsat3 | 11 | 13,317,305 | 13.3 | 0.825993 |
| mon | ct | hsat4 | 2 | 13,317,305 | 13.3 | 0.150181 |
| mon | dhor | censat | 6 | 13,317,305 | 13.3 | 0.450542 |
| mon | dhor | gsat | 1 | 13,317,305 | 13.3 | 0.075090 |
| mon | gsat | ct | 2 | 13,317,305 | 13.3 | 0.150181 |
| mon | gsat | dhor | 3 | 13,317,305 | 13.3 | 0.225271 |
| mon | gsat | gsat | 7 | 13,317,305 | 13.3 | 0.525632 |
| mon | hor | censat | 1 | 13,317,305 | 13.3 | 0.075090 |

| | | | | | | |
|---|---|---|---|---|---|---|
| mon | hor | hsat3 | 9 | 13,317,305 | 13.3 | 0.675812 |
| mon | hor | hsat4 | 1 | 13,317,305 | 13.3 | 0.075090 |
| mon | hsat1 | hsat3 | 1 | 13,317,305 | 13.3 | 0.075090 |
| mon | hsat2 | hsat2 | 1 | 13,317,305 | 13.3 | 0.075090 |
| mon | hsat3 | ct | 12 | 13,317,305 | 13.3 | 0.901083 |
| mon | hsat3 | hor | 7 | 13,317,305 | 13.3 | 0.525632 |
| mon | hsat3 | hsat3 | 20 | 13,317,305 | 13.3 | 1.501805 |
| mon | hsat4 | ct | 1 | 13,317,305 | 13.3 | 0.075090 |
| mon | hsat4 | hsat4 | 7 | 13,317,305 | 13.3 | 0.525632 |
| **Total mon** | | | **122** | 13,317,305 | **13.317** | **9.161013** |

**Table S17**. [*Table_S17.CENPA_estimates.tsv*] **CENP-A enrichment boundaries in CHM13v1.0.** Table listing enrichment sites as informed by boundaries of regional specific markers. (columns 0-3), estimated position of centromere dip region (CDR) in methylation profile (columns 4-6). Notation if centromere is expected to be positioned over the young HOR-hap (column 7). Estimated span of CENP-A region (column 8), Estimated span of CDR region (column 8)(noting that these regions often contain collections of one or more CDRs and provide a general estimate). Difference in span (column 9)

**Table S18. [***Table_S18.T2T_KGP4_MAC10_chrX_n0_FINAL_12_haps_props.txt*.] **CenHap Supplement table** 1599 males genotyped using published short-read sequencing data (86) into 12 cenhaps (with 98 individuals remaining unclassified).

**Table S19**. **X-array sizing**. [*Table_S19.all_male_DXZ1_lengths.csv*] Array sizing of 1599 males using kmer based approach.

### References

1. X. She, J. E. Horvath, Z. Jiang, G. Liu, T. S. Furey, L. Christ, R. Clark, T. Graves, C. L. Gulden, C. Alkan, J. A. Bailey, C. Sahinalp, M. Rocchi, D. Haussler, R. K. Wilson, W. Miller, S. Schwartz, E. E. Eichler, The structure and evolution of centromeric transition regions within the human genome. *Nature*. **430**, 857–864 (2004).

2. M. R. Vollger, X. Guitart, P. C. Dishuck, L. Mercuri, W. T. Harvey, A. Gershman, M. Diekhans, A. Sulovari, K. M. Munson, A. M. Lewis, K. Hoekzema, D. Porubsky, R. Li, S. Nurk, S. Koren, K. H. Miga, A. M. Phillippy, W. Timp, M. Ventura, E. E. Eichler, Segmental duplications and their variation in a complete human genome, , doi:10.1101/2021.05.26.445678.

3. N. Altemose, K. H. Miga, M. Maggioni, H. F. Willard, Genomic characterization of large heterochromatic gaps in the human genome assembly. *PLoS Comput. Biol.* **10**, e1003628 (2014).

4. A. F. A. Smit, R. Hubley, P. Green, RepeatMasker (1996).

5. S. Hoyt, From telomere to telomere: the transcriptional and epigenetic state of human repeat elements. *bioRxiv (in review)* (2021).

6. D. Olson, T. Wheeler, ULTRA: A Model Based Tool to Detect Tandem Repeats. *ACM BCB*. **2018**, 37–46 (2018).

7. A. M. Mc Cartney, Chasing perfection: validation and polishing strategies for telomere-to-telomere genome assemblies. *in prep* (2021).

8. S. Nurk, S. Koren, A. Rhie, M. Rautiainen, A. V. Bzikadze, A. Mikheenko, M. R. Vollger, N. Altemose, L. Uralsky, A. Gershman, S. Aganezov, S. J. Hoyt, M. Diekhans, G. A. Logsdon, M. Alonge, S. E. Antonarakis, M. Borchers, G. G. Bouffard, S. Y. Brooks, G. V. Caldas, H. Cheng, C.-S. Chin, W. Chow, L. G. de Lima, P. C. Dishuck, R. Durbin, T. Dvorkina, I. T. Fiddes, G. Formenti, R. S. Fulton, A. Fungtammasan, E. Garrison, P. G. S. Grady, T. A. Graves-Lindsay, I. M. Hall, N. F. Hansen, G. A. Hartley, M. Haukness, K. Howe, M. W. Hunkapiller, C. Jain, M. Jain, E. D. Jarvis, P. Kerpedjiev, M. Kirsche, M. Kolmogorov, J. Korlach, M. Kremitzki, H. Li, V. V. Maduro, T. Marschall, A. M. McCartney, J. McDaniel, D. E. Miller, J. C. Mullikin, E. W. Myers, N. D. Olson, B. Paten, P. Peluso, P. A. Pevzner, D. Porubsky, T. Potapova, E. I. Rogaev, J. A. Rosenfeld, S. L. Salzberg, V. A. Schneider, F. J. Sedlazeck, K. Shafin, C. J. Shew, A. Shumate, Y. Sims, A. F. A. Smit, D. C. Soto, I. Sović, J. M. Storer, A. Streets, B. A. Sullivan, F. Thibaud-Nissen, J. Torrance, J. Wagner, B. P. Walenz, A. Wenger, J. M. D. Wood, C. Xiao, S. M. Yan, A. C. Young, S. Zarate, U. Surti, R. C. McCoy, M. Y. Dennis, I. A. Alexandrov, J. L. Gerton, R. J. O'Neill, W. Timp, J. M. Zook, M. C. Schatz, E. E. Eichler, K. H. Miga, A. M. Phillippy, The complete sequence of a human genome. *bioRxiv* (2021), p. 2021.05.26.445798.

9. V. A. Shepelev, L. I. Uralsky, A. A. Alexandrov, Y. B. Yurov, E. I. Rogaev, I. A. Alexandrov, Annotation of suprachromosomal families reveals uncommon types of alpha satellite organization in pericentromeric regions of hg38 human genome assembly. *Genom Data*. **5**, 139–146 (2015).

10. A. Krogh, M. Brown, I. S. Mian, K. Sjölander, D. Haussler, Hidden Markov models in computational biology. Applications to protein modeling. *J. Mol. Biol.* **235**, 1501–1531 (1994).

11. T. J. Wheeler, S. R. Eddy, nhmmer: DNA homology search with profile HMMs. *Bioinformatics*. **29**, 2487–2489 (2013).

12. V. A. Shepelev, A. A. Alexandrov, Y. B. Yurov, I. A. Alexandrov, The evolutionary origin of man can be traced in the layers of defunct ancestral alpha satellites flanking the active centromeres of human chromosomes. *PLoS Genet.* **5**, e1000641 (2009).

13. L. I. Uralsky, V. A. Shepelev, A. A. Alexandrov, Y. B. Yurov, E. I. Rogaev, I. A. Alexandrov, Classification and monomer-by-monomer annotation dataset of suprachromosomal family 1 alpha satellite higher-order repeats in hg38 human genome assembly. *Data Brief*. **24**, 103708 (2019).

14. J. S. Waye, H. F. Willard, Chromosome-specific alpha satellite DNA: nucleotide sequence analysis of the 2.0 kilobasepair repeat from the human X chromosome. *Nucleic Acids Res.* **13**, 2731–2743 (1985).

15. K. H. Miga, Y. Newton, M. Jain, N. Altemose, H. F. Willard, W. J. Kent, Centromere reference models for human chromosomes X and Y satellite arrays. *Genome Res.* **24**, 697–707 (2014).

16. I. Alexandrov, A. Kazakov, I. Tumeneva, V. Shepelev, Y. Yurov, Alpha-satellite DNA of primates: old and new families. *Chromosoma*. **110**, 253–266 (2001).

17. Karen H. Miga and Ivan A. Alexandrov, Variation and evolution of human centromeres: A field guide and perspective. *Annu. Rev. Genet.* (2021).

18. H. Li, Aligning sequence reads, clone sequences and assembly contigs with BWA-MEM. *arXiv [q-bio.GN]* (2013), (available at http://arxiv.org/abs/1303.3997).

19. H. Li, B. Handsaker, A. Wysoker, T. Fennell, J. Ruan, N. Homer, G. Marth, G. Abecasis, R. Durbin, 1000 Genome Project Data Processing Subgroup, The Sequence Alignment/Map format and SAMtools. *Bioinformatics*. **25**, 2078–2079 (2009).

20. A. R. Quinlan, I. M. Hall, BEDTools: a flexible suite of utilities for comparing genomic features. *Bioinformatics*. **26**, 841–842 (2010).

21. S. Levy, G. Sutton, P. C. Ng, L. Feuk, A. L. Halpern, B. P. Walenz, N. Axelrod, J. Huang, E. F. Kirkness, G. Denisov, Y. Lin, J. R. MacDonald, A. W. C. Pang, M. Shago, T. B. Stockwell, A. Tsiamouri, V. Bafna, V. Bansal, S. A. Kravitz, D. A. Busam, K. Y. Beeson, T. C. McIntosh, K. A. Remington, J. F. Abril, J. Gill, J. Borman, Y.-H. Rogers, M. E. Frazier, S. W. Scherer, R. L. Strausberg, J. C. Venter, The diploid genome sequence of an individual human. *PLoS Biol.* **5**, e254 (2007).

22. C. Jain, A. Rhie, H. Zhang, C. Chu, B. P. Walenz, S. Koren, A. M. Phillippy, Weighted minimizer sampling improves long read mapping. *Bioinformatics*. **36**, i111–i118 (2020).

23. T. Dvorkina, A. V. Bzikadze, P. A. Pevzner, The string decomposition problem and its applications to centromere analysis and assembly. *Bioinformatics*. **36**, i93–i101 (2020).

24. R. Patro, G. Duggal, M. I. Love, R. A. Irizarry, C. Kingsford, Salmon provides fast and bias-aware quantification of transcript expression. *Nat. Methods*. **14**, 417–419 (2017).

25. C. Soneson, M. I. Love, M. D. Robinson, Differential analyses for RNA-seq: transcript-level estimates improve gene-level inferences. *F1000Res.* **4**, 1521 (2015).

26. S. Anders, P. T. Pyl, W. Huber, HTSeq--a Python framework to work with high-throughput sequencing data. *Bioinformatics*. **31**, 166–169 (2015).

27. H.-D. Li, GTFtools: a Python package for analyzing various modes of gene models. *bioRxiv* (2018), p. 263517.

28. G. A. Logsdon, M. R. Vollger, P. Hsieh, Y. Mao, M. A. Liskovykh, S. Koren, S. Nurk, L. Mercuri, P. C. Dishuck, A. Rhie, L. G. de Lima, T. Dvorkina, D. Porubsky, W. T. Harvey, A. Mikheenko, A. V. Bzikadze, M. Kremitzki, T. A. Graves-Lindsay, C. Jain, K. Hoekzema, S. C. Murali, K. M. Munson, C. Baker, M. Sorensen, A. M. Lewis, U. Surti, J. L. Gerton, V. Larionov, M. Ventura, K. H. Miga, A. M. Phillippy, E. E. Eichler, The structure, function and evolution of a complete human chromosome 8. *Nature*. **593**, 101–107 (2021).

29. B. L. Ng, N. P. Carter, Factors affecting flow karyotype resolution. *Cytometry A*. **69**, 1028–1036 (2006).

30. S. R. Eddy, Profile hidden Markov models. *Bioinformatics*. **14**, 755–763 (1998).

31. W. J. Kent, C. W. Sugnet, T. S. Furey, K. M. Roskin, T. H. Pringle, A. M. Zahler, D. Haussler, The human genome browser at UCSC. *Genome Res.* **12**, 996–1006 (2002).

32. B. J. Raney, T. R. Dreszer, G. P. Barber, H. Clawson, P. A. Fujita, T. Wang, N. Nguyen, B. Paten, A. S. Zweig, D. Karolchik, W. J. Kent, Track data hubs enable visualization of user-defined genome-wide annotations on the UCSC Genome Browser. *Bioinformatics*. **30**, 1003–1005 (2014).

33. R. C. Edgar, MUSCLE: multiple sequence alignment with high accuracy and high throughput. *Nucleic Acids Res.* **32**, 1792–1797 (2004).

34. S. Chen, Y. Zhou, Y. Chen, J. Gu, fastp: an ultra-fast all-in-one FASTQ preprocessor. *Bioinformatics*. **34**, i884–i890 (2018).

35. M. Kokot, M. Dlugosz, S. Deorowicz, KMC 3: counting and manipulating k-mer statistics. *Bioinformatics*. **33**, 2759–2761 (2017).

36. K. H. Miga, T. Wang, The Need for a Human Pangenome Reference Sequence. *Annu. Rev. Genomics Hum. Genet.* (2021), doi:10.1146/annurev-genom-120120-081921.

37. M. Ventura, F. Antonacci, M. F. Cardone, R. Stanyon, P. D'Addabbo, A. Cellamare, L. J. Sprague, E. E. Eichler, N. Archidiacono, M. Rocchi, Evolutionary formation of new centromeres in macaque. *Science*. **316**, 243–246 (2007).

38. O. Capozzi, S. Purgato, P. D'Addabbo, N. Archidiacono, P. Battaglia, A. Baroncini, A. Capucci, R. Stanyon, G. Della Valle, M. Rocchi, Evolutionary descent of a human chromosome 6 neocentromere: a jump back to 17 million years ago. *Genome Res.* **19**, 778–784 (2009).

39. E. M. Black, S. Giunta, Repetitive Fragile Sites: Centromere Satellite DNA As a Source of Genome Instability in Human Diseases. *Genes* . **9** (2018), doi:10.3390/genes9120615.

40. A. V. Bzikadze, P. A. Pevzner, Automated assembly of centromeres from ultra-long error-prone reads. *Nat. Biotechnol.* **38**, 1309–1316 (2020).

41. Y. Suzuki, E. W. Myers, S. Morishita, Rapid and ongoing evolution of repetitive sequence structures in human centromeres. *Sci Adv*. **6** (2020), doi:10.1126/sciadv.abd9230.

42. H. F. Willard, J. S. Waye, Hierarchical order in chromosome-specific human alpha satellite DNA. *Trends Genet.* **3**, 192–198 (1987).

43. C. Alkan, M. Ventura, N. Archidiacono, M. Rocchi, S. C. Sahinalp, E. E. Eichler, Organization and evolution of primate centromeric DNA from whole-genome shotgun sequence data. *PLoS Comput. Biol.* **3**, 1807–1818 (2007).

44. V. Sevim, A. Bashir, C.-S. Chin, K. H. Miga, Alpha-CENTAURI: assessing novel centromeric repeat sequence variation with long read sequencing. *Bioinformatics*. **32**, 1921–1924 (2016).

45. S. M. McNulty, B. A. Sullivan, Alpha satellite DNA biology: finding function in the recesses of the genome. *Chromosome Res.* **26**, 115–138 (2018).

46. Olga Kunyavskaya, Tatiana Dvorkina, Andrey V. Bzikadze, Ivan Alexandrov, Pavel A. Pevzner, HORmon: automated annotation of human centromeres. *in prep* (2021).

47. T. Lassmann, E. L. L. Sonnhammer, Kalign--an accurate and fast multiple sequence alignment algorithm. *BMC Bioinformatics*. **6**, 298 (2005).

48. F. Madeira, Y. M. Park, J. Lee, N. Buso, T. Gur, N. Madhusoodanan, P. Basutkar, A. R. N. Tivey, S. C.

Potter, R. D. Finn, R. Lopez, The EMBL-EBI search and sequence analysis tools APIs in 2019. *Nucleic Acids Res.* **47**, W636–W641 (2019).

49. R. Ihaka, R. Gentleman, R: A Language for Data Analysis and Graphics. *J. Comput. Graph. Stat.* **5**, 299–314 (1996).

50. C. Ginestet, ggplot2: elegant graphics for data analysis. *JOURNAL-ROYAL STATISTICAL SOCIETY SERIES A*. **174**, 245–245 (2011).

51. K. Tamura, D. Peterson, N. Peterson, G. Stecher, M. Nei, S. Kumar, MEGA5: molecular evolutionary genetics analysis using maximum likelihood, evolutionary distance, and maximum parsimony methods. *Mol. Biol. Evol.* **28**, 2731–2739 (2011).

52. A. Rzhetsky, M. Nei, Theoretical foundation of the minimum-evolution method of phylogenetic inference. *Mol. Biol. Evol.* **10**, 1073–1095 (1993).

53. Professor of Population Genetics Both of the Center for Demographic and Population Genetics Masatoshi Nei, M. Nei, S. Kumar, Evan Pugh Professor of Biology Masatoshi Nei, *Molecular Evolution and Phylogenetics* (Oxford University Press, 2000).

54. N. Saitou, M. Nei, The neighbor-joining method: a new method for reconstructing phylogenetic trees. *Mol. Biol. Evol.* **4**, 406–425 (1987).

55. J. Thakur, S. Henikoff, Unexpected conformational variations of the human centromeric chromatin complex. *Genes Dev.* **32**, 20–25 (2018).

56. M. Martin, Cutadapt removes adapter sequences from high-throughput sequencing reads. *EMBnet.journal*. **17**, 10–12 (2011).

57. O. K. Smith, C. Limouse, K. A. Fryer, N. A. Teran, K. Sundararajan, R. Heald, A. F. Straight, Identification and characterization of centromeric sequences in Xenopus laevis. *Genome Res.* **31**, 958–967 (2021).

58. G. Marçais, C. Kingsford, A fast, lock-free approach for efficient parallel counting of occurrences of k-mers. *Bioinformatics*. **27**, 764–770 (2011).

59. A. Gershman, M. E. G. Sauria, P. W. Hook, S. J. Hoyt, R. Razaghi, S. Koren, N. Altemose, G. V. Caldas, M. R. Vollger, G. A. Logsdon, A. Rhie, E. E. Eichler, M. C. Schatz, R. J. O'Neill, A. M. Phillippy, K. H. Miga, W. Timp, Epigenetic Patterns in a Complete Human Genome. *bioRxiv* (2021), p. 2021.05.26.443420.

60. H. Cheng, G. T. Concepcion, X. Feng, H. Zhang, H. Li, Haplotype-resolved de novo assembly using phased assembly graphs with hifiasm. *Nat. Methods*. **18**, 170–175 (2021).

61. A. Mikheenko, A. V. Bzikadze, A. Gurevich, K. H. Miga, P. A. Pevzner, TandemTools: mapping long reads and assessing/improving assembly quality in extra-long tandem repeats. *Bioinformatics*. **36**, i75–i83 (2020).

62. Sergey Aganezov, Stephanie M. Yan, Daniela C. Soto, Melanie Kirsche, Samantha Zarate, A complete reference genome improves analysis of human genetic variation. *bioRxiv (in review)* (2021).

63. M. Byrska-Bishop, U. S. Evani, X. Zhao, A. O. Basile, H. J. Abel, A. A. Regier, A. Corvelo, W. E. Clarke, R. Musunuri, K. Nagulapalli, S. Fairley, A. Runnels, L. Winterkorn, E. Lowy-Gallego, The Human Genome Structural Variation Consortium, P. Flicek, S. Germer, H. Brand, I. M. Hall, M. E. Talkowski, G. Narzisi, M. C. Zody, High coverage whole genome sequencing of the expanded 1000 Genomes Project cohort

including 602 trios. *bioRxiv* (2021), p. 2021.02.06.430068.

64. R. D. Finn, J. Clements, S. R. Eddy, HMMER web server: interactive sequence similarity searching. *Nucleic Acids Res.* **39**, W29–37 (2011).

65. H. Li, Minimap2: pairwise alignment for nucleotide sequences. *Bioinformatics*. **34**, 3094–3100 (2018).

66. W. Huang, L. Li, J. R. Myers, G. T. Marth, ART: a next-generation sequencing read simulator. *Bioinformatics*. **28**, 593–594 (2012).
